# Combining Machine Learning and Multiplexed, *In Situ* Profiling to Engineer Cell Type and Behavioral Specificity

**DOI:** 10.1101/2025.06.20.660790

**Authors:** Michael J. Leone, Robert van de Weerd, Ashley R. Brown, Myung-Chul Noh, BaDoi N. Phan, Andrew Z. Wang, Kelly A. Corrigan, Deepika Yeramosu, Heather H. Sestili, Cynthia M. Arokiaraj, Bettega C. Lopes, Vijay Kiran Cherupally, Daryl Fields, Sudhagar Babu, Chaitanya Srinivasan, Riya Podder, Lahari Gadey, Daniel Headrick, Ziheng Chen, Michael E. Franusich, Richard Dum, David A. Lewis, Hansruedi Mathys, William R. Stauffer, Rebecca P. Seal, Andreas R. Pfenning

## Abstract

A promising strategy for the precise control of neural circuits is to use *cis*-regulatory enhancers to drive transgene expression in specific cells. However, enhancer discovery faces key challenges: low *in vivo* success rates, species-specific differences in activity, challenges with multiplexing adeno-associated viruses (AAVs), and the lack of spatial detail from single-cell sequencing. In order to accelerate enhancer discovery for the dorsal spinal cord—a region critical for pain and itch processing—we developed an end-to-end platform, ESCargoT (*Engineered Specificity of Cargo Transcription*), combining machine learning (ML)–guided enhancer prioritization, modular AAV assembly, and multiplexed, *in situ* screening. Using cross-species chromatin accessibility data, we trained ML models to predict enhancer activity in oligodendrocytes and in 15 dorsal horn neuronal subtypes. We first demonstrated that an initial enhancer, Excit-1, targeted excitatory dorsal horn neurons and drove reversal of mechanical allodynia in an inflammatory pain model. To enable parallel profiling of a 27-enhancer-AAV library delivered intraspinally in mice, we developed a Spatial Parallel Reporter Assay (SPRA) by integrating a novel Golden-Gate assembly pipeline with multiplexed, *in situ* screening. Regression adjustment for spatial confounding enabled specificity comparisons between enhancers, demonstrating the ability to screen enhancers targeting diverse cell types (oligodendrocytes, motoneurons, dorsal neuron subtypes) in one experiment. We then validated two candidates, targeting Exc-LMO3 and Exc-SKOR2 neurons, respectively. In a companion paper by Noh *et al*, our colleagues show that the functional specificity of the Exc-SKOR2-targeting enhancer, unlike Excit-1, is capable of blocking the sensation of chemical itch in mice. These enhancers were derived from the macaque genome but displayed functional sensitivity in mice. This platform enables spatially resolved, multiplexed *in vivo* enhancer profiling to accelerate discovery of cell-targeting tools and gene therapy development.

## Introduction

Understanding functional circuits within the nervous system requires isolating specific neuron subtypes and interrogating their roles in complex sensory processing and behavior. Recent advances in single-cell genomics have allowed detailed classification of cell types and neuronal subpopulations based on spatial, transcriptomic, and epigenomic profiles^1,2^. These advances have also led to the development of tools that selectively target these subpopulations, namely the use of adeno-associated viruses (AAVs)^3,4^ to deliver transgenic cargo under the control of *cis*-regulatory elements, with the greatest cell type specificity achieved using distal enhancers^5–8^. However, prioritization and screening of enhancer-AAV candidates is highly resource-intensive with limited cross-species compatibility, hampering translational studies and the development of potential cell-targeting therapies. Progress has been especially limited for the spinal cord compared to the brain^9–13^. Spinal motoneuron-targeting enhancers have recently been identified^14–16^, but enhancer discovery in the dorsal horn, a key hub for somatosensory processing^17^, remains unexplored, thus hindering sensory research and the development of next-generation therapies for chronic pain.

Machine learning (ML) sequence-to-function models help prioritize enhancer candidates with greater success rates than using chromatin accessibility or transcription factor motif occurrences alone^6,18^, but prior approaches focused on single species, limiting translation. Multiplexing has the potential to accelerate enhancer screens, with success demonstrated in cell lines^19^ and recently for blood cells^20^, but tissue-specific *in vivo* approaches^8,10,21,22^ remain limited for reasons such as low expression of virally derived transcripts relative to cellular transcripts^8,21,23^, and challenges with cloning and delivery of pooled AAVs^24,25^. Single-cell sequencing-based approaches also lack critical spatial details about injection variability, AAV tropism, and enhancer-specific patterns, and there can be significant uncertainty when labeling rare neuron populations^26^.

To accelerate the discovery of enhancers with spatial- and cell-type-specific activities, we developed an end-to-end framework, ESCargoT (Engineered Specificity of Cargo Transcription), combining machine learning (ML)-guided enhancer prioritization, pooled AAV assembly, and *in situ,* multiplexed spatial profiling. Using single-nucleus chromatin accessibility datasets from macaque and mouse^27^, we trained cross-species ML models to predict enhancer activity across 15 dorsal horn neuron subtypes. An initial candidate enhancer, Excit-1, showed specific expression in excitatory neurons in the superficial lamina of the mouse dorsal horn. When Excit-1 was applied to selectively inactivate those neurons through chemogenetic inhibition^28^, symptoms of mechanical allodynia were abated, a condition where non-painful mechanical stimuli, like light touch, are perceived as painful. To identify additional cell type-specific dorsal horn enhancers to target a greater variety of behaviors, we developed a Spatial Reporter Assay (SPRA) by combining a novel Golden Gate (GG)-based cloning pipeline with multiplexed, *in situ* profiling. We assayed a 27-element AAV library (Xen1) in parallel, each with 3 unique barcodes (81 barcodes total). Spatially aware confounder adjustment revealed enhancer-specific signatures of spatial expression patterns and cell-type specificity. Here, we further validated the cell type specific expression of two enhancers from this assay, Excit-1 and SKOR2.103, for targeting the Exc-LMO3 and the Exc-SKOR2 cell type respectively. In the companion paper (Noh *et al.*), our colleagues utilized these enhancers to investigate these two transcriptomically distinct, yet closely related, cell types, to causally link each subtype to a distinct behavioural output. Overall, the framework we introduce here accelerates spatially resolved enhancer discovery, enabling scientists to probe specific neuron subtypes of neural circuits across species, and to develop new targeted therapeutics for neurological and psychiatric diseases.

## Results

### Macaque single-nucleus open chromatin profiling and conserved subtype harmonization

In order to identify candidate *cis-*regulatory enhancers that will maintain cell-type-specific, species-conserved function as part of an enhancer-AAV, we built upon a validated, primate-centric taxonomy of conserved dorsal horn subtypes based on mouse, macaque, and human snRNA-seq and mouse open chromatin (snATAC-seq)^27^. In this study, we performed single-nucleus open chromatin profiling of the lumbar spinal (L2/3) dorsal horn of two young adult rhesus macaques, and labeled nuclei with the previously described taxonomy, establishing concordance between mouse and macaque open chromatin profiles (**Figure 1a**). Following sequencing and pre-processing, quality control^29^ revealed consistent within-sample and within-animal nucleosome periodicity, cell clustering, fragment counts and TSS enrichments (**Figure S1a-d**). We retained 80,728 high-quality macaque snATAC-seq nuclei, which we clustered (**Figure 1b**), revealing major cell type organization of astrocytes (astrocyte.1 and .2), oligodendrocytes (oligo.1 and .2) and oligo-precursor cells (OPCs), endothelial cells, microglia, stromally derived meningeal cells (stromal), and neurons. Rare spinal cell types including Schwann cells and ependymal cells were not labeled, though the import of obtaining these cell types is lessened by their rarity and naturally lower tropism for enhancer-AAVs^30^. Relative open chromatin accessibility of known marker genes showed the expected cell-type-specific patterns (**Figure 1c**), and 8,533 nuclei (11%) were of neuronal identity (**Figure 1d**), within the typical range of un-biased single-cell profiles of the spinal cord^27,31,32^. Nuclei exhibited consistent signatures of epigenomic quality (TSS enrichment > 10, unique fragments per nucleus >10^3.5^). Next, we re-clustered the neuronal nuclei and co-embedded them with macaque snRNA-seq from the published reference^27^ (**Figure 1f**), enabling label transfer of snRNA-seq nearest neighbors to the snATAC-seq nuclei (**Figure S1e**). We validated conserved neuron subtype identity by first assessing marker gene accessibility (**Figure 1g**), which showed a high degree of specificity of each marker gene for its corresponding neuron subtype, especially for excitatory neurons. For additional validation, we compared macaque cells to those of the published mouse snATAC-seq^27^, first by the proportion of shared open chromatin peaks between subtypes (**Figure S1g**), then by the subtype specificity of transcription factor (TF) motif enrichments using chromVar^33^ (**Figure 1e**). For neuron subtypes that were successfully labeled in both species, we found that TF profiles of a neuron subtype in one species were most correlated with the homologous subtype of the other species, and furthermore recapitulated the structure of closely related subtypes. Specific species-conserved TFs with enriched motifs are also shown in **Figure S1h**. For example, Exc-LMO3 and Exc-SKOR2 were highly correlated based on TF profiles across species, consistent with their transcriptomic similarities^27^, and are specifically enriched for the glucocorticoid receptor (NR3C1; **Figure S1h**), a previous observation in mouse snATAC-seq^27^. The species-conserved cell type-specificity of TF profiles reflects building blocks of the conserved regulatory grammar of these neuron subtypes, which can be learned in greater detail by machine learning models^34–37^.

**Figure 1.**
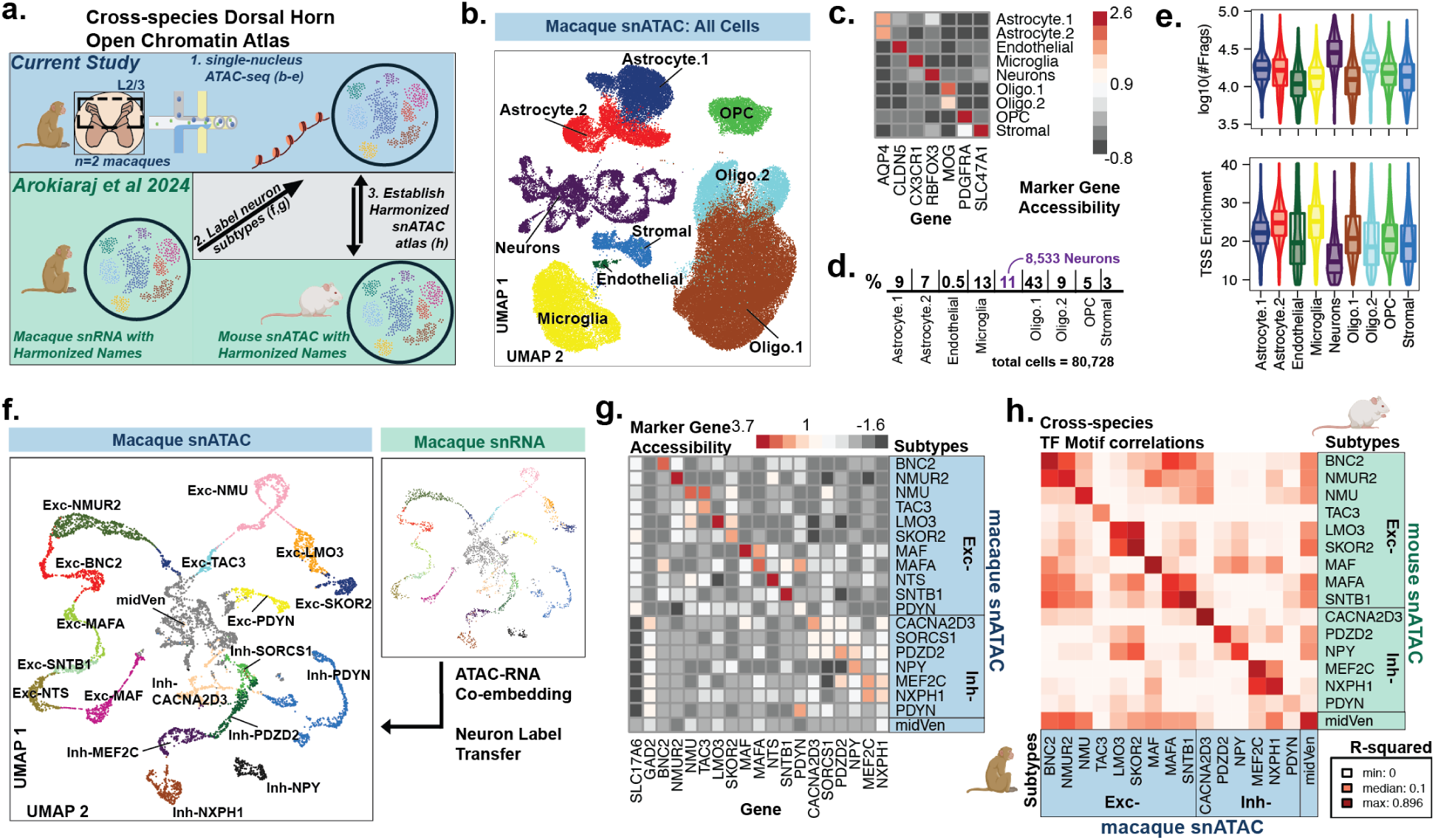
Cross-species Open Chromatin Atlas of the Lumbar Dorsal Horn. a. Overview of strategy to establish cross-species atlas. 1. The current study performed snATAC-seq on the lumbar dorsal horn of n=2 macaques. Neurons were first labeled and then harmonized with homologous mouse dorsal neurons using a previously published resource that provided 2. macaque snRNA-seq and 3. mouse snATAC-seq, respectively. b. Two-dimensional UMAP representation of processed macaque snATAC-seq. Dots represent individual nuclei colored by its major cell identity. c. Average relative accessibility of marker genes of each major cell identity. Gene score is normalized column-wise. d. Nuclei counts per major cell identity. Neurons make up 8,533 (11%) of the dataset. e. Violin and box plot distributions of quality control measures stratified by major cell identity. **Top:** the log10(number of total fragments) per nucleus. **Bottom:** fold-enrichment of transcription start sites (TSS) per nucleus. f. Labeling neuron subtypes with harmonized names. UMAP representations of co-embedding between macaque snATAC neurons (**Left**) and macaque snRNA from *Arokiaraj et al* (**Right**). Names of snRNA neurons are label-transferred to snATAC neurons. g. Average relative accessibility of marker genes of each neuron subtype. Gene score is normalized column-wise. h. Correlations of transcription factor (TF) motif enrichment between macaque snATAC and mouse snATAC (*Arokiaraj et al*) neuron subtypes. Motif enrichment per subtype and species calculated as chromVar deviations, then subtype-specific enrichments compared cross-species (R-squared).

### An excitatory dorsal-specific enhancer that drives chemogenetic reversal of mechanical allodynia in mice

Next, using the above described accessible chromatin data, we sought cell-type-specific dorsal horn enhancers that could be used to decipher the neural circuitry for pain and itch signaling. Due to their established relationship to acute pain, we first sought to target Exc-PDYN and related neuron subtypes^38,39^. An initial enhancer candidate (hereby referred to as Excit-1) was selected based on a combination of its accessible chromatin specificity patterns (**Figure 2a**) and an initial machine-guided priorization method applied to the macaque snATAC-seq, previously described^13^ for single-species enhancer prioritization. Overall, Excit-1 showed high accessibility in macaque neuron subtypes previously identified for their potential roles in mediating acute pain signals, especially Exc-PDYN^38,39^. The machine learning models trained on the new macaque open chromatin were able to reliably distinguish cell type-specific open chromatin (AUROC > 0.90; AUPRC > 0.87).

**Figure 2.**
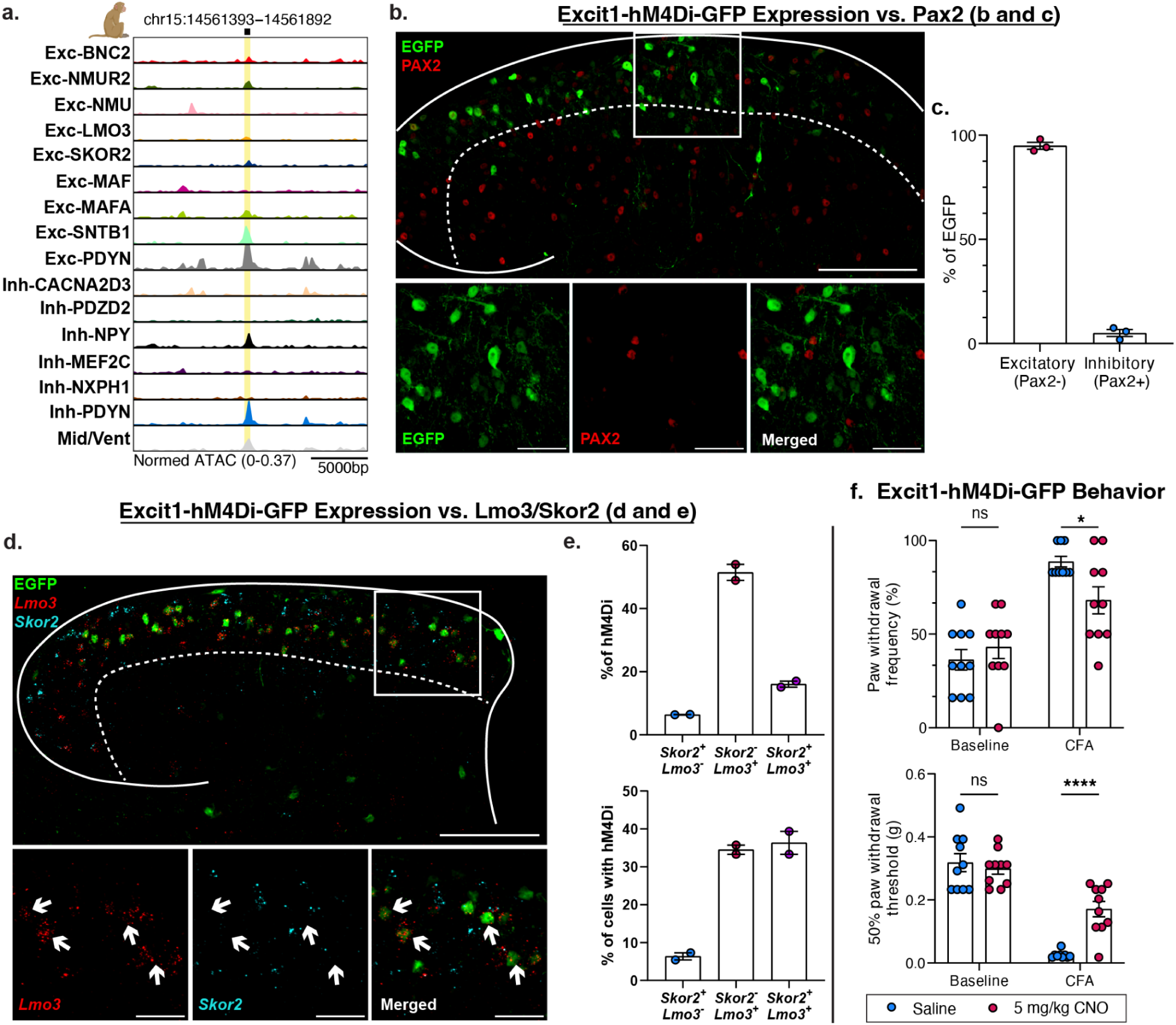
Excit-1, an excitatory, dorsal-horn-specific enhancer that drives chemogenetic reversal of mechanical allodynia. a. snATAC-seq trackplot of 500bp Excit-1 sequence in origin species (macaque). Excit-1 genomic position (x-axis; coordinates shown at top) is indicated by black squares and yellow rectangle highlights, with surrounding 10KB regions shown on each side. Peak height indicates normalized accessibility for a given dorsal horn neuron subtype (y-axis). b. Immunofluorescence assay: transverse lumbar spinal dorsal horn section showing eGFP (green) expressed by the AAV8.Excit1.hM4Di-2a-EGFP, and the inhibitory marker Pax2 (red) in a naïve mouse. White square (top) indicates zoom region (bottom; left, middle, and right). c. Quantification of GFP^+^ and Pax2^+^ immunoreactive cells in the superficial dorsal horn of naïve mice (n=3). Error bars indicate mean and standard error of the mean (s.e.m). d. Transverse lumbar spinal dorsal horn section showing EGFP (green, IF) expressed by the AAV8.Excit1.hM4Di-2a-EGFP, and cellular mRNA of the two marker genes *Lmo3* (red, FISH) and *Skor2* (cyan, FISH) in a naïve mouse. White square (top) indicates zoom region (bottom; left, middle, and right). IF=Immunofluorescence. FISH=Fluorescent in situ hybridization. e. hM4Di quantification (based on EGFP) in n=2 mice, in groups stratified by *Lmo3* and *Skor2* expression, and based on (**top)** percentage of total hM4Di in each group, and (**bottom)** percentage of cells of each group containing hM4Di. f. Paw withdrawal frequency (cotton swab test, top) and 50% paw withdrawal threshold (von Frey test, bottom) with saline or CNO (i.p.) in CFA mice intraspinally injected with PHP.eB.Excit-1-hM4Di-EGFP (n=10).Data are mean ± s.e.m. Statistical analysis was performed with two-way ANOVA with Bonferroni’s post-hoc test. *p < 0.05, ****p < 0.001.

We first explored the excitatory to inhibitory neuron distribution of this enhancer by immunofluorescent staining (IF) of n=3 mice injected intraspinally in the lumbar dorsal horn with AAV8 Excit1-hM4Di-2A-eGFP (**Figure 2b,c**). Comparing the expression of the reporter eGFP with the marker Pax2, which is highly selective for inhibitory neurons in the dorsal horn^31^, we found that 95% (s.e.m. 1.65%) of EGFP was consistently expressed in excitatory neurons (Pax2-). Thus, Excit-1 is highly specific for superficial, excitatory neurons. However, we found much broader expression by Excit-1 than one would expect if it was selective for the rare Exc-PDYN neurons. To assess expression by Excit-1 within particular subtypes, we performed RNAscope in n=3 mice, and measured the overlap of the GFP reporter with the species-conserved marker genes *Lmo3* and *Skor2* (**Figure 2d,e**), two prevalent and closely related neuron subtypes that matched the laminar distribution of Excit-1. Excit1-driven GFP expression overlapped with *Lmo3+* cells moreso than *Skor2+* cells (**Figure 2d**), as confirmed by cell quantification (**Figure 2e**). These results are consistent with what was observed by Noh *et al* (companion paper) using mouse-specific marker gene classifications for these closely related neuron subtypes, Exc-LMO3 (ie. *Grpr+/Tac1+*) and Exc-SKOR2 (ie. *Grpr+/Tac1-*). Noh *et al* also discovered that these distinct subtypes mediate mechanical allodynia and itch, respectively. Based on these experiments, which strongly indicate distinct circuit roles for these neurons, we hypothesized that behaviorally, Excit-1 was likely to influence mechanical allodynia (via Exc-LMO3).

Thus, we evaluated the behavioral effects of chemogenetically silencing neurons transduced by the Excit-1 enhancer in a complete Freund’s adjuvant (CFA) model of inflammatory injury (**Figure 2f**). To test the behavioral specificity of these cell types, we created a plasmid vector in which Excit-1 controlled the expression of hM4Di, a chemogenetic receptor that can inhibit neural activity when activated by the drug clozapine-N-oxide (CNO)^28^. Indeed, we found that CNO administered (5mg/kg, i.p) to mice injected with the PHP.eB Excit-1.hMDi-eGFP in the lumbar dorsal horn significantly attenuated both dynamic and static mechanical allodynia without affecting the baseline mechanical thresholds.

In summary, we generated macaque snATAC-Seq accessibility profiles to train ML models and identified Excit-1, which was predicted to be active in excitatory neurons of the superficial dorsal horn. Immunohistochemical characterization of Excit-1-driven eGFP shows specificity for excitatory neurons, and using *in situ* hybridization, we further show that Excit-1 is specific for *Lmo3*+ over *Skor2*+ neurons. We then used Excit-1 to inhibit the activity of its target cells based on the DREADD^28^ hM4Di, reversing the effects of an inflammatory model of mechanical allodynia. Notably, Noh *et al* (co-submitted) show Excit-1 drives reversal of mechanical allodynia from spared nerve injury and diabetic neuropathy as well, while baseline sensitivity and itch are unaffected. These results showcase the ability of combining genomics with machine learning to develop cell type-specific viruses that can selectively modulate specific behaviors.

### Enhancer Prioritization by Cross-Species, Sequence-to-Function Machine Learning

A higher precision of subtype specificity would enable a more detailed disentangling of the cellular roles of spinal circuits than is currently understood. Thus, we sought to prioritize several candidate open chromatin peaks for further screening, that would be most likely to function as subtype-specific enhancer-AAVs *in vivo* and across species. Previous studies have shown that flexible machine learning (ML) models such as convolutional neural networks (CNNs) can learn complex, species-conserved features of the cis-regulatory grammar^35,40^, and ML models can improve the success rate of enhancer prioritization beyond considering ATAC openness or TF motif identification alone^6,18^. To prioritize enhancers likely to drive cell-type-specific expression across rodents and primates (**Figure 3a**), we built off of the approach Specific Nuclear-Anchored Independent Labeling (SNAIL), described previously for mouse^6^ and macaque^18^. A schematic and description of the cross-species SNAIL procedure are described in **Figure 3a**. Several subtype-specific motifs were identified using MEME-Suite^41^. SVM^42^ and CNN model performance for four representative neuron subtypes are summarized in **Figure 3b**, and the remaining subtypes in **Table S1**. Separate SVMs were trained on macaque and mouse training data to compare species differences, while CNNs were trained on the combined data. Overall, models showed high performance based on auROC and auPRC (y-axes) across off-target cell groups (x-axis), for both mouse and macaque training data (SVMs, **Figure 3b**, top), and across five cross-validation folds (CNNs, **Figure 3b**, bottom). Additional performance metrics are provided in **Table S1**.

**Figure 3.**
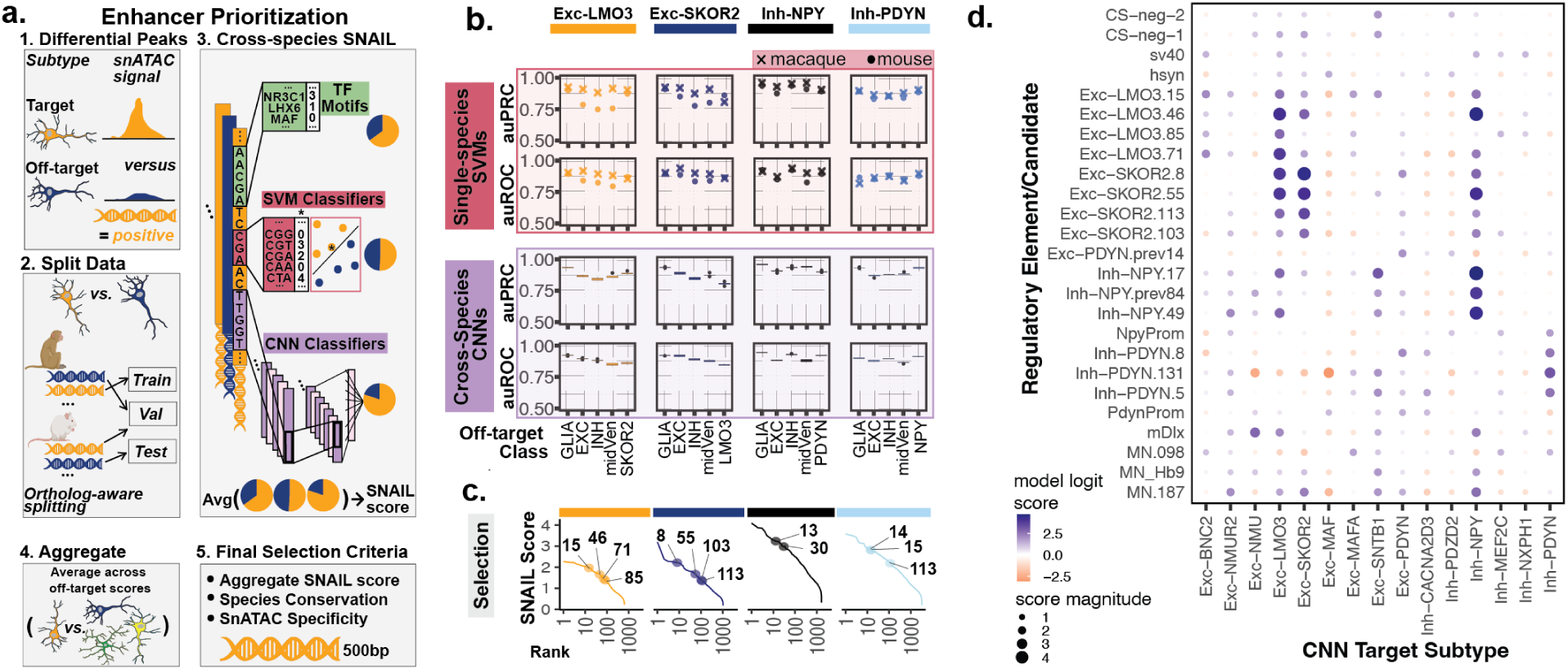
SNAIL Enhancer Prioritization and CNN Evaluation of Candidates/Controls. a. Overview of SNAIL pipeline. (1) Differential peaks are first computed for sets of target (positive) versus off-target (negative) cell groups for use in binary classification. (2) These peaks are assigned for each of five cross-validation folds into Train, Validation (Val), and Test sets, such that cross-species orthologs of peaks are assigned together. (3) SNAIL combines motif enrichment counts from MEME, gapped k-mer support vector machine (SVM) classifier scores, and convolutional neural network (CNN) classifier scores to compute a score for enhancer specificity. (4) For a given target neuron subtype, the SNAIL scores against each off-target class are averaged. (5) Enhancer selection considers the aggregate SNAIL score, conservation across mouse and macaque, and specificity in the snATAC experiments. b. Performance of SNAIL ML models for four representative neuron subtypes. The results for each target subtype is a color-coded column, while the x-axis of each figure is the off-target group for the model: GLIA, excitatory (EXC) inhibitory (INH) neurons, or a closely related subtype (*ie*. Exc-LMO3/SKOR2, or Inh-NPY/PDYN). Y-axis represents either area under the: Precision-Recall Curve (auPRC) or Receiving Operating Characteristic (auROC). **Top:** Performance of SVMs trained on either macaque or mouse peaks, where point shape represents species data. **Bottom**: Box plots of five-fold cross-validation for cross-species CNNs each trained and evaluated on the combination of macaque and mouse peaks, with dots representing outliers. c. Enhancer selection shown for the four neuron subtypes. The y-axis is the aggregate SNAIL score for each peak, one of the criteria for selection, and the x-axis is the ranking of putative enhancers by SNAIL scores. Individual numbers indicate the selected enhancers’ rankings. Consideration of species conservation and snATAC-seq specificity are in addition to SNAIL ranking. d. CNN evaluation of final candidate and control set that comprise the screening panel. The foreground/target subtype of the models shown on the x-axis, and the candidate or control element on the y-axis. The color of a particular point represents the ensemble-averaged logit score for the model set indicated by x, predicted on the candidate or control sequence indicated by y. A more positive score/blue color indicates higher predicted specificity for the target, while a more negative score/orange color indicates lower predicted specificity. Dot size indicates the magnitude of the ensemble logit score.

To uncover motifs most important to the CNN predictions, we used TF-MoDISco^43^, which performs aggregated motif importance scoring while comparing the highest scoring sequences for a given subtype to its negative set of sequences. We matched the resulting motifs to their nearest known transcription factors with Tomtom^44^ (**Table S2**), finding similar patterns as previously described from chromVar (**Figure 1h**), such as again finding the NR3C1 receptor as positively associated with specificity to Exc-LMO3 and Exc-SKOR2 neurons. Certain motifs differentiated between the subtypes as well, such as ZFHX3 (Exc-LMO3 specificity) and POU3F3 (Exc-SKOR2 specificity).

Additionally, we performed cross-species SNAIL to identify oligodendrocyte-targeting enhancers. Given that oligodendrocytes are found across the CNS, we analyzed a collection of published brain snATAC-seq datasets covering four species (human^45,46^, rhesus macaque^13^, rat^47^, and mouse^48–50^). Details about the datasets and labeling of cell types are described in the respective publications. We validated cross-dataset cell type harmonization of oligodendrocytes using chromVar (**Figure S2a**). We trained SNAIL^13^ models to prioritize oligodendrocyte-specific enhancers. Datasets for ML models and their performance are described in **Table S1**.

Finally, we used the dorsal horn cross-species CNNs to retrospectively predict the specific activity of the previously described Excit-1 candidate. We found that model predictions corresponded well with the cross-species snATAC-seq signals and were similar to *in vivo* patterns, with greatest predictions for mostly excitatory neurons overall, Inh-PDYN, Exc-NMUR2 and Exc-PDYN among the highest predicted subtypes, with positive prediction of Exc-LMO3 activity, as well. Inhibitory subtypes such as Inh-CACNA2D3 and Inh-NXPH1 were among the lowest predicted (**Figure S1j**).

### Design of Regulatory Element Library and Control Panel for Screening

The ML algorithms were able to prioritize dozens of candidate cell type-specific enhancers across our fifteen neuron subtypes. Testing these candidates individually, as is commonly done^9,12,13^, would require considerable time, animals, personnel, and resources. Thus, we sought to create a library of top candidates that we could screen in parallel.

We constructed a library of regulatory element candidates and controls as follows. We tested two random negative controls: we generated random 500-base-pair (bp) elements that can be inserted in place of a true regulatory element, with required GC content within 40-60%, and ranked for low predicted activity in neurons using previously trained cross-species sequence-to-function models^40^. As positive controls, we selected SV40^51^ and hSyn^52^, with hSyn expected to show greater pan-neuronal specificity than SV40. We found that there was a dearth of already identified and characterized spinal-targeting regulatory elements in the literature, especially for the dorsal horn. Thus, while truncating or extending to maintain chromatin-accessible 500bp regions (see Methods), we selected the following: (1) the mDlx promoter^53^, expected to show specificity for inhibitory neurons, (2) the NPY^54^ and PDYN^17^ promoters, because of their expected preference for inhibitory neurons and Exc/Inh-PDYN neurons respectively, and (3) MN.Hb9^15^ and MN.098^16^, each validated for motoneuron-specific activity, with the strongest evidence provided for MN.098. Finally, from the same study we treated MN.187 as a naturally occuring negative control sequence, originally intended to target motoneurons but with evidence that it does not significantly express *in vivo*^16^.

For enhancer candidates to target dorsal horn neuron-subtypes, we used SNAIL rankings (**Figure 3c**) to select four enhancer candidates for the neuron subtypes Exc-LMO3 and Exc-SKOR2, two for Inh-NPY, and three for Inh-PDYN. One additional Inh-NPY candidate and one Exc-PDYN candidate, ranked in a pilot iteration of SNAIL, were also added. In total, fifteen candidates were prioritized to target five dorsal neuron subtypes. We also included two SNAIL-ranked candidates targeting oligodendrocytes, human.oligo.41 from the human genome and mouse.oligo.18 from the mouse genome, and both were found to have oligodendrocyte-specific accessibility in the mouse dorsal horn snATAC-seq (mouse ortholog of human.oligo.41; **Figure S2b-e**).

ML-based, subtype-specific activity predictions are shown for the full collection of 27 enhancer candidates and controls in **Figure 3d**, with oligodendrocyte-targeting candidate predictions given in **Table S1**. Overall, highest model predictions were of the dorsal horn enhancer candidates for their targets, as expected from ranking, although model predictions demonstrate residual predicted off-target in some cases. The mDlx, NPY and PDYN promoters were noted to have non-specific predictions, consistent with the principle that promoters are generally less cell-type-specific than enhancers, but with the caveat that models trained on distal enhancers may not accurately interpret the regulatory grammar of promoters^40^. For similar neuron subtypes, including LMO3 and SKOR2, even the top scoring candidate enhancers had predicted off-target effects (**Figure 3d**), a potential limitation of finding maximally specific natural enhancer sequences.

### Modular assembly of pAAV library vectors for high-throughput profiling of non-coding regulatory regions

To facilitate high-throughput analysis of the regulatory activity of non-coding genomic regions in neuron-specific cell types, we developed the modular platform SPRA for the rapid assembly of pAAV vectors. We strategically engineered SPRA pipeline to integrate three core technologies underpinning the ESCargoT project: (1) our systemically delivered Massively Parallel Reporter Assay (MPRA) for barcoded, *in vivo* assessment of candidate cis-regulatory elements^22^; (2) high-resolution spatial transcriptomics for cell-type-specific expression profiling within intact tissues^55^; and (3) Fluorogen-activating protein (FAP) biosensor technology^56–59^ with high sensitivity required to detect reporter expression in transduced cells.To get the SPRA pipeline operational, we first re-designed our sysMPRA pAAV vector^22^ to create the final SPRA vector. This redesigned vector was adapted for Golden Gate (GG) assembly and divided into four modules, containing the essential elements required for a high-throughput library vector pipeline (**Figure 4a** and **S3**). Next, we engineered SPRA into two sequential steps (**Figure 4** and **S4**): Step 1, pAAV_A vectors: integration of CREs (enhancers) and the barcoded library (**Figure 4b**), followed by Step 2, pAAV_B vectors: the incorporation of the fluorescent reporter (**Figure 4c**).

**Figure 4.**
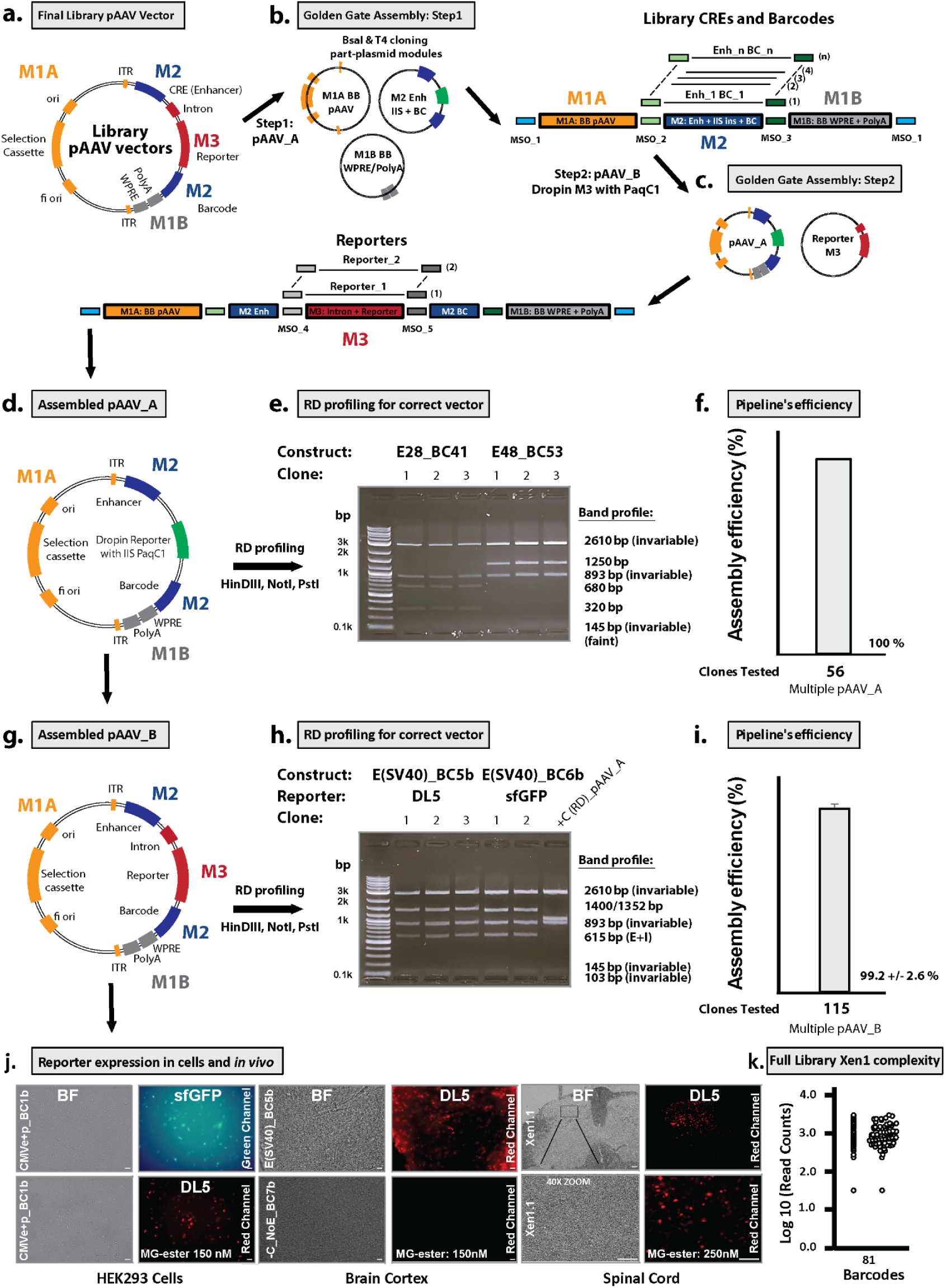
Design and Validation of the SPRA Golden Gate (GG) assembly pipeline for studying non-coding genomic regions (enhancers) *in vivo*. The SPRA cloning pipeline for pAAV library vectors is a two-step, high-throughput process designed for modular incorporation of CREs and barcode libraries. It uses part-plasmid modules and offers flexibility to integrate any fluorescent reporter of choice into the platform. An extensive technical overview of the cloning details of the SPRA pipeline is presented in figure S4. (a) Cartoon representation of final pAAV_B library vectors comprising four modules: M1A (orange), M1B (gray), M2 (blue) and M3 (red). (b) Step 1: GG assembly of pAAV_A Vector using three part-plasmid modules: M1A (backbone (BB), orange), M1B (elements WPRE and PolyA, gray), and M2 (CRE (enhancer), IIS enzyme drop-in sites and barcode, blue). Enzyme BsaI generates Module Specific Overhangs (MSOs, squares in different colors) that guide directional ligation via T4 ligase. Each module contains two unique MSOs that dictate the assembly direction: MSO_2 and MSO_3 (light and dark green) align M1A, M2, and M1B, while MSO_1 (blue) ensures proper circularisation of the vector (see details Figure S4). (c) Step 2: GG assembly of the final pAAV_B vector by inserting reporter module M3 (red) into pAAV_A utilizing enzyme PaqC1 mediated GG cloning. PaqC1 generates MSO_4 and MSO_5 (light and dark gray squares) to enable directional assembly and correct integration of the reporter module M3 into pAAV_A to create the final pAAV_B vector. (d) Cartoon representation of the assembled pAAV_A vector of step 1. (e) Restriction digest (RD) analysis utilizing *HinDIII*, *NotI*, and *PstI*-digested pAAV_A vectors of multiple clones. The pAAV_A vectors show consistent profiles with invariant and variable bands matching the expected band patterns; representative constructs E28BC41 and E48BC53 are shown.(f) SPRA pipeline assembly of pAAV_A vectors achieved 100% efficiency across 56 constructs in several independent GG reactions (see details Table S4).(g) Cartoon representation of the assembled pAAV_B vector of step 2. (h) As in (E), RD analysis of final pAAV_B vectors shows expected cutting patterns, with variable bands corresponding to unique enhancers, barcodes and reporters. Representative constructs E(SV40)-DL5-BC5b and E(SV40)-sfGFP-BC6b are shown. (i) As in (f) SPRA pipeline assembly of pAAV_B vectors achieved 99.2 +/- 2.6 % efficiency across 115 constructs in several independent reactions, demonstrating robust pipeline performance (see details Table S5). (j) Keyence fluorescent images of reporter expression: HEK293 cells transduced pAAV_B vectors with CMVe+p-BC1b driven by sfGFP (left, top) and DL5 (Left, bottom) show significant reporter expression. *In vivo*, E(SV40)-BC5b driven DL5 is expressed in the mouse cortex (middle, top), while no signal is detected with control construct no-enhancer (middle, bottom). Robust Xen1.1 library driven DL5 expression is detected in the spinal cord (right, top), with 40x zoom image (right, bottom) demonstrating reliable performance of the SPRA cloning pipeline in libraries assembly.(k) NGS barcode read counts for the final Xen1 pAAV_B library showing representation of all 81 library member combinations (27 enhancers times 3 unique barcodes). FAP/fluorogen fluorescence was achieved with 150-250 nM MG-ester and measured at E_excitation_ = 636nm/ E_emission_ = 664nm. Scale bars 30µM. Abbreviations: WPRE: Woodchuck Post-transcriptional Regulatory Element; M: module; E: Enhancer; P: Promoter; BC: Barcode; BB: backbone, RD restriction digestion; ITR: Inverted Terminal Repeats; FAP: Fluorogen Activating Protein; GG: Golden Gate; CMV: Cyto-Megalo-Virus; SV40; Simian Virus 40.

In Step 1, the vector system is designed using three interchangeable modules: M1A (orange), M1B (gray), which contain the core elements required for the pAAV vector, and M2 (blue) which includes the enhancer-barcoded library (CREs and barcodes) and a drop-in IIS PaqCI site (**Figure 4b**) that enables the insertion of the fluorescent reporter M3 (red), in Step 2 (**Figure 4c**). For full design part-plasmid modules see **Figure S5**. For the GG assembly process, we employed, empirically verified high-fidelity module-specific overhangs (MSOs) from prior work^56^, ensuring efficient and reliable assembly of the pAAV vectors (**Figure 4, S4** and **Table S3**). The pAAV_A vector is assembled via a three-component GG reaction using the IIS restriction enzyme (RE) BsaI and T4 ligase. In short, the part-plasmid modules are cleaved to generate the MSOs and these overhangs enable directional ligation by T4 ligase (**Figure 4b**, squares) and subsequently circularization (MSOs, blue) to produce the first pAAV_A vector (**Figure 4d**; detailed design/assembly workflow **Figure S4a–d**). Similarly, the final pAAV_B vector is assembled via a two-component GG reaction with PaqCI and T4 ligase (**Figure 4c**). Vector pAAV_A is combined with Module M3 to directionally ligate the fluorescent reporter on the MSOs, producing the final pAAV_B vector (**Figure 4c**; Detailed design/assembly workflow **Figures S4e–g**).

First, we constructed part-plasmid modules M1A, M1B, M3 reporters (sfGFP and DL5) and two test M2 enhancer-barcode modules using entry vector pYTK001^60^ that would allow us to test the SPRA cloning pipeline. Indeed, the insert sizes of the part-plasmid modules matched the expected band patterns (**Figure S6**), and sequence analysis confirmed these part-plasmid modules. We then employed Step 1 of the cloning pipeline to assemble pAAV_A vectors and verified correct assembly by utilizing restriction digestion (RD) profiling, a rapid method for early assessment for correct vector assembly, as previously described^56^.We confirmed the first two vectors CMVe+p_BC1b and CMVe_BC2b, as their RD profiles matched the expected band patterns (**Figure S7a**). Subsequently, we generated multiple pAAV_A plasmids to rigorously evaluate the SPRA assembly pipeline, including positive (SV40 enhancer) and negative (no enhancer) controls (**Figure S7b,c**) as well as several single library elements of the spinal cord (**Figure 4e**). RD profiling analysis of all tested plasmids showed expected band patterns, indicating correct vector assembly (**Figure S7** and **4e**). Since the robustness of the SPRA cloning pipeline depends on highly efficient library-scale assembly, we assessed assembly efficiency by testing 17 constructs and randomly screening 56 clones, achieving 100% correct assembly in the initial cloning step (**Figure 4f, Table S4**). Notably, GG assembly relies on type IIS restriction enzymes, enabling high fidelity, sequence-specific overhang ligation while avoiding PCR-induced errors, effectively eliminating the need for resequencing verified modules^56,61,62^. To confirm the accuracy of the SPRA initial cloning step, we performed whole-plasmid sequencing of multiple pAAV_A vectors and found a match between the expected structure, including the 4 bp overhangs and sequenced plasmid.

We then performed the final step of the SPRA pipeline by assembling pAAV_B vectors through the integration of fluorescent reporter modules M3 (sfGFP and DL5) into pAAV_A plasmids using GG assembly with type IIS enzyme PaqCI (**Figure S8**). RD profiling of representative constructs, including E(CMVe)-DL5-BC2b, E(CMVe+p)-DL5-BC1b, E(SV40)-DL5-BC5b, and E(SV40)-sfGFP-BC6b, revealed band patterns matching expected sizes (**Figure 4H, S8a,b**). Consistently, NoE-DL5-BC7b and NoE-sfGFP-BC8b displayed the predicted band profiles (**Figure S8b**). As in Step 1, we assembled 54 constructs and randomly screened 115 clones, achieving an assembly efficiency of 99.2% ± 2.6% (**Figure 4i, Table S5**). Again, whole-plasmid sequencing confirmed correct overhang ligation and thus, validating the accuracy, efficiency and reliability of SPRA’s final cloning step.

Subsequently, we validated reporter expression from SPRA-assembled constructs by transducing HEK293 cells with positive control vectors containing the CMV enhancer/promoter (CMVe+p) driving either sfGFP or DL5. Fluorescence microscopy confirmed robust expression of both reporters, indicating efficient transcriptional activity and proper vector function (**Figure 4j**, left). *In vivo*, DL5 expression was detected in the mouse cortex following delivery of an SV40-driven enhancer construct, while no signal was observed with a no-enhancer control, confirming enhancer-dependent expression (**Figure 4j**, middle). These results demonstrate that SPRA enables consistent reporter expression *in vitro* and *in vivo*, validating the platform’s utility and functional integrity.

To test the library-scale performance of the SPRA pipeline, we first constructed the Xen1.1 spinal cord library, comprising 27 enhancers each with a single barcode. As an initial validation, we also assembled a smaller subset (Xen1-small, 9 elements), generating both pAAV_A (**Figure S9a**) and pAAV_B (**Figure S9b**) vectors. RD profiling revealed the expected pattern: consistent invariant bands alongside variable fragments reflecting enhancer and barcode diversity, including elements lacking restriction sites, confirming successful library assembly. Next-generation sequencing verified full representation of all Xen1.1 elements (**Figure S10**). Following intraspinal delivery of PHP.eB carrying the Xen1.1 library, we observed widespread DL5 reporter expression throughout the spinal cord (**Figure 4j**, right and **S9c**), indicating functional activity of the CREs *in* vivo. These findings demonstrate that the SPRA pipeline enables efficient, high-fidelity construction and *in vivo* delivery of complex AAV libraries, at least for Xen1.1 library.

Finally, we assembled the full Xen1 library (27 enhancers, each with three barcodes) for spatial transcriptomics. As with Xen1.1, we validated pAAV_A and pAAV_B constructs via RD profiling, which showed the expected banding patterns, confirming correct assembly (**Figure S11a,b**). To assess library complexity, we RD-profiled individual clones randomly selected from LB plates. This simple yet effective approach provided an early indication of SPRA’s cloning performance, revealing structural diversity and integrity within the library. As such, we first calculated the ratio of the expected RD cutting profile of the 81 element Xen1 library (**Table S9**). The RD profile predicted distribution of the Xen1 library consists of 43 uncut elements (53.1%, orange) and 38 cut variants (46.9%, green), containing one or more HinDIII or PstI sites within CREs or barcodes (**Figure S12a, Table S10**). Next, we analyzed two Xen1 libraries; Xen1-DL5 (9 clones) yielded 5 distinct cut profiles and 4 uncut; Xen1-sfGFP (10 clones) showed 4 cut and 6 uncut profiles (**Figure S12b**). The combined profile ratio (10:9, uncut:cut) matches, on average, the expected 53:47 percentile distribution, including the 9 distinct cutting profiles (cut, green), providing a first indication that the SPRA cloning pipeline likely preserves library diversity and structural integrity. To minimize potential batch effects and enhance library complexity, we pooled two independent pAAV_B assemblies across 10 cloning batches (see Methods). NGS analysis confirmed robust representation of all 81 constructs (27 CREs x 3 barcodes) in the Xen1 library, except for a single underrepresented barcode, likely due to an GG overhang mutation during G-block synthesis or pipetting error upon library assembly (**Figure 4k**). Overall, our results establish SPRA as a robust and efficient platform for highly-fidelity library construction enabling effective *in vivo* delivery of complex AAV libraries.

### Multiplexed, *In Situ* Viral Expression of Single Cells

We sought to measure the expression of the regulatory elements of the above-described library in a multiplexed, *in situ* assay. We designed a custom Xenium panel with probes measuring endogenous cell-type-specific RNAs^27^ probes to detect the synthetic, 500 bp DNA barcodes delivered by the AAVs. We pre-screened the custom barcode panel to determine potential off-target detection of the synthetic barcodes by the probes. First, we imaged and analyzed two naive mice, without injection of AAV, using Xenium. Our results show that five of the 81 barcodes had spuriously high expression in the virus-free mice (**Figure S13a**), and as such were excluded from the subsequent study of the Xen1 library in Xenium.

To assess the *in vivo* expression of the pooled Xen1 viral library (**Figure 5a**), we intraspinally injected PHP.eB-Xen1-DL5 in C57BL/6J mice (n=3) at 1.62 x 10^10^ and 2.7 x 10^10^ vector genomes (vg) total. After 4 weeks, the L3-L5 lumbar segment of the spinal cords were dissected and imaged with Xenium using the custom panel. We performed iterative cell labeling as described previously in ^27^ for each mouse, with one update. We noted in our mouse Xenium results (**Figure S13e**), and retrospectively found in macaque snRNA-seq (**Figure 1e of** ^27^), that Exc-SKOR2 neurons tend to split into two separable sub-clusters, which we re-named Exc-SKOR2.1 and Exc-SKOR2.2, the latter corresponding to the cells most similar to Exc-LMO3 (**Figure S13e**; Exc-SKOR2.1 cells in light grey, Exc-SKOR2.2 in dark blue). Overall, cells with a label transfer confidence score > 0.5 for major cell types and/or neuron subtypes were kept (**Figures S13c,f**). Three small clusters of neurons were excluded based on unclear identity after integration across the three mice. After these quality control steps, there were 362,006 total cells, including 63,610 (18%) neurons among the three mice (**Figures S13b,e**), and confirmed with individual marker gene specificity (**Figure S13d,g**). Example slices (**Figure 5b**) are shown with labeled major cell types (**Top**) and neuron subtypes (**Bottom**). Distributions of laminar depth (**Figure 5c**) and cell area (**Figure 5d**) were also compared across neuron subtypes, with spatial depth patterns reproduced as in ^27^.

**Figure 5.**
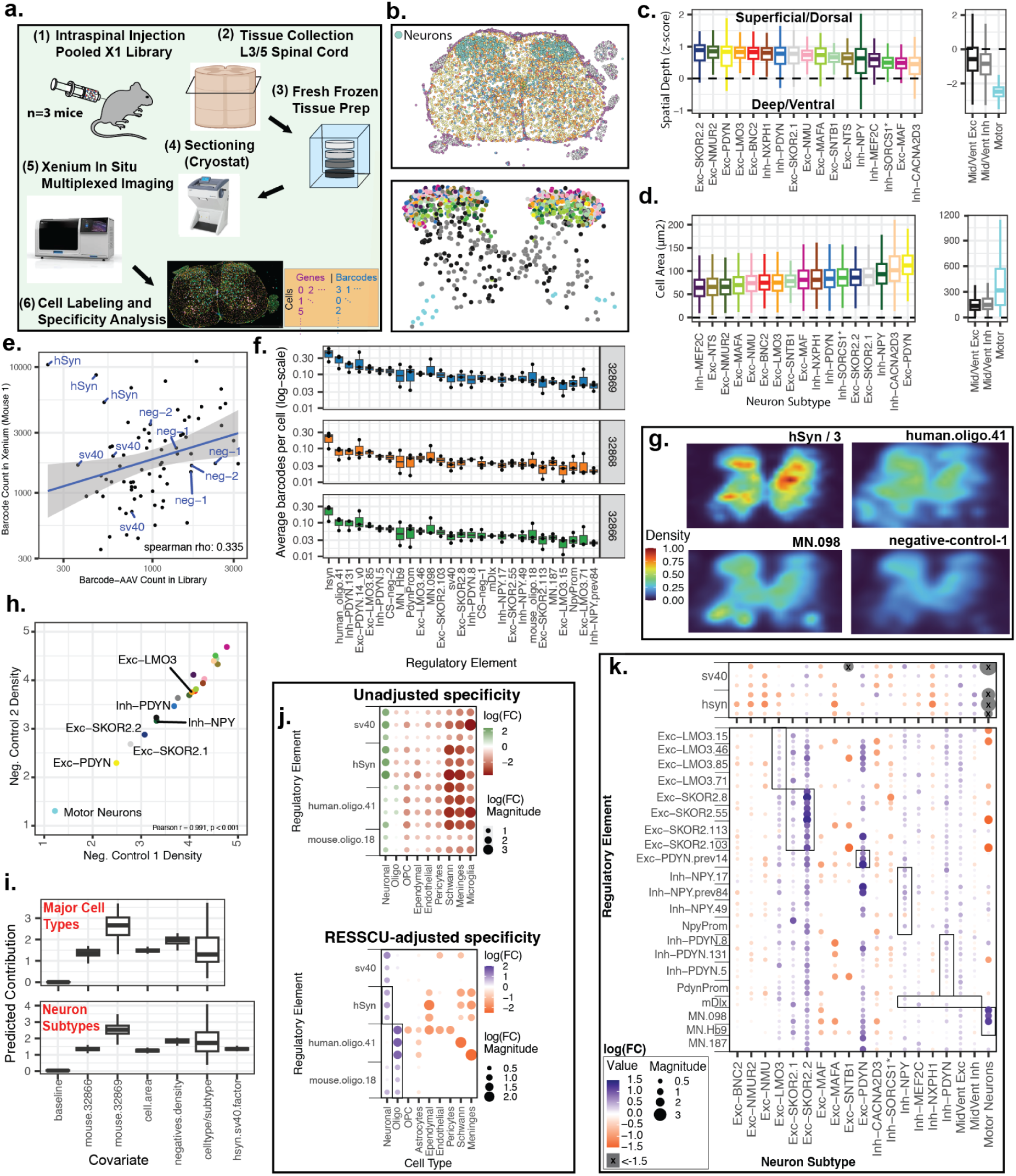
Single-Cell Analysis of Multiplexed, *In Situ* Viral Expression in Mouse Spinal Cord. a. Schematic of experiment and analysis procedure. b. Representative Xenium image of spinal slice with segmented, labeled major cell types (Top) and neuron subtypes (Bottom). Neuron subtype, same color scheme as in c and d. c. Box plot distributions of spatial depths stratified by neuron subtype, normalized as z-score of vertical position. Positive (negative) z-score represents more superficial (deep). Separate y-axis scale shown for Mid/Ventral neurons and motor neurons. d. Box plot distributions of cell area stratified by neuron subtype. Separate y-axis scale shown for Mid/Ventral neurons and motor neurons. e. Scatterplot of Barcode-AAVs for representative mouse where each dot is an individual barcode: library representation of barcode based on sequencing (x-axis) versus total detected barcode count in Xenium (y-axis). hSyn, SV40, and negative controls (neg-1/2) highlighted. Spearman correlation shown in bottom right. f. Boxplot distributions, stratified by each of the n=3 mice, of each regulatory element (x-axis; data points for the box plot are each of the 2-3 barcodes for that element) and its per-cell average detected count (y-axis; log scale). g. Example images of spatial density distributions of detected barcodes from each of four regulatory elements (hSyn, human.oligo.41, MN.098, negative-control-1). One third of detected hSyn signal shown due to strong expression and to compare across images with the same density scale. h. Scatter plot of spatial density of negative-controls-1 and 2 (x and y axes respectively) stratified by neuron subtypes (each individual point), showing regulatory element-independent bias in spatial spread of barcode signals. Pearson r correlation in bottom right. i. Coefficients from regression models for major cell types (Top) and neuron subtypes (bottom). Covariates shown on x-axis. Predicted contribution to barcode counts from each covariate, *ie*. exp(model coefficient), shown on y-axis. Boxplots aggregated across barcode count prediction models for each covariate. j. Specificity fold change (FC) results for regulatory elements (y-axis) targeting each major cell type (x-axis); unadjusted raw log(FC) of counts (Top) and log(FC) estimated from RESCU coefficients (Bottom). Top and Bottom: size of circles indicate magnitude of Log(FC). Top: green (red) circles indicate higher (lower) unadjusted log(FC) specificity for each cell type. Bottom: blue (orange) circles indicate higher (lower) regression-estimate log(FC). Boxes highlight expected target of regulatory element. k. Specificity fold change (FC) results for regulatory elements (y-axis) targeting each neuron subtype (x-axis); Size of dots indicate magnitude of Log(FC). Grey circles with x indicate very low specificity of log(FC) < −1.5. Blue (orange) dot indicates higher (lower) RESCU-estimated log(FC) of subtype specificity. Boxes highlight expected target of regulatory element.

Overall detection levels of each regulatory element were compared to their proportion of representation in the Xen1 library (**Figure 5e**), with hSyn showing consistently high detection despite relatively low library representation, which is expected based on its function as a strong neuronal promoter. Variability of the regulatory elements across their barcodes and three mice (**Figure 5f**) showed overall trends of varying degrees of barcode variability, and although mice had different absolute levels of viral load, the relative detection of barcodes was highly consistent across mice (correlations between mouse #32869 versus other two mice, of per-barcode expression, Pearson r = 0.988 and 0.994). As in the four examples of spatial spread of regulatory elements (**Figure 5g**), it was observed that regulatory elements tended to have a strong shared component of spatial extent, likely reflecting perfusion biases of intraspinal injection, and/or tropism bias of the PhP.eB^63,64^ AAV, and sources of noise such as from off-target detection of single-stranded DNA^65^ or from enhancer crosstalk, a known phenomenon whereby the presence of multiple AAV viral genomes in a cell can result in an enhancer from one AAV genome driving expression of the cargo of another^25^. However, unique characteristics of the regulatory elements were also observable, such as the more ventral expression of MN.098^16^, which one would expect given that motor neurons are the most ventrally positioned neurons of the spinal cord (**Figure 5g**, **bottom of MN.098**). Similarly, as expected, we found greater expression of human.oligo.41 extending beyond the grey matter to where oligodendrocytes are higher density (for reference, see **Figure 5b Top**, yellow cells = oligodendrocytes). To better isolate the shared spatial component, we compared the spatial densities of the two random negative control sequences, finding nearly perfect correlation at the neuron subtype level (pearson r = 0.991, **Figure 5h**), demonstrating a consistent spatial pattern of detection that is independent of any particular regulatory element.

### Cell-type-specific Characteristics of Regulatory Elements

To address confounding factors in our *In Situ* assay, we used a multiple regression approach, Regulatory Element Specificity after Spatial Confounder Untanglement (RESSCU), to estimate the expression of enhancer-AAV barcodes at the single-cell level that are attributable to the major cell type or subtype identity of the cell. For regulatory elements targeting either neurons or oligodendrocytes, we predicted barcode expression of a cell based on (1) the mouse from which the cell is derived, to account for global viral load differences (**Figure 5f**), (2) the previously described spatial density average (**Figure 5h**) of the two random negative controls, (3) the cross-sectional area of the cell which influences exposure to AAV and uptake (**Figure 5d**), and finally, (4) the cell type of the cell, (Neuron, Oligodendrocyte, *etc*). For the neuron subtype-targeting candidates, analysis was the same as above, except the subtype identity was considered (Inh-NPY, Exc-LMO3, *etc*.); and, an additional covariate was added to adjust for potential enhancer crosstalk, based on the combined barcode counts of hSyn and SV40, given that these strong promoters could drive significant expression of other barcodes.

For each coefficient of the fitted model, exp(coefficient) represents the contribution to counts from the covariate (**Figure 5i**). Coefficients for cell/subtype identity showed the highest variability across regulatory elements, suggesting cell-type-specific properties of regulatory elements are isolated by the model.

The raw, unadjusted log(fold-change) for each cell type/subtype is defined as the average log(barcode count) in the target cell/subtype relative to their counts in the off-target cells, weighted by their cell counts, and is shown in **Figure 5j-Top** (major cell types) and **Figure S1h** (neuron subtypes). The RESSCU-adjusted log(fold-change) is defined the same but instead of log(counts), the model coefficient for the target and off-target cell/subtypes are used, shown in **Figure 5j**-Bottom (cell types) and **Figure 5k** (neuron subtypes).

The positive controls SV40 and hSyn, both unadjusted and RESSCU-adjusted, showed a preferential enrichment for neurons (**Figure 5j-Top versus Bottom**) as expected based on promoter and AAV tropism^66,67^. After RESSCU adjustment (**Figure 5j, Bottom**), hSyn showed greater neuron specificity than SV40. The two oligodendrocyte-targeting candidates showed neuronal preference in unadjusted counts; after applying RESSCU (**Figure 5j-Top versus Bottom**), both showed greatest specificity for oligodendrocytes across barcodes, with human.oligo.41 showing greater specificity than mouse.oligo.18. The remaining enhancer candidates were compared for their relative specificities for particular neuron subtypes (**Figure 5k**), together with SV40 and hSyn controls. hSyn was highly non-specific for motoneurons^14^. Overall, the inhibitory candidates targeting Inh-NPY, Inh-PDYN were non-specific, and Exc-LMO3 also showed weak specificity. There were also remaining biases in expression even after RESSCU adjustment, with broadly high expression in Exc-SKOR2 clusters and Exc-PDYN and broadly low expression in Exc-SNTB1 and Exc-MAFA. Several candidates showed *in vivo* specificity for their targets: multiple candidates were on-target-specific for the Exc-SKOR2 cells (Exc-SKOR2.1 and Exc-SKOR2.2) especially for the more numerous Exc-SKOR2.2 subtype, which had higher representation in the original snATAC-seq training data. Exc-PDYN.prev14 also showed preference for Exc-PDYN. MN.098 showed the most consistent and specific expression across all three of its barcodes, whereas Hb_9 and MN.187 had relatively lower motoneuron specificity, as expected. Spatial specificity estimation before and RESSCU adjustment are visualized in **Figure S13i** for the example candidates human.oligo.41, Exc-SKOR2.103, and Exc-LMO3.71.

### Validation of Individual Enhancer Candidates

To test if inhibitory-targeting candidates are non-specific outside of the Xenium multiplexing, we selected two candidates, Inh-NPY.17, and Inh-PDYN.8, for individual validation. We found based on *in situ* hybridization (RNAscope) that neither candidate (**Figure S15**) was specific for *Npy*+ or *Pdyn*+ neurons, respectively, demonstrating concordance with the non-specificity observed in Xenium.

Two other candidates, Exc-LMO3.71 and Exc-SKOR2.103, were selected for one-at-a-time validation as potentially specific enhancers, based on the combination of ML model predictions, species conservation, and screening results with Xenium. snATAC-seq profiles are shown for each (**Figure 6a,b**) for macaque and mouse orthologs. Exc-SKOR2.103 showed greater conservation than Exc-LMO3.71 in snATAC-seq profiles, whereas both candidates were predicted to be cross-species, cell-type-specific based on the ML models (**Figure 3d**). Each candidate was packaged in PhP.eB with a GFP reporter and delivered intraspinally to 3 CL57BL6/J mice. *In situ* hybridization (RNAscope) and immunofluorescence (IF) was performed, using a primary antibody to GFP and probes for the species-conserved markers *Lmo3* and *Skor2*. As seen in **Figure 6c,d**, GFP expressed by both candidates was confined to the superficial layers of the dorsal horn, where Exc-LMO3 and Exc-SKOR2 neurons predominate, especially Exc-LMO3.71. Cell quantification (**Figure 6e,f**) revealed that both candidates were most active in *Skor2*+ and *Lmo3*+ neurons. As expected based on the SNAIL model predictions and Xenium screening, Exc-SKOR2.103 driven expression of GFP had more *Lmo3*-/ *Skor2*+ cells compared to Exc-LMO3.71 (**Figure 6e**). The specificity of Exc-LMO3.71 for its target neurons exceeded that estimated by Xenium screening and by the snATAC-seq signal of its mouse ortholog.

**Figure 6.**
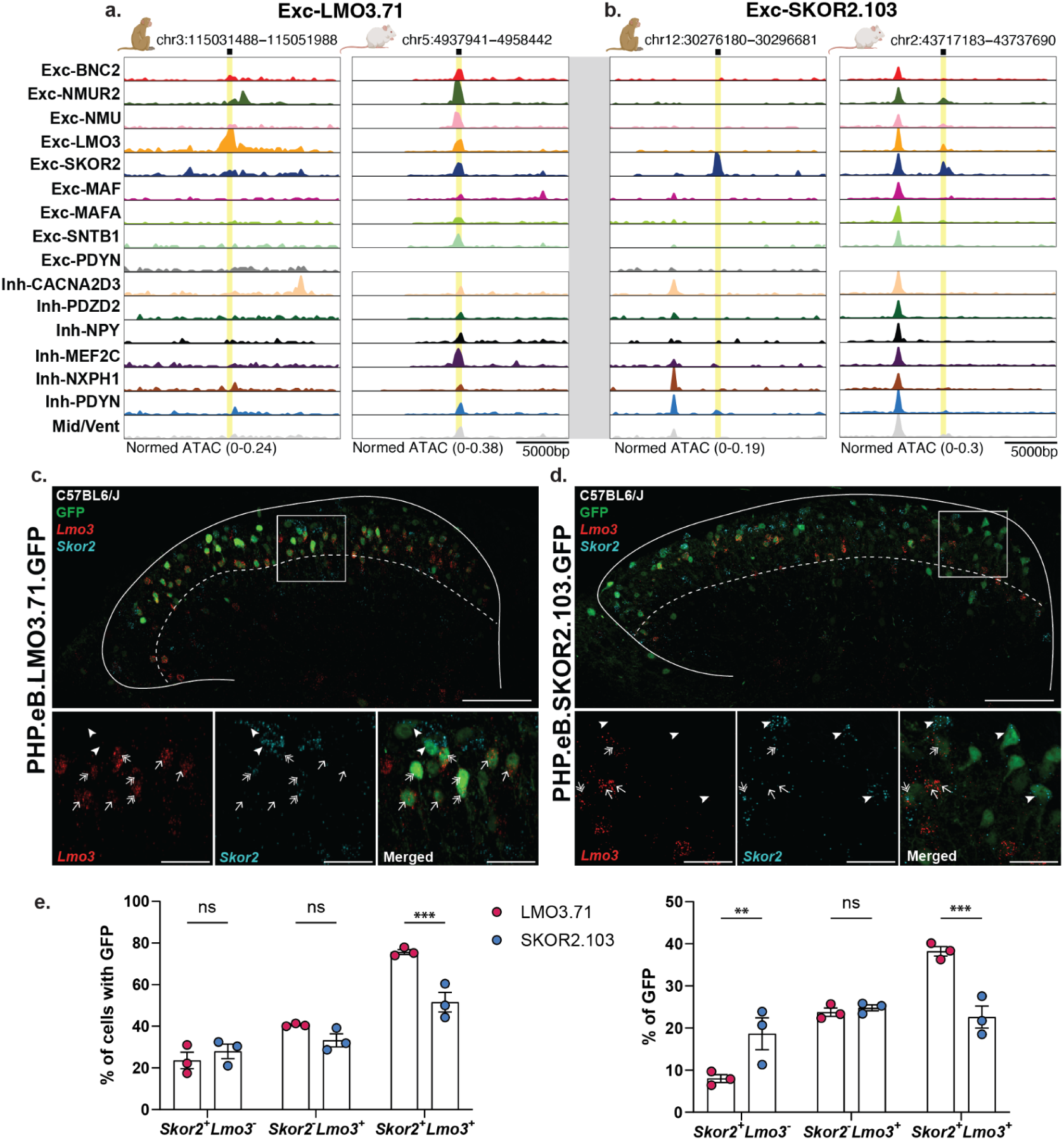
Validation of Two Dorsal Spinal Enhancers. **a and b.** snATAC-seq trackplots of selected regulatory elements (**a.** Exc-LMO3.71 and **b.** Exc-SKOR2.103). Element’s 500bp sequence in macaque (left) and its ortholog in mouse (right). Sequence genomic position (x-axis; coordinates shown at top) is indicated by black squares and yellow rectangle highlights, with surrounding 10KB regions shown on each side. Peak height indicates normalized accessibility for the given subtype (y-axis). **c and d**. Lumbar dorsal horn sections showing sfGFP (green, IF), *Lmo3* (red, FISH), and *Skor2* (cyan, FISH) expression following intraspinal delivery of PHP.eB.LMO3.sfGFP (left) or PHP.eB.SKOR2.103.sfGFP (right). Arrows indicate sfGFP cells expressing *Lmo3*, arrowheads indicate sfGFP cells expressing *Skor2*, and double-arrows indicate sfGFP cells expressing both *Lmo3* and *Skor2*. Scale bars: 100 μm (main images), 30 μm (insets). IF=Immunofluorescence. FISH=Fluorescent in situ hybridization. **e.** Quantitative results from cell counting of RNAscope images (see **c** and **d;** n=3 mice each). Sensitivity based on % of cells from each subtype that are GFP+ (Left) and specificity based on of % of GFP found in each subtype (Right). Data are mean ± s.e.m. Statistical analysis was performed with two-way ANOVA with Bonferroni’s post-hoc test. **P < 0.01, ***P < 0.001.

Work in our companion study (Noh *et al.* co-submitted) shows that Skor2+ neurons mediate chemical itch. Enhancers with higher specificity toward this cell population, such as Exc-SKOR2.103, enable targeted modulation of itch circuits. Indeed, while chemogenetic inhibition via both Excit-1 and SKOR2.103 enhancers attenuates mechanical allodynia, only SKOR2.103 suppresses chemical itch. These results clearly demonstrate that spinal cord enhancers with different cell type-specificities are able to selectively modulate specific behaviors.

## Discussion

In this study, we developed ESCargoT, a cross-species, spatially resolved framework for discovering and profiling of cell type-specific enhancers. We applied that framework in the spinal dorsal horn, a key region for somatosensory processing. By integrating single-nucleus chromatin accessibility data from macaque and mouse with ML-based enhancer prioritization, we identified enhancer candidates predicted to target species-conserved neuron subtypes. We developed a spatial parallel reporter assay, SPRA, by combining a modular AAV assembly pipeline with multiplexed, *in situ* profiling (Xenium) to evaluate enhancer spatial and specificity patterns of expression.

We used an initial machine learning approach to identify the Excit-1 enhancer, which demonstrated dorsal excitatory specificity and reversed mechanical allodynia but not itch *in vivo*. Additional candidates found through cross-species ML prioritization and SPRA screening, Exc-LMO3.71 and Exc-SKOR2.103, showed distinct specificities, and Exc-SKOR2.103 blocked both mechanical allodynia and itch *in vivo*, demonstrating the potential of our platform to link specific neuron populations to behavioral effects.

Sequence-to-function ML prediction provides an important criterion when prioritizing enhancer candidates for testing. For example, for the enhancer Exc-LMO3.71, cross-species machine learning more accurately predicted its validated specificity in mouse, compared to the snATAC-seq signal of its mouse ortholog. Overall, we screened enhancer candidates from the macaque genome that nonetheless showed activity in mouse. Incorporating non-human primate snATAC-seq into ML model training is crucial for identifying enhancers with translational relevance. Enhancer prioritization based on rodent-only data often results in candidates that can work in mice but not primates. Adding in data from multiple species, phylogenetic data augmentation, can substantially improve the performance of sequence-to-function models^35,68,69^. ML models trained on both mouse and primate data could also more effectively learn to overcome subtle cross-species differences in the regulatory grammar. This enables the study of the same enhancers in mouse and primate, including interrogating the relationship between cell-type-specificity and behavioral effects across species.

The neuron subtypes most focused on in this study, namely Exc-LMO3, Exc-SKOR2 and Exc-PDYN, are very closely related based on transcriptomics, epigenomics and their spatial distributions^27^, resulting in an inherently difficult biological problem of targeting one subtype and not the others. Although ML prioritization led to the identification of enhancers with cell-type-specific properties, the precision of this specificity was limited; for example Exc-SKOR2.103, though inhibiting itch as expected, also reduced mechanical allodynia likely because of off-target effects (Noh *et al.* co-submitted). This motivates the development of synthetic enhancers with cell-type and behavioral specificity that could surpass that of naturally occurring sequences. Machine learning models can serve to guide the design of these enhancers^19,70^.

Beyond enhancer prioritization, a major goal of this study was to improve upon multiplexed strategies for enhancer screening *in vivo*. Currently, massively parallel reporter assays (MPRAs) show the greatest success in cell lines^19,71^, which has enabled significant progress in the screening of non-coding loci for causal alleles and the foundational study of ML-guided design of synthetic enhancers^19^. However, these assays do not present enhancers to the fully realistic *in vivo* cellular context, nor do they provide the ability to screen enhancers for their *in vivo* spatial patterns and/or their activity within very closely related neuron subtypes such as Exc-LMO3 and Exc-SKOR2. There is also progress in the technology for multiplexed screening *in vivo*^22,72,73^, but with limitations in sensitivity of detecting viral transcripts^8,21^, and issues with deconvolving signals due to enhancer crosstalk^10,25^.

Xenium screening, in addition to providing insight into properties of specific enhancers, has the sensitivity to reveal biological patterns that influence measures of enhancer specificity, such as injection effects, AAV tropism, cell size which affects AAV uptake, and possibly intrinsic transcription rates. For example, Exc-NTS and Exc-SNTB1, the Prkcg cells in mouse, showed low barcode expression across all enhancers, which aligns with previous observations of failure of these cells to express AAV unless paired with particular promoters (e.g. CAG but not hSyn)^74^. The ability to multiplex allows for the assessment of enhancer candidates relative to inert sequences (negative controls), enabling better normalization across experimental conditions and accounting for injection-specific variability, analogous to the RNA vs. DNA comparisons in cell line MPRA analysis^75^. Multiplexing introduces noise, such as enhancer crosstalk and AAV chimerism^24^, but our regression approach RESSCU partially deconvolves these signals by accounting for each cell’s exposure to the injection, the cellular size that would increase the likelihood of AAV uptake, and the presence of strong promoters (hSyn and SV40) within the cell that could cause the most extreme crosstalk effects. Multiplexing large numbers of candidates at once, as we performed here, also reduces crosstalk effects, by combinatorially reducing the number of interactions of any particular candidate pairs. Finally, we sought to screen enhancer candidates intended to target different neuron subtypes, because crosstalk could otherwise inflate specificity estimates if candidates only target the same subtype^8^.

Our *in situ* profiling offers higher confidence when screening for enhancers that express in rare subtypes such as motoneurons, due to the ability to augment information about marker gene expression with information about cell location, cell size, and by avoiding the pitfalls specific to sequencing^26^. Accurate screening of motoneuron targeting was validated based on hSyn’s low motoneuron expression, a recapitulation of hSyn’s low tropism for cholinergic neurons^14^, and 3/3 barcodes for MN.098 were highly specific to motoneurons. The platform also supports ranking candidates within the same subtype, as demonstrated with motoneuron, oligodendrocyte, and Exc-SKOR2 enhancers.

ESCargoT enables a robust, high-throughput platform for high-confidence, cell-type-specific enhancer identification with exceptional sensitivity and resolution at the level of neuronal subtypes, as demonstrated by Exc-LMO3 and Exc-SKOR2 subtype-selective neuronal and behavioral responses. This level of precision is powerful; decoding enhancer function at the resolution of individual neuronal subtypes not only deepens our understanding of the non-coding regulatory landscape in neural circuits but also establishes a strong foundation for the next generation of precision gene therapies. A growing consensus across the field underscores that effective treatment of neurological or psychiatric disorders often depends on targeting narrowly defined neuronal populations or closely related subtypes, where even minimal off-target expression can result in excitotoxicity, immune activation, or behavioral side effects. Thus, targeting neuronal subtypes with high specificity is both biologically essential and therapeutically imperative. By tailoring genetic payloads to precise cellular contexts, the ESCargoT platform, integrated with emerging insights from the Cell Armamentarium^76^, can offer a promising paradigm for the development of precision therapies that are not only more effective but also better positioned to minimize off-target effects, thereby advancing the therapeutic landscape of genetic medicine.

Overall, ESCargoT offers a powerful framework for decoding enhancer function in complex neural circuits, opening new avenues for precisely targeting therapies in neurological disease.

### Limitations

Single-nucleus chromatin accessibility, the source of the training data for our machine learning models, while state-of-the-art for *in vivo* single-cell enhancer assays, is only moderately correlated with enhancer-driven expression. Intraspinal lumbar injection of AAVs in P21 mice is also state-of-the-art for enhancer screening, but alternate injection strategies, or the study of activity in cervical, thoracic, or sacral spinal cord, or testing in mice of different ages, was not explored.

There are remaining cell-type and subtype-specific biases in the experiment, and remaining difficulties targeting inhibitory neurons with regulatory elements. Overcoming these limitations may require improved machine learning approaches—such as designing synthetic enhancers— and/or alternative injection strategies. False positive detection contributes noise to the Xenium *in situ* screening. For example, in the Xen1 injected mice, barcodes for both random negative controls were detected at moderate levels. Potential other noise sources include: probes binding to single-stranded or episomal DNA, chimerism, residual crosstalk, ITR expression, and residual probe off-targeting of endogenous genes. While these signals can be adjusted for with spatially aware regression, further technological improvements would enable better comparison of weakly expressed cell-type-specific enhancers and more subtle differences between candidates. Finally, several candidates in the assay were not tested with one-at-a-time validation, and enhancer activity in macaque was not evaluated.

### Data Availability and Code Availability

Data and code will be made publicly available at time of publication.

## Methods

### Macaque single-nucleus ATAC sequencing

Fresh lumbar spinal cord (L3-5) was cut at the central canal and split into dorsal and ventral halves. The tissue was dounce homogenized with a loose pestle in 2 ml of Lysis Buffer containing in mM: 10 Tris-Cl, 10 NaCl, 3 MgCl2; ∼25 strokes, and filtered through a 70 mm cell strainer, followed by a centrifugation step at 400 x g for 5 min. at 4°C. After that, the pellet was resuspended in 2 ml of Nuclei Wash and Resuspension Buffer (1X PBS, 1% BSA and RNase inhibitor 0.2 U/mL) and filtered-through a 40 mm cell strainer to optimize myelin and debris removal. Finally, nuclei viability was assessed by trypan blue staining and processed at 10x Genomics Core at the University of Pittsburgh. The 10x libraries were prepared according to the manufacturer’s instructions. Completed libraries were sequenced on the Novaseq 6000 (Illumina).

### Macaque snATAC-seq Pre-Processing, Cell Labeling, and Peak Calling

Bcl files were converted to fastq format using the *cellranger mkfastq* command line tool. Fastq reads were then aligned to the rhemac10 genome using Chromap. Subsequent steps were done with the software suite ArchRv1.0^29^. Arrow files were generated from the aligned fragments and doublet scores were generated using ArchR’s built-in doublet algorithm. An ArchR project was then generated from the Arrow files and sample metadata from the sequencing run. Cells were filtered with minimum transcription start site (TSS) enrichment = 10, minimum number of unique fragments 10^3.5, and doublets filtered out with a cutoff enrichment threshold of 0.5. Fragment periodicity plots were also inspected for intact periodicity of nucleosomes **(Figure S1a**).

Un-biased clustering of cells was done using Iterative LSI, with variable features looped between 30 and 230 thousands of features. 200 thousand variable features were selected having a favorable resolution of clustering with minimal batch effects. Batch correction was performed with Harmony^77^.

We labeled major cell types based on ATAC-RNA integration using a published macaque snRNA-seq resource as the reference^27^. We performed unconstrained integration using ArchR’s implementation of Seurat’s label transfer algorithm using the command addGeneIntegrationMatrix(). We then verified that patterns of gene accessibility correspond to expected marker genes of major cell types, using the gene score, which is calculated from the accessibility at and around the transcription start site of the gene. We then created separate neuron and glia ArchR projects, and re-clustered neurons again with iterative LSI (60 thousand variable features), in order to determine the variability between neuron subtypes. We again performed integration with the neuron subtypes from the aforementioned macaque reference, and verified that marker genes for each neuron subtype were most accessible in the neurons that were labeled as that subtype. We also visualized the distributions of prediction scores for each neuron subtype, a score from 0 to 1 where a higher score corresponds to less ambiguity in the assignment of a label during ATAC-RNA integration (**Figure S1e**). We then re-combined the labeled neuron and glia ArchR projects and generated the coordinates of reproducible peaks using ArchR’s iterative overlap method, then determined presence of reproducible (501 bp) peaks for neuron subtypes using the statistical peak calling method MACS2^78^ called from ArchR’s addReproduciblePeakSet().

### Harmonization of Cell Labels in Macaque and Mouse Dorsal Spinal snATAC-seq datasets

This study utilizes the above described macaque snATAC-seq dataset and a previously published mouse dorsal horn snATAC-seq dataset for cross-species enhancer identification. Both datasets were labeled using the same macaque snRNA-seq reference^27^ with labels previously shown to correspond to neuron subtypes that are conserved across primate and rodent. To provide supplementary evidence of correspondence of the cell type labels between the datasets, we also calculated the relative proportion of ATAC-seq peaks that are shared by the cell types from each species. We first lifted over^79^ the macaque (rheMac10) subtype-specific reproducible peak sets to rheMac8 coordinates to be compatible with halLiftover^80^, which we used in conjunction with the tool HALPER^81^ to map the rheMac8 peaks to mouse (mm10) coordinates to be directly comparable to the reproducible peaks for the cell types in the mouse dataset. For both species, we limited to reproducible peaks unique to one cell type. We then calculated the proportion overlap of cell-type-specific peaks across species by cell type using subsetByOverlaps() and findOverlaps() from the GenomicRanges R package (**Figure S1g**). The proportion of shared peaks for cell types A and B was defined as (# shared peaks)/sqrt((# specific peaks in A)*(# specific peaks in B)).

As an additional way to verify consistent neuron subtypes in both species, we analyzed the enrichment of transcription factor motifs in individual cells using ArchR’s implementation of chromVar^33^ (ENCODE^82,83^ database). This generates a cell-by-motif deviations matrix that represents the enrichment of motifs in accessible peaks relative to expected by chance based on the average of all cells, and after adjusting for PCR and Tn5 tagmentation biases. Finally, using the vector of z-scores of the motif deviations per neuron subtype, we computed the R^2^ of each macaque versus mouse subtype and generated a macaque-subtype by mouse-subtype matrix (**Figure 1h**). We also identified specific TF motifs that are conserved within subtypes across species, by defining motifs with z-score > 1.5 in both species as conserved, aggregating motifs by TF, and TFs with 3 or more conserved motifs (**Figure 2h**). The latter step mitigates false positive risk that would be due to inaccuracies in linking motifs to transcription factors, by requiring that several different TF-associated motifs be conserved.

### Curation of Cross-species Differential Peak Training, Validation, Test Sets

As in^6,18^, we used an ensemble approach for learning sequence patterns of enhancer specificity, by training classification models to predict from differential peak sequences whether they are from the target (foreground) cell type versus from a background class of cells. Background cell classes (vs. GLIA/INH/EXC/MidVen) are defined in **Table S1**. For oligodendrocyte-targeting models, the backgrounds were vs. GLIA/INH/ITexc/nonITexc/Opc/Astro/Microglia/Vasc. INH = inhibitory neurons, EXC/ITexc/nonITexc = excitatory neurons (spinal), or IT and non-IT excitatory neurons (cortex). Whenever a foreground cell type would normally be part of the background class, the background was modified to exclude the foreground cell type. Because of the potential differences in the regulatory grammar of promoters versus distal enhancers^40^, peaks for training, validation, and testing were limited to distal and intronic peaks 20KB or more from their nearest TSS. Positive and negative differential peaks were then calculated using ArchR’s getMarkerFeatures(useGroups=foreground, bgd.groups=background, testMethod= “wilcoxon”). Positive (negative) examples were defined as the differential peaks satisfying FDR-adjusted significance < 0.05 and that were logFC > 0.5 (positive example) or logFC< −0.5 (negative example).

We generated .bed files of the positive and negative peaks, then split into training/validation, and test sets, whereby the training/validation sets were further organized into five-fold cross-validation 80/20 training/validation splits. To avoid contaminating validation and test sets with orthologues peaks from the training set of the other species, we used a synteny approach whereby we pre-defined the training, validation, and test sets in reference to human (hg38) chromosomes. Then, a macaque or mouse peak was assigned to a given category by determining which synteny block it belongs to and assigning the orthologous human category for that synteny block. Synteny blocks were determined using halSynteny^84^. The test set was pre-defined as human chr1 and chr2. Validation sets were defined for each fold as follows: *fold1 = chr6, chr13, chr21: fold2 = chr7, chr14, chr18; fold3 = chr11,chr17, chrX; fold4 = chr9, chr12; fold5 = chr, chr10*. The training set for a given fold was the set of human chromosomes not in the test set or in the validation set of that fold.

### Motif Enrichment Calculations

We identified cell-type-specific transcription factor binding motifs enriched in differentially accessible peaks using MEME suite with the human HOCOMOCOv11 reference motif database. We calculated *Z*-score normalized motif enrichment for each individual peak.

### Machine Learning Models to Predict Enhancer Specificity

For each combination of target cell type and background class, we trained the following binary classification models to predict, from DNA sequence, whether a differential peak is from the target cell type or the background class: (1) a support vector machine (SVM) using the mouse training set, (2) an SVM using the macaque training set, and (3) a cross-species convolutional neural network using the combined training sets. The SVMs were trained on fold 1 while the CNNs were trained on all five folds.

The SVMs were developed using the gkm-SVM^42,85^ software, which uses the frequency of gapped k-mers of the input sequences as the features for the SVM. We chose the following methods and hyperparameters using the command gkmtrain: t=4 (center-weighted gkm), k=7 (k-mer length), k=6 (number of informative columns), d=1 (mismatches to consider), M=50, H=50, m=10000, T=4, and -s. The hyperparameter w for class imbalance was set to the ratio of positives to negatives for the particular model. The regularization hyperparameter c was tuned between 0.1 and 1 and the best performing model was selected.

The CNNs were developed using custom software by the Pfenning Lab based on Keras 2. 500bp input sequences were one-hot encoded (500×4). Binary classification models trained and evaluated on one-hot-encoded sequences were designed as followed: with two 1-dimensional convolutional layers each followed by maximum pool layers of size 26 and stride 26, and with 500 convolutional filters each with width 11 units and a stride of 1, followed by a dense layer of 300 units, and finally a single output unit with sigmoid transformation. Models were trained with a dropout rate of 0.3 and L2 regularization across layers of 1e^-4. The following hyperparameters were used: batch size 512, adam optimizer with constant momentum (beta1 = 0.9, beta2 = 0.999, base momentum = 0.875, maximum momentum = 0.99), and an exponential learning rate (initially 4*10^-4, decay by factor 0.95 per epoch). Weights were initialized based on the Glorot Uniform distribution. Models were trained up to 50 epochs with early stopping criteria based on validation auPRC with a patience of 5 epochs.

For each fold of cross-validation, we evaluated the following performance metrics on the cross-species validation sets: F1 score, accuracy, precision, recall, sensitivity, specificity, negative predictive value, NPVSC^35^, areas under the receiving operating characteristic and precision-recall curves (auROC and auPRC). We also calculated the same metrics for the validation sets of each species separately (**Table S1**). For any metrics based on a single threshold to differentiate predicted positives from predicted negatives, we used a threshold of 0.5.

## Enhancer Prioritization

As in ^18^, for each target neuron subtype, differential peaks for that subtype 2KB or more from the nearest gene were ranked by the following procedure. Subtype-specific MEME motifs were counted and z-scored across all sequences. Then for any given sequence, its motif z-score, geometric-averaged CNN logit scores, and geometric-averaged SVM logit scores, were again averaged together (geometric mean). This provides an aggregate ranking score for each sequence. Finally, enhancer candidates were selected for experimental validation by searching top-ranked sequences and considering other factors such as species conservation of the peak, atac-seq specificity signals, and the presence or absence of nearby marker genes.

### Interpretation of CNNs

We performed CNN model interpretation using Deep SHAP version 0.37.0^43,86^. For each target subtype, with prioritized candidates as the foreground, we generated per-base importance scores and hypothetical importance scores relative to the reference set of negative sequences from the validation set. TF-MoDISco aggregates results from DeepSHAP and uses commonly important subsequences, seqlets, to construct the motifs that the model learned to prioritize. We ran TFMoDISco version 0.4.2.3 with the options sliding_window_size = 11, flank_size = 3, min_seqlets_per_task = 3000, trim_to_window_size = 11, initial_flank_to_add = 3, final_flank_to_add = 3, kmer_len = 7, num_gaps = 1, and num_mismatches = 1. The resulting motifs were filtered to remove rare patterns with 100 supporting seqlets. Then, the motifs were visualized and associated with known motifs using Tomtom^44^ version 5.3.3 with the HOCOMOCO v11 FULL database and default parameters. Motif results provided in **Table S2**.

#### Strains and growth conditions

*E. coli* strain 10-beta (DH10B, NEB) was cultured at 37°C in liquid LB medium or on LB agar plates (Difco), supplemented with 35 µg/ml chloramphenicol or 100 µg/ml ampicillin (RPI) for selection.

#### Cell lines

HEK 293 cells (CRL-1573 ATCC) were cultured in growth media DMEM (with 4.5 g/L glucose, l-glutamine, and sodium pyruvate) supplemented with 10% fetal bovine serum under standard manufacturer’s guidelines in a tissue culture incubator at 37 °C and 5% CO_2_. The AAV was produced in AAVpro(R) human embryonic kidney 293 cells (Takara Bio USA, #632273) and cultured according to manufacturer’s instructions.

#### Genetic constructs

**The part-plasmid modules.** The SPRA GG assembly pipeline of the pAAV_A and pAAV_B vectors were constructed using part-plasmid modules or a combination with part-plasmid modules and G-blocks (IDT or Twist Biosciences) directly. For the part-plasmid modules the GFP drop-out vector pYTK001 vector was used^87^. This vector contains IIS type BsmBI sites up and downstream of the GFP gene that allows for the construction of GG compatible part-plasmid modules for the SPRA pipeline with IIS enzymes BsaI or PaqC1. Building the part-plasmid modules for step 1 and step 2 in the SPRA cloning pipeline follows the general design principle. From 5’-3’, each G-block contains a type IIS BsmBI site followed by a IIS BsaI site (for the pAAV_A plasmids) or a IIS PaqC1 site (for pAAV_B vectors), 4N module specific overhang (MSO) sequence as previously published^56^ and the specific SPRA module sequences (see **Figure 4** and **S5**). All part-plasmid modules were constructed with synthesized G-blocks that were codon optimized for mammalian systems. The pipeline’s modules were built utilizing several different DNA templates and genetic cloning approaches. For the modules M1A and M1B that contain the core parts of the pAAV vector, DNA template of pAAV-EF1a-Sun1GFP-WPRE-pA was used as previously reported^4^ and adapted and designed for GG cloning with genetic engineering software Geneious prime version 2024 (Geneious). The DNA sequences of the part-plasmid M2 modules and G-blocks for the SPRA pipeline are listed in **Tables S11** (individual Enhancers/ barcodes) and **S12** (library).

Module M1A containing the core backbone pAAV ITR’s, the Ampicillin selection cassette and the origin of replication (ori) including the f1 ori and was constructed using two separate fragments. Fragment 1: From 5’ to 3’, IIS BsmBI site-IIS BsaI site-4MSO (AACC)-ITR-f1ori-and part of the AmpR cassette-overhang site (AGTG)-IIS PaqC1 site and IIS BsmBI site. Fragment 2: From 5’ to 3’, IIS BsmBI site-IIS PaqC1 site-overhang site (AGTG)- rest AmpR cassette- *E.coli* ori- ITR- MSO (TAGA)-IIS BsaI-IIS BsmBI. Subsequently, the 2 fragments of module M1A were cloned in a one pot GG reaction into the entry vector pYTK001 with IIS enzymes BsmBI and PaqC1. Module M1B containing the pAAV elements WPRE and PolyA was synthesized (Twist biosciences) and cloned using GG assembly with IIS BsmB1 into the entry vector. Module M2 contains 500bp Enhancer, IIS PaqC1 drop-in and 500bp Barcodes sequences (**Table S11** and **S12**). Here, multiple templates are used for building the single enhancer-barcode constructs. The standard enhancer DNA sequences SV40, CMVe and CMVe+p were used (**Table S11**). For the no enhancer constructs (NoE) the 500bp enhancer sequence was excluded. The library enhancer and barcode sequences were designed by using our computational models (see methods below). Module M3 contains the intron and the fluorescent reporters with a nuclear localisation signal (NLS). The sequence for sfGFP was derived from pAAV-EF1a-sun1-GFP-WPRE-pA^6^, and the DL5 sequence was obtained as previously described^56^ and subsequently synthesized (Twist Biosciences). The plasmids constructed for this study are presented in **Table S10**.

#### Golden Gate assembly of pAAV_A and pAAV_B vectors for SPRA

For this pipeline, we leveraged the well-established Golden Gate (GG) assembly strategy from New England Biolabs^88^ combined with the 13-piece GG vector system for the Nanobody discovery platform developed at Carnegie Mellon Institute^56^. The GG technology is modular, highly flexible and extremely efficient for library construction because it can be pushed to 100% efficiency if the GG assembled fragments are under 4 pieces. The pAAV_A vectors were assembled from BsaI-compatible part-plasmid modules (**Figure S5a**) by using an adapted version of the GG cloning as previously published^2^. In step 1 of the SPRA cloning pipeline, Golden Gate (GG) cloning was performed using a one-pot, three-piece assembly reaction. The part-plasmid modules M1A, M1B, and M2 were assembled into the pAAV_A vector. For library construction, G-blocks corresponding to Module M2 were directly utilized to assemble the pAAV_A vectors. The G-blocks of the spinal cord library were pooled into 3 pools; Xen1.1, Xen1.2 and Xen1.3, and were re-calibrated to 50 ug/ul (**Tables S6/S8**). For the full Xen1 library construction, the 3 pools (Xen1.1, Xen1.2 and Xen1.3) were combined. GG assembly reactions were set up with 75 ng of part-plasmid modules in a reaction mix containing 1× T4 DNA ligase buffer (NEB), 2.0 μL GG assembly mix (E1601, NEB), 0.5 μL T4 DNA ligase (NEB), and 0.75 μL BsaI-HFv2 (NEB), in a final volume of 20 μL. The pAAV_B vectors were assembled with the pAAV_A vectors from step 1 and PaqC1 compatible part-plasmid module M3 (**Figure S5c**). Step 2 of the SPRA cloning pipeline is using a one-pot, two-piece assembly reaction and were performed using 75 ng pAAV_A and 75 ng part-plasmid module M3 in GG reaction mixture 1× T4 DNA ligase buffer (NEB), 2.0 *μ*L T4 ligase (NEB), 1.25 *μ*L PaqC1-HF (NEB), 0.75 *μ*L PaqC1-activator (NEB) in a final volume of *V*final = 20 *μ*L. The reactions were carried out using a Mastercycler Nexus X2 (Eppendorf) with the following program: 10s at 37°C, 30 cycles of 2 min at 37°C and 2 min at 16°C, followed by 5 min at 60°C, and a final hold at 4°C. The GG reaction mixtures were subsequently used for *E. coli* transformations as described below. The part-plasmids constructed for this study are presented in **Table S10**.

#### Restriction digestion RD-profiling

The constructed part-plasmid modules M1A, M1B, M2 were analyzed for correct insert sizes by restriction digestion with BsaI-HFv2 (NEB) and M3 with PaqC1 (NEB). For the pAAV_A and pAAV_B vectors, correct GG assembly was analyzed by simultaneously using three RD enzymes HinDIII-HF, NotI-HF, and PstI-HF (all NEB). The RD-profiling experiments were executed according to manufacturer’s guidelines (NEB) with minor modifications as described previously^56^. RD reactions were conducted with in the reaction mixture Vfinal = 15μL in cut smart buffer (NEB) containing 0.25 μL enzymes, 300-600 ng plasmid DNA and incubated for 15-25 min at 37°C in a Mastercycler Nexus X2 (Eppendorf). Next, the enzyme reaction mixtures were analyzed using DNA gel electrophoresis on 2% TAE agarose gels (40 mM Tris, 10 mM EDTA, 20 mM acetic acid) containing 1.0 μg/mL Ethidium Bromide. Electrophoresis was performed in TAE buffer containing 1.0 µg/mL ethidium bromide using a mini gel electrophoresis system (Horizon 58, BRL/Life Technologies) and power source (Thermo Fisher Scientific) set to 100V, 50 mA for 30 min at RT.

#### Sequencing part modules and pAAV vectors

Part-plasmid modules and single constructed pAAV_A and pAAV_B vectors were diluted in TE buffer (IDT; PH = 8.0)) to a final concentration of 30ng/μL and sent out for whole plasmid sequencing (Eurofins Genomics). Subsequently, the sequence data was analyzed by using Geneious prime software (Geneious, version 2024) utilizing the reference genetic sequences of the part-plasmid modules or the full sequences of the pAAV_A/ pAAV_B vectors to confirm the correct genetic sequence. As a critical validation step in the SPRA cloning pipeline, the pAAV_A and pAAV_B vectors were rigorously analyzed for the presence and accuracy of module specific overhang (MSO) sequences, which form the cornerstone of the SPRA cloning strategy. As expected in GG strategy using high fidelity MSOs, 100% of the tested pAAV vectors exhibited correct MSO sequences, underscoring the precision and reliability of the cloning pipeline.

#### Library pAAV_A and pAAV_B Xen1 vectors spinal cord

Each library was constructed using independent GG assemblies and transformations for the pAAV_A and pAAV_B vectors. For the pAAV_A library, two independent GG assemblies were performed, and 5 x 150 µl of transformed cells (in outgrowth medium) were each inoculated into 5 ml of LB medium supplemented with 100 µg/ml ampicillin. After culturing, plasmid isolation was carried out on 1.5 ml of each culture, and the purified plasmids were pooled to maximize library complexity. For the pAAV_B library, three independent GG assemblies and transformations were performed. 6 x 150 µl of transformed cells were each inoculated into 5 ml of LB medium with 100 µg/ml ampicillin, cultured, and plasmid isolation was carried out on 1.5 ml of each culture. The purified plasmids were then pooled to ensure full library complexity.

#### Bacterial Transformations and HEK293 cell line Transfections

*E. coli* transformations were performed using high-efficiency competent 10-β cells (NEB, C3019) following an adapted protocol from NEB’s (5-Minute Transformation *Protocol using NEB 10-beta Competent E. coli (C3019H/C3019I)*; NEB). Briefly, 25 μL or 50 μL (Xen1 libraries) of frozen competent cells were thawed on ice for 10 minutes, then mixed with 3.5 μL of the GG assembly mix for pAAV_A, pAAV_B vectors or the part-plasmid modules, respectively. The cells were incubated on ice for 30 minutes, followed by a heat shock for 32S at 42°C (water bath). After a 5-minute incubation on ice, 600 μL of stable outgrowth medium (NEB) was added, and the cells were shaken at 250 rpm (Anova 40R shaker) at 37°C. Finally, the transformed cells were plated on 2% LB agar plates supplemented with 100 μg/mL ampicillin (for pAAV_A and pAAV_B vectors) or 35 μg/mL chloramphenicol (for part-plasmid modules) and incubated overnight at 37°C for clone selection, plasmid isolation (Qiagen) and downstream experiments. For the Xen1 libraries the cells were grown in LB liquid medium supplemented with 100 μg/mL ampicillin and these libraries containing cell culture were used for plasmid isolation (Qiagen) and downstream experiments. HEK293 Cell line (CRL-1573 ATCC) transfections were performed by using electroporation (Biorad Gene pulser Xcell) in 2 mm cuvettes (Genlantis, Zeus C902050) with 200 μL 1.0 X10^6^ cells in DPBS (Gibco) and 1.0 μg purified pAAV_B plasmids under the manufacturer’s standard pre-set Hek293 protocol. Post transfection, cells were cultured for 24 hrs and subsequently lifted and co-cultured on optical-bottom (Greiner, #655090) 96 wells-plates for 24 hrs to allow Keyence fluorescent microscopy. Cells were cultured for an additional 24 hrs prior to fluorescence imaging.

#### Keyences Fluorescent Imaging

Fluorescence of the reporters sfGFP and FAP DL5/MG-ester was measured using a Keyence BZ-X800 series all-in-one fluorescent microscope with a BZ-X810 light source. For sfGFP, the Keyence 49003_UF1 filter set (ET-EYFP, ET500/20x; ET535/30m, T515lp-UF1, Lot: C-220059) was used with a scan range of 480-560 nm. For FAP/fluorogen fluorescence, the Nikon Eclipse filter set (ET Cy5, ET620/60X, ET700/75m, T660pxr, Lot: C211482) was used. Imaging was performed with PanApo objective lenses: X10 (NA 0.45), X20 (NA 0.75), and X40 (NA 0.95). FAP fluorogen fluorescence was achieved by adding MG-ester to the cells or tissue at a final concentration of 150-250 nM. For imaging HEK293 cells, MG-ester was added directly to the culture 1 hour before analysis. For imaging the brain or spinal cord, MG-ester in PBS + 0.5% PF127 was applied directly to the tissue slices and incubated for 1 hour and subsequently washed 3 times with PBS + 0.5% PF127 before analysis. All images were processed and analyzed using Keyence BZ-H4A software and ImageJ.

#### Next generation sequencing

Next-generation sequencing was completed to analyze the barcode proportions of the libraries. Samples were amplified from 0.5 ng of plasmid DNA in a two step PCR protocol. First, the plasmid library was amplified for 10 cycles using NEBNext High-Fidelity PCR Master Mix (NEB) on the Proflex Thermo Cycler (Applied Biosystems) with 280-NGS_BC_Fw_3 and 281-NGS_BC_Rv_4 primers (**Table S13**). Each library was amplified in three reactions, then pooled and purified using the MinElute PCR Purification Kit (Qiagen). The second amplification step was performed for 26 cycles with NEBNext Master Mix and primers synthesized from Eurofins Genomics following the Illumina Nextera tagmentation format as previously published^89^. The second amplification phase was also completed with three separate reactions for each library. The PCR products were combined and purified using the AMPure XP Bead system (Beckman Coulter) with size selection at 0.55x and 0.85x ratios of beads. Library quality was assessed on a TapeStation using D1000 Screen-tapes (Agilent) and concentration was measured on a Qubit Fluorometer (Thermo Fisher Scientific). Each sample library was pooled and sequenced at 6 pM with 10% PhiX (Illumina, #FC-110-3001) on the Illumina MiSeq platform using a 300-cycle, Micro V2 Kit (Illumina #MS-103-1002). Sequencing runs on the MiSeq were performed following the manufacturer’s guidelines (Illumina).

#### pAAV Production

The PHP.eB capsid variant of Adeno-associated viruses (AAV) were all produced in the Pfenning lab. AAVpro® 293 cells (Takara, catalog #632273) were co-transfected with linear polyethylamine, 25 kDa (PEI), pAAV_B plasmids (single or the libraries), AAV helper plasmid, and pUCmini-iCAP-PHP.eB (gifted by Viviana Gradinaru; Addgene plasmid #103005, RRID)^63^. Media supernatant was collected after 72 and 120 hours and polyethylene glycol (PEG) was used to precipitate and collect the AAV from the media. The transfected cells were collected 120 hours post-transfection and lysed with Salt Active Nuclease (SAN, ArcticZymes). The cell lysate and AAV collected from the media supernatant was then combined and purified through an iodixanol gradient (OptiPrep, Sigma Aldrich) with 15, 25, 40, and 60% layers. Virus was collected from the 40-60% interface, then further purified and concentrated with Amicon Ultra 15s filters (Millipore). Viral titers were determined using the AAVpro(R) Titration Kit (Takara, catalog #6233) as previously described^90^. The final virus preparation was aliquoted and stored at −80°C until further use. AAV8.Excit-1.hM4Di-2a-EGFP was generated by subcloning Excit-1 gene fragment into an existing vector. Specifically, Excit-1 gene fragment was made by Twist Bioscience and subcloned into the pAAV.SYN1.HA-hM4Di (Addgene, plasmid# 121538)^91^ using MluI (New England Biolabs; R3198S) and KpnI (New England Biolabs, R3142S) restriction enzyme sites. HA-hM4Di sequence was replaced by hM4Di-2a-EGFP from AAV.EF1α.Flex/3’USS.hM4D-2A-EGFP(ATG mut) (Addgene, plasmid # 197892)^92^ using a PCR kit (New England Biolabs, M0530S) and restriction enzymes KpnI and HindIII (New England Biolabs, R3104S). Whole plasmid sequencing was performed by Plasmidsaurus, and the plasmid was packaged into AAV8 by Neurotools viral vector core (UNC).

#### Animals

Animal procedures were conducted with approval from Carnegie Mellon University and University of Pittsburgh IACUCs and followed the *Guide for the Care and Use of Laboratory Animals* (National Research Council, United States). Molecular and imaging studies utilized male and female C57BL/6J mice (strain #000664, The Jackson Laboratory). Adeno-associated virus (AAV) was injected into mice via retro-orbital injection or direct injection into the spinal cord (intraspinal injections) and used for library sequencing, fluorescent imaging and Xenium experiments. For Retro-orbital injections, mice were anesthetized with 3-4% isoflurane until respiration slowed and no pedal reflex was observed, then maintained with 1-2% isoflurane until injections were complete. 4×10¹¹ vector genomes (vg) total were injected at a volume between 50-100uL. Following the injection, 0.5% proparacaine hydrochloride ophthalmic solution was applied to the eye for comfort, and mice were monitored for any adverse side-effects post-procedure.

The intraspinal injection procedure and injection details were performed as previously described^93^ with minor modifications. Mice, 21-22 days old, were deeply anesthetized with 4% isoflurane and maintained with 1.5-2% isoflurane throughout the procedure. A midline incision was made to expose the thoracolumbar vertebral column. Without performing a laminectomy, the intervertebral space between T12-T13 (to target lumbar segments L3-L4^94^ or T13-L1 (to target lumbar segments) was exposed on the left side with blunt dissection. The vertebral column was stabilized by clamping at L1 using V-notch spikes (Kopf Instruments). After making a small tear of the dura, a glass microelectrode (tip diameter ∼50 µm) was inserted 350-400 µm lateral to the dorsal midline blood vessel to a depth of 200 µm from the dorsal surface using a stereotaxic frame (Stoelting or Kopf Instruments). AAV vectors were infused over a period of 5 minutes using a stereotaxic injector (Stoelting or Drummond Scientific). The micropipette remained in place for at least 3 minutes post-infusion before being slowly withdrawn. The latissimus dorsi muscle was closed with 6-0 nylon sutures, and the skin was closed with 6-0 silk sutures. Unless otherwise stated, behavioral and histological experiments were performed at least 3 weeks post-injection to allow for maximal and stable viral transgene expression.

The AAV library injections of Xen1.1 and Xen1 were executed with 300 nl or 500 nl of PHP.eB-X1.1-DL5 (2.95 x 10^13^ vg/ml) or PHP.eB-X1-DL5 (5.4 x 10^13^ vg/ml)^94^. AAV containing the library Xen1.1 or Xen1 were allowed to express for at least 4 weeks before spinal cords were collected for RNAScope™ or Xenium experiments. 300 nl of AAV8.Excit-1.hM4Di-2a-EGFP, PHP.eB.SKOR2.103.hM4Di-2a-EGFP, and PHP.eB.LMO3.71.hM4Di-2a-EGFP (1.00 x 10^12^ vg/ml) were injected intraspinally as described above.

### Chemogenetic Activation

To activate the hM4Di DREADD receptor, clozapine-N-oxide dihydrochloride (CNO; 5 mg/kg; Tocris, 6329) was dissolved in sterile saline and administered intraperitoneally (i.p.) 30 minutes prior to the start of each experiment. A crossover design was used, in which animals received both CNO and vehicle (sterile saline) injections on separate test days. All injections were performed in an identical manner, and the vehicle condition served as the control. The experimenter was blinded to treatment conditions throughout all phases of data collection and analysis.

### Behavioural Assays

Mice were tested as previously described^74^. Prior to treatment, they were habituated for at least one hour in opaque Plexiglas chambers positioned on a wire mesh table. Punctate mechanical allodynia was assessed using the von Frey test with a set of calibrated Semmes-Weinstein monofilaments, following the Up-Down method starting with a 0.4 g filament. Filaments were applied perpendicularly to the plantar surface of one or both hind paws for 3 seconds, or until a clear nocifensive response, such as paw withdrawal, shaking, or licking, was observed. Responses due to rearing or normal ambulation were excluded. Stimuli were applied at 5-minute intervals, and the 50% paw withdrawal threshold (PWT) was calculated for each mouse.

Dynamic mechanical allodynia was evaluated using a cotton swab, which was teased out to approximately three times its original size. The swab was gently stroked across the hind paw from heel to toe. A response was considered positive if the animal withdrew, shook, or licked the paw; absence of these behaviors was recorded as a negative response. Each paw was tested six times at 3-minute intervals, and the percentage of positive responses was calculated per animal. Complete Freund’s Adjuvant (CFA) model of inflammatory pain. Mice were anesthetized with 4% isoflurane for induction and maintained at 2.5% during the procedure. A 10 μL injection of a 1:1 emulsion of Complete Freund’s Adjuvant (CFA; Sigma, F5881) and sterile saline was administered into the glabrous surface of the left hind paw. Behavioral assays were conducted 3–5 days post-injection.

### Validation of Barcode Expression by RNAscope™

2 male C57BL/6J mice, one injected intraspinally and one injected retro-orbitally, were used to validate the expression of barcodes using RNAScope^TM^ as described previously^3^. A custom probe for barcode sequence driven by the 500bp hSyn promoter was used to validate the expression of the X1 library in the spinal dorsal horn. The custom barcode was amplified and visualized with TSA Plus Cy5 reagent (Akoya Biosciences; 1:1500). Spinal cords were imaged using a Nikon A1 confocal microscope.

### Xenium Spatial Transcriptomics Panel

Per 10x guidelines, we provided a reference snRNA-seq^31^ of the mouse spinal cord in order to evaluate candidate Xenium probes against genes naturally expressed in the target tissue. We then provided 100 targets for a custom panel; 6-8 probes per target were selected by 10X based on their bioinformatics procedure. The custom panel consisted of probes targeting 81 synthetic barcode sequences (**Table S12**), EGFP, mCherry, DL5, and 16 naturally occurring genes:

Mef2c, Rbfox3, Npy, Nxph1, Nmur2, Lmo3, Skor2, Bnc2, Maf, Nmu, Cacna2d3, Grpr, Mafa, Mog, Tac1, Sorcs1

The custom panel was also used in combination with the standard mouse brain panel, which consisted of probes for the following genes:

2010300C02Rik, Acsbg1, Acta2, Acvrl1, Adamts2, Adamtsl1, Adgrl4, Aldh1a2, Angpt1, Ano1, Aqp4, Arc, Arhgap12, Arhgap25, Arhgap6, Arhgef28, Bcl11b, Bdnf, Bhlhe22, Bhlhe40, Btbd11, Cabp7, Cacna2d2, Calb1, Calb2, Car4, Carmn, Cbln1, Cbln4, Ccn2, Cd24a, Cd300c2, Cd44, Cd53, Cd68, Cd93, Cdh13, Cdh20, Cdh4, Cdh6, Cdh9, Chat, Chodl, Chrm2, Cldn5, Clmn, Cntn6, Cntnap4, Cntnap5b, Cobll1, Col19a1, Col1a1, Col6a1, Cort, Cplx3, Cpne4, Cpne6, Cpne8, Crh, Cspg4, Cux2, Cwh43, Cyp1b1, Dcn, Deptor, Dkk3, Dner, Dpy19l1, Dpyd, Ebf3, Emcn, Epha4, Eya4, Fezf2, Fgd5, Fhod3, Fibcd1, Fign, Fmod, Fn1, Fos, Foxp2, Gad1, Gad2, Gadd45a, Galnt14, Garnl3, Gfap, Gfra2, Gjb2, Gjc3, Gli3, Gm19410, Gm2115, Gng12, Gpr17, Grik3, Gsg1l, Gucy1a1, Hapln1, Hat1, Hpcal1, Hs3st2, Htr1f, Id2, Igf1, Igf2, Igfbp4, Igfbp5, Igfbp6, Igsf21, Ikzf1, Inpp4b, Kcnh5, Kcnmb2, Kctd12, Kctd8, Kdr, Lamp5, Laptm5, Ly6a, Lypd6, Mapk4, Mdga1, Mecom, Meis2, Myl4, Myo16, Ndst3, Ndst4, Necab1, Necab2, Nell1, Neto2, Neurod6, Nostrin, Npnt, Npy2r, Nr2f2, Nrep, Nrn1, Nrp2, Nts, Ntsr2, Nwd2, Nxph3, Opalin, Opn3, Orai2, Paqr5, Parm1, Pcsk5, Pde11a, Pde7b, Pdgfra, Pdyn, Pdzd2, Pdzrn3, Pecam1, Penk, Pglyrp1, Pip5k1b, Pkib, Plch1, Plcxd2, Plcxd3, Plekha2, Pln, Pou3f1, Ppp1r1b, Prdm8, Prox1, Prph, Prr16, Prss35, Pthlh, Pvalb, Rab3b, Rasgrf2, Rasl10a, Rbp4, Rfx4, Rims3, Rmst, Rnf152, Ror1, Rorb, Rprm, Rspo1, Rspo2, Rxfp1, Satb2, Sdk2, Sema3a, Sema3d, Sema3e, Sema5b, Sema6a, Shisa6, Siglech, Sipa1l3, Sla, Slc13a4, Slc17a6, Slc17a7, Slc39a12, Slc44a5, Slc6a3, Slfn5, Slit2, Sncg, Sntb1, Sorcs3, Sox10, Sox11, Sox17, Spag16, Spi1, Spp1, Sst, Stard5, Strip2, Syndig1, Syt17, Syt2, Syt6, Tacr1, Tanc1, Th, Thsd7a, Tle4, Tmem132d, Tmem163, Tmem255a, Tox, Trbc2, Trem2, Trp73, Trpc4, Unc13c, Vat1l, Vip, Vwc2l, Wfs1, Zfp366, Zfp536, Zfpm2.

### Tissue Preparation and Xenium Spatial Transcriptomics Imaging

3 female and 3 male P60-75 CB57BL/6J mice were intraperitoneally injected with 100 mg/kg ketamine and 20 mg/kg xylazine. Mice were perfused with ice-cold RNAse-free PBS and the L3-L5 lumbar segment of the spinal cords were dissected. The spinal cords were then embedded in optimal cutting temperature (OCT) compound and frozen following the 10x Genomics’ Xenium In Situ for Fresh Frozen Tissues - Tissue Preparation Guide. L3-L5 segment of the spinal cords were cryosectioned at 10 μm with at least 40 μm between each section. Spinal cord slices were sectioned onto Superfrost Plus^TM^ microscope slides for RNAScope^TM^ validation or Xenium slides for spatial transcriptomics analysis with Leica CM1950 cryostat. Slides were stored at −80 ℃ until the RNAScope^TM^ or Xenium assay was performed. 2 males and 2 female C57BL/6J mice injected intraspinally. The Xenium experiments were performed following the manufacturer’s guidelines: https://cdn.10xgenomics.com/image/upload/v1710785024/CG000584_Xenium_Analyzer_UserGuide_RevE.pdf.

### Xenium Cell Labeling

Neurons were labeled based on a previously described and validated approach^27^ with minor modifications. As in ^27^, a mouse snRNA-seq dataset from Russ et al. (2021)^31^, previously ascribed species-conserved neuron subtype labels from ^27^, was used as a reference to annotate the Xenium dataset. The reference was hierarchically organized into three annotation levels: major cell type, neuron subclass, and neuron subtype. At the cell type level, cells were broadly categorized as neuron or various glial and immune cell types. Within the neuronal subclass, neurons were divided into mid/ventral versus dorsal horn versus Exc-PDYN. Neuron subtypes included mid/ventral excitatory, inhibitory, and motorneurons, and the dorsal subtypes as described in ^27^.

Xenium cells were labeled based on iterative integrations across the three annotation levels with the reference snRNA-seq described above. snRNA-seq genes were limited to those also measured by Xenium, and both assays were log-normalized. First, labels were transferred at the major cell type level using Seurat v5’s FindIntegrationAnchors() and IntegrateData() based on canonical correlation analysis (CCA), and finally TransferLabels(). CCA was used because of major differences in the dynamic range of measured genes for snRNA-seq versus Xenium. Following cell-type-level annotation, both the reference and query datasets were subsetted to neurons. A second round of integration was conducted to assign neuron subclass annotations. This process was repeated once more within each subclass subset to assign precise neuron subtypes.

Based on the observation that Exc-SKOR2 forms two distinct clusters in both Xenium and snRNA-seq, we split this subtype into Exc-SKOR2.1 and Exc-SKOR2.2. Finally, only cells with an integration predicted confidence score above 0.5 were kept for further analysis.

### Estimation of Spatial Injection Spread

The spatial coordinates of the negative control barcodes were gathered for each slice of each run. These were 3/3 barcodes corresponding to CS-neg-1, and 2/3 barcodes corresponding to CS-neg-2 (1 removed for high expression following virus-free experiment). Each slice was broken up into a grid where each square was 75 x 75 um across, such that an average of ten cells were present in each square. For each of the two negative controls, using kde2d() from the R package MASS, the kernel density of the barcodes was estimated across the grid squares. The densities were normalized such that the sum of the densities corresponded to the sum of negative control barcodes found in the slice of interest. Then each cell in the slice was assigned to the corresponding density estimate of the grid square to which it belongs, for each of the two negative controls. After comparing the similarity of distributions of each negative control, the final density assigned to each cell was the average of the densities for both negative controls.

### Estimation of Crosstalk

Enhancer crosstalk is the phenomenon when a barcode or reporter of one AAV is driven by the regulatory element from a different AAV, and is a concern when multiple, different AAVs are tested in the same animal and region that can lead to inaccurate measures of specificity and activity for each individual enhancer^25^. To mitigate the impact of this source of noise when assessing enhancer specificity, we (1) created a relatively large library of 27 unique enhancers and 81 unique barcodes, such that pairwise interactions of any two unique enhancer-AAVs was minimized. We also (2) measured crosstalk risk using the counts of hSyn and sv40 as a proxy within each individual cell. The crosstalk risk *x*_*crosstalk i*_ for cell *i*:

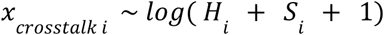

Where *H*_*i*_ and *S*_*i*_ are the summed hSyn and SV40 counts found in cell *i*, respectively. Although crosstalk can be driven by enhancers other than hSyn and SV40, there were multiple enhancer candidates for the same subtype in the experiment; it would be expected that these could have true specificity for the same cells, which would confound estimates of crosstalk between them.

### Regulatory Element Specificity after Spatial Confounder Untanglement (RESSCU)

Given the inherent differences in regulatory element strengths and unclear choice of threshold for deeming a cell as positive or negative for a barcode, modeling barcode counts was preferred over binary classification. For each individual barcode *j*, the total counts of the barcode in cell *i* were modeled as negative-binomially distributed, given that they are discrete counts and based on the observation of overdispersion. The total counts 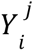 of barcode *j* in cell *i* was represented by the following generalized linear model:

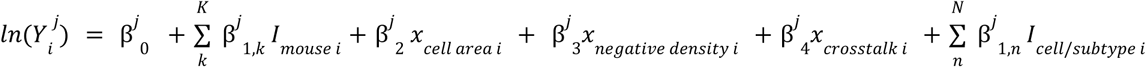

where *ln* is the natural log link function, β are continuous, learned model coefficients, *x*_*negative* *density* *i*_ and *x*_*crosstalk i*_ are continuous covariates and defined in the previous two sections. *I*_*mouse i*_ and *I*_*cell*/*subtype* *i*_ are binary indicator functions such that *I*_*mouse i*_ =1 when cell belongs to the dummy variable for mouse identity *k*\* ∊ {1, 2, 3} and 0 otherwise, *I*_*cell*/*subtype* *i*_ = 1 when cell/subtype *i* belongs to the *n*\* *th* cell type or subtype dummy variable and 0 otherwise. The *x*_*crosstalk*_ term was omitted when fitting models at the major cell type level and only included for estimates of specificity in neuron subtypes. The β^*j*^_1,*n*_* are of ultimate interest, interpreted as the *ln*(counts) of the barcode in a given cell *i* that are attributable toβ that cell’s cell type or subtype identity, having adjusted for the remaining covariates.

β^*j*^_0_ the intercept corresponds to baseline *ln*(counts) when cell size, crosstalk, and negative density are zero and when the dummy variables for mouse identity and cell/subtype corresponding to a chosen reference. Given the observation that microglia (cell type) and Exc-NTS (subtype) have generally lowest counts across barcodes, and mouse 32868 had the lowest expression of the three X1 mice, these were chosen as the reference cell type/subtype and mouse. Mouse identity was modeled as a fixed effect rather than a random effect because of the low number of mice (n=3). We used the R package glmmTMB (https://journal.r-project.org/archive/2017/RJ-2017-066/index.html) to perform maximum likelihood estimation of model coefficients, with the default dispersion model and parameter *family = nbinom2* chosen to parametrize the negative binomial such that the variance scales quadratically with the mean.

Model coefficients and accompanying statistics are provided in **Table S14-S15**, (Sheet 1,2).

### Evaluation of RESSCU models

In this setting of sparse, highly skewed counts data with several sources of experimental and biological variability, predictive power of regression models is limited, and the primary purpose of RESSCU models is statistical inference, especially the relationship of cell and subtype to barcode expression. In order to evaluate the model fit for each barcode, we first binarized cells as being positive or negative for the barcode, and then computed the area under the Receiving Operating Characteristic (auROC) and area under the Precision-Recall Curve (auPRC). We compared these performance metrics to the proportion of positive cells (**Figure S14**). We also performed the same comparison with McFadden’s pseudo R^2^ which compares the log-likelihood of a null model (intercept only) and the full model:

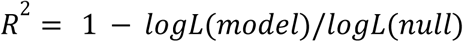

McFadden’s pseudo R^2^ follows a different scale than the coefficient of determination (R^2^); above 0.2 of pseudo R^2^ is considered an “excellent fit”.^95^ Comparison of McFadden’s pseudo R^2^ and proportion of positives for each barcode is shown in (**Figure S14**).

Finally, we assessed for zero inflation, over-dispersion, non-uniformity, and skewness using the DHARMa package (https://github.com/florianhartig/DHARMa). We reported those statistics along with mean absolute error (MAE) in **Table S16-S17, Sheet 3,4**. Overall, models were generally not zero-inflated (p<0.05 for 4% of barcodes), were consistently over-dispersed (p<0.05 for 88% of barcodes), and were mixed in terms of non-uniformity (p<0.05 for 48% of barcodes). Overdispersion and non-uniformity are partially explained by low proportions of positives (mean 0.1) and very high skewness (mean skewness 10.68), consistent with sparse enhancer expression data. MAE modestly improved over the null model, and moreso when limited to positive examples, also typical of sparse, overdispersed data (**Table S16-S17, Sheet 3,4**).

### Calculation of fold specificity for particular cell/subtype and barcode combinations

For a given barcode *j,* the log fold*-*change (FC) specificity for cell/subtype category *n* was calculated separately as an unadjusted measure as well as a RESSCU-adjusted estimate. The unadjusted measure for barcode *j* and cell/subtype *n* was defined as:

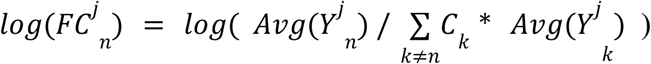

Where 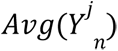 and 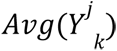 are weighted averages of barcode counts for cell/subtype categories *n* and *k* respectively, and *C*_*k*_ is the total number of cells in category *k*. Thus, the denominator is the weighted average of counts for each cell/subtype *k*≠*n*.

For regression-adjusted estimates, the calculation was similar but the exponentiated coefficients for cell/subtype were used in place of unadjusted average counts:

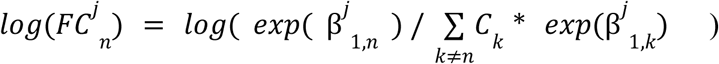

## Acknowledgments

Authors like to thank Jonathan W. Jarvik for the HEK293 cell line and helping with the plasmid transfections into HEK293. We thank Michael Janeček for helpful comments on the manuscript. Also we like to thank Biocognon for providing us the fluorogen MG-ester. One or more of the authors of this paper self-identifies as a member of the LGBTQ+ community. NIH UH3MH120094 (ARP), NIH UF1MH130881 (ARP), NIH R56NS133364 (ARP and RPS), NSF Career 2046550 (ARP), the Cure Alzheimer’s Fund (ARP), F30 DA053020 (BNP), and F31NS134318 (MJL).

