## Supplemental Figures and Tables S3-S13 for "Combining Machine Learning and Multiplexed, *In Situ* Profiling to Engineer Cell Type and Behavioral Specificity"

Supplementals SI

**Figures:**

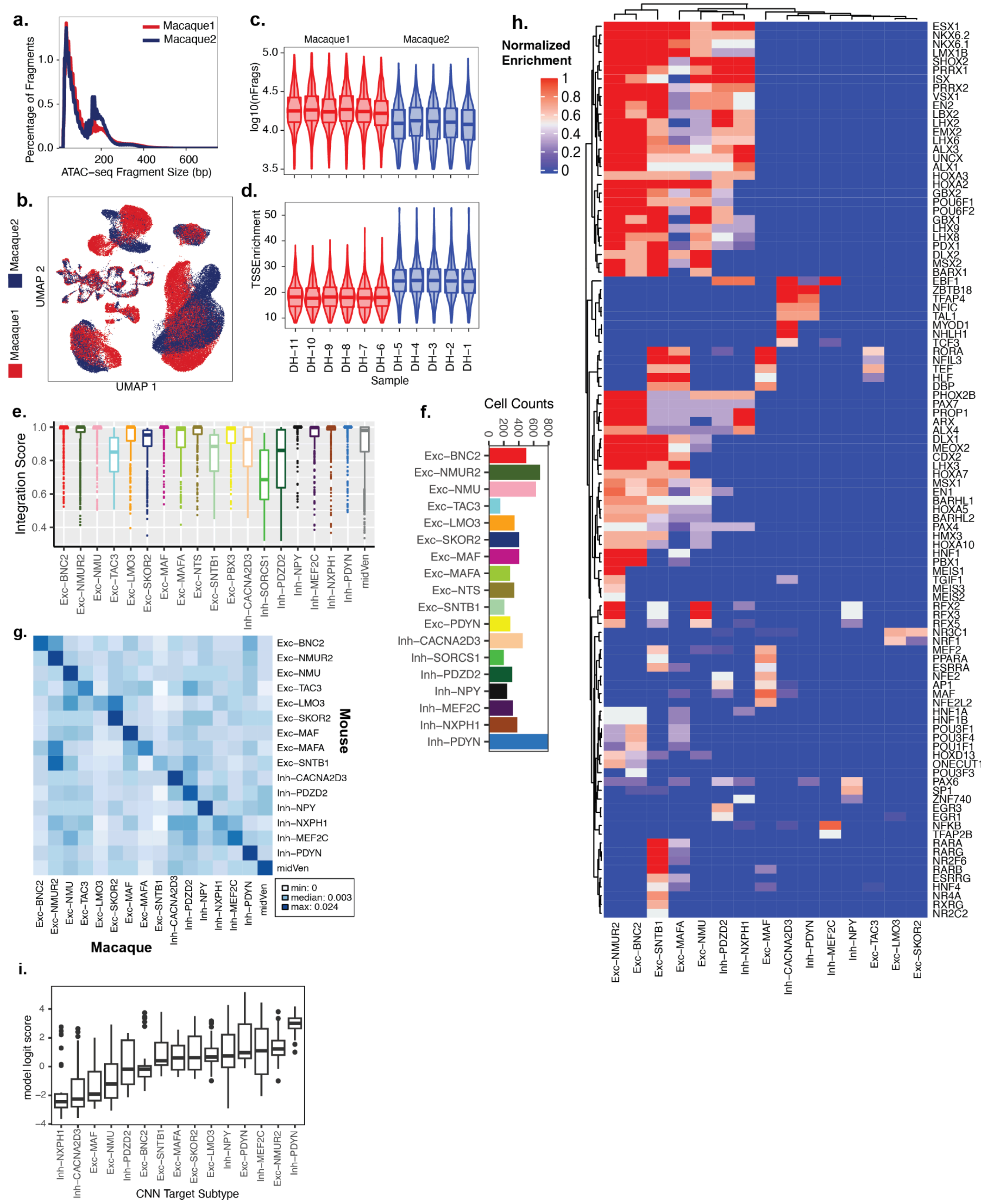

Figure S1. **Establishment of Cross-species Open Chromatin Atlas of the Dorsal Horn.**

a. Fragment size periodicity plots showing depletion at multiples of nucleosome size (~140bp). Color indicates animal.

b. Two-dimensional UMAP representation of processed macaque snATAC-seq stratified by animal.

c. Number of unique fragments (nFrags) by individual technical sample, indicated by the x-axis in **d**. Color indicates animal.

d. Transcription start site (TSS) enrichment by individual technical sample. Color indicates animal.
e. Box plots stratified by neuron subtype of the distribution of confidence scores of cell label following integration with reference macaque snRNA-seq.
f. Bar plot of cell counts for each dorsal neuron subtype.
g. Heatmap of shared peak proportions between neuron subtypes from macaque (x-axis) and mouse (y-axis). Peaks specific to each subtype were individually determined, and the fraction of macaque peaks and mouse orthologs of macaque peaks was calculated for each cross-species subtype pair.
h. Heatmap of specific transcription factors (TFs) and their motif enrichments from chromVar, that were found to be conserved in specificity between macaque snATAC-seq and mouse snATAC (*Arokiaraj et al*) neuron subtypes. Normalized enrichment across subtypes shown.
i. Cross-species SNAIL CNN predictions of Excit-1 enhancer. A higher model logit score (y-axis) represents a higher predicted specificity by the CNN for its target subtype (x-axis). Box plots are distributed over the ensemble of models for each off-target background, and over cross-validation fold models.

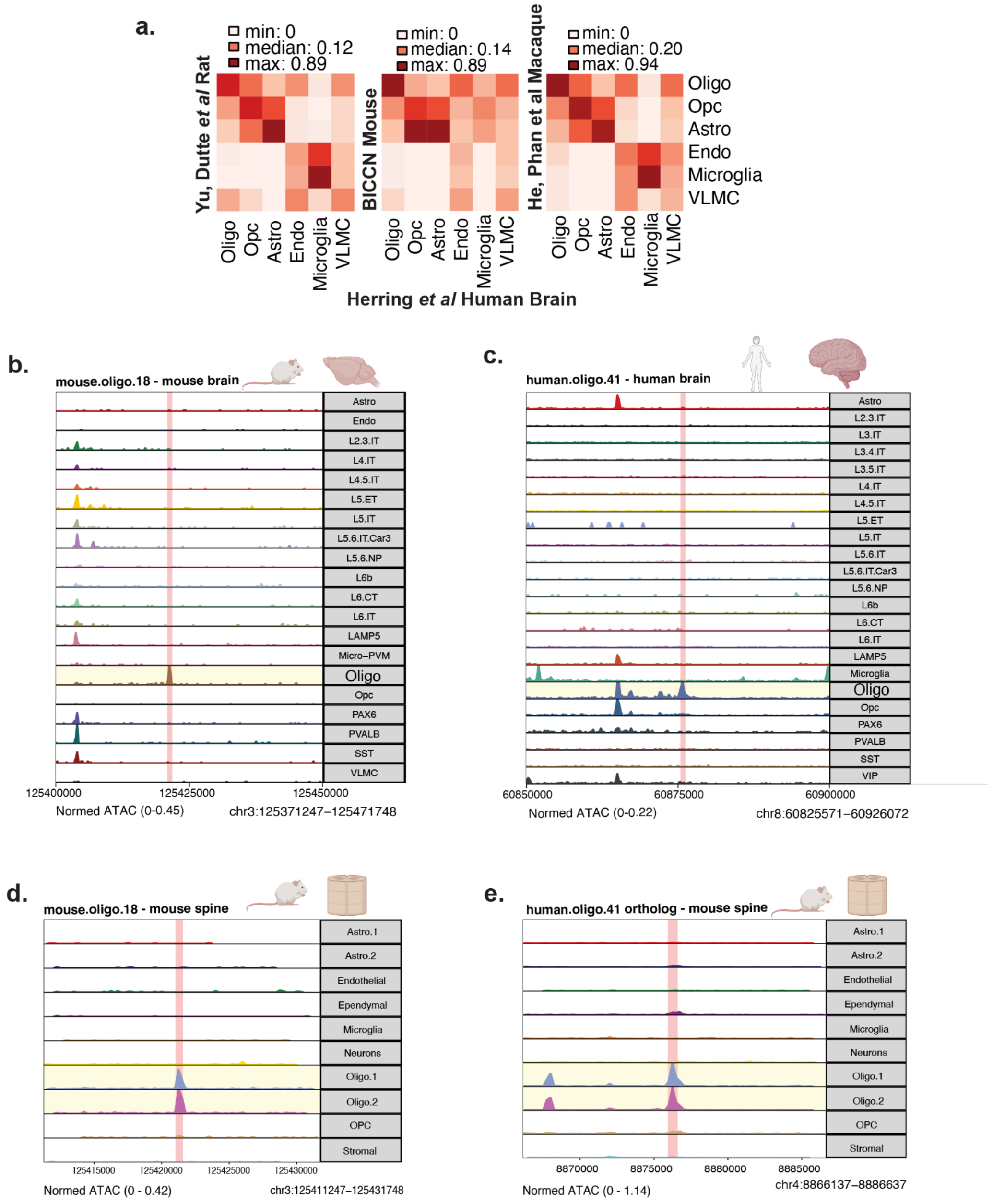

Figure S2. **Oligodendrocyte-targeting Enhancer Candidates**
**a.** Heatmaps of chromVar deviations of major cell types from snATAC-seq of Human brain (x-axis) versus major cell types in Rat brain (Left), Mouse brain (middle), and Macaque brain (right).
**b,c,d,e**. Track plots of 500bp oligodendrocyte-targeting sequences in mouse brain (**b**), human brain (**c**), and mouse spinal cord (**d,e**) (mouse.oligo.18, **b,d** and human.oligo.41, **c,e**). Genomic position (x-axis; coordinates shown at top) is indicated by black squares and red rectangle highlights, with surrounding regions shown on each side. Peak height indicates normalized accessibility for a given cell type (y-axis). In **e**, the accessibility of the mouse ortholog of human.oligo.41 is shown.

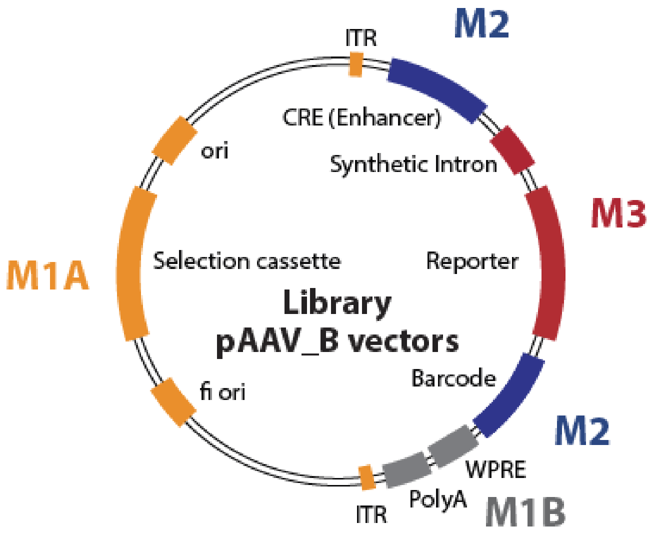


| Module | Vector Components |
|---|---|
| M1A | Selection cassete AMP, *E.coli* ori, f1 ori, ITR |
| M1B | WPRE and PolyA |
| M2 | CRE Enhancer (non-coding region) and BC library |
| M3 | Intron and fluorescent reporters |

Figure S3. **Schematic representation of library non-coding genomic region pAAV final plasmid for SPRA.** The Cis-Regulatory Elements (CREs, enhancers) and barcodes are highlighted in blue; the fluorescent reporter with the intron is outlined in red; selection cassette ampicillin, origin of replications (*E. coli* and f1) and ITR are depicted in orange and post-transcriptional regulatory element (WPRE) and PolyA are shown in grey. M, modules of the Golden Gate assembly pipeline

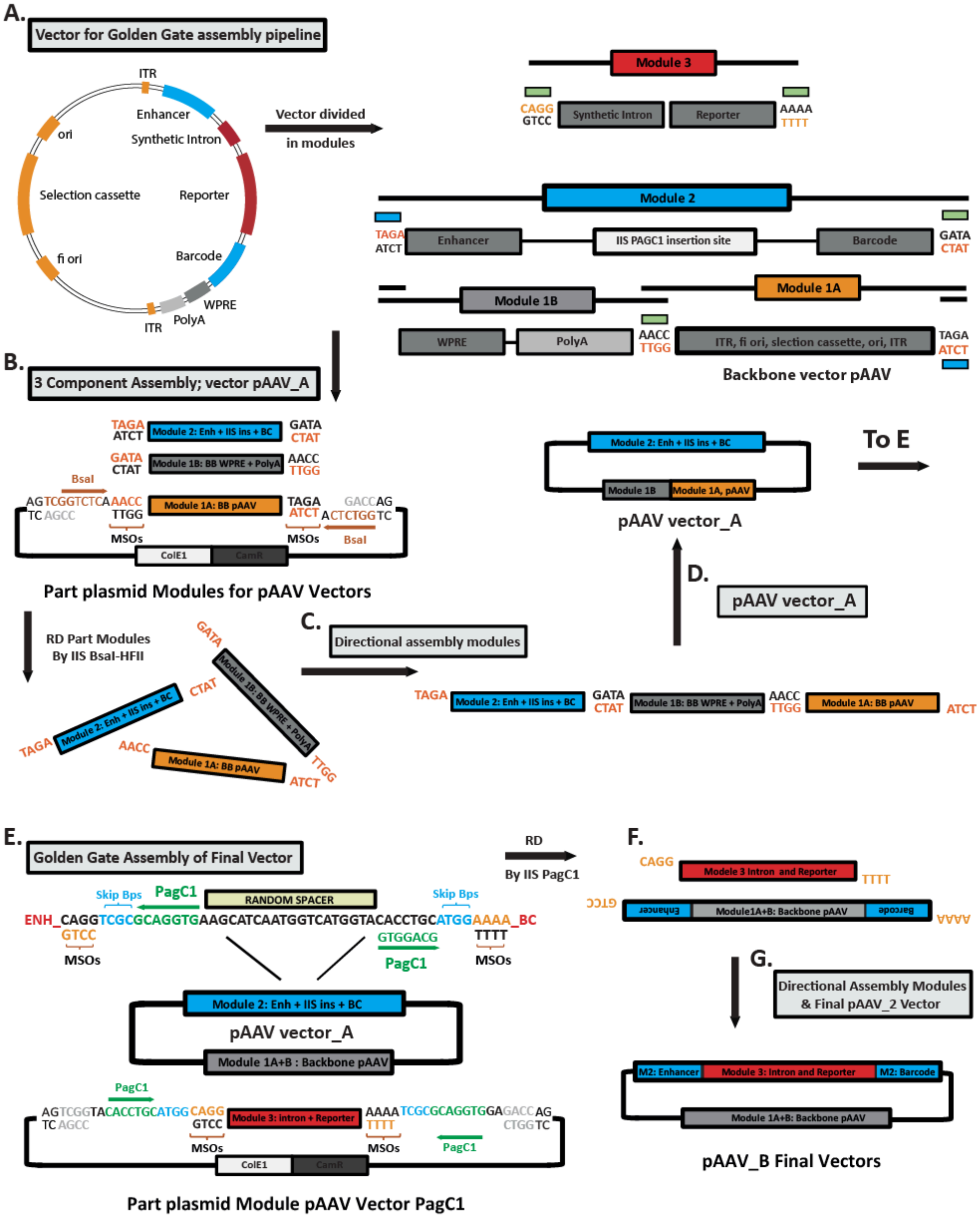


Figure S4. **Design of the SPRA Golden Gate assembly pipeline for studying non-coding genomic regions (enhancers) *in vivo*.** The SPRAGolden gate assembly pipeline is designed for the sequential construction of the ultimate enhancer library vectors through a two-step process. (1) The initial step involves crafting the fundamental library enhancer-barcode vector, enabling the simultaneous synthesis and linkage of both the enhancer and barcode in a single operation. (2) The subsequent step entails integrating the intron and fluorescent reporter to assemble the final library vectors. (a) Step one, the vector for the assembly pipeline is divided into modules (Modules 1A, 1B and 2) and Module 3 is strategically designed as an integration point in module 2 (IIS PaqCI insertion site). The Part-plasmid modules, composed of four components (M1, 1B, 2 and 3), are tailored for executing the Golden Gate reaction. The design of the module specific overhangs MSOs are crucial for high efficiency and directional Golden Gate assembly of the vectors (shown in green and blue) (b) The part-plasmid modules M1A, M1B and M2 are used in a Golden Gate reaction. These modules are released from the part-plasmid through restriction digestion using IIS BsaI-HFv2. (c) The released modules are then directionally assembled on module specific overhangs (MSOs, depicted in orange and black sequences) to generate (d) the initial pAAV_A vector in step 1. (e) For step two, the pAAV_A vector is employed to integrate module 3 (synthetic intron and reporter) into module 2. This integration transpires between the enhancer and barcode of module 2 and is facilitated by the Golden Gate IIS restriction sites PaqC1. Similarly, the final vector pAAV_B is constructed with the pAAV_A and part-plasmid module 3 using Golden Gate assembly with IIS PaqC1 to (f) liberate the modules and (g) directional assembly of the final enhancer-barcode library vectors.

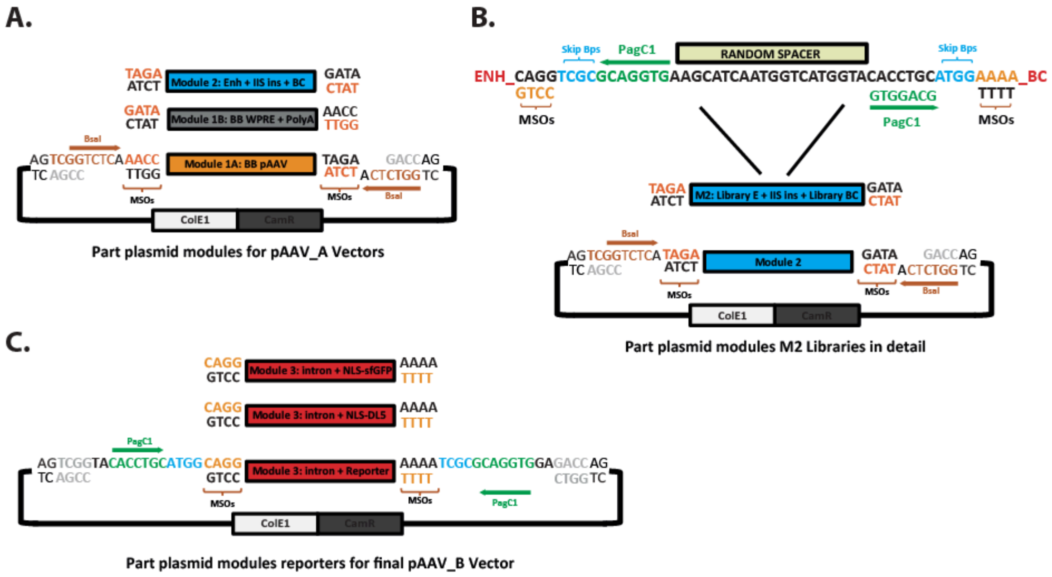


Figure S5. **Part-Plasmid Modules for SPRA**. Overview of the part-plasmid modules used in this study. (a) Schematic of part-plasmids used in constructing the pAAV_A vector, showing the module-specific overhangs (MSOs) and restriction enzyme IIS BsaI-HFv2 sequences. (b) Detailed view of part-plasmid module 2 (libraries), including sequences between the enhancer (E) and barcode (BC) regions. Module 2 contains the drop-in sites for module 3 (fluorescent reporters) and MSOs and restriction enzyme IIS PaqC1 sequences. (c) Schematic of part-plasmid module 3, representing the fluorescent reporters, with MSOs and restriction enzyme IIS PaqC1 sequences, used to construct the final pAAV_B vector.

**Validation of part-plasmid Modules for GG assembly pipeline. Restriction digestion with IIS RE BsaI-HFv2 .**

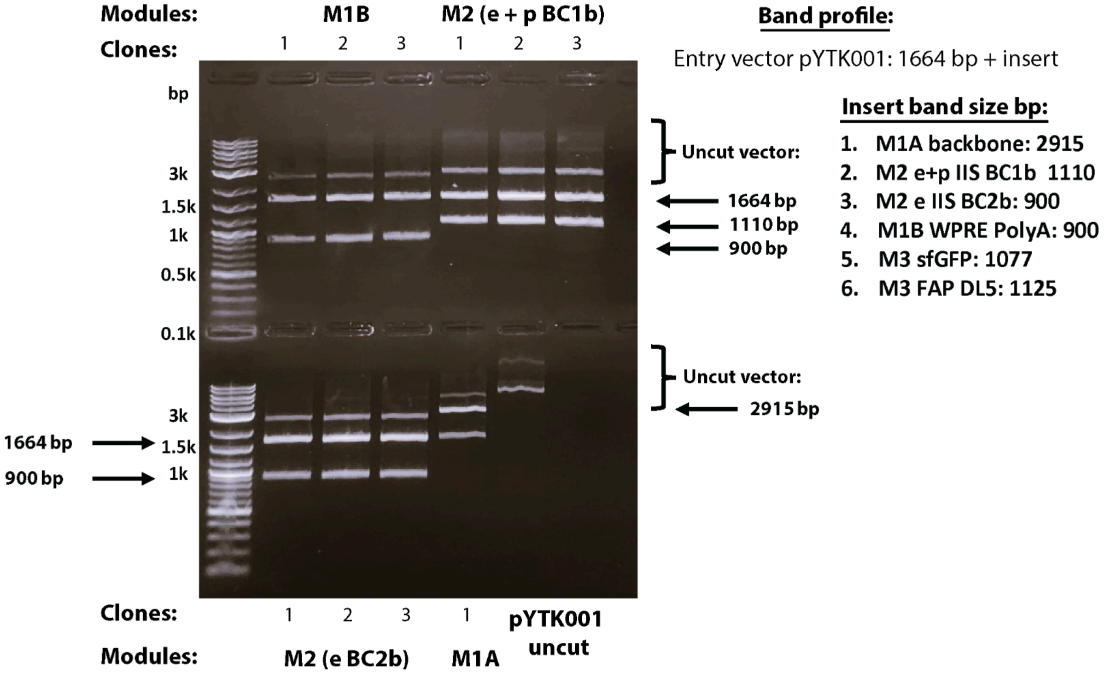


**Validation of part-plasmid Modules M3 Reporters for GG assembly pipeline. Restriction digestion with IIS RE PaqC1-HF .**

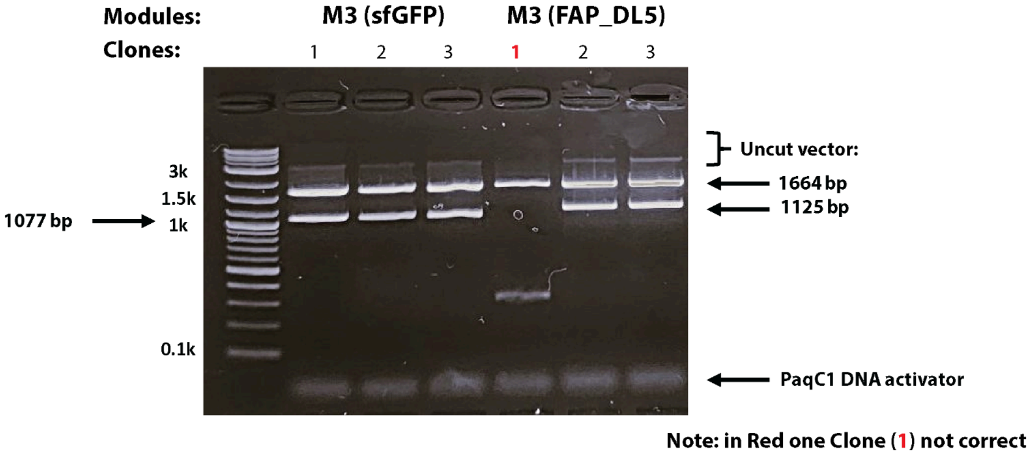


Figure S6. **Validation of part-plasmid modules for the SPRA Golden Gate (GG) assembly platform.** Modules M1A (backbone part of the pAAV vector), M2 (Enhancer, IIS PaqC1 drop-in and Barcodes) and M1B (WPRE and PolyA) are digested with IIS restriction enzyme BsaI-HFv2 to check the correct insert. Modules M3, the reporters with sf-GFP and FAP DL5 are digested with IIS restriction enzyme PaqC1-HF. All modules are confirmed and have the correct insert. Subsequently, all the modules are verified by sanger sequencing. Abbreviations: WPRE: Woodchuck Post-transcriptional Regulatory Element; E: Enhancer; P: Promoter; BC: Barcode; FAP: Fluorogen Activating Protein; RE: Restriction Enzyme.

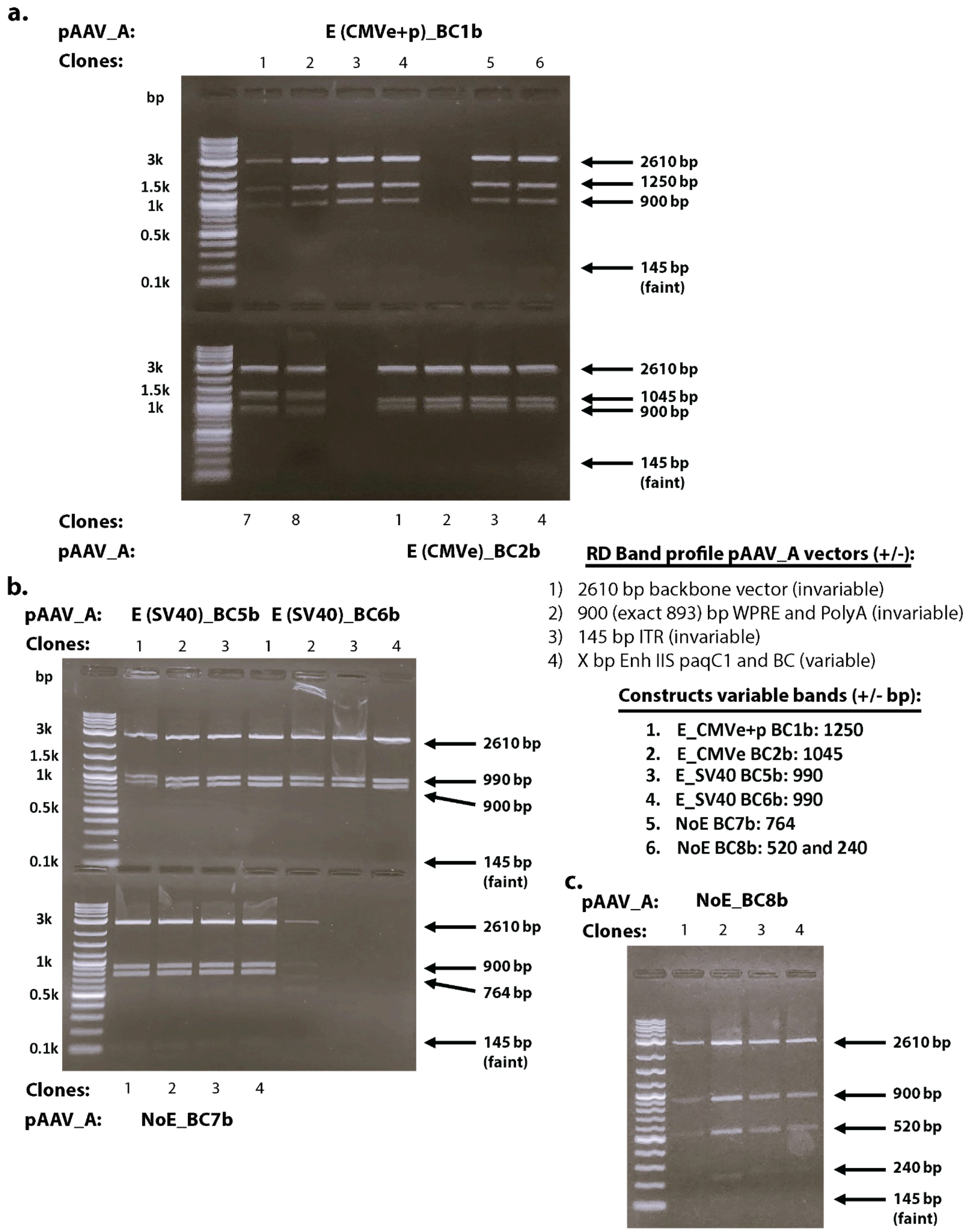


Figure S7. **Initial validation of the SPRA Golden Gate (GG) assembly pipeline for the pAAV_A vectors by restriction digestion profiling with REs HinDIII, NotI and PstI.** The pAAV_A vectors are constructed with part-plasmid modules M1A, M1B and M2 by utilizing the GG assembly pipeline and GG cloning with BsaI-HFv2 and T4 ligase (see methods). The assembled pAAV_A vectors are profiled for a specific cutting pattern by using REs HinDIII, NotI and PstI. All constructed pAAV_A vectors contain invariable RD cutting bands, 2610bp, 900bp and 145bp, corresponding to the backbone pAAV vector, WPRE/PolyA and ITR, respectively. The band profile is variable on the M2 module, the Enhancer and Barcode elements, as these can contain single and multiple RE sides of HinDIII, NotI and PstI. This feature makes it possible to track the performance of the GG assembly pipeline and determine relatively easily, in the early stages of cloning, if the pAAV_A vectors are assembled correctly. Depicted is a panel of the first set of constructed pAAV_A vectors to validate the GG assembly platform. (a) DNA electrophoresis of restriction enzyme digested pAAV_A vectors E (CMVe+p) BC1b and E(CMVe) BC2b where the RD profile matches the expected band profile of all clones tested. (b) DNA electrophoresis of restriction enzyme digested pAAV_A vectors E (SV40) BC5b, E(SV40) BC6b and NoE BC7b where the RD profile matches the expected band profile. (c) DNA electrophoresis of restriction digested pAAV_A vector NoE BC8b and matches the expected band profile. The GG assembly pipeline works efficiently for constructing the pAAV_A vectors. Note: The first panel of pAAV_A vectors are additionally confirmed with whole plasmid sequencing to ensure the correct functioning of the cloning pipeline. Abbreviations: WPRE: Woodchuck Post-transcriptional Regulatory Element; E ,e or Enh: Enhancer; p: Promoter; BC: Barcode; RD restriction digestion; RE: Restriction Enzyme, ITR: Inverted Terminal Repeats: CMV: Cyto-Megalo-Virus; SV40; Simian Virus 40.

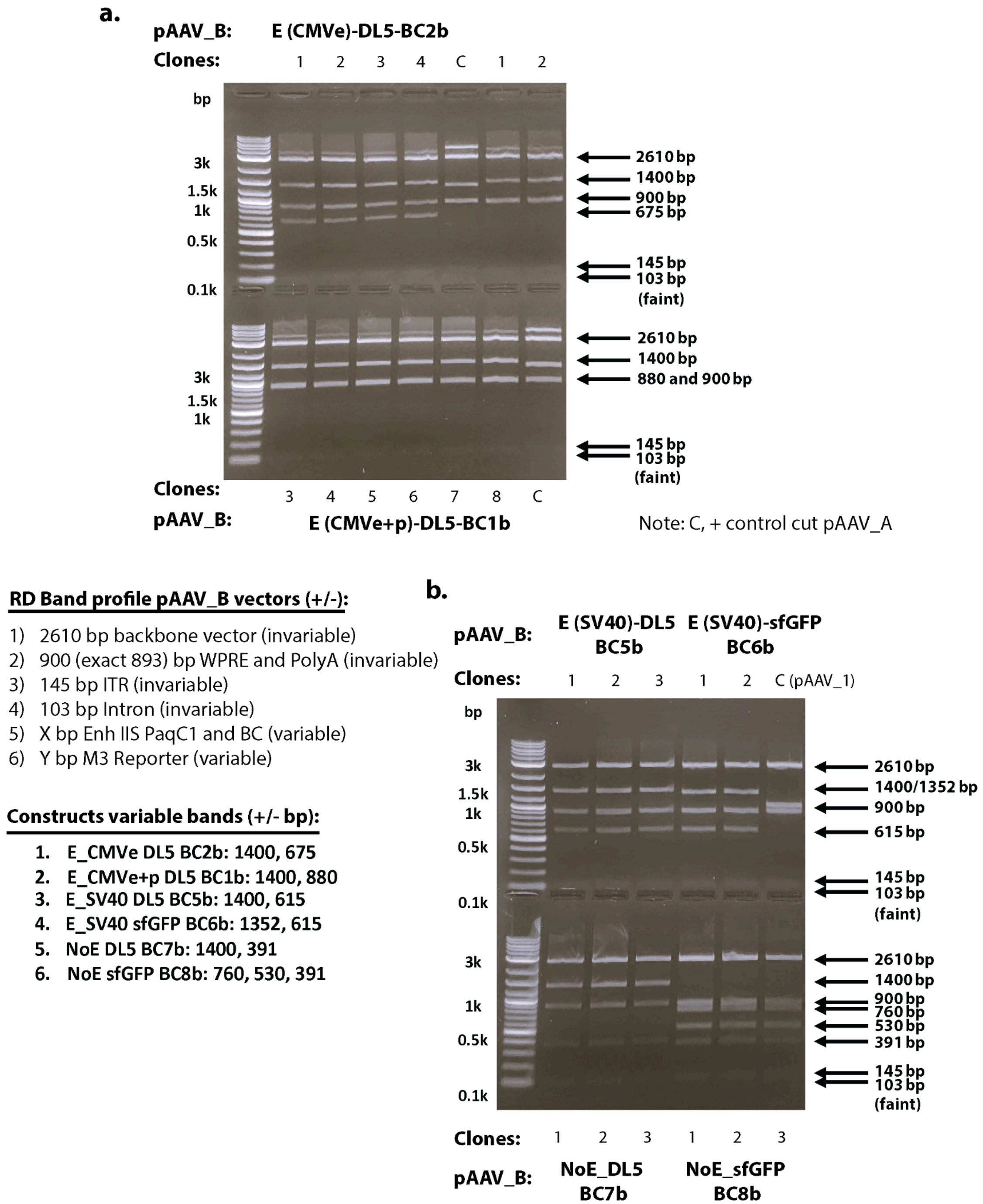


Figure S8. **Initial validation of the SPRA golden Gate (GG) assembly pipeline for the final pAAV_B vectors by restriction digestion profiling with REs HinDIII, NotI and PstI.** Step 2 of the GG assembly platform where the pAAV_B vectors are constructed by using the pAAV_A vectors (first step of the assembly pipeline) and module 3 and utilizing restriction enzyme PaqC1 and T4 ligase (see methods). The assembled pAAV_B vectors are profiled for a specific cutting pattern by using REs HinDIII, NotI and PstI. All constructed pAAV_B vectors contain 4 invariable RD cutting bands, 2610bp, 900bp, 145bp and 103bp corresponding to the backbone pAAV vector, WPRE/PolyA, ITR and part of the Intron, respectively. Similar as by RD profiling of pAAV_A vector (Figure S6), the differences in the RD profile arises from module M2 (Enhancers and barcodes, variable) and the additional module M3 (the reporters, variable). Depicted is a panel of the first set of constructed final pAAV_B vectors using two different reporters FAP DL5 and sfGFP to validate the GG assembly platform. (a) DNA electrophoresis of restriction enzyme digested pAAV_B vectors E(CMVe)-DL5-BC2b and E(CMVe+p)-DL5-BC1b where the RD profile matches the expected band profile in all clones tested. (B) DNA electrophoresis of restriction enzyme digested pAAV_B vectors E (SV40)-DL5-BC5b, E(SV40)-sfGFP-BC6b, NoE-DL5-BC7b and NoE-sfGFP-BC8b where the RD profile matches the expected band profile. Additionally, this panel of vectors were confirmed with whole plasmid sequencing and indicates that the GG assembly platform is highly efficient and operates accordingly. Abbreviations: WPRE: Woodchuck Post-transcriptional Regulatory Element; E: Enhancer; P: Promoter; BC: Barcode; RD restriction digestion; RE: Restriction Enzyme, ITR: Inverted Terminal Repeats; FAP: Fluorogen Activating Protein; GG: Golden Gate; CMV: Cyto-Megalo-Virus; SV40; Simian Virus 40.

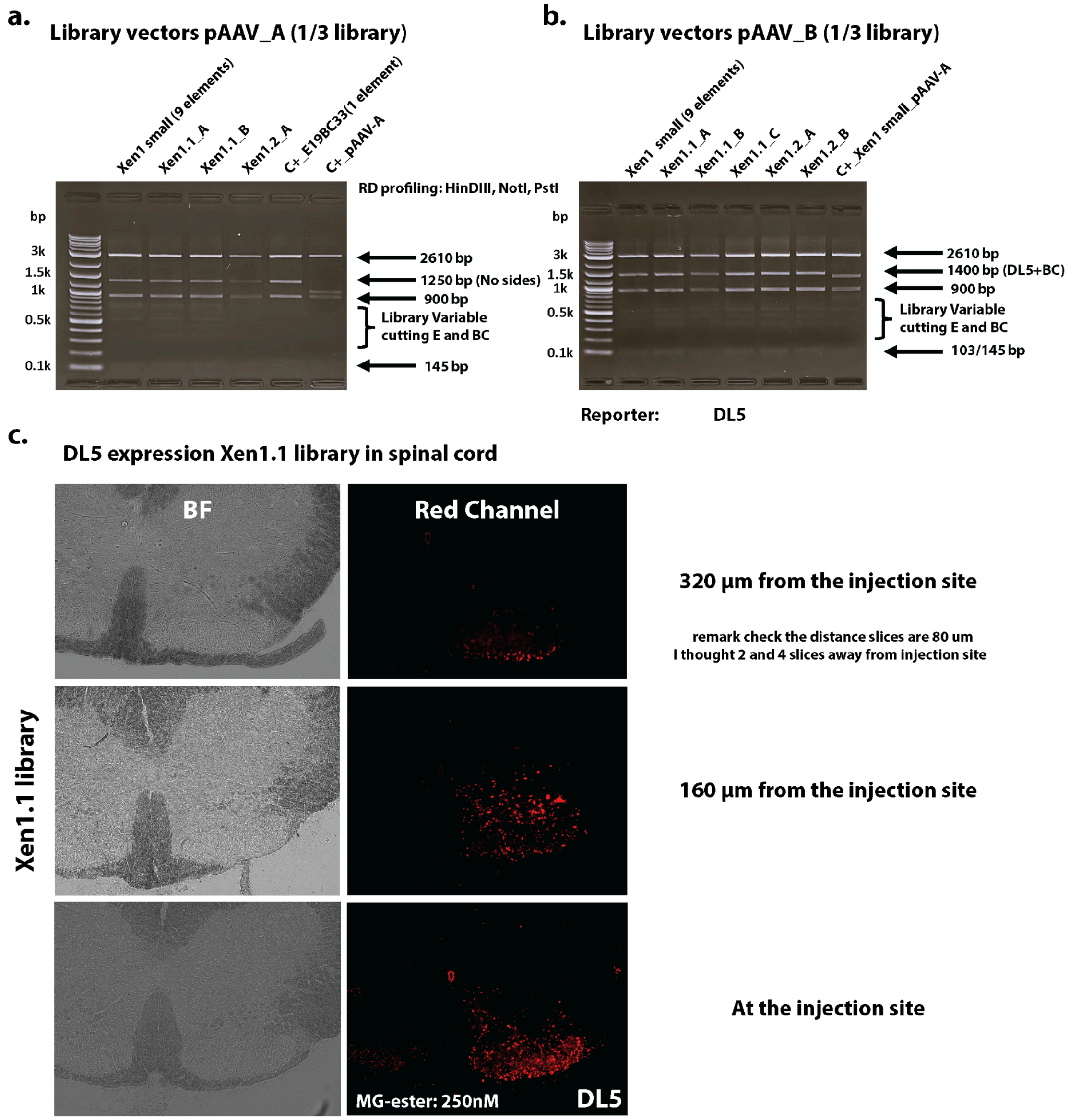


Figure S9. **Validation of the SPRA Golden Gate pipeline for library vectors and reporter expression in Spinal Cord.** The full spinal cord library consists of 27 enhancers paired with 81 unique barcodes, divided into three pools. Each pool (e.g., Xen1.1, Xen1.2, Xen1.3) contains all 27 enhancers combined with a distinct set of 27 barcodes (see Tables S5/7). Library vectors, pAAV_A and pAAV_B were assembled using the SPRAGolden Gate pipeline. Virus containing the Xen1.1 library was injected into mouse spinal cords and analyzed for FAP DL5 expression using a Keyence fluorescent microscope. Restriction digest (RD) profiling was conducted with enzymes HindIII, NotI, and PstI, followed by DNA electrophoresis for validation. (a) Multiple assemblies, including Xen1.1_A, Xen1.1_B, Xen1.2_A, a test library Xen1 small (9 enhancer-barcode elements), and a single enhancer-barcode control (E19BC33), were analyzed. All constructed pAAV_A vectors contain invariable RD cutting bands, 2610bp, 900bp and 145bp, corresponding to the backbone pAAV vector, WPRE/PolyA and ITR, respectively. Library elements lacking enzyme recognition sites for HindIII, NotI, and PstI displayed a 1250 bp band (No Sides). For library elements with one or more RE cutting sites in the enhancer or barcode, the 1250 bp band is fragmented into smaller pieces (library variable cutting E and BC). All pAAV_A vectors have the correct RD band profile. (b) RD-Profiling of 1/3 Library Vectors (pAAV_B with DL5). Similar to figure (a), several Xen1.1 assemblies (Xen1.1_A, Xen1.1_B, Xen1.1_C, Xen1.2_A, Xen1.2_B), the Xen1 small test library, and control E19BC33 with DL5 are shown. All pAAV_B vectors show consistent RD bands at 2610 bp, 900 bp, 145 bp, and 103 bp, corresponding to the backbone, WPRE/PolyA, ITR, and intron, respectively. Library elements without HindIII, NotI, or PstI sites in the barcode display a 1400 bp band (DL5 + BC), while those with RE sites in the barcode or enhancer show smaller variable fragments (library variable cutting E and BC). All pAAV_B vectors with DL5 have the correct RD band profile. (c) Fluorescent imaging of FAP DL5 expression from the Xen1.1 library in the mouse spinal cord. The pAAV_B library vectors Xen1.1_A, Xen1.1_B, and Xen1.1_C were combined to the Xen1.1 library and analyzed for FAP DL5 expression. Robust DL5 fluorescence in the spinal cord is detected and confirms the successful expression of the DL5 reporter by the Xen1.1 library. The FAP DL5 protein is visualized by adding the fluorogen MG-ester at a final concentration of 250 nM. FAP/fluorogen fluorescence was measured at $E_{excitation}$ = 636nm/ $E_{emission}$ = 664nm. Abbreviations: BF: brightfield; WPRE: Woodchuck Post-transcriptional Regulatory Element; E: Enhancer; BC: Barcode; RD restriction digestion; RE: Restriction Enzyme, ITR: Inverted Terminal Repeats; FAP: Fluorogen Activating Protein.

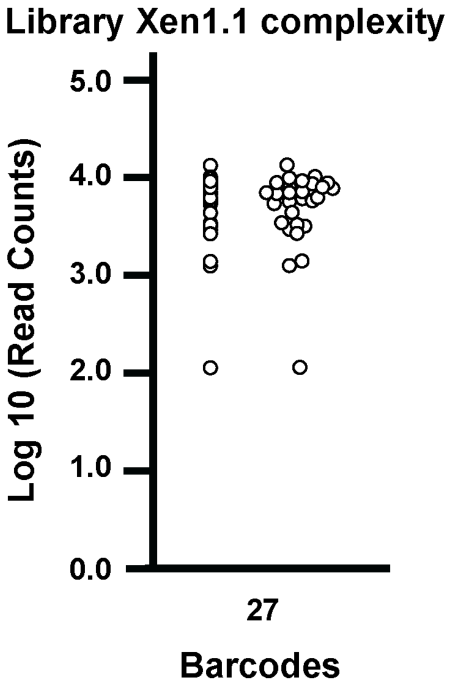


Figure S10. **Library complexity coverage of the 27 barcodes in the (⅓) Xen1.1-Dl5 library.**

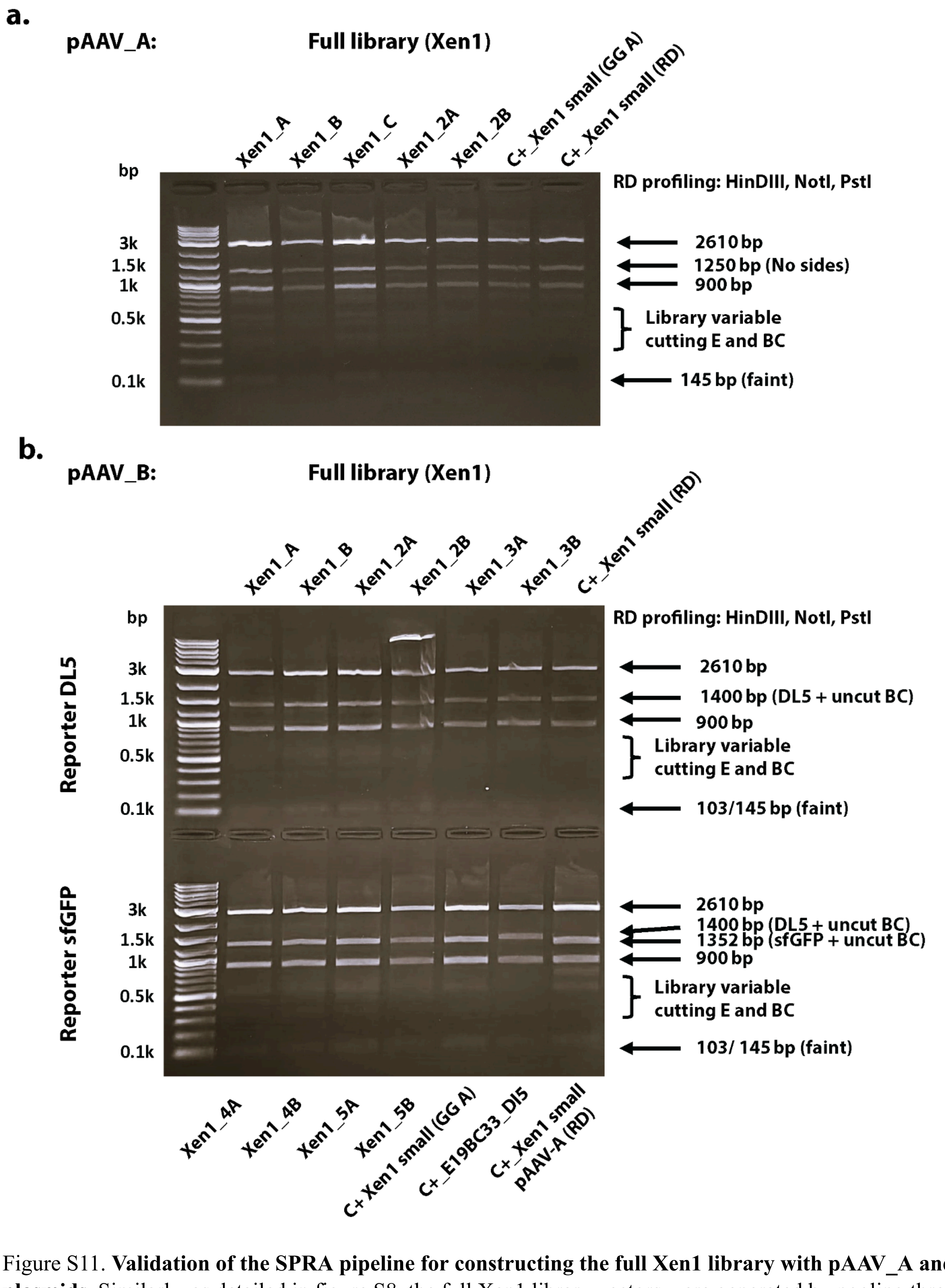


Figure S11. **Validation of the SPRA pipeline for constructing the full Xen1 library with pAAV_A and pAAV_B plasmids.** Similarly, as detailed in figure S8, the full Xen1 library vectors were generated by pooling three 1/3 libraries Xen1.1, Xen1.2, and Xen1.3 (see Tables S5/7, pools 1, 2, and 3) to create a complete library with 27 enhancers and 81 barcodes. RD profiling was performed using restriction enzymes (REs) HinDIII, NotI, and PstI, followed by DNA electrophoresis analysis. (RD profiling of the full pAAV_A library vectors shows multiple Golden Gate (GG) assemblies, including Xen1_A/C and Xen1_2A/B, along with the GG assembly control Xen1 small (GG A) and RD control Xen1 small. The band profile of the full libraries aligns with the expected band sizes. (B) RD profiling of the full pAAV_B library vectors, with reporters DL5 and sfGFP, displays multiple GG assemblies, including Xen1_A/B, Xen1_2A/B, and Xen1_3A/B with DL5, as well as Xen1_4A/B and Xen1_5A/B with sfGFP. It also includes the GG assembly control Xen1 small (GG A), RD control Xen1 small, and the single E/BC control E19-BC33 with DL5. The band profile of the full libraries pAAV_B with DL5 and sfGFP matches the expected band size profile confirming that the SPRA GG cloning pipeline operates efficiently. Abbreviations: E: Enhancer; BC: Barcode; RD restriction digestion; RE: Restriction Enzyme; GG: Golden Gate.

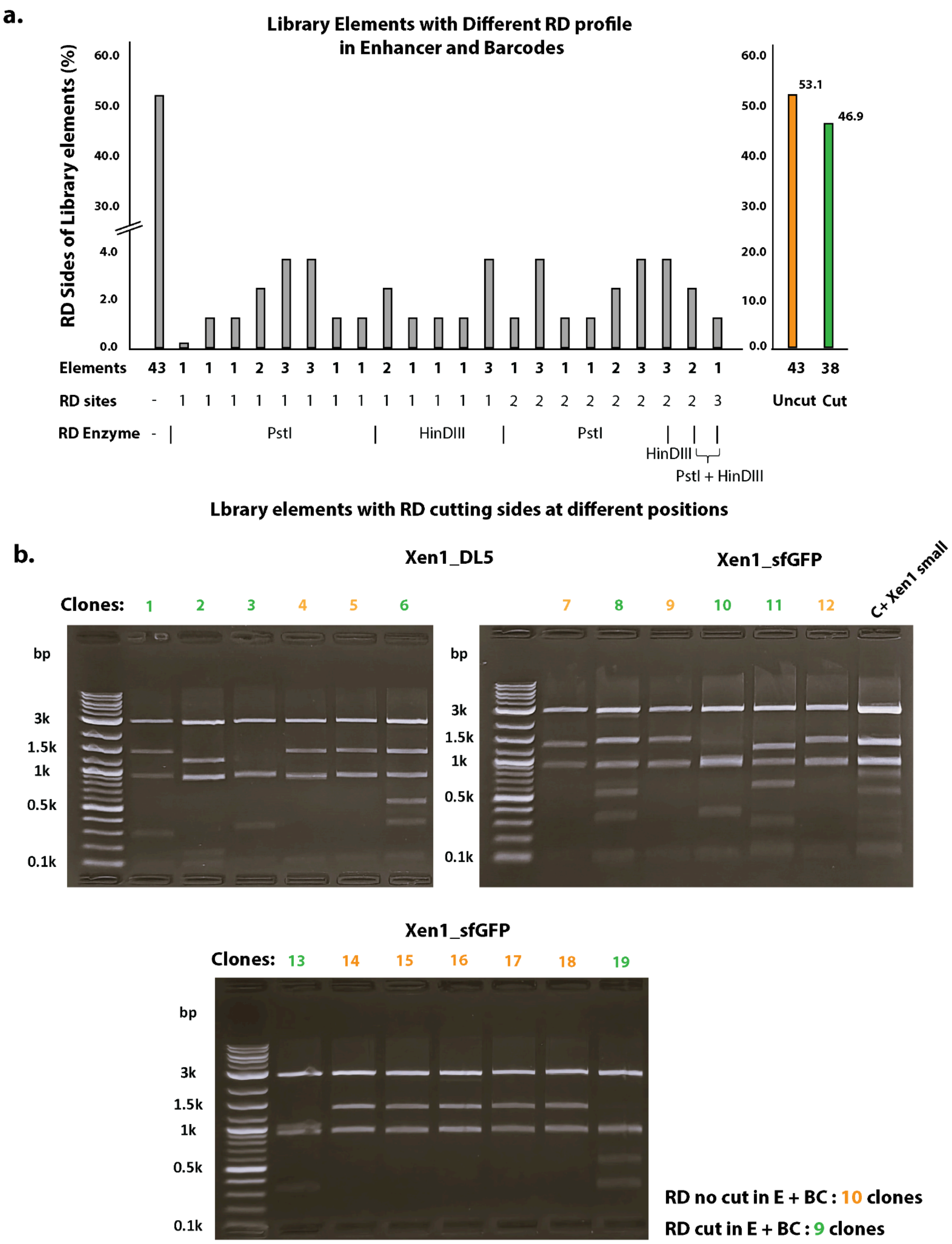


Figure S12. **Preliminary assessment of library complexity conducted through RD profiling of individual library Xen1 elements.** Individual clones from two independent libraries, Xen1_DL5 and Xen1_sfGFP, were randomly selected and RD profiled to assess the complexity of the Xen1 library. The RD profiles of the 81 library elements, which include single or multiple restriction enzyme sites for HinDIII and PstI, facilitate an early and efficient preliminary evaluation of library diversity during the SPRAcloning process. The enhancer and barcode RD profile for the 81 elements in the library contains single or several RD sides with HinDIII, PstI. The ratio determination of expected RD profiles in the Xen1 library is calculated and evaluated (see details Table S8).(a) Bar diagram RD sides of the 81 individual elements in the Xen1 library. The RD sides in the library are represented in ratio of existence (%). RD profiles expected are 43 library elements (orange, 53.1%) contain no RD sides, and 38 (green, 46.9%) library elements contain 1, 2 or 3 RD sides at 31 distinct positions. (b) RD profiling of 19 individual Xen1 library elements using HinDIII, PstI, and NotI revealed that 10 clones (orange) exhibited no RD sites, while 9 clones showed distinct RD patterns. This suggests that the complexity of the 19 tested elements aligns with the expected RD profile of the full library.

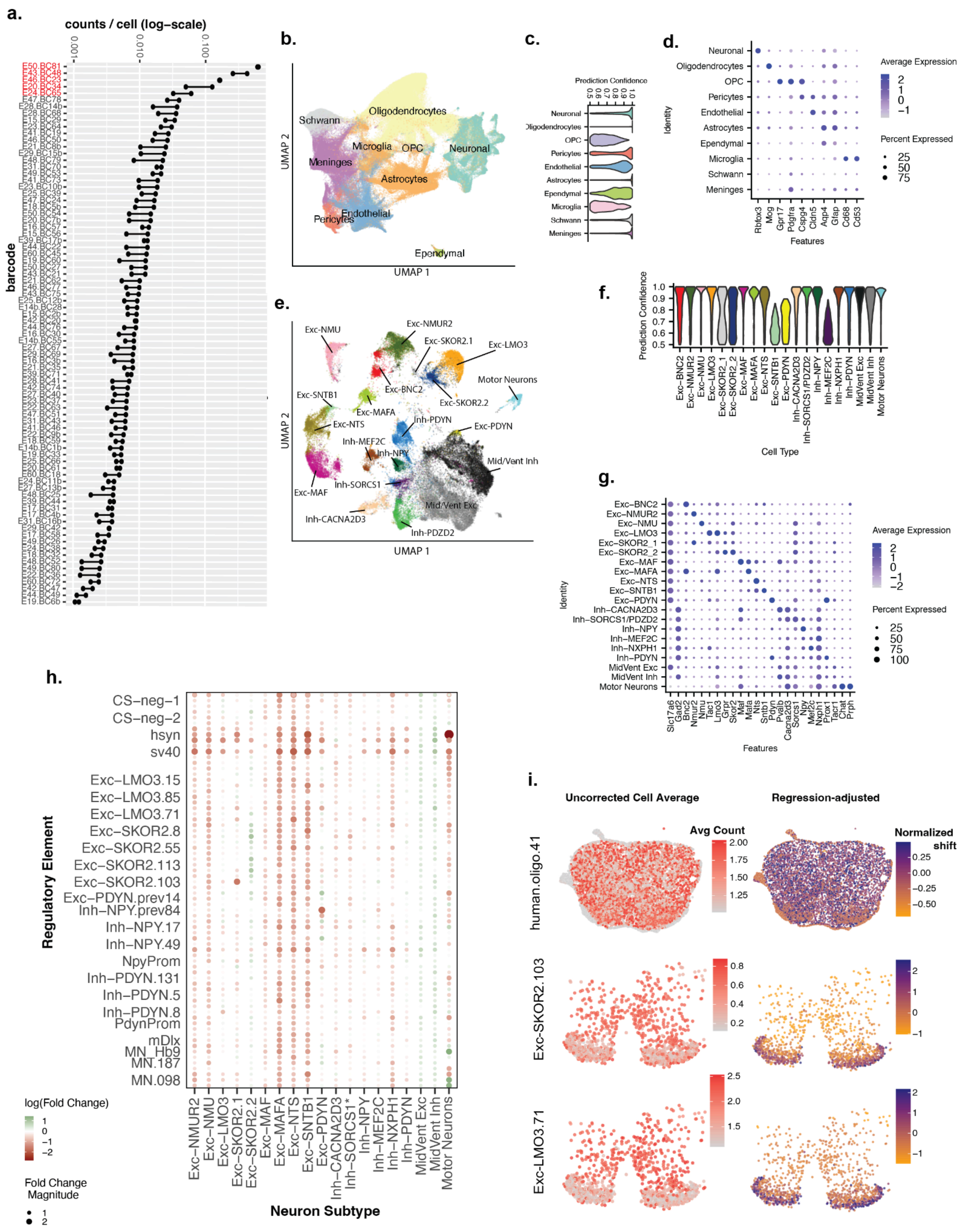
a.
counts / cell (log−scale)
barcode
b.
UMAP 2
UMAP 1
Schwann
Oligodendrocytes
Microglia
OPC
Neuronal
Meninges
Astrocytes
Pericytes
Endothelial
Ependymal
c.
Prediction Confidence
d.
Average Expression
Percent Expressed
Identity
Features
e.
f.
Prediction Confidence
Cell Type
g.
Average Expression
Percent Expressed
Identity
Features
h.
Regulatory Element
Neuron Subtype
log(Fold Change)
Fold Change Magnitude
i.
Uncorrected Cell Average
Regression-adjusted
Avg Count
Normalized shift
human.oligo.41
Exc-SKOR2.103
Exc-LMO3.71

Figure S13. ***In Situ* Single-Cell Analysis of Multiplexed Viral Expression**

a. False-positive detection of barcodes in Xenium assay for n=2 mice that received no viral injections. Counts per-cell (x-axis; log-scale) are shown for individual barcodes (y-axis). Each dot is corresponding to one mouse. Barcodes highlighted in red were removed from subsequent analysis in X1 library due to excessive false-positive detection.
b. Two-dimensional UMAP representation of individual cells labeled by major cell type from X1 (n=3 mice) integrated across mice.
c. Violin plots stratified by X1 major cell type of the distribution of confidence scores of cell label, following integration with reference mouse snRNA-seq.
d. Dot Plot showing expression of individual gene markers (x-axis) for major cell types (y-axis).
e. Two-dimensional UMAP representation of individual cells labeled by neuron subtype from X1 (n=3 mice) integrated across mice.
f. Violin plots stratified by X1 neuron subtype of the distribution of confidence scores of cell label, following integration with reference mouse snRNA-seq.
g. Dot Plot showing expression of individual gene markers (x-axis) for neuron subtypes (y-axis).
h. Unadjusted specificity fold change (FC) results for regulatory elements (y-axis) targeting each neuron subtype (x-axis); Size of dots indicate magnitude of Log(FC). Grey circles with x indicate very low specificity of log(FC) < -1.5. Blue (orange) dot indicates higher (lower) regression-estimate log(FC) of subtype specificity.
i. Cell/subtype average of barcode counts before regression adjustment (Left), and proportional shift of counts attributable to cell/subtype after adjusting for spatial confounders. Example barcodes for human.oligo.41 (Top), Exc-SKOR2.103 (Middle), and Exc-LMO3.71 (Bottom) are shown. Blue (orange) shift indicates greater (less) predicted specificity of the barcode for those cells after regression adjustment.

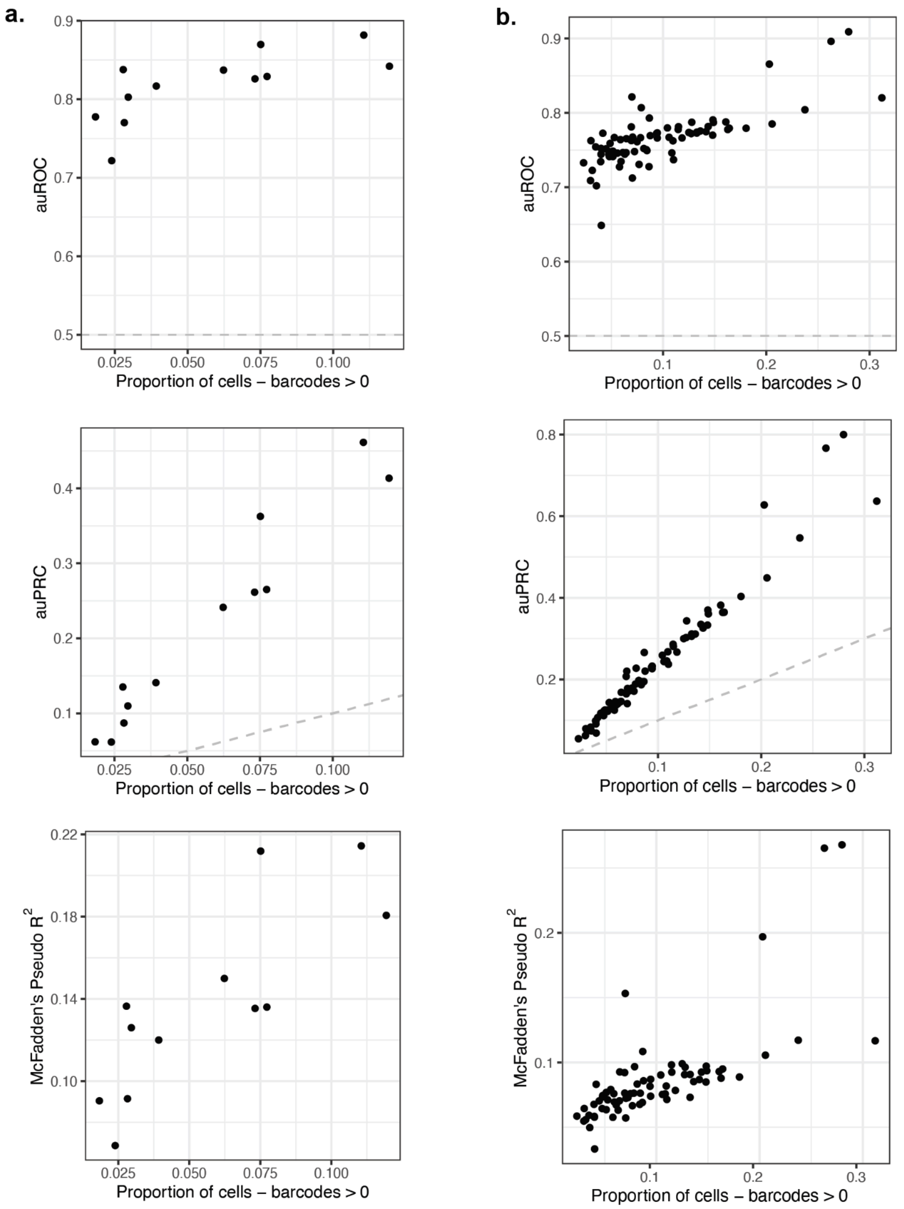


Figure S14. **Evaluation of RESSCU Models.** Scatterplots where each point represents a model for a particular barcode, and position on the x-axis by the proportion of positive cells for that barcode. **a.** Models for cell type, corresponding to Figure 5j in Main Text. **b.** Models for neuron subtype, corresponding to Figure 5i in Main Text. **Top y-axis:** Area under the Receiver Operating Characteristic (auROC) for discrimination of barcode-positive versus negative cells. Dotted gray line represents random chance auROC = 0.5. **Middle y-axis:** Area under the Precision-Recall Curve (auPRC) for discrimination of barcode-positive versus negative cells. Dotted gray identity line represents random chance auPRC = proportion of positive cells. **Bottom y-axis:** McFadden's pseudo R-squared.

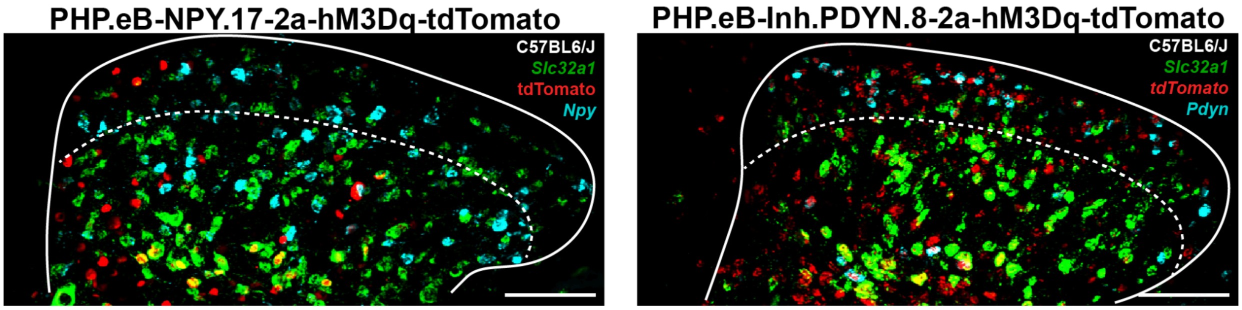


Figure S15. **Fluorescence in situ hybridization (RNAscope) of Inh-NPY.17 and Inh.PDYN.8 candidates.** **a,** Transverse lumbar dorsal horn section showing *Slc32a1* (green), *Npy* (cyan) mRNA transcripts with tdTomato reporter (red) following intraspinal delivery of PHP.eB.NPY.17-hM3Dq-2a-tdTomato. **b,** Transverse lumbar dorsal horn section showing *Slc32a1* (green), *tdTomtao* (red), and *Pdyn* (cyan) mRNA transcripts following intraspinal delivery of PHP.eB.PDYN.8-hM3Dq-2a-tdTomato.Dashed lines delineate lamina II/III border. Scale bars: 100 µm (main images), 30 µm (insets).

**Supplemental Tables:**

**link: TableS1_SNAIL_dataSummary_models_performance**

TableS1. **SNAIL DataSummary Models Performance**

**link: TableS2_modisco_motifs_summary.xlsx**

Table S2**. Modisco Motifs Summary**

| Module (part-plasmid) | MSOs_1 sequences (top/bottom strand) | MSOs_2 sequences (top/bottom strand) | MSOs in Module (start-module-end) | Vector Component |
|---|---|---|---|---|
| Module 1A | AACC/TTGG | TAGA/ATCT | AACC----Module 1A----TAGA | Ampicillin cassette, ori's and ITRs |
| Module 1B | GATA/CTAT | AACC/TTGG | GATA----Module 1B----AACC | WPRE + polyA |
| Module 2 | TAGA/ATCT | GATA/CTAT | TAGA----Module 2-----GATA | Enhnancer/IIS PaqC1/Barcode (or ExBCy library) |
| Module 3 | CAGG/GTCC | AAAA/TTTT | CAGG----Module 3----AAAA | Fluorescent reporters |

Table S3. **Module specific overhangs for the SPRA Golden Gate cloning pipeline**.

| pAAV_A Vector Assembly | Clones Tested | RD Profiling Correct | GG Assembly Efficiency | Different pAAV_A Constructs |
|---|---|---|---|---|
| 1 | 12 | 12 | 100 | 4 |
| 2 | 11 | 11 | 100 | 3 |
| 3 | 4 | 4 | 100 | 1 |
| 4 | 6 | 6 | 100 | 2 |
| 5 | 9 | 9 | 100 | 3 |
| 6 | 2 | 2 | 100 | 1 |
| 7 | 12 | 12 | 100 | 3 |
| Total Clones | 56 | 56 | - | 17 |
| Total Efficiency Pipeline (%) | | | 100 | |
| stdev | | | 0 | |

Table S4. **Golden Gate Assembly Pipeline Efficiency of the pAAV_A Plasmids.**

| pAAV_B Vector Assembly | Clones Tested | RD Profiling Correct | GG Assembly Efficiency | Different pAAV_B Constructs |
|---|---|---|---|---|
| 1 | 12 | 12 | 100.0 | 2 |
| 2 | 12 | 12 | 100.0 | 4 |
| 3 | 12 | 11 | 91.7 | 4 |
| 4 | 12 | 12 | 100.0 | 4 |
| 5 | 10 | 10 | 100.0 | 10 |
| 6 | 1 | 1 | 100.0 | 1 |
| 7 | 20 | 20 | 100.0 | 20 |
| 8 | 18 | 18 | 100.0 | 6 |
| 9 | 12 | 12 | 100.0 | 1 |
| 10 | 6 | 6 | 100.0 | 2 |
| Total Clones | 115 | 114 | - | 54 |
| Total Efficiency Pipeline (%) | | | 99.2 | |
| stdev | | | 2.6 | |

Table S5. **Golden Gate Assembly Pipeline Efficiency of the Final pAAV_B Plasmids.**

| ID | Pool | Nanodrop Spec (ng/µL) | | | Average (ng/µL) | H2O to 50 ng/µL | Final Concentration (ng/µL) | Volume for pool (µL for 100 ng) |
|---|---|---|---|---|---|---|---|---|
| | | 1 | 2 | 3 | | | | |
| E14b.BC1b | Xen1.1 | 69.3 | 69.1 | 68 | 68.80 | 5.19 | 50.00 | 2.00 |
| E15.BC2b | Xen1.1 | 55.3 | 54.9 | 53.7 | 54.63 | 1.28 | 50.00 | 2.00 |
| E16.BC3b | Xen1.1 | 66.3 | 65.9 | 64.4 | 65.53 | 4.29 | 50.00 | 2.00 |
| E17.BC4b | Xen1.1 | 70.7 | 69 | 67.9 | 69.20 | 5.30 | 50.00 | 2.00 |
| E18.BC5b | Xen1.1 | 87.5 | 84.4 | 87.6 | 86.50 | 10.07 | 50.00 | 2.00 |
| E19.BC6b | Xen1.1 | 74.7 | 73.1 | 78.2 | 75.33 | 6.99 | 50.00 | 2.00 |
| E20.BC7b | Xen1.1 | 80.9 | 80 | 78.5 | 79.80 | 8.22 | 50.00 | 2.00 |
| E21.BC8b | Xen1.1 | 86.6 | 86.5 | 84.4 | 85.83 | 9.89 | 50.00 | 2.00 |
| E22.BC9b | Xen1.1 | 46.5 | 45.7 | 45.3 | 45.83 | -1.15 | 45.83 | 2.18 |
| E23.BC10b | Xen1.1 | 67.5 | 65.5 | 65.5 | 66.17 | 4.46 | 50.00 | 2.00 |
| E24.BC11b | Xen1.1 | 61.1 | 61.5 | 60.5 | 61.03 | 3.05 | 50.00 | 2.00 |
| E25.BC12b | Xen1.1 | 65.2 | 63.4 | 62.8 | 63.80 | 3.81 | 50.00 | 2.00 |
| E27.BC13b | Xen1.1 | 50.1 | 49.6 | 49.7 | 49.80 | -0.06 | 49.80 | 2.01 |
| E28.BC14b | Xen1.1 | 79.1 | 78 | 77 | 78.03 | 7.74 | 50.00 | 2.00 |
| E29.BC15b | Xen1.1 | 64.5 | 62.3 | 61.1 | 62.63 | 3.49 | 50.00 | 2.00 |
| E31.BC16b | Xen1.1 | 62.9 | 60.6 | 60.1 | 61.20 | 3.09 | 50.00 | 2.00 |
| E39.BC17b | Xen1.1 | 60.5 | 58 | 58.1 | 58.87 | 2.45 | 50.00 | 2.00 |
| E41.BC19 | Xen1.1 | 62.9 | 59.1 | 59.9 | 60.63 | 2.93 | 50.00 | 2.00 |
| E42.BC20 | Xen1.1 | 10.6 | 10.5 | 10.3 | 10.47 | -10.91 | 10.47 | 9.55 |
| E43.BC21 | Xen1.1 | 10.8 | 10.8 | 11.2 | 10.93 | -10.78 | 10.93 | 9.15 |
| E44.BC22 | Xen1.1 | 74.7 | 73.6 | 72.2 | 73.50 | 6.49 | 50.00 | 2.00 |
| E46.BC23 | Xen1.1 | 17 | 17 | 17.2 | 17.07 | -9.09 | 17.07 | 5.86 |
| E47.BC24 | Xen1.1 | 79.6 | 78.6 | 77.4 | 78.53 | 7.88 | 50.00 | 2.00 |
| E48.BC25 | Xen1.1 | 11.9 | 11.8 | 11.3 | 11.67 | -10.58 | 11.67 | 8.57 |
| E49.BC26 | Xen1.1 | 73.2 | 72.8 | 72.7 | 72.90 | 6.32 | 50.00 | 2.00 |
| E50.BC27 | Xen1.1 | 81.1 | 79.5 | 78.1 | 79.57 | 8.16 | 50.00 | 2.00 |
| E60.BC18 | Xen1.1 | 72.4 | 72.3 | 69.6 | 71.43 | 5.92 | 50.00 | 2.00 |

Table S6. **Recalibration of individual elements of spinal cord library; Pool 1, Xen1.1.** Library of the spinal cord Xen1 is divided into 3 pools Xen1.1, Xen1.2 and Xen1.3. For pool 1, X1.1 the G-blocks of individual elements of the spinal cord library (CREs and Barcodes, $E_xBC_y$) are recalibrated by measuring the concentration of the G-blocks and calculated to 50 ng/µl final concentration. This ensures that every individual element of the library is 50 ng/µl and is subsequently used for constructing the full library with the GG assembly pipeline.

| ID | pool | Nanodrop Spec (ng/µL) | | | Average (ng/µL) | H2O to 50 ng/µL | Final Concentration (ng/µL) | Volume for pool (µL for 100 ng) |
|---|---|---|---|---|---|---|---|---|
| | | 1 | 2 | 3 | | | | |
| E14b.BC28 | Xen1.2 | 86.5 | 87.9 | 85.2 | 86.53 | 10.08 | 50.00 | 2.00 |
| E15.BC29 | Xen1.2 | 72.6 | 71.9 | 69 | 71.17 | 5.84 | 50.00 | 2.00 |
| E16.BC30 | Xen1.2 | 79.7 | 73.2 | 77.4 | 76.77 | 7.39 | 50.00 | 2.00 |
| E17.BC31 | Xen1.2 | 77.6 | 76.5 | 75.7 | 76.60 | 7.34 | 50.00 | 2.00 |
| E18.BC32 | Xen1.2 | 73 | 71.7 | 71.5 | 72.07 | 6.09 | 50.00 | 2.00 |
| E19.BC33 | Xen1.2 | 97.3 | 84.1 | 79.6 | 87.00 | 10.21 | 50.00 | 2.00 |
| E20.BC34 | Xen1.2 | 96.6 | 95.9 | 90.8 | 94.43 | 12.26 | 50.00 | 2.00 |
| E21.BC35 | Xen1.2 | 76.2 | 74.3 | 73.2 | 74.57 | 6.78 | 50.00 | 2.00 |
| E22.BC36 | Xen1.2 | 83.1 | 76.9 | 82.3 | 80.77 | 8.49 | 50.00 | 2.00 |
| E23.BC37 | Xen1.2 | 61.2 | 58.5 | 59 | 59.57 | 2.64 | 50.00 | 2.00 |
| E24.BC38 | Xen1.2 | 70.7 | 69 | 69.1 | 69.60 | 5.41 | 50.00 | 2.00 |
| E25.BC39 | Xen1.2 | 74 | 72.3 | 71.7 | 72.67 | 6.26 | 50.00 | 2.00 |
| E27.BC40 | Xen1.2 | 75.5 | 72.9 | 72.9 | 73.77 | 6.56 | 50.00 | 2.00 |
| E28.BC41 | Xen1.2 | 65.8 | 65.8 | 65.2 | 65.60 | 4.31 | 50.00 | 2.00 |
| E29.BC42 | Xen1.2 | 77.5 | 74.8 | 74.5 | 75.60 | 7.07 | 50.00 | 2.00 |
| E31.BC43 | Xen1.2 | 74.2 | 71.3 | 70.7 | 72.07 | 6.09 | 50.00 | 2.00 |
| E39.BC44 | Xen1.2 | 87.2 | 84.6 | 83.6 | 85.13 | 9.70 | 50.00 | 2.00 |
| E41.BC46 | Xen1.2 | 64.8 | 63.3 | 62.4 | 63.50 | 3.73 | 50.00 | 2.00 |
| E42.BC47 | Xen1.2 | 75.7 | 75.6 | 74.5 | 75.27 | 6.97 | 50.00 | 2.00 |
| E43.BC48 | Xen1.2 | 68.8 | 68.5 | 65.9 | 67.73 | 4.89 | 50.00 | 2.00 |
| E44.BC49 | Xen1.2 | 60.6 | 58.7 | 59.5 | 59.60 | 2.65 | 50.00 | 2.00 |
| E46.BC50 | Xen1.2 | 72.6 | 71.9 | 68.8 | 71.10 | 5.82 | 50.00 | 2.00 |
| E47.BC51 | Xen1.2 | 56.4 | 56.2 | 55.8 | 56.13 | 1.69 | 50.00 | 2.00 |
| E48.BC52 | Xen1.2 | 63.1 | 62.2 | 61.2 | 62.17 | 3.36 | 50.00 | 2.00 |
| E49.BC53 | Xen1.2 | 41.1 | 40.5 | 40.2 | 40.60 | -2.59 | 40.60 | 2.46 |
| E50.BC54 | Xen1.2 | 45.7 | 43.3 | 43.8 | 44.27 | -1.58 | 44.27 | 2.26 |
| E60.BC45 | Xen1.2 | 61.1 | 59.3 | 60.2 | 60.20 | 2.82 | 50.00 | 2.00 |

Table S7. **Recalibration of individual elements of spinal cord library; Pool 2, Xen1.2.** Library of the spinal cord X1 is divided into 3 pools Xen1.1, Xen1.2 and Xen1.3. For pool 2, X1.2 the G-blocks of individual elements of the spinal cord library (CREs and Barcodes, $E_xBC_y$) are recalibrated by measuring the concentration of the G-blocks and calculated to 50 ng/µl final concentration. This ensures that every individual element of the library is 50 ng/µl and is subsequently used for constructing the full library with the GG assembly pipeline.

| ID | pool | Nanodrop Spec (ng/µL) | | | Average (ng/µL) | H2O to 50 ng/µL | Final Concentration (ng/µL) | Volume for pool (µL for 100 ng) |
|---|---|---|---|---|---|---|---|---|
| | | 1 | 2 | 3 | | | | |
| E14b.BC55 | Xen1.3 | 46.8 | 45.1 | 45.1 | 45.67 | -1.20 | 45.67 | 2.19 |
| E15.BC56 | Xen1.3 | 43.6 | 43.8 | 42.4 | 43.27 | -1.86 | 43.27 | 2.31 |
| E16.BC57 | Xen1.3 | 56.2 | 55.6 | 54.8 | 55.53 | 1.53 | 50.00 | 2.00 |
| E17.BC58 | Xen1.3 | 45.5 | 43.6 | 43.2 | 44.10 | -1.63 | 44.10 | 2.27 |
| E18.BC59 | Xen1.3 | 49.6 | 48.3 | 48.1 | 48.67 | -0.37 | 48.67 | 2.05 |
| E19.BC60 | Xen1.3 | 86.5 | 85.1 | 84.3 | 85.30 | 9.74 | 50.00 | 2.00 |
| E20.BC61 | Xen1.3 | 53.5 | 51.2 | 52.4 | 52.37 | 0.65 | 50.00 | 2.00 |
| E21.BC62 | Xen1.3 | 50.6 | 48.6 | 49.3 | 49.50 | -0.14 | 49.50 | 2.02 |
| E22.BC63 | Xen1.3 | 51.5 | 49.8 | 49.6 | 50.30 | 0.08 | 50.30 | 1.99 |
| E23.BC64 | Xen1.3 | 47.9 | 46.8 | 47 | 47.23 | -0.76 | 47.23 | 2.12 |
| E24.BC65 | Xen1.3 | 42.3 | 41.1 | 40.9 | 41.43 | -2.36 | 41.43 | 2.41 |
| E25.BC66 | Xen1.3 | 54.4 | 52.6 | 54 | 53.67 | 1.01 | 50.00 | 2.00 |
| E27.BC67 | Xen1.3 | 70.1 | 71.1 | 70.5 | 70.57 | 5.68 | 50.00 | 2.00 |
| E28.BC68 | Xen1.3 | 58.6 | 57.2 | 56.8 | 57.53 | 2.08 | 50.00 | 2.00 |
| E29.BC69 | Xen1.3 | 52.4 | 51 | 51 | 51.47 | 0.40 | 50.00 | 2.00 |
| E31.BC70 | Xen1.3 | 46.9 | 45.2 | 44.1 | 45.40 | -1.27 | 45.40 | 2.20 |
| E39.BC71 | Xen1.3 | 60.4 | 58.6 | 58.9 | 59.30 | 2.57 | 50.00 | 2.00 |
| E41.BC73 | Xen1.3 | 42.8 | 41.6 | 42.1 | 42.17 | -2.16 | 42.17 | 2.37 |
| E42.BC74 | Xen1.3 | 31.2 | 32.6 | 32.7 | 32.17 | -4.92 | 32.17 | 3.11 |
| E43.BC75 | Xen1.3 | 63.3 | 61.8 | 60.8 | 61.97 | 3.30 | 50.00 | 2.00 |
| E44.BC76 | Xen1.3 | 46.2 | 45 | 45 | 45.40 | -1.27 | 45.40 | 2.20 |
| E46.BC77 | Xen1.3 | 73 | 71.9 | 73.5 | 72.80 | 6.29 | 50.00 | 2.00 |
| E47.BC78 | Xen1.3 | 49.7 | 49.2 | 47.7 | 48.87 | -0.31 | 48.87 | 2.05 |
| E48.BC79 | Xen1.3 | 53.8 | 55.1 | 54.8 | 54.57 | 1.26 | 50.00 | 2.00 |
| E49.BC80 | Xen1.3 | 43.1 | 42.3 | 41.1 | 42.17 | -2.16 | 42.17 | 2.37 |
| E50.BC81 | Xen1.3 | 45.4 | 45.4 | 44.4 | 45.07 | -1.36 | 45.07 | 2.22 |
| E60.BC72 | Xen1.3 | 74.3 | 73.7 | 72.4 | 73.47 | 6.48 | 50.00 | 2.00 |

Table S8. **Recalibration of individual elements of spinal cord library; Pool 3, Xen1.3.** Library of the spinal cord X1 is divided into 3 pools Xen1.1, Xen1.2 and Xen1.3. For pool 3, X1.1 the G-blocks of individual elements of the spinal cord library (CREs and Barcodes, $E_xBC_y$) are recalibrated by measuring the concentration of the G-blocks and calculated to 50 ng/µl final concentration. This ensures that every individual element of the library is 50 ng/µl and is subsequently used for constructing the full library with the GG assembly pipeline.

| Different cut sides $E_{27}BC_{81}$ | RD cut sides in $E_{27}BC_{81}$ | Enzyme | Position | Number Library Elements | Ratio | RD cut sides representation (%) | RD uncut/cut (%) | Expected clones uncut/cut (19 clones) |
|---|---|---|---|---|---|---|---|---|
| 1 | No cut | N.A. | N.A. | 43 | 0.5309 | 53.1 | 53.1 | 10 |
| 2 | I cut | PstI | 1 | 1 | 0.0123 | 0.12 | 46.9 | 9 |
| 3 | I cut | | 2 | 1 | 0.0123 | 1.23 | | |
| 4 | I cut | | 3 | 1 | 0.0123 | 1.23 | | |
| 5 | I cut | | 4 | 2 | 0.0247 | 2.47 | | |
| 6 | I cut | | 5 | 3 | 0.0370 | 3.70 | | |
| 7 | I cut | | 6 | 3 | 0.0370 | 3.70 | | |
| 8 | I cut | | 7 | 1 | 0.0123 | 1.23 | | |
| 9 | I cut | | 8 | 1 | 0.0123 | 1.23 | | |
| 10 | I cut | HinDIII | 9 | 2 | 0.0247 | 2.47 | | |
| 11 | I cut | | 10 | 1 | 0.0123 | 1.23 | | |
| 12 | I cut | | 11 | 1 | 0.0123 | 1.23 | | |
| 13 | I cut | | 12 | 1 | 0.0123 | 1.23 | | |
| 14 | I cut | | 13 | 3 | 0.0370 | 3.70 | | |
| 15 | I cut | | 14 | 1 | 0.0123 | 1.23 | | |
| 16 | 2 cuts | PstI | 15+16 | 3 | 0.0370 | 3.70 | | |
| 17 | 2 cuts | | 17+18 | 1 | 0.0123 | 1.23 | | |
| 18 | 2 cuts | | 19+20 | 1 | 0.0123 | 1.23 | | |
| 19 | 2 cuts | | 21+22 | 2 | 0.0247 | 2.47 | | |
| 20 | 2 cuts | | 23+24 | 3 | 0.0370 | 3.70 | | |
| 21 | 2 cuts | HinDIII | 25+26 | 3 | 0.0370 | 3.70 | | |
| 22 | 2 cuts | PstI and HinDIII | 27+28 | 2 | 0.0247 | 2.47 | | |
| 23 | 3 cuts | PstI and HinDIII | 29+30+31 | 1 | 0.0123 | 1.23 | | |
| | | | **Total elements library:** | **81** | | | | |

Table S9. **Library Xen1 complexity ratios and clones exhibiting RD profiles with or without cutting, as expected in the RD profiling test.**

| Name | Vector ESCARGOT pipeline | Description |
|---|---|---|
| pYTK001 | part-plasmid | Golden Gate assembly entry vector (moclo kit) |
| pYTK001-M1A-ITR2/AmpR/ori | Module M1A part-plasmid | pAAV backbone (Includes: 5' and 3' AAV ITRs (Inverted Terminal Repeats), ampicillin resistence gene, and Ori (origin of replication)) |
| pYTK001-M1B-WPRE/polyA | Module M1B part-plasmid | WPRE (Woodchuck Post-transcriptional Regulatory Element) and PolyA |
| pYTK001-M2-CMVe+p/BC1b | Module M2 part-plasmid | CMV (Cyto-Megalo-Virus) enhancer plus promoter sequence paired with DNA barcode #1 |
| pYTK001-M2-CMVe/BC2b | Module M2 part-plasmid | CMV enhancer sequence paired with DNA barcode #2 |
| pYTK001-M3-intron-GFP | Module M3 part-plasmid | Clontech intron/nuclear localized GFP (Green Fluorescent Protein) reporter |
| pYTK001-M3-intron-DL5 | Module M3 part-plasmid | Clontech intron/nuclear localized DL5 reporter |
| pYTK001-M2-SV40/BC5b | Module M2 part-plasmid | SV40 (Simian Virus 40) enhancer sequence paired with DNA barcode #5 |
| pYTK001-M2-SV40/BC6b | Module M2 part-plasmid | SV40 enhancer sequence paired with DNA barcode #6 |
| pYTK001-M2-NoE/BC7b | Module M2 part-plasmid | Module M2 with no enhancer sequence paired with DNA barcode #7 |
| pYTK001-M2-NoE/BC8b | Module M2 part-plasmid | Module M2 with no enhancer sequence paired with DNA barcode #8 |
| pAAV-CMVe+p-BC1b | pAAV_A | pAAV backbone plasmid with CMV enhancer plus promoter sequence-DNA barcode #1 |
| pAAV-CMVe-BC2b | pAAV_A | pAAV backbone plasmid with CMV enhancer sequence-DNA barcode #2 |
| pAAV-CMVe+p-GFP-BC1b | pAAV_B | pAAV backbone plasmid with CMV enhancer plus promoter sequence-Clontech intron/nuclear localized GFP reporter-DNA barcode #1 |
| pAAV-CMVe+p-DL5-BC1b | pAAV_B | pAAV backbone plasmid with CMV enhancer sequence-nuclear localized DL5-DNA barcode #1 |
| pAAV-CMVe-GFP-BC2b | pAAV_B | pAAV backbone plasmid with CMV enhancer sequence-Clontech intron/nuclear localized GFP reporter-DNA barcode #2 |
| pAAV-CMVe-DL5-BC2b | pAAV_B | pAAV backbone plasmid with CMV enhancer sequence-Clontech intron/nuclear localized DL5 reporter-DNA barcode #2 |
| pAAV-SV40-BC5b | pAAV_A | pAAV backbone plasmid with SV40 enhancer sequence-DNA barcode #5 |
| pAAV-SV40-BC6b | pAAV_A | pAAV backbone plasmid with SV40 enhancer sequence-DNA barcode #6 |
| pAAV-NoE-BC7b | pAAV_A | pAAV backbone plasmid with no enhancer sequence-DNA barcode #7 |
| pAAV-NoE-BC8b | pAAV_A | pAAV backbone plasmid with no enhancer sequence-DNA barcode #8 |
| pAAV-SV40-DL5-BC5b | pAAV_B | pAAV backbone plasmid with SV40 enhancer sequence-Clontech intron/nuclear localized DL5 reporter-DNA barcode #5 |
| pAAV-SV40-GFP-BC6b | pAAV_B | pAAV backbone plasmid with SV40 enhancer sequence-Clontech intron/nuclear localized GFP reporter-DNA barcode #6 |
| pAAV-NoE-DL5-BC7b | pAAV_B | pAAV backbone plasmid with no enhancer sequence-Clontech intron/nuclear localized DL5 reporter-DNA barcode #7 |
| pAAV-NoE-GFP-BC8b | pAAV_B | pAAV backbone plasmid with no enhancer sequence-Clontech intron/nuclear localized GFP reporter-DNA barcode #8 |
| pAAV-Xen1-small | pAAV_A | pAAV backbone plasmid with 9 M2 enhancer/DNA barcode pairs |
| pAAV-Xen1.1 | pAAV_A | pAAV backbone plasmid with Xen1.1 library: each of the 27 enhancers, paired with one set of DNA barcodes (#1-27) |
| pAAV-Xen1.2 | pAAV_A | pAAV backbone plasmid with Xen1.2 library: each of the 27 enhancers, paired with one set of DNA barcodes (#28-54) |
| pAAV-Xen1.3 | pAAV_A | pAAV backbone plasmid with Xen1.1 library: each of the 27 enhancers, paired with one set of DNA barcodes (#55-81) |
| pAAV-E19-BC33 | pAAV_A | pAAV backbone plasmid with enhancer #19 (GABA3.13)-DNA barcode #33 |
| pAAV-Xen1-small-DL5 | pAAV_B | pAAV backbone plasmid with Clontech intron/nuclear localized DL5 and 9 M2 enhancer/DNA barcode pairs |
| pAAV-Xen1-small-GFP | pAAV_B | pAAV backbone plasmid with Clontech intron/nuclear localized GFP and 9 M2 enhancer/DNA barcode pairs |
| pAAV-Xen1.1-DL5 | pAAV_B | pAAV backbone plasmid with Clontech intron/nuclear localized DL5 and Xen1.1 library: each of the 27 enhancers, paired with one set of DNA barcodes (#1-27) |
| pAAV-Xen1.2-DL5 | pAAV_B | pAAV backbone plasmid with Clontech intron/nuclear localized DL5 and Xen1.2 library: each of the 27 enhancers, paired with one set of DNA barcodes (#28-54) |
| pAAV-E19-DL5-BC33 | pAAV_B | pAAV backbone plasmid with enhancer #19 (GABA3.13)-Clontech intron/nuclear localized DL5-DNA barcode #33 |
| pAAV-Xen1 | pAAV_A | pAAV backbone plasmid with Xen1 library: each of the 27 enhancers, each paired with 3 DNA barcodes (#1-81) |
| pAAV-Xen1-DL5 | pAAV_B | pAAV backbone plasmid with Clontech intron/nuclear localized DL5 and Xen1 library: each of the 27 enhancers, each paired with 3 DNA barcodes (#1-81) |
| pAAV-Xen1-GFP | pAAV_B | pAAV backbone plasmid with Clontech intron/nuclear localized GFP and Xen1 library: each of the 27 enhancers, each paired with 3 DNA barcodes (#1-81) |

Table S10 **Plasmids that are constructed for this study.** Abbreviations: WPRE: Woodchuck Post-transcriptional Regulatory Element; P: promotor, E: Enhancer; BC: Barcode; ITR: Inverted Terminal Repeats; Amp: Ampicillin; Ori; origin of replication; NLS: Nuclear localization signal; GG: Golden Gate; CMV: Cyto-Megalo-Virus; SV40; Simian Virus 40.

| ID | Enhancer | Barcode |
|---|---|---|
| **CMVe+p. BC1b** | GCGTTACATAACTTACGGTAAATGGCCCGCCTGGCTGACCGCCCAACGACCCCCGCCCATTGACGTC<br>AATAATGACGTATGTTCCCATAGTAACGCCAATAGGGACTTTCCATTGACGTCAATGGGTGGAGTATTT<br>ACGGTAAACTGCCCACTTGGCAGTACATCAAGTGTATCATATGCCAAGTACGCCCCCTATTGACGTCA<br>ATGATTATGCCCAGTACATGACCTTATGGGACTTTCCTACTTGGCAGTACATCTACGTATTAGTCATCGC<br>TATTACCATGGTGATGCGGTTTTGGCAGTACATCAATGGGCGTGGATAGCGGTTTGACTCACGGGGA<br>TTTCCAAGTCTCCACCCCATTGACGTCAATGGGAGTTTGTTTTGGCACCAAAATCAACGGGACTTTCCA<br>AAATGTCGTAACAACTCCGCCCCATTGACGCAAATGGGCGGTAGGCGTGTACGGTGGGAGGTCTAT<br>ATAAGCAGAGCT | GCAGCATACGTTCTTCGCGCCAACTGATCTGGTAATGCACGTCCAGCAGCCTATATCTTGATG<br>ATATTGGTCGTGAGATCCTACTGGTCCGTCTACGAAGGATACCGAACATTGATAAGCGAACC<br>TCTGCTCTGTTCGAATAATGCCGCCACTCCTAGCAAGAAGCCGAACCGCATAAGCCTTAGGA<br>CTCTTATGGGTCACTCAACTGCCGCCTCGTCTTCACTCACTATTGATTCCAGCGCCTATATCTG<br>CGAATGAATGCACGCGAAGGTCAACAAGTCGCTGGGTGCTCCTAACTTCTGACCAACCGAT<br>GACCCAGCCATAATCCGGTTAACCTGCATTACAATGCACCAATCCGTAAAGCAACTAACACTC<br>CACCATGTTCGTTGTGAGTAGTCTTCCACCAGGAGTCGTTCTTCAGGTATGCTCACTCAGACC<br>GCAATAATGTTCGATCGGAATATGTCATGCTGATTACTGAACCATGGAAGACCGCACAGGTG |
| **CMVe.BC2b** | GCGTTACATAACTTACGGTAAATGGCCCGCCTGGCTGACCGCCCAACGACCCCCGCCCATTGACGTC<br>AATAATGACGTATGTTCCCATAGTAACGCCAATAGGGACTTTCCATTGACGTCAATGGGTGGAGTATTT<br>ACGGTAAACTGCCCACTTGGCAGTACATCAAGTGTATCATATGCCAAGTACGCCCCCTATTGACGTCA<br>ATGATTATGCCCAGTACATGACCTTATGGGACTTTCCTACTTGGCAGTACATCTACGTATTAGTCATCGC<br>TATTACC | TATCTACACTTGACCAAGATGAGTACGAACGTCACCGCCTATACCACTATACCACAGAA<br>GTGACACGCGAATCCAGCTCCACGGTAGAAGTATCATCGAACAGTCATGTTGAGTACCT<br>AGAACAGTATCTCGCAATGCACCATCCAGCCAATACAGAGGAGTCATACCACACCGAC<br>AAGGAGTCTCCGTATTGGTAGCTCCTCCTATTGATAGTGGTTGTTGGAACGACTGTTCTGA<br>CGCCCGGACCACTTCGATAATGGTATGCTTCGGGTCATTGCGTCTGGTTTACCGACATG<br>AGCCTAGTGACCATCTCGCATTGACAGCTTCCGACGGTACTCTATCAACCCACTCACG<br>CTCTAGGTTCGGACCATAGGATCGACTTCACCGCCTTACATGCAGACACCTCCGCCTG<br>ACTTGAATCATCGCTCTACATAACACTTAATGTTGACCGCGAACCTATTGCAGTACTCGTT<br>ACCTATCAATGGACGAGATGCCTGCAC |
| **SV40.BC5b** | TAAGAGGATCCCTGGTTGAGCGTGCAACCCCTTTCAATTGAAACCCAAAATACACTAGGGTACCGTGG<br>CACGCAAGACCGAGGAAGCAAACAGATTCTGCCGAGGCCGGGGTAGACAGTCTCACGCAACGTGC<br>GCATAAGAGCTTAGCGATTTTACATTAATTATCTTCTCACCCAATGTACTCCCGTAAGTTTCTTGAAATTT<br>ATGGGGTCTTCTAAGTCTGGGTATTCTCATCACGAAAGCTATACAAAGGAGCACGAAACTTACTGAAG<br>TACCGGTGTGGAAAGTCCCCAGGCTCCCCAGCAGGCAGAAGTATGCAAAGCATGCATCTCAATTAGT<br>CAGCAACCATAGTCCCGCCCCTAACTCCGCCCAGTTCCGCCCATTCTCCGCCCCATGGCTGACTAATTT<br>TTTTTATTTATGCAGAGGCCGAGGCCGCCTCGGCCTCTGAGCTATTCCAGAAGTAGTGAGGAGGCTTT<br>TTTGGAGGCCTAGGCTTTTGCAAA | GAGCCAAGGAGGTCAGAATACTACAGGTCGCCGATAGCATTAGTCAACCTGACCACATCCC<br>TCCATCGGTAAGGTCCACATCTAGGACACAGAACAGAGTACATCCGCGTTAGTGTTATTGCC<br>ATACAATACTAGTACTTGTGCGAGCGGAACGCTACAGGAATATGCGTCAAGCTCAGGTAGA<br>ACCACAAGCCTATCGAACGCGGTATCTCTACAGGTCGAATACACGCCAAGTACATTGAGTTG<br>ACCATTGTTAATTGAGAACTGTCCCACAATCATCTTACCTGTCTTACTGTCTGGGAATACTCAG<br>AATCCTTAGGAGCATGGACCTAACCAAGTAGAACGCGTACCAATAAGGAAAGCGCATACGG<br>TGTGAAGTCACGCCTAATCCAGTCCACACGTTCGTTACTGCAAGCGTTAGATAAGCTCCGTC<br>GACCTACGCCTGAATGGATCCTACCTATTCTGCATGCGTTGTGGACTTCTTGCCATTCGTATGT<br>CGG |
| **SV40.BC6b** | TAAGAGGATCCCTGGTTGAGCGTGCAACCCCTTTCAATTGAAACCCAAAATACACTAGGGTACCGTGG<br>CACGCAAGACCGAGGAAGCAAACAGATTCTGCCGAGGCCGGGGTAGACAGTCTCACGCAACGTGC<br>GCATAAGAGCTTAGCGATTTTACATTAATTATCTTCTCACCCAATGTACTCCCGTAAGTTTCTTGAAATTT<br>ATGGGGTCTTCTAAGTCTGGGTATTCTCATCACGAAAGCTATACAAAGGAGCACGAAACTTACTGAAG<br>TACCGGTGTGGAAAGTCCCCAGGCTCCCCAGCAGGCAGAAGTATGCAAAGCATGCATCTCAATTAGT<br>CAGCAACCATAGTCCCGCCCCTAACTCCGCCCAGTTCCGCCCATTCTCCGCCCCATGGCTGACTAATTT<br>TTTTTATTTATGCAGAGGCCGAGGCCGCCTCGGCCTCTGAGCTATTCCAGAAGTAGTGAGGAGGCTTT<br>TTTGGAGGCCTAGGCTTTTGCAAA | TCTTCTCGTCCTTACTAATTGCTCGTGCCACAAGGAACATCGACCTATGTGGTACGATGG<br>TATCGAGAACTACAGAGAGCTGATTGTTCGCGGTGTCCTGAGTACCATTACGAAGCGGT<br>TAGTGCAATCTCAATGAGACAAGTGGATAACCAAGTATTAGCCATGCCTAGAGCAGATC<br>CTCCTCTAGCACAATGGTTGCGAATGAGTATCCAACATAGTCCAGCCGTAAGTTCACGG<br>ACAAACATGGACGACTTAATCACCTATTCGCGTCCACATATCCGAATTGTTCTGGTAGGT<br>AAGCCTGCGAGTTGATGCAGGAACTTGTATGCAATCTTGTGACGTAACGCATGTCAACG<br>ATTAGTGTATGCCGAGGAAATCCGTGACTTACTTCGCCGGATAACCATGCCATCTACTG<br>CTGAACTAGTGGATATAAGCAACATACATCGCATCACCAGTAAGTAGTCTCTTCCAATCG<br>CTTATGAAGTATCCGTTGCATCTAC |
| **NoE.BC7b** | no enhancer | GTCGCTCTCATCATCGGATCTTACCGTGCCTGAAGATAGCTGCAACGACCAAGTAGGAACTC<br>GTGTTGAAGCTCCCTCCACCTTAATTGTCATGTACGCGCACCAGTACCGGACCACAGACGTA<br>CAACCTCACCTCTCGCCTACCACCATGAACGACCGATGATGCCGTAGACTGGTGTTCACAAC<br>CTGACCTGCATGGTATGTCACTCTCACGTCTGGAGGTAATGATGAACTGGTCGGAGGTCCG<br>AAGAAGCTAGGACGTGTGTTCGATCTAATACGACCGATATAGTCGTCGGTCATCCTCCGCGT<br>AATGGTAAGTGTAGAACCAGATTGACTGTGAGGATCTCTGGACTATGTTGGAGAGTAACAG<br>AGCGGGATTCGCAGTGTTACCCTTAACGAACATGGTCGACCATATTGTAGCAGGACTAGGAT<br>TGGAGTCTGGAATGAGTATCAGGTATCAATGGTTCGAGCGTTAATCTTACCTTATACGTGAAC<br>ATCAT |

| ID | Enhancer | Barcode |
|---|---|---|
| **NoE.BC8b** | no enhancer | GCTTCTGCAATACTCGAGCCGCACCAATACTTCACACTGGACTGGGATGCAGACTATAC<br>TGGTGATATGTTGATGACGCCATAATCCTGTTGCGTTATACCAGAAACCTGTTCCTAATTC<br>AACACGTCCGATTCCATACGTGTGGAGTCGCTCCACGCTAACTACACTCGTCCAACAA<br>GACGTACTTAGGATATGGAAGTCGATCCAATTCCGCATTACCTTGACCTCCTGATAAGCG<br>TCAAGAAGAGCAGTTGTCACTGATGACCAACGATCGCTTATTCACTCTCAAGCAGTCGTT<br>ACTCCGGTATGATCCAAACTCCTAACCTGCGGACGCTTCCAACACTCTCTGTCCTCCGT<br>TCGTCCGACGATGATGCGAACTCAGGTGTAGCAGCCAACTCTAGGTGCGAAGACCGC<br>ACAAGGTCACTAACTCCGCTTGGTCGATTCCGTGTACTATGCGAGCGGATCAGTATTGAT<br>CGTTCCATCTCTTGACGAATGCGAGT |

Table S11**. Individual Enhancer and Barcode sequences that are used in this study.**

| ID | Enhancer . | Barcode |
| --- | --- | --- |
| **E14b.BC1b**<br>SV40e-ext:<br>Ubiquitous | TAAGAGGATCCCTGGTTGAGCGTGCAACCCCTTTCAATTGAAACCCAAAATACACTAGGGTACCGTGGC<br>ACGCAAGACCGAGGAAGCAAACAGATTCTGCCGAGGCCGGGGTAGACAGTCTCACGCAACGTGCGC<br>ATAAGAGCTTAGCGATTTTACATTAATTATCTTCTCACCCAATGTACTCCCGTAAGTTTCTTGAAATTTATGG<br>GGTCTTCTAAGTCTGGGTATTCTCATCACGAAAGCTATACAAAGGAGCACGAAACTTACTGAAGTACCG<br>GTGTGGAAAGTCCCCAGGCTCCCCAGCAGGCAGAAGTATGCAAAGCATGCATCTCAATTAGTCAGCAA<br>CCATAGTCCCGCCCCTAACTCCGCCCAGTTCCGCCCATTCTCCGCCCCATGGCTGACTAATTTTTTTTATTT<br>ATGCAGAGGCCGAGGCCGCCTCGGCCTCTGAGCTATTCCAGAAGTAGTGAGGAGGCTTTTTTGGAGG<br>CCTAGGCTTTTGCAAA | GCAGCATACGTTCTTCGCGCCAACTGATCTGGTAATGCACGTCCAGCAGCCTATATCTT<br>GATGATATTGGTCGTGAGATCCTACTGGTCCGTCTACGAAGGATACCGAACATTGATAAG<br>CGAACCTCTGCTCTGTTCGAATAATGCCGCCACTCCTAGCAAGAAGCCGAACCGCATA<br>AGCCTTAGGACTCTTATGGGTCACTCAACTGCCGCCTCGTCTTCACTCACTATTGATTCC<br>AGCGCCTATATCTGCGAATGAATGCACGCGAAGGTCAACAAGTCGCTGGGTGCTCCTA<br>ACTTCTGACCAACCGATGACCCAGCCATAATCCGGTTAACCTGCATTACAATGCACCAA<br>TCCGTAAAGCAACTAACACTCCACCATGTTCGTTGTGAGTAGTCTTCCACCAGGAGTCG<br>TTCTTCAGGTATGCTCACTCAGACCGCAATAATGTTCGATCGGAATATGTCATGCTGATTA<br>CTGAACCATGGAAGACCGCACAGGTG |
| **E15.BC2b**<br>Inh-PDYN.8<br>(spinal cord) | CCGGGAGTAGTGGACTTATCCAGGTTCCCAGGCTCAGGGAAGAGGCTTACGACAGCCCAGCCTCACA<br>CCTGGGCAAGTGGCCCCCATCAACCTGCCCAGCTGCCCTGTGTCTGATTTGCCCCTGGCAGCAACCTAG<br>TTCCCATGCAATTGGCCAGATAGTTTTCCCCAAAAGCAAGAACCAACCCATCCTTCCACTGTGAAATGCCT<br>CCCACCCTTGCTTGGGTGACTCGGTGCCAGGGTAGGCTGATATGCCAAAAACTGAAGCCTAAAGTGTC<br>GCAGAGAACCACAAACGCCATCTGTCAACAGGGAGTTAAGGCACGCTAATCTAAACACATCCCTGGAG<br>ATAGCTCTCTACAGCTCATAAATATATACACACAAAACCCTCTCCCAGAGGCAACAAAGCAGGACTGGCT<br>TGAGGCCACGACCAGGCCCGCCTCAGACACCCTGCCTCTCTCTCCTTCCTCTTACCAGGGACTTGGGCA<br>CCAGGGACCCTGTCCATT | TATCTACACTTGACCAAGATGAGTACGAACGTCACCGCCTATACCACTATACCACAGAA<br>GTGACACGCGAATCCAGCTCCACGGTAGAAGTATCATCGAACAGTCATGTTGAGTACCT<br>AGAACAGTATCTCGCAATGCACCATCCAGCCAATACAGAGGAGTCATACCACACCGAC<br>AAGGAGTCTCCGTATTGGTAGCTCCTCCTATTGATAGTGGTTGTTGGAACGACTGTTCTGA<br>CGCCCGGACCACTTCGATAATGGTATGCTTCGGGTCATTGCGTCTGGTTTACCGACATG<br>AGCCTAGTGACCATCTCGCATTGACAGCTTCCGACGGTACTCTATCAACCCACTCACG<br>CTCTAGGTTCGGACCATAGGATCGACTTCACCGCCTTACATGCAGACACCTCCGCCTG<br>ACTTGAATCATCGCTCTACATAACACTTAATGTTGACCGCGAACCTATTGCAGTACTCGTT<br>ACCTATCAATGGACGAGATGCCTGCAC |
| **E16.BC3b**<br>Inh-PDYN.131<br>(spinal cord) | ATGAAAGACTCAGACTCTTCTCTGGGATCTTAATTTTTATGTGGCTTCTTATTACTTTTTTATCCCTTCTAAAA<br>CTCTAAATCCCTTCAACATAGCTTTGGCCCCTTCCACAGCTGAACTGGCAGCAGGCTGATCTTTTTATTGG<br>TGGGTAATGGATTTAGTTTGATTGCTGACACGGGAACAATGAGAGGATAAACAATTCCATTTGTGGCCAT<br>GAAATTCCCCATCTGTTGAGCGCGTTCAGGACGGCAGAGCACTGGGCAGAGACAGATGAATGAGCCC<br>GCCACTCCTGATGTTAATGACATCCGTCATACAAGCTGAACATGTGGCTTCATAAATCAACAGCTAATAG<br>CAACTCTGGTTGCTCATGTCATTAATAAGCCATCCGTTTAAATACATAAGAGTTTGCCATAACATTCAGCCT<br>AGTGACCTGACTTGCAATTCTGATCTCACCAAAAAACCAGAAGTAGTAAAGGATGCCGAAACAGCATTCT<br>TCTAAG | ACACCTTACATGGACGCTACTTACATTGAGAATGTGTAGTGCCAGATGACCTTGAACGTA<br>ATCCTATGTGAGAGAACGTCGGTCCTGAGAACAACGGTACGATGGACAACATACACTCA<br>GCCAACATAAGCGATATTCGGAGCAAGCCTGATCATGTCTTACTCGTTGTCAGCATCGT<br>CAACCGTAAGTCATCAATAGTGAGGTGGTGCCACACATCTCTTCTCAGGTAGATCCTTAG<br>AGGTGATTATGTAGCGCGTCATTAAGGAAGGTATATCACCATCCTCACATTCGTGGATAG<br>ACAACGTCAAGTCACACATTGTTGGTCATCCTTATAAGTACGCGAACTATCCACTGCGTA<br>GCAACACCTAAGCCTAACGCAGTTGGTCACATGCGCGGACCACATATCCAGTGTTCCA<br>GCGAACATGACACTGCTTAGTTGTCCATATAGCTGTAACTTCATCAACCCGAGGTGATTC<br>TATTTGGTATGTCGCAAGAGCCTA |
| **E17.BC4b**<br>Inh-NPY.17<br>(spinal cord) | GGCCTATAATTTCTAAGAAGCCAGGATACATTTCTCTGACAACAGGCAATTACTTTTGTGCGTGTGTTGAA<br>GGTGACATGATCTTCTTTAATTTAAAGGCACGGGAAGACTGCTTTATGGAAAGGTGAATCACCTTTCCCTT<br>TCCTCCTCCTCGCGACTTGACTTGTACCCCAGATTTGCTTCGCTAATTCATTCCTTGCTTGTCTGTCATGGA<br>AAATGGTTTGTTACGTTAATATCATTCTTATGTTTTTCAGTTATGAATAGCTTTTCAAGGCTTAATGTAGCTTT<br>TGAAAATTCTCTTATTACTGATCTTGGTATTATCACCGTAGGAGTTAAAGTCATGAGACAATTTTATGGGG<br>GTAATTTCTTAGTAAAGTAAAATTTTGTGCTGAAATCCTGCTGGGTTCTACACATGTTGATGGATGACCCA<br>CTGTTTAATAGGACATTTTACTGTTCATATATGTGAGGAATGGGTGTGCTAATTAGAGACTTAGAAGACA | ATACTTCGATGGTCCAACCATTCAGCTTACTAGCCTCGAATAATGAGAGAAGTGGTCAAT<br>ATAGAATGGAATCGCCTAATGCAGATGATGAACCTTCCAACCTGCGAAGACGGATCGAC<br>CATAAGGTAATGAGCGATGACCTATATGGACCACCAGTCGTAGCAAGAATTGGTTACAC<br>AGTATGCAGTGTGGTCACTCTGTGCGTATAAGACGCTCCAACTATAACAGTCCGGTCAG<br>GTTGAGGATTCGGATACTACCTCAGATGTAAGAAGAGATGAACAACGTAGCTAACTGATA<br>AGGTCTAGCGAGTTGAAGGAGTATCTTCTTGGTTCGTGATTGCAGTTAAAGATGCACCAG<br>CGACTAACCTTGAACATGTGAGCCTGTTGGAGAGCTTCTGCAATACTCACTGAACGCAT<br>AGGTCGCCAATATCGTCGTGTCGAGCGTACTTGAGCTTCGTATCGGTTCAGGTCAGCCA<br>ACGTGTTCGCAACCTGTACCCAACG |
| **E18.BC5b**<br>Inh-NPY.0<br>(spinal cord) | GTCTCCAGATCTTCTCAGCACCCCCATCCTCCACCCCCACTCCTGGCTCAGTGTTGGCGTTCAGAAATGTT<br>AAATAACAATAATAATAATAACACATCTGAGTGTGACGCTTTAATAGCCAATCTGTCATTTTCCTACTTTATT<br>TCTGAGATTTTAAGAACAGAGCCCTTCCTGCACAAAACATCAGGGTGCATTTATATTAAAAAATGAAACTA<br>ATTAAAATGTTTGAAAACCATAAAGCACACCACCATAACACAGCAGCAGCCCTATTTTTCCATTCGAAAAA<br>GATTTTTCTTCTTCATTTTGTGGCCTGCCCTGATGAGAAAACTGTTGGCGCTGGGAGTCACTACAGGTGT<br>GAGGCTGATAGGAGCCGTTTTGCCCCTGAGCACTGCCTTTGCTCCTAGACCCTGGCATGGCGTCTGAG<br>GCCACAAAGCTGGAGCAGCCAGAGGGTTGGACTTGGATGGAGAAGGCTGGAGGACAAGCTACCCTC<br>CAGGTAATCA | GAGCCAAGGAGGTCAGAATACTACAGGTCGCCGATAGCATTAGTCAACCTGACCACAT<br>CCCTCCATCGGTAAGGTCCACATCTAGGACACAGAACAGAGTACATCCGCGTTAGTGTT<br>ATTGCCATACAATACTAGTACTTGTGCGAGCGGAACGCTACAGGAATATGCGTCAAGCT<br>CAGGTAGAACCACAAGCCTATCGAACGCGGTATCTCTACAGGTCGAATACACGCCAAG<br>TACATTGAGTTGACCATTGTTAATTGAGAACTGTCCCACAATCATCTTACCTGTCTTACTGT<br>CTGGGAATACTCAGAATCCTTAGGAGCATGGACCTAACCAAGTAGAACGCGTACCAATA<br>AGGAAAGCGCATACGGTGTGAAGTCACGCCTAATCCAGTCCACACGTTCGTTACTGCA<br>AGCGTTAGATAAGCTCCGTCGACCTACGCCTGAATGGATCCTACCTATTCTGCATGCGT<br>TGTGGACTTCTTGCCATTCGTATGTCGG |

| ID | Enhancer . | Barcode |
| --- | --- | --- |
| **E19.BC6b**<br>Inh-NPY.49<br>(spinal cord) | GTCTAAGGCACATCTTCTCCAAATGAGAATTGTATTCATTTTTATTAGTAGTAGGATAGTTAAATCACCTTC<br>ATAGAATAACTTACGGGTTTTTAAAACAAATCTGAAAATTTATTTGAAAGTTATTAATGCTAATTTTATTATTA<br>AAATTGTAATAGTCTTACTCTAAAACATTGATGATATGGTGCATGACCAATATTATCATAGCTGCAAGATTA<br>ATTCTACAGTCACTGTATTAAACATTGGCAACAGTACTAAATAGACACACTCAAGTGGCACATTAATGCTA<br>TTGCTAACATTTTTGTCTATAATAGAATTTGAATAGGAAGAAAAATTTAAATTACATGTTTTCTAAAAGAAG<br>GTAGAAGCAGGGATTGTAAATTGGATAAATATAAAAAATCCTTCAACTACTGAAAATGGTTAGCAGTGTA<br>CAAAATGATTTTGACAAGCGTGGTGTTTAAAATGGTAACTTCTGTAGACTCACACCTATAAGGAAATGT | TCTTCTCGTCCTTACTAATTGCTCGTGCCACAAGGAACATCGACCTATGTGGTACGATGG<br>TATCGAGAACTACAGAGAGCTGATTGTTCGCGGTGTCCTGAGTACCATTACGAAGCGGT<br>TAGTGCAATCTCAATGAGACAAGTGGATAACCAAGTATTAGCCATGCCTAGAGCAGATC<br>CTCCTCTAGCACAATGGTTGCGAATGAGTATCCAACATAGTCCAGCCGTAAGTTCACGG<br>ACAAACATGGACGACTTAATCACCTATTCGCGTCCACATATCCGAATTGTTCTGGTAGGT<br>AAGCCTGCGAGTTGATGCAGGAACTTGTATGCAATCTTGTGACGTAACGCATGTCAACG<br>ATTAGTGTATGCCGAGGAAATCCGTGACTTACTTCGCCGGATAACCATGCCATCTACTG<br>CTGAACTAGTGGATATAAGCAACATACATCGCATCACCAGTAAGTAGTCTCTTCCAATCG<br>CTTATGAAGTATCCGTTGCATCTAC |
| **E20.BC7b**<br>mDlx-prom<br>(500bp):<br>Inhibitory<br>neurons | CTTTGGGTTCGTAGCTCCCCAGGATCAATTCTGAACAAAGCCTCCAGCTGCAGTGCCATCCAATTTGAAG<br>CAGACATTGGGGACAATTTAAGGTTTTTATCCACAAGAAGGTTTTTTTCCATTCTCTTAAATGCAGCCATAA<br>TTAGAGTAATTTTTCATGTAGCCCGCTGATTACAGCGTTTTTACCGTCAAAGATAATTACCTGTAATTTTCTT<br>CCACTTTTAATACTAAAAAGCCATCTTTATTTAGATTCAGGAACAGGAAAGGCGAAACAAAAGAGGGAAA<br>TTATTCTGTTATTCATACACAAATTGCAGAGACATAGGACCTAAAATTGAAAATTAACCAAAATTATAATGC<br>TGGAAAGAATGGAAGAGGCTGCAGAAGTACAGCTGGTAGTAGGGCTTTGCTGAATTATTCAAATTATGT<br>TAGAGAAAAAGCTACCATATATCGAAACCAACACTAATTTTTCCTTAAAAAATTTCACCAAATATATGCCAA<br>AC | GTCGCTCTCATCATCGGATCTTACCGTGCCTGAAGATAGCTGCAACGACCAAGTAGGAACTC<br>GTGTTGAAGCTCCCTCCACCTTAATTGTCATGTACGCGCACCAGTACCGGACCACAGACGTA<br>CAACCTCACCTCTCGCCTACCACCATGAACGACCGATGATGCCGTAGACTGGTGTTCACAAC<br>CTGACCTGCATGGTATGTCACTCTCACGTCTGGAGGTAATGATGAACTGGTCGGAGGTCCG<br>AAGAAGCTAGGACGTGTGTTCGATCTAATACGACCGATATAGTCGTCGGTCATCCTCCGCGT<br>AATGGTAAGTGTAGAACCAGATTGACTGTGAGGATCTCTGGACTATGTTGGAGAGTAACAG<br>AGCGGGATTCGCAGTGTTACCCTTAACGAACATGGTCGACCATATTGTAGCAGGACTAGGAT<br>TGGAGTCTGGAATGAGTATCAGGTATCAATGGTTCGAGCGTTAATCTTACCTTATACGTGAAC<br>ATCAT |
| **E21.BC8b**<br>MN.098:<br>motor neurons-<br>Harvatin (spinal<br>cord) | GGCTTCATTTCTGAGCTGCAAACTGCCTGGGTGTCTGGAGAAGCAGGTTGGCGTGGTGGTTAGCAGTG<br>CGTGGGCGGGGTTGCCCGCTCTTGATTTATGATTTCTTTGTCTCTGTGGAAGCACTTAAGTGCAGGCTTT<br>AGTTCCAATGACACTCAGGAGCCTCTGGATTCCAGCACTGGGGATGGGGGTGGGGTAGAACGTTCTCA<br>GGCCTCACCAACCCCTCCCCTGTGTGCTGCCTTTGGGAGAGTCCCAAGGCTTCAGCATTACTTAATTAAT<br>TAGGCCTCTACTGCTACATAGGCTCAGATTCAAAAGAACAGAGTGGCCCACGTCAGCCATTCCCGGAAA<br>AGTCTGATGGCTGGAAGCCAGAGGACTATGTGTCTGCCTTGCTGCCCTTGGCCAGCCCATCCTGAATGC<br>CCAGACTCGGACAATGGAGTAGGTACAGAAGGGTAAAGACAGTGTCTTCTGTACCAGTAAGTGGGCCC<br>TGATCTGCTCTCTACAGC | GCTTCTGCAATACTCGAGCCGCACCAATACTTCACACTGGACTGGGATGCAGACTATACTGG<br>TGATATGTTGATGACGCCATAATCCTGTTGCGTTATACCAGAAACCTGTTCCTAATTCAACACG<br>TCCGATTCCATACGTGTGGAGTCGCTCCACGCTAACTACACTCGTCCAACAAGACGTACTTAG<br>GATATGGAAGTCGATCCAATTCCGCATTACCTTGACCTCCTGATAAGCGTCAAGAAGAGCAG<br>TTGTCACTGATGACCAACGATCGCTTATTCACTCTCAAGCAGTCGTTACTCCGGTATGATCCAA<br>ACTCCTAACCTGCGGACGCTTCCAACACTCTCTGTCCTCCGTTCGTCCGACGATGATGCGAAC<br>TCAGGTGTAGCAGCCAACTCTAGGTGCGAAGACCGCACAAGGTCACTAACTCCGCTTGGTC<br>GATTCCGTGTACTATGCGAGCGGATCAGTATTGATCGTTCCATCTCTTGACGAATGCGAGT |
| **E22.BC9b**<br>Pdyn-prom<br>(500bp):<br>Inh and<br>Exc PDYN<br>(spinal cord) | AGGACTGTTCAGGGCATGGTTTCCAGAGGACAGGCAGCTCTTTGGACTCTCTGAGCTTCTGTAAGCTCA<br>CCTTGAAACTATGATTTCCCTGTGCTGAGTAACCAGACCCAAACCTAACTTCCCCCGTCCAAGCTCCCACC<br>GGCGCCCCTGCATAGCACCAGGAGCACATGCATAAAGCCAGAGAGCCTGAGCGCCCTGCGGATCTTA<br>CCTGCTTACTGCTTTGTCCCTGGCAGGCTTCTGGCTGATCAGGTGGTAGGGTGGCCGATCCAAGATTCA<br>GAAGGTACCCTGACTCTAGCTCAGAGAAGTGCCACAGAGGAAGCGACGAGGGCTACCAAATCCCCCT<br>ATGGCAGTTCTGGAGCCTTATTCCCCTTTTATAGATGAGAGTCTCCCCCTGTCGCTCAGCCCTCTCAGCG<br>CTCGCTCACAGAGCACAGGCTGCCAGGAGCCAATGGGGGAGTGGCAGAGCAGTCAACGCTCAGACC<br>CCCGGGTCCTTCCACTCAACT | ACACTCCTGCGACGAAATCCTGGTCAACTCTAAGGACCATTCAGGTAATCGCACCACGGAGC<br>GCCTCTGCAACAAACACATCCGGTAATAGTCATTGTTCCTGTAACCAACTGACTCTGTCAGAC<br>GCATATCGAGCGTGTGAATGCGAGTTGGATATTGACCTGGGAGCTGGTGAACTGGTCCTGT<br>GGTGGTACGATACTCAGATAGCCTAGTCGAGCAGATCCGTGACTTATTAGGTGGTCAATCTC<br>AGGAATCTGTTGCCTTATATGGAATGAAGGATCGCTATTCTAATTCAGCACCAGTGAACGATC<br>TGGACAAGTCCAGTACCTATCGCGGATGATGTACCTCCGTAGACCGTAAATACAAGTCGCAA<br>TCATATAGCTCTCTTCCTGGTACGAGATGTTAACGCACAGCATCATAACTGTGTTATGGACCAT<br>TCGACATATCCTCGAGTGATACCTTCGTTGAGCGTGAGTTAGGGTCCAATGATAACGGTGAG<br>C |
| **E23.BC10b**<br>Npy-prom<br>(500bp):<br>Inh and<br>GABAergic<br>(spinal cord) | AAGCCACTGGGACCTTGCGGTCCCGCACAGCTGCACCGGGAGCGGGAAAAACCTCGCTCGCCTGCCT<br>CCCTCCCAGAGCGCGCAGTTATCCCGCGGGCCCCAGATATGCTCCCCAAGTAGAATATTCAGGTCCCTA<br>CAGACCCGCACGCAGAGAGCTGACAATTCCCGCGACAGGCAACCTAAGCGGTCCCGCTTCATCTCCCT<br>AGCGGGACTCGACGGTCGGGGATAGAGAGAGGGCTCCGAGGGTGCTCACTCTTGACTGTTCCCTTCT<br>CCCCGCAGAGCGGGCCGCTTAGAATTGGGGTGTGGGTGGCTCCAGACGCCGCCACTCGAGCGGCAG<br>TGGCTCCAGCCTCCTCCCCCGCCGCTGGGGGCGGGAAGTGGCTGTGGGAGTCACCCGGGCGTGACT<br>GCCCCCGAGGCCCCTCCTGCCGCGACAAGGGCGCTCCATAAAAGCCTGTGGGCGACCCGCTCTCCGC<br>ATCCCACCGGTGGATCTCTTCTCTCACAGA | CGACCTCCAATGTATAATCGGTGCGCTCAGATGATCGTATGGAGGAAGTGGTCATACGG<br>ACCGTTATGACAGTTCCCGAATTACGATATGGGTAGATTGACGCGGGTTCCTCCTAGTC<br>CATGCGCCAGTCCAGATGCATCCACAACTTCCGGATTGGTCTACTAATAACACCATTCA<br>CGTGGTAGCAGTCCGATCGGAACCGCGAATTGCCGACGTGATAACTGACCTGGAACCT<br>GGATTTCGACGGTGACTTGGGATTCTGATTGACGAGGAGATGCCGTCAGTGAGTTCCAT<br>CTTCATTATCATCGAACAGTCTTGTGTAAGGAAGACCCAAGCGGAGGATAAAGCACCTG<br>TCGATGGAACTTACCTCCAACGGTGCCTTGCGCTGTTGTGCGAGTAGTAATGTATCTGGT<br>CTTATCCCACCGATATTGATTCCTTACGGACACCACATTGACGCAGATGATAGATAAGGA<br>CTACAGAGTAGGATTAGGACGGTGTTG |

| ID | Enhancer | . | Barcode |
|---|---|---|---|
| **E24.BC11b** MN_Hb9: motor neurons (spinal cord) | AGAGTGGTTAGCTGATGAATTGACAAAAACTAATCAGCTTTATTGGGAAACAGGTTTAAGGGCACGGACGTGTCAATAACGCTCAGCCTGACCCCCTCTTCCATTAGCTAGGCAGGCTGATTAGAGTAAGAAGACCCTCTCGCCGATTGCACCGAAATTGCTTTCATAAACTCACAATATTAGCAGTCGATGTGAGCAGCTAATTTAGAGAAGACATCCAGGCCCGTCACATCGATCATCGGGTTTATGTATCATTATCTCATTTACAGAATGTCGCAGTGTAATACATGTTTTTGGCAATAGGCTTGGCCGCTCGCCGGAGCCGCGCGCTGTTCCTTTCGGCCACACTGCAGGGGACTTCCTCAACAATCTACGATGTCTCTTCGCGTGAACAAGGAGGCACTTTGCACACATATGCACACTTGCGCCCAGGCTCTCAGTGCGCGCTCAGTGCACACATGCACGTGGACACATGCTCCGACATACTACCTGAGCTGCCT | | GTTAGAAGACCAGTTTCCATCGTCGTAGGTTTGACACATTCGTACGTGATATACGAACTCACGCAAGGTTAGGTGTACCTAGTTGCCTCCACTATACGGAATGAGGTGTGTCAGAGCGGTGCCGTACCTGCTAGTGGTGTGGAACGAGTGCAACCGAACGAACCAACTAACCTCGTGAACAGCCGGTAAGATTGAGATCCGTGTCGCCATTCCATTATAGCAAGGTCGACCTGACGTGAAACACTTCTCGCCTCGATGACACGCCTCTCGCCATCTTGACCACACTTGCGACACATACTCGATGTGCCTACGCGACGGTACAACTATAACATGGATCGTCTTACCTTCCTTGACTTGATCTCTGGTGCTGGATGCAGTATATTGCGCGTACATCAATATCCGCACCACCAGAACTAATACCCAATCAGGACTACCTCTTAATGAACACGGTCCGTAGGTAAGCAAATGTCCAGTGAGCAAACTGTTCGGATCGGTAAGCC |
| **E25.BC12b** mm_oligo.18: oligodendrocytes | ATTAGATATTTTCTTTACTTACATTTCAAATTTTATCCCCTTTCCTCATTTCCCCTTTGAAACTCCCCCATCCCATCCCTCTTCCCCTGCTCACTAGCCCACCCACTCCTGCTTCTAGCACCACTCCTTGAAAAGACTTGCTCCACAGATCTGCTTTTGTTCCCTTGTCAAGGAATAATTTTCCACAGAGACTTTGTTTAGGAACTCACTATTCTCGTCTATTGATCTTTGTGTCTGTCCTTTCAGTAGACACTAATACAATTGTTGCATCTGTCCAGTGAGCTTCGAAGTGTATTTGTTTCCATTGTTCTTTTGCAATACTGAATGTCCATTATATAGGATTAGTGTACTGTGCTGCTGATCTGCAAAAAATGAGCAGTAACTGATAACTGAATTCTCAACATGGATTTTCTTGGTGACCTGTCTAGAGATACTGAACATGATACTCTGAAATGTTTTCTGCCCTAGGAGAAGGAATGAGGTGATTTGTTATGGGGTGA | | GCAGTAGTAACGCTCATCGTAGAGCTTCAACATAGGACAGGTGCAGAGGTCATTGGAACAGCCGGAATCTTGTATTATCTATGACCTTGGACCAGACGCCAACACATGGAATTGCCTAGAAAGCAATCAACGCAATCAAGACACGCTCCACCGCTTAGAGTATATTAGCTGATGTCTTAGCACTGTCGGTTAAGGATCTGAACGCGAGCTCAGAGAGAGTGATGGTTAGGTCAACGTCATTTATGATCTCCGTCCAACCTTCATCCACAAGCGCCTACTGAATATATAGGTTGTCTCTCGCCGATAATAGTGGAAAGAGTCGTAATATGCATTATGCTGGACTAGCTACCGAGACTAAGGATGAGTTAGTCGATCTTAGGACTTCACAACGATTGGATTCTCACGGTCAGTCGATATAAGGGACACGATAAGCTCCCAGTTAATCGCCATTAATACCGACCATACGCGTGCAGGAGATGTTGCACCGTCAAGCCTATAGA |
| **E27.BC13b** hu_oligo.41: oligodendrocytes | AGGTCAGTCCTGGAAGAAGTAAAACCATGCCAAGCTGTCTTCTACCCCAAATGCTTGTCCTGTGCATCAGTCGCCATTCTATTGTGTCCCTGAGCCTTGAGCGCACCTCCTTTCTCCTGCCCAAAGTGGGGGGTAGAAGAAGCCCCCTGCATGGCAGCCTGCTGCCAGCACTGAATGGCTCATGCTTAGATCCACTTGGCACACTGGCACACAGCATACACAATGTCCGCTGTTGTGCCTACTGTCCCTGCTTGGTGACAGTACCGCAGGGAACTCTGTCCCCATTGTGTCTGTCCAGTGCTTTGTGAGCATGTGTTTGTTTAGGATAGAATGAAGGGTTTTTCCTCACTCATGAGACTGGGAGAAAATCAGGCAATCTAATAAGCCTATAGTCTCTTCCCTGCAGCACTCAGACCTGCAAGACACAAACAAGACCTTTCTTACCCGCCTGGCTTCTGGCAGCTGCCACAGAACCTGCATGGTGGCCCTGCCAAACTAAC | | CTACCGAGTGATCCAGTCCGCATAAGATACCACACATTGCGAAGGTTACCATTGACCTTGTCTCCGATATTCCTTATCTGAGGAATTACCTGTCCTGCAATTAGACCGTTAGCGTACTCGCTGTCATAGGAGCCGCTCTGAACCACCGGAGCCGGTACTAGGTTGGAATAGACCATACGTCTGACTCGTTGTGTGTCTCGCTCAGGAACCACTTCAGGATTATGGACTACTCATCTGTCGGTCAAGGATAAGTTCGACCTGAAGTTAGAACCAAGTAGCCACGGTCACAGGTTAAGATTCAGTGTATCCTCGCAGACCTTCGCAACCAGACTTGAGTTCTTATCCACGAGTGTGTAGTGCCATTCTACACTACAGCGGTCAACAAGGTGCCCTGTCTTAATTGGTTGCGCGTGTTGACTCCAATTCACGTTCGCCACCGTTATCTCTGTGAGCGAGAACTGCCATAGTGCCTATGGTCAGTTAGCAACCGCACGCGTTAT |
| **E28.BC14b** MN.187: motor neurons-Harvatin | CAAAGCAGCTGCTTCCCACACTCAGAGTGCAAGAGCTCACACCTCCCCCAGCTCTGGGGAAGGACAATGACATTTTGAAAGTACTGTAGCAGCTACTCACTGTTCCTGAAGCTTCCACCCTTCTGCCCTGAACCCTGAACTCTTTTTTCTTTGCTAGCAGTTTTCCTAACATCATTCAAACCAGATGGATTTTTCATCAATTATGTGACATATCACATTGCCCTCATATGTCATAAATTTCATGGATTTCTCAATATCTTGTCATGATGAACCCAAATTTAGCAATATTATCAGACAGTGGCTCTGCATATCAAGACACTTAATGTCTGTGCCAATAACCAAGATTTATCATTTCTTGAATTTAACAGCCAGTGCCATTGGTTCTTTTAAATTGTTTCAGCGTGGGCTAAGCTTTCAGAAATAAAGAAATTAAACTTTTACTGCTCTCTCTCTTATCTCTCTCTCTCTCTCTCTCTCTCTCTCTCTCTCTCTCT | | TGCAGTGTCGAGTGGAAGAACGAAGAGAAGTCATTCCTCGATACCCACACCGTAATGCTTAATACGCGACGATACGTAAGGAATAGTTGGAACATTAGTCACGTCCGACGGTGCTAACTCGATATGATGCGGATGAACTATGCCACTCACCCGCTGTGCTGCTATCCGAGTTCTAGGTGCAATGCAGGTATGAAGTCTAACCTTCTTCCTATAGCGCGTAACTTCAGTTGAAGACGGTACGTCCAGCAGTCTACCTACGAGGTGGTCATTTCCAACCACCTAGACACGTGTATATTGACCAGAGGTACAGTGCGCAGTAGCTGGTTCAAGCGGAATGTTATCGCCGCCAAGTCTGCTGTCTCTCTGTATATCCGGTTAACCGACCTTAGACGCGGTAAGGTATGGTTCCACGGTCGCTATCTAATCCAAGAACCAACGCTAAGGAGATGGACACTTGGTAGACGGTACGCTGGTCCTTGGAGCGTGGATTTAGACCAGCG |
| **E29.BC15b** Inh-PDYN.5 (spinal cord) | CATGCTTCCTCCTCCGTGCCCTCCTTCCATGCCCTCCAAGGAGAGCACGTAGGGCCGTGGAAGGGGCCTCGGCAAGTAAGGGACCTGCTGCTTAGGCTGCATCAGCAGATGGCATCTGCTTACAAAGCCATCCTTGGCTGTCATGACCCTTTCTGAAAGAGAAGACGGATGAGGGATGTTGTTTTTCTTTCCTGAGCCTATAATACAGCAACAGAGATGAGATGTGTAAATGCCTAGCCTAGAGTCTGTCTCATGGTAACTGTTCAGTGACTGGCAGCTACTATGGTTATTAGTAAATTACCATCTGCTTTAATCATTCTCCTGTGAGTTTCATCTGAACCTATTAGTTTCCCAGAGCCAGCTTGCCCTGGGGAAAAGTCACAAAGCTAAGAACAACTTTAATGTTGGCCTCCAGACTCAGTCAGATACCTGTTTGCTAGAACACTTTGAACCAGGGTGTGGGTCCTGCTCCATGAACATGGTGTGAATTACCCTGTAAA | | GTGTAGTGACCATGTGTACCGTTAAGATTCCAATTCGCATATACGCGCATAGCCGCAAGATATACCAGCCAATCTGGACTATCGTTCGTGATACCGCTTACTCCGACACACATCGGAGAACCACGACCTGCTAACACCAACAACCACCGGAACAACAAGAGGTGCCAAGTCCAATACGTCAGACCTTACTTGTTCTCACATCACAATAGGGTCCAACTGTCGCTTCTTATGAACGACTACTAGCTAGACGTGTTCGATAGGTAGTTCCAAAGACAACAGCAATAGAACTGGTATAAGAGGTGCTACTAACATACGTGTTCCAATGAGTTCACTTACGTTGGTGACAACCATGCTTCGATTCATTCAGTTAGTTCCCTATCCGTACCTTAACCTACCTAAGCCGTTAACTCTGGACTATGTTGTTGGTATGATGGAGAATGGTTGTCCTAATATGCAAGCTAGTGACTGCTCGTCGTGCTACAATTAAGAACCAGCGTCGT |

| ID | Enhancer | . | Barcode |
|---|---|---|---|
| **E31.BC16b** hsyn-prom (500bp): neurons | AGTATCTGCAGAGGGCCCTGCGTATGAGTGCAAGTGGGTTTTAGGACCAGGATGAGGCGGGGTGGGGTGCCTACCTGACGACCGACCCCGACCCACTGGACAAGCACCCAACCCCCATTCCCCAAATTGCGCATCCCCTATCAGAGAGGGGGAGGGGAAACAGGATGCGGCGAGGCGCGTGCGCACTGCCAGCTTCAGCACCGCGGACAGTGCCTTCGCCCCCGCCTGGCGGCGCGCGCCACCGCCGCCTCAGCACTGAAGGCGCGCTGACGTCACTCGCCGGTCCCCCGCAAACTCCCCTTCCCGGCCACCTTGGTCGCGTCCGCGCCGCCGCCGGCCCAGCCGGACCGCACCACGCGAGGCGCGAGATAGGGGGGCACGGGCGCGACCATCTGCGCTGCGGCGCCGGCGACTCAGCGCTGCCTCAGTCTGCGGTGGGCAGCGGAGGAGTCGTGTCGTGCCTGAGAGCGCAGCTGTGCTCCTGGGCACCGCGCAGTCC | | TCCAAGTAACGATGCTAATGAGAAGAGTGCGTCCACATCCTCCTGATGACCACTCTTGCGAGCAGGTAGTCGATCAGGTGTCTATGAGTATGCCGTAGCTGCCAAATACCGTGCTAACCGATTGAGGACACCACGCGTCCATTCCGGTAACTGATGTTCGACTCCAGGACTTGCTAATTGATTGTCATAGCGCGATGTCGGTGGTCCTGTCCACGGATTATTACGACCTGATACTTGATGCTCAGGATATTCTACTGGTGGTCATCTGCCTATATGCAATCCGGTATTGAGGAGAGAACATTAAGCATAGAGCCTTATGTCGGATCGCTGAACGCACCTCCAACTGAACGTGATGTAGGTTCCACAATCCATGAGCTACGCCTTCGGTCTACCACTGTCGTCCTTACAGACGATTATACTAGACCTACGCACTTACGTAATGGTGCGACTGTCAACTCGTCCACCAACTCTATGTGCAACGAATGGAAGCGCCTCTGCAG |
| **E39.BC17b** Exc-SKOR2.8 (spinal) | AGAGCCTTCAGGTGTCTCCTTCTTACTCAGAGTTGCTGAAAGAAGTAGGAAAACACATGGGGTTCTATATACGTTTTGGGGATTGCTGATGTGATTTATTGAAGATTTTCGTAGCCTTGAAAAATTCAATTTTATACAAAATCAAGTATTTTGCATAATAAAACTGGTAGAAAATGTGCATATGTTTTACTTTTATGTTCCGCTTTTATGAACACTGTCAGACTGAAAACTGAATTTATGAAGAATTTACCCAACACACCAAGAAAATTGCTGCCAAGAAATCTTGTATTTACACAGTGTGATAGAAAGAGGCTGAGAGGCATTTAACACTGCAGTCCAGTTTATATTCAGCCACACTCTATATTTAACTTTTTCTTTTCCAAAACACGAGAAGCTAAAGGAAAAAATCTTTATAGAAGAAAAAGGCCTTCTTTGAGATAAAAATTGTATAATTCAGAATAAAACTGCAAAGGAGAGAGAAAGAATAATCTCAAGACGAT | | GCAAGAGGACCGATTAATTGCTGTTGTACCGAGGAGTTCTGATGACATATTAGCACAAGGTATGTCCTACCTGATGGACGTGTGAGTGAATGAGTCTAAGGAGCGATGCAACTTACCACTACCGTCAAGTTCTGGAGACAATCATGGTGGGTAATAACCGACTGGACGACCTTGGATGCCGAGAAGATAGTCCGATCGCCAGTCCTCCATTCTATGGATTCAAGGACTCATAACTAACCGCTAATCGCTGTGAGAAAGTTGGTATCATGGTAGCTTAGTGGAATGGGATAGTTGGAGGTATCGCACGGTCTTCCTTACGCAGCACCGAATAAGAGCGTGACTACCAGGTCTACAACACTGCTCATCGTTACATACACGTGGAGAACCTACACGACCTCAGATCAAATTAAGATCAAGCCGCGTGCCATAATCTCGAGTTACACATGACACAGTTGCACGATCCGTGTCTCACCTCTCTTGCCGAACACCATCTCCAACAC |
| **E60.BC18** Exc-PDYN.0 | TTGACCATTTGACACACAGCACTGCAATGGGTGATGGGTAGtttttttttAttttttttcttAAAGTGTAGCTTGTTAAATTCTTACTGAAAATAGACTCCAAACTACATTCTTGGCAGGGCACGAGAGAAAGGCCTGACTGTTATCTGCTGAAGCCATGAAACTGCTCTCAGGAAGCTGTTGGCTGCCAGGAGAGCCCCAGCTGAGGAGAGCGGCTGAGAGAATTACCTCCTCGGGCCCACAGAGGGAGGGAGGCCTGCAGGTCGAAGCCAGCCAATGCTCAGAGCAGCGTGGGCGGCCTGGAGGGTCCACACAGCACTTGGCTACCGAGTGGCAGCATCATTCCTAGCAGGCCATCTTCTCATTCCACTTGTCTGGCATCGGCAGGGCTGCACAGAAGCCCAAAGACTGAGCCAACTGGGGTGGCAGAGAGGAGCAGTAAGAACAGAATGAAAAAAAAGAAAGAAGAAAAAACTCCCTCTAGCTAAATGCTATCCTAC | | ATAAGAGCACCGCTGTGCTCGTCTGTTACTGACCGTCACCGGGGACTAGTCAAGAAAACGTGGTACATATAGTGGAGCATCTGGCGTAGACACAAGGTGTTACCAACCGCAGATGCGTGCCCTAGCGCCTGATTCTTCCCTGAGTTGTAACAAATTGCTCCTCGGCAGCAAGTAGTCCAACCATACAGTGAGTGATCCGGCTTGGACAACACAATAATTCGGAGAACTTCTCTCCCTTATCCGTACGTGTACTCACTTTTGACAGGAGTCCGAGCAAGCCTACATCTTCAAGTCGAAAGCCGCTGGATAGGTAAGCGAATGATATACTCGGGTTTGCAGCGGCACAAGCAGTGCATGGGGGTAGTGGTCAACGTAATTGACTCTTCGCGTCAACTGAAGATGCTGGGAATAGGCTAGCGCCGTGGAGTTTAGTCCATTAGAGGAGGGTTGACCCTGAGTTAACTCTAATGGTGGTTTCGGCAACGACTATGCTGCGGAAG |
| **E41.BC19** Exc-SKOR2.55 (spinal) | GAAGATAGTGTTTGTCAAGGATCAGCAACAAAGCCTGGTACATAGTAGGCAGTCAATAATTACTATATCATATTGATTCTAAAATGCACACTTATTTTCACATTTTAACATCTCTGAAATTGAAATGGATCTATATTGTGTTATGGTTTAATGACAGTTTTTTCTCTTTCTTAATGTGCATAAAATAACAGTACATCTTATAGTCAGTGGCATCTTAAGTTGATGAAATGCAGTAATTTCTTCTCTCTCTTACTACTGTATCCAAGAACATATTTTACTCCCTCACAAACCTGACAAGTAACACAGAAAAAATAAAAGCAGCACTATAACAAGTGGTGCCTTACAACCATTTAGAATAATCAAGAACAATTTTTTCCCATTTCACCTGACTATTTCACTTTTAGAGCTGTGATAAGAGCAACTGAGATTTTTGTGACATAACAGGATTGTTTAAAAGATCTCTTTTTGCCTTTTTCACAGACAGGCCTTGTTTAAATTAA | | CCACTGTCACATCGAAGTGCACTGGGGTGCCAGGCCGTGCGCGAAAAACGCCCCAGCATAGTCTGTGGCAGCAAGTATAGCTGATCCTTTGATCTTATGCAAAGTAGGGTTTCCGATCCAAATGCACAGCCAACTGAATCCGGCTATGATTAGAACTACGTGTGACCTTCTCTTACCGTAGGGATTACATTATCTCGTATTTGTCTCCCAAGGCATCACTACTGAAATGGTCCTAAGAATTGGTGGTCAAAGGGCTGTTGGGGAACGCAAAAGGCTCTAAGTTTGAACTAGCTTCGCACTACCTATCCCGATTGAAGTGTGTAGAATCCTAATACCCAATGAGCCGACTTTCCCACCACTGTTCGGCGGAGCCAAAGCCAGTAACCTTTCACGCTGTTGAAGCGCTTCCTTAAATTGTCATTTCTGCGACGCAATGCCCGGTCTGCATACCAAATCTATGGGCGAAAGGTGGACCTGCGGGATCTCTCAAAACGCGTATT |
| **E42.BC20** Exc-SKOR2.113 (spinal) | AAGATTATTGGCTTTCTGACTAGTGCCTTAAAGCTGGAGTTGGATATATCAAAGCGCTGTTGCATGATTATTTATGATTTCATTTTACTGCAGTGTAAATTTTTGTAATGTCAGCATTTAAGAAATATTATTCCCGTCAACTTTACCAAGCCCTGAGTGGCTTCTTATATATTACTTTTCAGCATATTTGCTAATTTCCTCAAGCAGTGCTAGATATTCACTTTGGAATTTGCCTTTTGATGTCTCTTCTGTGATGAGCAGCCACCAAACAGCCCCCACCACATTCTAATTCTTTATTGCAAACAGTAAAATATGGAGCACTGCATCTGAAAACCACTGTGCGAGACAATCTCATTTTTACTTTCTTTCTGCTTAAATATGCAATAAATATGCATTCTGATAAGTACAGTTCTAGAATAAGATAGTTTCGTGTAACATCTTGGAGTAAAAAAGAGAAAGAAAAATTTTTTTTTTTGAGAAGGAGTCTTGCTTTGTTCCC | | GATAATTGTTGGCACTCCCCGGACCTTGGGCATAGTTCGAGGGCGTACCCCAGCCTATGACCACCCAACGAACCGTACCTAGTTCATTCCGTCGTCACACTCTGTTCGTACTCGCGGAGCTAACATCATTAGCTATTAACGAAGAAGACGAAGCGATAAACTTATATGGTTGGCCACGTCTACCACCCAATTACCTTTCCGCACGTTGATGGACCCTCAATTAGAAAGGTATACTGCGGTTGAATCTTAGGCAGCGGGCTACGCGAAATTGTGGAGACACACCGTTCTTAATCCGGACCATGTGGGAGTTTCGAGAAAGATGGTCGGAATACTTTCGCCCGCTATCTTTTCGAAATAAGTCGTATTAACGGTCGGTGGAAAAACCCCGGAGGTATTGAAATCATTTTCACCGCATTACAATCAGGGCAGGCGCGTCATTAGGGGTAGATCGTGAAGAGCCAGCAAGCGGTCCAGTTCAAACCCAGTCGAGGGTCCACAAA |

| ID | Enhancer | Barcode |
|---|---|---|
| **E43.BC21**<br>Exc-SKOR2.103<br>(spinal) | ATGTAACTCAGCTGAAGAATCATTATAGGATTTTTTTAAAAACTGGAAACTTCCCCCCTGCCCCCAGCTGT<br>CTACAGAAATAGATTATTTCTTCTAATTTCCTGATTTAGTTATAGTCATAGTGTACATAAAAACAGATTTGAT<br>TTTCTTGGAATCAAACATAAAATGAAAGCCAAGGCCTTATCATTGAAACAAGACAGCAGTTTTGCAAAGC<br>TGTGTACATCCTTTGAACTCAGATGACATTGTCTCAGAGGAAAAGATGTTTGATGAGTTTTTTACACTTTAA<br>AGCTCTTCCTAAGGAAAATGAGCTGGGAATCTCCTTTGGTAGAATTGAAGGGGGGTGGATCATTTCGTC<br>TTCAAAAGATGAGGGGAAAAGTGTCATTTTCTATTTCCCTTAAGTATCATTCTTTCAAATACTTGTGTGACA<br>AGAGGTTCAACAGGCTTAGATTTGGGTTTTTAGTTTTCCAGAAATGCATGGAGGACTTAAATGGCCCTGT<br>ACA | TTTTTTTACGTCTGTCTAGTCATCTCGCATCCAGTAATCCCGAGGCATGAACTTTGCCCTCCGT<br>CCTTGTCTCGCGGTTCGGTACTTCTCCTAATCGCTATATGACCCACCTATGTTAGTCACGTGTC<br>TTGACTTGACAGCAAGGGAACACCAAGTTAATACGGTTTTGCATGCACAACCCCGTGTATCG<br>ACCTGTTAGAATCTGGATCGTGAACGACGGCATGTTCTTGGATGGCAGAGGTTATGAGAATT<br>CCCATCCCCGGTACCCCATGCAATCATGACATGCGACGCATTCTGGACTTCAGGACACGCAC<br>GACGAGATAACTACCGTACAGATCATAAATGCCGTGGTACGCGTTATGTACCCATTGCACTAT<br>TGGATGGCGGTTCAAGACTGCATGGTAGCATGTCATAATTGACCTAGCATACGTGAGGAAG<br>ATCCTAAGGCGTAAGGTTTGATGGGTGGATTCCTGAAGATTTGAGTAGTCAAATCGTTGCTG |
| **E44.BC22**<br>Exc-LMO3.15<br>(spinal) | TCGAGAATTTACTACAAAATATACTAGTGTGAGGTGTTTTACAGAATACTCTTTTCCTAGACTCTTAAAAAC<br>AAATCCTTCTGTGTTTTTCCATTTCCCATACAAAGTATAAAAACTTAGAATGGGTGTTCTGAAGTTATTCTGT<br>TCCATGTTCCTCTTCTTAGCTCAAAGTGATTAACAGATATTACTGTAACTAATGAGAACACATTGTGCAGGT<br>CGGAGAATATTAGATAGAAATAGAACATGAATTAATGACATCTGAAAACACTGATGGATGTTCATCAGTT<br>GCTAGGGAAACTTTAGTTTCAGTACTCATGCCAATGAAACAGGAATGCTGTCTATGCACAAGGCAGCTA<br>GGATCTATCTCAGAAATGGAAATGTTGTGCCTGGGAAATGATGATCAACAGGATAAGGCTGTGAATATC<br>AGGATCTTCCAAACAACATAACCCAATAGCTACCCAATATGATGCTGATTGTGTTACCAGAACAGGGTCC<br>TGATC | ACACGCAATCGCAGTAGTGTGCATAGAGCTCTTGTCCTTAGGCTGATGAGGTCCTAAATCCC<br>CACCCTTAACGGTCGTGCCGCAGCTACCGGACGGTGCCTCCGGCTATCGTACGGCGGTATA<br>CCTGACCATAAGATAGAACGCCTGGTGGATATACGCCGTGCCCCCAATTTGGAGTGGCTAC<br>AACAAGTTTAAAAGGTACTTACACACTACAAGAACGGGGTTGTACGCTCAATCAACATGCGC<br>AGGTTGATGACAATTAGTAAATGGCTACTATCTGCGTGGTAGGGGTCGTCAGATATTTCATG<br>CCACTCAGTTGAATACGGTATCCTCACTCGAAACGCTCCCTACCACAGTATTTCCCATCTATCT<br>CGAATGTAGACGTGAATGATTTCAAGACAATAACGCGCTCATTCACATCTTTCACGCTATCTCC<br>ACATTTAAACCCATACTACCTACAGCTGAATGAACATCAATAATCCACTAGGCACAGCGTCTG<br>T |
| **E46.BC23**<br>Exc-LMO3.46<br>(spinal) | GGTTTGAAATGATGGTGAGAAAGCAAGTAAGTAGTGATAACCTAATGACAGACACCAGATGGAAATTAA<br>AAGGAAGGGATCTGTTGACTCATCTGCATTAAATGATAGAAGTTATTTTTAGATGAAATATTTTATTTAATTT<br>TAGTGGATAAATTTCTTTTTCAAAAATTAGATTTCTTCTTCACTAAGACAGAACATTGCTTTCAATTCCAATTT<br>TGGATGCTTTCAATTGCAATTTTGGGAGCCTTAAAAGTTTGAAGTAAATAATTTCATGATCATTTATATAAAT<br>TCACCAGAAAAACATGCAAATCTCATTGAATATATACTTATAATTTGTTTTATTGCTTAAGAGATAATTTTCCT<br>ATTACAGTAAGAATAAATAAATACAAATTTAGATTCATTTAGGCAAATTCTCTGCTCTTACTTGGAGTTAAG<br>GATTATTCATATATCACTGTAGTATTGTAAATATTCTGTAGTTTTGCTATTCAATCTGTATCAAT | AGCCGGTAGACCGGCCCCGCACCTCACCGTCTATGGCACCAGGCTGGCTTGGCATACCGTC<br>CTTATAGCTTATGGTGGGATCCTCATTCACAAGTTGTGCCCGTAACATAGGCGTTCATCGAGC<br>CTACACCAGGTAATGTTCTTGCACTCGTTCTTATCCGGACGGCTGATCGCACGCTCGAATTGG<br>GGGCCCCGACAACGCTAAAGTAGAAATAACGAAGTGGGCATGATGTGAGATCGTCTAACCC<br>GGCATGTACGCGGTGAGCTGTAGTAGAGTCCGGATAGACCTCGTCTAAGCGCTCTTTAGGA<br>AGAGAGAAGACGCGTGTAGGATAGAGAGTTCCCTTACGTCGATTCTTGCATGCTCGGATTC<br>GGTCACCACCTAGTGTCCAATTCCCATTGACAACAGCAACCGCAAGTGCGTTAGGACGCGTA<br>CGACTCTAATGTCTGTATACGTGGAACGGGACAGTGATCAGTACGCAAGAAGTCATTCCAGA<br>GTGAGG |
| **E47.BC24**<br>Exc-LMO3.85<br>(spinal) | AATTGGAATCTCTCCCAGAAGCGTTTCCTGTGATGGTCTGAGCCATACCTGTGCCCGTGCTATGTGCAGT<br>TGGAGAAGTCAACAGCCCCTCCCACCCCACCACCAATTCCCTGAATATTTGCAAACTGTGTTTAGAGAGA<br>AAGAGGAAGCTTTCTTGATAACTTCCTTGTTTATACTAATAGCAAAACTCCAACAACTTTGATGGGATCTG<br>CACCAGCAATCTTTCCCTAGGGCTGATGCAGGATATTTAGGGAAGCTCTGTGTTCTAGGCAGGTAAGAT<br>AGCCAGGTTCCTAACGGGAATCAACATATCTCTTTAATTCATATTTGAAGTAAAAGAAACATCTCTCTTCCT<br>CCCATTTGGCTCAGGAATTCATCTCTAAAAATCTATCCTGAGACAAAATTTTATGCCCAAAGATAACTCATG<br>CATTATTTTTAATAGCAAAAAAAATGGAAACAATCTTTGCCCAACATTAGCAAGATGATCGAATGAATTAT<br>GGCAT | TCTGCGACCTCTGTTTCGTCGGGGATTTAAAGAGCTCATTATGGAAGAGATATCTCTACTTAA<br>GGCAGGCATGGTTATAGACTGAACACACAGGCTCTGATTTATGTCACCTATCGGAACTCAAC<br>ACCTAGATTCTCTTTATCGTGTCATTTCTTTCAGCATCACAAATTCGATCCACCGCTCTAGTCAG<br>ACACCAGACCCTACGTACAAGGATTATAGCAAGAATTTTACCCGGAGGTCACGGGCAGGTT<br>ACGGCCCACATGCGACTTGTCCATCGTAGCTGACTCTCTCGATACAGAGTACCTCTAAAGGA<br>GGTGCAACCGACCTGTACTTACGTTGCTCTCCCATACGAGTCGCGGAATCTAAAACAAGGGA<br>TTCCATGCGATATCACATACGCGATAAATATAGGGTAGCCGGACTCTCACACTTTTCCTTAGC<br>CTGAGACATTGAAAGTGAACCTCTAGTCGGAGGGGTAGCAAGTTGAGCAGTCTCAGTCGGA<br>G |
| **E48.BC25**<br>Exc-LMO3.71<br>(spinal) | AGAACACGTGGATCCGTTAGAGGATGCCTCTTGGTTGCTTAAAAAAAGAGTCAGTCCATTTAAAATTGTT<br>AGTTTTCTATAGTCTTTCAGAACATAGAATTCATTTAGAATTTCCAGCAAAAGTACAGAGAGTCTTCAGGG<br>AAGTTGAAGAAGATAAAATAATTCTGAACACACTGTACACTTTCAGAAGAAAAATATTAAGAGTAAAAATA<br>GTTTCCCAACTCAGTCATTCTTTGGAACTGATAATTTATTCCTGATGGTTCCCCAAAAGAGGGGAAGAGA<br>AGCAAAATAATATATTTGGCTCCTCTAATAGATATTATTCATAGCTCTTTTCCCTCATAGGATAATAGACTAT<br>TAGAGCTTCTATAAAATCCCTTTCAGTAATTTTATGGTGGGTCTGCATTTTGTAAGCCATGTTTTATAACTGA<br>GAAAAGAAAAAAAAAAAACACAACCACCAAGAGAATAATTTTTACAATAATTCATTCCACTGATTTTATTAT | ATTAGGAGTTCATTCTCGACTCCTTTACAGGATGACCGGATGAGCTTATAGGAGCTATATCAT<br>TAATTATCGCGGCGTGATTACGGCTCGGCGCCAAGCACTGGGGGTGAGGATGCATACAAAT<br>TTTCTAGAATACGTCACCAGTTCCGTCGTTTGTTTACGATTATGCCATCAAAAGAGATCTTAAG<br>GTTGACTGGTCGTGATTCTCATGTGTGTAACGAACGTGGGAAGGTGTAGCATTCCCGCCCCT<br>TAGTCCTTCCCGACGGTGTAAACAGTACCCTACCTTTAGTCGTTCATGCGGACTTCTTGAAATA<br>CAAGCCCTGCAATCGGAGCCTGAGGGAAGCTTTGTCGCGCTGAGATCAGATCAGGGCAGA<br>AGACGGACGCTTAGTGTGGTGTTATACTGACTGGCCCGCAGTCTCGATGGGTTTAAAGAAG<br>CCGGTCGTTTTGTGGCGCGGGGCTTGGTATCTTCAAGCAGACAGAACGTGCTCGTGACGCC<br>CTAC |

| ID | Enhancer | Barcode |
|---|---|---|
| **E49.BC26**<br>CS-neg-1:<br>negative control | CGGTCGGCCGGTATGTTGGACCGTCTTTATGTGTGAACAGCATCCGCAAATCTGACTACAGCGAGAGTA<br>CGGGGCATTGGTCCGGGAACGGTCGTCAATACTTCTAGCTAACGAGCACTCTTGTTCTCACTAATATGTG<br>CAGCACGAAACATATGTCCAAAAGAACCAAACAACGTGTTCCCAATATTGTATACTCATAGGGGCGCAG<br>AAAGGCCATAATCGATCCTACCTAGCGGTTCTACGTCAGTGCGATCCCCATTATGCCCGACCGTCGCGT<br>AGTATGACGGCACGCTCTGGAGCAAGTGACTTGGGAGAGCACTGCGGACGAGCTACTCTGGTGCCGT<br>CCCGCTAGCAAGTGCGTGACGGCAGTCTTAGCAGTACTTGCTGGCATAGCTTAAAGCTCGCGCTCGAC<br>GAGACAGGGCGCTACGCTCGTGCTCACACCGCCACAATTCCTCCGGCAAGGCGTGGTGGAATTGCCA<br>AGCGAGCTTTGGGCAGGCCAA | AAGTCTTCAGGAACCTTTCTGACCATCGTCACATTTGCTGACACAGTATAAGGGTTGGCTACA<br>ACCCTACAGCTTCGGTAGCCTGTCGCCCAGACATGGTGTCATAGCGTAACGTTCCGCCCGCC<br>ACGCCACGGGGCAATCGGTAGTGAGAAGTGCGCCTATATTAGAGTCCTAGTCACGGCGTTA<br>GCGGCCGTACGGCGCGGTGTTTCGTGGCCGAGAGCATACGGCAGCCTGGCCCAGTCCACT<br>CACGTGGAAAAACATACCCTAGACCGAGATCTTGAATCCGTACGGGACGCGGGGACGCCA<br>GTGCCTGAATACTCGACTGTTCAACTCGATCCTTTCCTTTTTTCTATCGGCTTGCACTCTCGAGA<br>GTTCCCTCCAAATGCGTGACCCATGCTATGGGCAAGCGCACCTAAACTTCTTTTTCTCGAAAA<br>TCTCCTTGCTCCGATACTGTTGTGACTACACCATGGCCTTACGTTGGTGTATCTCAAGTTCGGA<br>AT |
| **E50.BC27**<br>CS-neg-2:<br>negative control | GGGTTGCGTAGTCGGAGCAACGATTTGCTTCGGAAAATCGAAGACCGCCTTTGCGCGGACGAAGGAA<br>ACTTTATGGCCTCAGTCCGTTCTGGTTAGAGCTTCGTCTGCTTAAACATAGTGACGTCAACCCACTACTGG<br>ATAAACGTATACATGTGTCTCGGCCTTGTAATTGGATCGGAGGAACAGCTTTTACTCCCAAAGAAAGATC<br>GGCTCTTACCGATCCTCGGCTGGCGGAAAGGGGCTACGGATGACAACGTGGGCAAGCTCCCCCTTGC<br>TAACCTAGGTGACCGAAGGCGCCGTAACACAGTCACGTATCTGTTGCGCCTGAGTGTGTCGTTCATTAC<br>TTTCTAGCTCCAACGATCCTGTGGGAACATCCGAGGAACAAGTTGGTCCTCTACAATAAGAACAGAATG<br>GCCAAGTGGGCGCAGATAGCAAACTCCGTTTGGCAGACGCATCCCTTACCCACCATTGGTAGAGGCCC<br>GGAAGGGGCTATTCCGTAT | GAGCGGCTATTTCTAAAGTGATAGTTTGACGCGCTCTATGGATGTTTGGACGCCTGGCAAAC<br>ACGTAAGGCGGTTGGGCCCCGCAAATTTTCAGCCAATGGCACTGGGAACATTATGCGCTCA<br>AAGATAGTTAAACCAAGACTCTCCACAAGCTCCAGCATCTAGCCAAGACGCCGGGAAGCTTC<br>GTTCATATTCCGTAGGTCGTACGTTAGGGGATCGCTGATTACTGGTAACGCGTAGCGGCAAT<br>GTGTATTCTGCGAGTTTCGCCAGCCAAGCCCGCCCAGGTAAGACACATCTCTGCAATTTGCTT<br>AGAGTAATATGGGTTATCACGTGCTTGGATTGCCCTTTAGACCTCGGTCAGCCAATAGCAAA<br>ATGCGGGTTATACTTTTAAATAATGCGAATGTCGTCCGCGACATACTCCCAAAGGGGAATTTG<br>TTCCCGTTAGACAGCGTCGTACTTAAAGCATCGAGTATAAATCTAAACACGTGTAGACGCTGT<br>AC |
| **E14b.BC28**<br>SV40e-ext:<br>Ubiquitous | TAAGAGGATCCCTGGTTGAGCGTGCAACCCCTTTCAATTGAAACCCAAAATACACTAGGGTACCGTGGC<br>ACGCAAGACCGAGGAAGCAAACAGATTCTGCCGAGGCCGGGGTAGACAGTCTCACGCAACGTGCGC<br>ATAAGAGCTTAGCGATTTTACATTAATTATCTTCTCACCCAATGTACTCCCGTAAGTTTCTTGAAATTTATGG<br>GGTCTTCTAAGTCTGGGTATTCTCATCACGAAAGCTATACAAAGGAGCACGAAACTTACTGAAGTACCG<br>GTGTGGAAAGTCCCCAGGCTCCCCAGCAGGCAGAAGTATGCAAAGCATGCATCTCAATTAGTCAGCAA<br>CCATAGTCCCGCCCCTAACTCCGCCCAGTTCCGCCCATTCTCCGCCCCATGGCTGACTAATTTTTTTTATTT<br>ATGCAGAGGCCGAGGCCGCCTCGGCCTCTGAGCTATTCCAGAAGTAGTGAGGAGGCTTTTTTGGAGG<br>CCTAGGCTTTTGCAAA | CTGAAGCGATAAAGTACATTTGTTGGGCTCCTCAGCTCGGTGGTTTCCGAATCAGGACGCTA<br>TGTCCGCTACCTTACTGATATTGGGTTGTGCCCCAAAGCCAATTGTCATGCCAATGGTAGTGC<br>TACCGTACCAGCATACTATCCGTTATTCGCAATGATACGTTGGGCAGTGTATTTAGGAACGCA<br>GGCATGGTAGCACGTCGACCGCGATGGTACACGGCACCACTACCATTGAGGTGCACTTCGT<br>AGCAGCCTAGCGGACCCCGGCCTTCTGTAAAGCGTTAAACCTTTCTCCGTTGTCTGTTGGATC<br>TTCGATAAGTACATCTCGCCCGACCATGTTGGTGTTACATGCGCCGATCTACTAGTGGCGGA<br>TAGGTCCGGGCCGTCAGTATGGGAGGATACAGTTCGCATTAGTGACAAGATAATTGACAAC<br>GAAGCTGAAGTCAAACGATGTCACGCGTGCCAACATGGCTTATGCGTGCTCCTGCACTAAAT<br>GCT |
| **E15.BC29**<br>Inh-PDYN.8<br>(spinal cord) | CCGGGAGTAGTGGACTTATCCAGGTTCCCAGGCTCAGGGAAGAGGCTTACGACAGCCCAGCCTCACA<br>CCTGGGCAAGTGGCCCCCATCAACCTGCCCAGCTGCCCTGTGTCTGATTTGCCCCTGGCAGCAACCTAG<br>TTCCCATGCAATTGGCCAGATAGTTTTCCCCAAAAGCAAGAACCAACCCATCCTTCCACTGTGAAATGCCT<br>CCCACCCTTGCTTGGGTGACTCGGTGCCAGGGTAGGCTGATATGCCAAAAACTGAAGCCTAAAGTGTC<br>GCAGAGAACCACAAACGCCATCTGTCAACAGGGAGTTAAGGCACGCTAATCTAAACACATCCCTGGAG<br>ATAGCTCTCTACAGCTCATAAATATATACACACAAAACCCTCTCCCAGAGGCAACAAAGCAGGACTGGCT<br>TGAGGCCACGACCAGGCCCGCCTCAGACACCCTGCCTCTCTCTCCTTCCTCTTACCAGGGACTTGGGCA<br>CCAGGGACCCTGTCCATT | AATGTCACTTCGCGCGCCTAGTCAGATAAAAACTCCCCTACTATCATTCAACACCCGAGCGGC<br>CTAGACAACGGGTATGATCTTCGGACCCAGGACTCCAAATGTCGCAACGTGTGATAAATCCC<br>GTATTTTTTAACGACTGATGAGACTGCTGTGTAATTACAAATGAAGGCGGGTACCCATCACCT<br>GTAGAAGACTCGAATCGCCGAAGTTACCTTTTTACCACGTTCGGGACTCTTAGATGTAAATCA<br>CGTGCTGCCCAGTGATAGCAACTACTTAATTTTCACGATCCTCTGTGGACAGGTCTGTTGAGC<br>ACTGGCACGCGTGGCTATTGGCAGGTTCTCGCGCGCAGCCTATCCATGCCACATGAAGGAC<br>TGTTTTTCTTCGCCCGATCATTTTCATGACGCGGGCTGGCCCTAAATCCCATGAAGCCATGGA<br>CGCTTGTAGTTTTATCCTGTACACAGGAGCATTTGCGCCAGCCGGTTGCAGTACTTGAGGGG |
| **E16.BC30**<br>Inh-PDYN.131<br>(spinal cord) | ATGAAAGACTCAGACTCTTCTCTGGGATCTTAATTTTTATGTGGCTTCTTATTACTTTTTTATCCCTTCTAAAA<br>CTCTAAATCCCTTCAACATAGCTTTGGCCCCTTCCACAGCTGAACTGGCAGCAGGCTGATCTTTTTATTGG<br>TGGGTAATGGATTTAGTTTGATTGCTGACACGGGAACAATGAGAGGATAAACAATTCCATTTGTGGCCAT<br>GAAATTCCCCATCTGTTGAGCGCGTTCAGGACGGCAGAGCACTGGGCAGAGACAGATGAATGAGCCC<br>GCCACTCCTGATGTTAATGACATCCGTCATACAAGCTGAACATGTGGCTTCATAAATCAACAGCTAATAG<br>CAACTCTGGTTGCTCATGTCATTAATAAGCCATCCGTTTAAATACATAAGAGTTTGCCATAACATTCAGCCT<br>AGTGACCTGACTTGCAATTCTGATCTCACCAAAAAACCAGAAGTAGTAAAGGATGCCGAAACAGCATTCT<br>TCTAAG | CGTCGTAGGGAGGGCATTGGATCTCATACTATGTGCTGGTATGAAAGGGTTGCCTTGCCGTT<br>GACGTATAGAGTCTTTTCGCTACGCCATCGTAGACGCCTGGACATACATGGTGGCTGGAAG<br>CCACAGACAATGATGTTAGTGCTCTTTCCTGTCAGTTAAGCCTTCCAAAAGATGAGTTTAATGA<br>ACGAGTCGTGATACCACGTGCGCTAAGCTGAGTTCGCTGGGGCTGGATTTTGCATGGTACC<br>GAACCGTGTATATCGAAGACTGTTCGATACATTTATGTCAACGGAGGGCAAAAACACGGTGC<br>GCTATTCTCGACTTAAAACCCCTGATCGATGTGTGCTACAGGGTCACAAACACTCATTACATG<br>TTACTGAATTCAATTGGGTACTGCACCCGCAGGGCAACTCAGGTCCTCTAAAACCCGACACA<br>GCCCTGACCATCCATCCTACTGGATGACAGGAGTTAAGTATAATCCCAAGTAGGCACGTAAT<br>GTT |

| ID | Enhancer | Barcode |
| --- | --- | --- |
| **E17.BC31** Inh-NPY.17 (spinal cord) | GGCCTATAATTTCTAAGAAGCCAGGATACATTTCTCTGACAACAGGCAATTACTTTTGTGCGTGTGTTGAAGGTGACATGATCTTCTTTAATTTAAAGGCACGGGAAGACTGCTTTATGGAAAGGTGAATCACCTTTCCCTTTCCTCCTCCTCGCGACTTGACTTGTACCCCAGATTTGCTTCGCTAATTCATTCCTTGCTTGTCTGTCATGGAAAATGGTTTGTTACGTTAATATCATTCTTATGTTTTTCAGTTATGAATAGCTTTTCAAGGCTTAATGTAGCTTTTGAAAATTCTCTTATTACTGATCTTGGTATTATCACCGTAGGAGTTAAAGTCATGAGACAATTTTATGGGGGTAATTTCTTAGTAAAGTAAAATTTTGTGCTGAAATCCTGCTGGGTTCTACACATGTTGATGGATGACCCACTGTTTAATAGGACATTTTACTGTTCATATATGTGAGGAATGGGTGTGCTAATTAGAGACTTAGAAGACA | GGTTAGCTGCCTTTCTTTTGGCAACACCTGTATTAAAATGCGCATCACGGAAATTCAAGTCAGGATCGCCCCGGCGGCTCCTGAATGGAGCTTCTCTGGCATGCGTGCAGGTAGCGTGTTTCTAACAGCGAAAGCCTGGTCGCTGCGATGCCGTGGTTAGCTGAGGACTATGACATGCGGTCTGACCGGGCGTACCGGGAGAAGGGGAAACCGCAAGCGTGCTTTCTACAACTACCTGTCGCCTTGCCATACCACAGTGGTGAAGGCACTGGCTAGATGCTATGTCTTAAGTTTGTTAGGACTGTGGTAGGTTGAAACACTCCACCCTTGACGAAGTTACCCTAAACGTATGCTCGATGTGTGCTCAAGTTGGATGGTCACTTGTAGCGATTATACTTCCTAACTGGGATTCCGGGACACAAGGGGCCACGTAACGACCCCGATCTCTGTAGATTTTGACCAGGACTTGTGCAGGAGAAGAGGCACCTGGCACAGTTCGA |
| **E18.BC32** Inh-NPY.0 (spinal cord) | GTCTCCAGATCTTCTCAGCACCCCCATCCTCCACCCCCACTCCTGGCTCAGTGTTGGCGTTCAGAAATGTTAAATAACAATAATAATAATAACACATCTGAGTGTGACGCTTTAATAGCCAATCTGTCATTTTCCTACTTTATTTCTGAGATTTTAAGAACAGAGCCCTTCCTGCACAAAACATCAGGGTGCATTTATATTAAAAAATGAAACTAATTAAAATGTTTGAAAACCATAAAGCACACCACCATAACACAGCAGCAGCCCTATTTTTCCATTCGAAAAAGATTTTTCTTCTTCATTTTGTGGCCTGCCCTGATGAGAAAACTGTTGGCGCTGGGAGTCACTACAGGTGTGAGGCTGATAGGAGCCGTTTTGCCCCTGAGCACTGCCTTTGCTCCTAGACCCTGGCATGGCGTCTGAGGCCACAAAGCTGGAGCAGCCAGAGGGTTGGACTTGGATGGAGAAGGCTGGAGGACAAGCTACCCTCCAGGTAATCA | GCCTTGACACTGTGTAAGGGGAGCGTTGCTCTGGGGTGCAGTGTAAGAAACAACACGTTTCCTATGAGGCATGCAATTGACCCTGATTGAATATTTTCCAGGGCACTTTTGCCTCCTGCAATAGGCCCGCCGCCTAAAAGGGTCTAACACCTGAAACCGGACTCCAGAGGTTCGGAGGACACTGGTCGGCTGCATCCCTGCACTCCATGAGTTAATTGATTTATAAAGTTTTCTGGCGCCTGTGACACGGTGAAGTTACGAGCCGACCACCTTTGGCCCGTCCATCCTTGGGGTTGGGCCTGTACAATCTCGCAGAATGCGCCTAAAAGTGAATACTTGAAGAACTCGTTCCGAGGGAGCGGCCCTGACAAAGGAACCAGAGAAGACTTTCACTAAACGTGAACTGAGATGGTATGTCCGGATCCAACGCATAGATGGACGGCCTGCTTAACGCGGTGAAAGTATACCGTCGCCAATACTAGCGTGGTTATTCTAGTTGA |
| **E19.BC33** Inh-NPY.49 (spinal cord) | GTCTAAGGCACATCTTCTCCAAATGAGAATTGTATTCATTTTTATTAGTAGTAGGATAGTTAAATCACCTTCATAGAATAACTTACGGGTTTTTAAAACAAATCTGAAAATTTATTTGAAAGTTATTAATGCTAATTTTATTATTAAAATTGTAATAGTCTTACTCTAAAACATTGATGATATGGTGCATGACCAATATTATCATAGCTGCAAGATTAATTCTACAGTCACTGTATTAAACATTGGCAACAGTACTAAATAGACACACTCAAGTGGCACATTAATGCTATTGCTAACATTTTTGTCTATAATAGAATTTGAATAGGAAGAAAAATTTAAATTACATGTTTTCTAAAAGAAGGTAGAAGCAGGGATTGTAAATTGGATAAATATAAAAAATCCTTCAACTACTGAAAATGGTTAGCAGTGTACAAAATGATTTTGACAAGCGTGGTGTTTAAAATGGTAACTTCTGTAGACTCACACCTATAAGGAAATGT | GGGGTTGCAAGATGCATCCTGCTATGGCGCTTAACGGCCCGGCGAATAATCCCCAGCAGTCGGAGGGCGGGCCACGGCCAGCGACTCGAGCATATAAGGCAAACGGTAAGGCTACTATGGTGCGGGTCAGTTCCAGCCGTGCCGCTCGGTAGGGGGCATCTCCGCGCATTTCAACAGGGCGGCAAGGGCATATTAAGAACGTTCTCCTAAGACTCCCAACTATTGGCTGTCCCGATGGGAATAAATGTCGACTATTTTGTATGAGTCTGACCACGTTCGAGTCCTAAACCATCTCTGATCGGTTAGGACCTTACTCGTGTCCCTCTTTTTTCCAACCACGGTGTATTTCTACCACTACAGGCTAATATATCCAATAACCTGGGCCGACTACGGGCATGTCCGCAGTGCGATTGAACTGTACTTTGCGTCGAGAACGTACGAAGTCATCAGCTATGATTAAGCCAATTTCGGCGGTGGTGCTACGCTGGCCTGTTCCTAAT |
| **E20.BC34** mDlx-prom (500bp): Inhibitory neurons | CTTTGGGTTCGTAGCTCCCCAGGATCAATTCTGAACAAAGCCTCCAGCTGCAGTGCCATCCAATTTGAAGCAGACATTGGGGACAATTTAAGGTTTTTATCCACAAGAAGGTTTTTTTCCATTCTCTTAAATGCAGCCATAATTAGAGTAATTTTTCATGTAGCCCGCTGATTACAGCGTTTTTACCGTCAAAGATAATTACCTGTAATTTTCTTCCACTTTTAATACTAAAAAGCCATCTTTATTTAGATTCAGGAACAGGAAAGGCGAAACAAAAGAGGGAAATTATTCTGTTATTCATACACAAATTGCAGAGACATAGGACCTAAAATTGAAAATTAACCAAAATTATAATGCTGGAAAGAATGGAAGAGGCTGCAGAAGTACAGCTGGTAGTAGGGCTTTGCTGAATTATTCAAATTATGTTAGAGAAAAAGCTACCATATATCGAAACCAACACTAATTTTTCCTTAAAAAAATTTCACCAAATATATGCCAAAC | CTTAAGGCTTCACCGCCAGAGTATTTTGCCCATAACCATAATTCTCTCATGATCCTGTTTATCTAGGGTGGTCCATTACATCGAGGCTCCTTGTCTTAGCTTGTGCACATGACTTTTATGTCGATTGGCGCTAATTGTTAAAGATCTGAGAAATTGTAACGCCGTTCTTATGCAACAAGCACAAGAAGTGACTACAATTATTGATGATGTGGATGGTCGCAATTCACACTAGACCAGGTGCAGTACTCCTGAACTGGTGAACCCGTAGCAGCTCGATATATTGACTGTGCTCCTGGAGCATTTTTTTCTGAGCAGGAACCAGGTTTTTTCAACTGCGACAATACGAACACTTTCTTTTAAGAATTGCGCCGCAGCTTTCAAGACCTGGCCGCCGTGATGGACCGGCTTGATCATTCTAGCATGTTTGTCTGTGACCGCCGAGCAGGCCACACGAGACAAAATCGCCTAGAAATCCCTTCAAATGCAAAAGCATAAGTGAT |
| **E21.BC35** MN.098: motor neurons-Harvatin (spinal cord) | GGCTTCATTTCTGAGCTGCAAACTGCCTGGGTGTCTGGAGAAGCAGGTTGGCGTGGTGGTTAGCAGTGCGTGGGCGGGGTTGCCCGCTCTTGATTTATGATTTCTTTGTCTCTGTGGAAGCACTTAAGTGCAGGCTTTAGTTCCAATGACACTCAGGAGCCTCTGGATTCCAGCACTGGGGATGGGGGTGGGGTAGAACGTTCTCAGGCCTCACCAACCCCTCCCCTGTGTGCTGCCTTTGGGAGAGTCCCAAGGCTTCAGCATTACTTAATTAATTAGGCCTCTACTGCTACATAGGCTCAGATTCAAAAGAACAGAGTGGCCCACGTCAGCCATTCCCGGAAAAGTCTGATGGCTGGAAGCCAGAGGACTATGTGTCTGCCTTGCTGCCCTTGGCCAGCCCATCCTGAATGCCCAGACTCGGACAATGGAGTAGGTACAGAAGGGTAAAGACAGTGTCTTCTGTACCAGTAAGTGGGCCCTGATCTGCTCTCTACAGC | CCTCCAGTTTTTATGACGGCGCGCGACGACAGGGCCCATTATGGGGAATGCCTAGTGGGGCAGAATCTACCAGTGGCGTCTACCGGGTGGAGGACTTGTAGGGTAACCTTGATACGCATTGCTTCAGACCCGCTTCCGGACTGACTCTAAGTTAGCGCCCGGGAAATTACTACGCGACACATCATGATTATGCATAGGTGCTAGGGGTCATGAGGGCAGTTGCGTCAAATGGTTTTCACCTAAGCAGTAAACGCAAAGGGAGTCGTCTAATATCAATCTGCCGTAGCACCATCGGTGATTACCAACATAGGGGGTGGGCGAGGGAAATGGGTCGCGGACAATCCGTATCCGCAACAAGGACCCGAGAAGCCACGTGTCGTCCAGACTGGTTTAAGCGGGTGGGGACCTCTAAGGATTTAAGGACTCACTGTGTTTCGGACACCGCCATAGTCACGAGTCCATGATTTCAAAAGTATCGCATGCCTGGATGTGATGACTGC |

| ID | Enhancer | Barcode |
| --- | --- | --- |
| **E22.BC36** Pdyn-prom (500bp): Inh and Exc PDYN (spinal cord) | AGGACTGTTCAGGGCATGGTTTCCAGAGGACAGGCAGCTCTTTGGACTCTCTGAGCTTCTGTAAGCTCACCTTGAAACTATGATTTCCCTGTGCTGAGTAACCAGACCCAAACCTAACTTCCCCCGTCCAAGCTCCCACCGGCGCCCCTGCATAGCACCAGGAGCACATGCATAAAGCCAGAGAGCCTGAGCGCCCTGCGGATCTTACCTGCTTACTGCTTTGTCCCTGGCAGGCTTCTGGCTGATCAGGTGGTAGGGTGGCCGATCCAAGATTCAGAAGGTACCCTGACTCTAGCTCAGAGAAGTGCCACAGAGGAAGCGACGAGGGCTACCAAATCCCCCTATGGCAGTTCTGGAGCCTTATTCCCCTTTTATAGATGAGAGTCTCCCCCTGTCGCTCAGCCCTCTCAGCGCTCGCTCACAGAGCACAGGCTGCCAGGAGCCAATGGGGGAGTGGCAGAGCAGTCAACGCTCAGACCCCCGGGTCCTTCCACTCAACT | AATCGCCTTCCTAAATCATAGGTTCCGAAAATCACCGGAGCGCACGGGGTCCACACACTGCAGATCCGGCAGATTGTATCCATCGCCGTGTGCATGGTGGGTCGGAATTGACTGAGCCGACCTCAGAAGATCTCCAGTGTTAATAGCGTGGTGCATGAAGAAACTTGTGGTCGTTGTAGACATTATGGTAATGATTCCGGGGCGAGTTTAACCAAAACTGACGTATGGATAGGTAACAAGTGCTACGACCGGGCTATCTTCAATTCAAAGGAGCGTTCGTGTCACGAGATGAAACCGTGAAGTCTACGCGGTGCGCAGTATTCTCTTATGCGTCATTTGTTTGGCGATAATAACGTAGGCTTGCCTCGAACGTCGGTTTGTGGCCACATAAGGTACTTGTGGTTTTGTGTCCCTCAGAGCCGTCGCCAACTCTCAATTACCTTGGACATCAAGTATCAGGCATAACTGCACTCATGTCTTTCCACGCTATAAAAAGTTTC |
| **E23.BC37** Npy-prom (500bp): Inh and GABAergic (spinal cord) | AAGCCACTGGGACCTTGCGGTCCCGCACAGCTGCACCGGGAGCGGGAAAAACCTCGCTCGCCTGCCTCCCTCCCAGAGCGCGCAGTTATCCCGCGGGCCCCAGATATGCTCCCCAAGTAGAATATTCAGGTCCCTACAGACCCGCACGCAGAGAGCTGACAATTCCCGCGACAGGCAACCTAAGCGGTCCCGCTTCATCTCCCTAGCGGGACTCGACGGTCGGGGATAGAGAGAGGGCTCCGAGGGTGCTCACTCTTGACTGTTCCCTTCTCCCCGCAGAGCGGGCCGCTTAGAATTGGGGTGTGGGTGGCTCCAGACGCCGCCACTCGAGCGGCAGTGGCTCCAGCCTCCTCCCCCGCCGCTGGGGGCGGGAAGTGGCTGTGGGAGTCACCCGGGCGTGACTGCCCCCGAGGCCCCTCCTGCCGCGACAAGGGCGCTCCATAAAAGCCTGTGGGCGACCCGCTCTCCGCATCCCACCGGTGGATCTCTTCTCTCACAGA | ATTTACAGTCCTTCTGGACATTATCTCCAAAGCTAACATCCTGGGAGTGCCGCGGTCGTCGACAGCCGCCCCGGTAAGTTCGACCGTACTGAAAAACTAGCGTTCGGTGAAGCTATATTCGGTCATCTATTGGCGTCACAAGTCCATACATAGTCGAGAACCAAAGTGATAACGCAGCTACCATGCGTACTCAAAAGATGACCTATTAGGGGCAGCTATTAAGGGTGGTCCGAAGCTCGGACCGAGAACGCAGAGTCGCATCTTCGCTCTCCGAACCTTGCACCCGCAGTCCCTAGTTTAACCGAACTTTGACCGCGCAATGGACATGCTGCGACCACCTCTTATTTTCGCCCGCGTATTGTGGATGCAACCCGAGAGTCAAGGTCGGAGGCACTTTCGTTCATTCACTTGAAATTAAACTATATCCTTATACCGAGTGTGAGTATAAGAGAAATCTTAACCAAGCTCATGAAGGGTTAAAGCGATCCTATACGAATCGC |
| **E24.BC38** MN_Hb9: motor neurons (spinal cord) | AGAGTGGTTAGCTGATGAATTGACAAAAACTAATCAGCTTTATTGGGAAACAGGTTTAAGGGCACGGACGTGTCAATAACGCTCAGCCTGACCCCCTCTTCCATTAGCTAGGCAGGCTGATTAGAGTAAGAAGACCCTCTCGCCGATTGCACCGAAATTGCTTTCATAAACTCACAATATTAGCAGTCGATGTGAGCAGCTAATTTAGAGAAGACATCCAGGCCCGTCACATCGATCATCGGGTTTATGTATCATTATCTCATTTACAGAATGTCGCAGTGTAATACATGTTTTTGGCAATAGGCTTGGCCGCTCGCCGGAGCCGCGCGCTGTTCCTTTCGGCCACACTGCAGGGGACTTCCTCAACAATCTACGATGTCTCTTCGCGTGAACAAGGAGGCACTTTGCACACATATGCACACTTGCGCCCAGGCTCTCAGTGCGCGCTCAGTGCACACATGCACGTGGACACATGCTCCGACATACTACCTGAGCTGCCT | TCTGGGTCGCGGCCTTAGGATATGCACCTGGGCCCGAGATCGCGCGTACCGTTGTACCCCAGGAGCATTGATCGTTTGTCTAAGGCACTTTGTCTTCGTCGTATCCTCTGGGACTTGGGCGATTCGCCCAATCTGGGATCGGCTCTACATTGAATTCCTGGTCGACTTGGACCCCCTGGTTCCCGGTGGTAAGTCCAATTGATCACTTATTATATGGCCTGAAAACCCGCTAGAGCTTCAATGTTCATGAATAGGATACTGAATGGCTGCTTTTGTCCGAACGGTGCACTACCCTGGGTTGGCCGGGCCTTACGTTTCCTCGGCAGCTAGTTGTACAATATGGGGTTGCTGTGTCCAACGGTTCGTTGTGATTGCACGCCTAACTCAGGGGATGCTTTCCAACGATACGTTGATGTAATGAACACATGGGACCTCCACGGACAACAATCCACGTGATATGTTCCGGGCCGCGTAGAAACTCTACTGTAAATAGATCATAC |
| **E25.BC39** mm_oligo.18: oligodendrocytes | ATTAGATATTTTCTTTACTTACATTTCAAATTTTATCCCCTTTCCTCATTTCCCCTTTGAAACTCCCCCATCCCATCCCTCTTCCCCTGCTCACTAGCCCACCCACTCCTGCTTCTAGCACCACTCCTTGAAAAGACTTGCTCCACAGATCTGCTTTTGTTCCCTTGTCAAGGAATAATTTTCCACAGAGACTTTGTTTAGGAACTCACTATTCTCGTCTATTGATCTTTGTGTCTGTCCTTTCAGTAGACACTAATACAATTGTTGCATCTGTCCAGTGAGCTTCGAAGTGTATTTGTTTCCATTGTTCTTTTGCAATACTGAATGTCCATTATATAGGATTAGTGTACTGTGCTGCTGATCTGCAAAAAATGAGCAGTAACTGATAACTGAATTCTCAACATGGATTTTCTTGGTGACCTGTCTAGAGATACTGAACATGATACTCTGAAATGTTTTCTGCCCTAGGAGAAGGAATGAGGTGATTTGTTATGGGGTGA | GCGCAAGGGGTGTTAGTCAGCCCGTCCCTAGAAAACATGACCCTAGTAGATTAACCTTCTGATCTAATTATCTGCGATTTTCAACTCCATTGCCTAAACTGAAGACCGTGAGCTTACGGTTGCTGCTGCGTCCGGAACACCTAATGGGCCCATGTAAGCCTGATACTTCACTATGGAAGTAGTCCCCTGTATCAGCTGATTATGCCACTCATCCACCGGCATGCCGGGACATGTAGAGAACGTTGATCGGGCGGGGGGTTTTCAGTTGGGTCAGACTGTCGAGCCTAAGGCTACCCCTAGCCTGTATCCTCCGACAGCGCCTGAAGGCCTTCGCATTCTATAAGTTGGTCCGTACAGTACAGGCAGATCGCATGCCACAACTTGAACCCCACGGTGTCCCCAAGGTAACCTAACAATCCTTATTTATGTGTCAAACAGCGTACAGTAGCTTGTAATCAGCACGTTGCTGTATCTCCTGCACGGATTACTAGTTACAGTCC |
| **E27.BC40** hu_oligo.41: oligodendrocytes | AGGTCAGTCCTGGAAGAAGTAAAACCATGCCAAGCTGTCTTCTACCCCAAATGCTTGTCCTGTGCATCAGTCGCCATTCTATTGTGTCCCTGAGCCTTGAGCGCACCTCCTTTCTCCTGCCCAAAGTGGGGGGTAGAAGAAGCCCCCTGCATGGCAGCCTGCTGCCAGCACTGAATGGCTCATGCTTAGATCCACTTGGCACACTGGCACACAGCATACACAATGTCCGCTGTTGTGCCTACTGTCCCTGCTTGGTGACAGTACCGCAGGGAACTCTGTCCCCATTGTGTCTGTCCAGTGCTTTGTGAGCATGTGTTTGTTTAGGATAGAATGAAGGGTTTTTCCTCACTCATGAGACTGGGAGAAAATCAGGCAATCTAATAAGCCTATAGTCTCTTCCCTGCAGCACTCAGACCTGCAAGACACAAACAAGACCTTTCTTACCCGCCTGGCTTCTGGCAGCTGCCACAGAACCTGCATGGTGGCCCTGCCAAACTAAC | ATGTCATACCGTCACGTAGCCACCAGTAATGGTGATTCCCCGGAATTAGAGGATGCGTCGGGTGCCTAGTCCAGCCAGAACGGACAATCTTTCCTTGGTAAGCACCGGGAGGTGAGTCAGACTGTTCCTTGGCAGACAGCACTCAATACTAGGCAGATATACCAAATGTACGTGACTTTCAACGATGTGTATCACGGGGCAGCGCGCCTTTCTCTCCTTGTGGCCTGGATTCTCGTGAGCATATACCCGCCAAACCCCTGGTTTCCATAGGTATAAAGGGGTTAGAAGTTTATCTACCACGCATAACACTCTGAAGAACTATCCTGCTGAGCCTCCATTCATTAGATGACATAGAACCATGAACACATAGGTACCCGACTCAGAGATGGCAGACTGTTCTTTCCTGTTGTACTTGCTGGATCTTCAGTGGTGCTAGTCGCGCGACGCAGCTGTTTGGTCACACGTGTCCTGCTCTGTACTTCCACACCGAAAGCAGAGGT |

| ID | Enhancer . | Barcode |
|---|---|---|
| **E28.BC41**<br>MN.187: motor neurons-Harvatin | CAAAGCAGCTGCTTCCCACACTCAGAGTGCAAGAGCTCACACCTCCCCCAGCTCTGGGGAAGGACAAT<br>GACATTTTGAAAGTACTGTAGCAGCTACTCACTGTTCCTGAAGCTTCCACCCTTCTGCCCTGAACCCTGAA<br>CTCTTTTTTCTTTGCTAGCAGTTTTCCTAACATCATTCAAACCAGATGGATTTTTCATCAATTATGTGACATAT<br>CACATTGCCCTCATATGTCATAAATTTCATGGATTTCTCAATATCTTGTCATGATGAACCCAAATTTAGCAAT<br>ATTATCAGACAGTGGCTCTGCATATCAAGACACTTAATGTCTGTGCCAATAACCAAGATTTATCATTTCTTG<br>AATTTAACAGCCAGTGCCATTGGTTCTTTTAAATTGTTTCAGCGTGGGCTAAGCTTTCAGAAATAAAGAAA<br>TTAAACTTTTACTGCTCTCTCTCTTATCTCTCTCTCTCTCTCTCTCTCTCTCTCTCTCTCTCTCTCTCT | CTCTTCGAAGTTGAGCTGCCACAATCGGCCCATATTTTAGATGCAATCCGGAACGTCCAGGC<br>CTATCTACGTCCAAAAGGTGCGAATCTGGTTATTTGTTCCAGCAATAGCTTTGCATTCTGCTCC<br>GCTTCACCTATGGCCGAGGGGAAAACAACCACATCTAGGAGCCTTAGGTGCCCTCCGTAAT<br>CGAGATCACATTACCGAGCCGGAAGACTGTACTGCTCACCAGGTATGTCGCAACATTAGACA<br>CCATCATACGTGGTGAACCGGTTTTCTGTGGAACCATTGTATACAGCTTTTATTCCCGGAAAAT<br>ATTCGGTCCTAACTCATGTTGGGAGATCTCATGTTATCGTAGACTGAGTAGCTTTACAGCCCC<br>TCCCATTCCAACGATTGCACAAGTCAGGCTATTTCTTCTCACGGGAGCAGTGATGTCATTGTT<br>GGCCCCGCCCCCAGAGATGAGTAACTTCCACCTCAACGGTAAGGATGTTTGCTACCAGTCG |
| **E29.BC42**<br>Inh-PDYN.5<br>(spinal cord) | CATGCTTCCTCCTCCGTGCCCTCCTTCCATGCCCTCCAAGGAGAGCACGTAGGGCCGTGGAAGGGGCC<br>TCGGCAAGTAAGGGACCTGCTGCTTAGGCTGCATCAGCAGATGGCATCTGCTTACAAAGCCATCCTTGG<br>CTGTCATGACCCTTTCTGAAAGAGAAGACGGATGAGGGATGTTGTTTTTCTTTCCTGAGCCTATAATACA<br>GCAACAGAGATGAGATGTGTAAATGCCTAGCCTAGAGTCTGTCTCATGGTAACTGTTCAGTGACTGGCA<br>GCTACTATGGTTATTAGTAAATTACCATCTGCTTTAATCATTCTCCTGTGAGTTTCATCTGAACCTATTAGTTT<br>CCCAGAGCCAGCTTGCCCTGGGGAAAAGTCACAAAGCTAAGAACAACTTTAATGTTGGCCTCCAGACTC<br>AGTCAGATACCTGTTTGCTAGAACACTTTGAACCAGGGTGTGGGTCCTGCTCCATGAACATGGTGTGAA<br>TTACCCTGTAAA | GAATTTCTTAGCTTATTACATTAACCTAGTCAAATAACTCACTCTGGATGAAGTCATGCGCAAT<br>AGCTAGTACCAGGATGGCGACGTGAAAGTCGTGTCCTGACTTGTATCGTACGCTAAGTCCG<br>CCGCTTATAAGAGCTCTATGCGTGGCCATTCACGCAGGCTTTATAGCGGGTATATAAGTATG<br>AGCCTTTAACGTATTGAAATCTGATATTATTGTTACCGTGATATCCCAGAAGTAGCCTGGTCCA<br>ACTGTGCCCGCATGGAACTTCGCGATAAGATGTGAGCGTTTGTGTTTTACGGAATTACGAGC<br>AGGATCTCATAAGGTTGCTTGGGAATCTTTGGACAGAAGTGAGTCAGAGCCGGAAGTGCAT<br>TACGTTGCAAAATCTTGACCTAGTGTTTGTAAGACTCCCGTTCAAAGTCTTCGACATTGTCACA<br>ACGACGCAGGATCGAACTCTTTCGGACGCCGATTCATGGGTGGAGAGAGGACCCACCGAG<br>CG |
| **E31.BC43**<br>hsyn-prom<br>(500bp):<br>neurons | AGTATCTGCAGAGGGCCCTGCGTATGAGTGCAAGTGGGTTTTAGGACCAGGATGAGGCGGGGTGGG<br>GGTGCCTACCTGACGACCGACCCCGACCCACTGGACAAGCACCCAACCCCCATTCCCCAAATTGCGCAT<br>CCCCTATCAGAGAGGGGGAGGGGAAACAGGATGCGGCGAGGCGCGTGCGCACTGCCAGCTTCAGC<br>ACCGCGGACAGTGCCTTCGCCCCCGCCTGGCGGCGCGCGCCACCGCCGCCTCAGCACTGAAGGCGC<br>GCTGACGTCACTCGCCGGTCCCCCGCAAACTCCCCTTCCCGGCCACCTTGGTCGCGTCCGCGCCGCCG<br>CCGGCCCAGCCGGACCGCACCACGCGAGGCGCGAGATAGGGGGGCACGGGCGCGACCATCTGCGC<br>TGCGGCGCCGGCGACTCAGCGCTGCCTCAGTCTGCGGTGGGCAGCGGAGGAGTCGTGTCGTGCCTG<br>AGAGCGCAGCTGTGCTCCTGGGCACCGCGCAGTCC | AAATTGCAATAGAATGTCGCCAATCTTGCGCATCCCTTGGGGCCTGAAGACATCGACGGTGT<br>GCACCGCGGGTATCGCGATATCGAGGCATGAGTAAGGGTATTCGTCCGTACGGTGTAGCC<br>CGATGTGGTGTGTCGAGAAAACAGTCCGAATAATTTTTGAGAGAGTCGGCCTCCGATTCGC<br>GACTTTGTGGATGGCGGTAGATAATTGACCCCTAGATAAGAACTCCTATAGACATGGATATTT<br>CCGAATTCGGGCGCAGTGGAATGCTTTTGGCAGACTAAATGCTTTGGTAACGCCTGCTTAAT<br>ACTCATGCAATTCTACATCCCCGTTGTCAAGCGCTCGAGTAGATCGACTAACGCCAGCTGGT<br>CCAGTCACCGAGCGCGTAATACGCACCTCACATGGGACAGATTCAATGCGTGATTTGCAGA<br>CTTATCTTTACAGGAGTAGTGAGAGCGGCAGCCCAATCTGGTTCGACCCACAGCACTTGACT<br>TACCCCC |
| **E39.BC44**<br>Exc-SKOR2.8<br>(spinal) | AGAGCCTTCAGGTGTCTCCTTCTTACTCAGAGTTGCTGAAAGAAGTAGGAAAACACATGGGGTTCTATAT<br>ACGTTTTGGGGATTGCTGATGTGATTTATTGAAGATTTTCGTAGCCTTGAAAAATTCAATTTTATACAAAAT<br>CAAGTATTTTGCATAATAAAACTGGTAGAAAATGTGCATATGTTTTACTTTTATGTTCCGCTTTTATGAACAC<br>TGTCAGACTGAAAACTGAATTTATGAAGAATTTACCCAACACACCAAGAAAATTGCTGCCAAGAAATCTT<br>GTATTTACACAGTGTGATAGAAAGAGGCTGAGAGGCATTTAACACTGCAGTCCAGTTTATATTCAGCCAC<br>ACTCTATATTTAACTTTTTCTTTTCCAAAACACGAGAAGCTAAAGGAAAAAATCTTTATAGAAGAAAAAGGC<br>CTTCTTTGAGATAAAAATTGTATAATTCAGAATAAAACTGCAAAGGAGAGAGAAAGAATAATCTCAAGAC<br>GAT | CCCACCTCCGCGGCGGAGTACGCGGTCGCGCCCAGTCCCTCCATTGAGAGGCCCAGGGCT<br>ACTTGGGTTCCCCAATCCGAAAATAGAAATGATGTGTGGAGTTGCTGCCAACACAATTTTGCC<br>TAACGCGTAATTAAGGCTGACGCTGAGGTCAATAAGGCTTCGGCTGGGCACGCCGAGTCC<br>GAGCCCTGCCGCAACCTCGCCCACATTCCACACTAAGTGTAAATGTAGTGTTTACCCGCGAG<br>GGCAGCACGCCGGCCCGATGCCGGTCATCCCTGCATCAAAGATGATTATCTGAAGGGTTTT<br>CATACTGATATAATGGCAGAGGCTACATCGTGCGTGTTGTGTATTGCTAATGCACCTTCTCCG<br>TCGAGTAAAAAAGGGGCTTCCCAACTACAGGAAGTGACCCGATTCGAAACGCCGCATGCTG<br>GTATATGGATTCCTTATTACCCCACAATATCCTGACATTACTCACGGTGCACGACCTTTCGTGT<br>CACCGA |
| **E60.BC45**<br>Exc-PDYN.0 | TTGACCATTTGACACACAGCACTGCAATGGGTGATGGGTAGttttttttAttttttttcttAAAGTGTAGCTTGTT<br>AAATTCTTACTGAAAATAGACTCCAAACTACATTCTTGGCAGGGCACGAGAGAAAGGCCTGACTGTTATC<br>TGCTGAAGCCATGAAACTGCTCTCAGGAAGCTGTTGGCTGCCAGGAGAGCCCCAGCTGAGGAGAGCG<br>GCTGAGAGAATTACCTCCTCGGGCCCACAGAGGGAGGGAGGCCTGCAGGTCGAAGCCAGCCAATGC<br>TCAGAGCAGCGTGGGCGGCCTGGAGGGTCCACACAGCACTTGGCTACCGAGTGGCAGCATCATTCCT<br>AGCAGGCCATCTTCTCATTCCACTTGTCTGGCATCGGCAGGGCTGCACAGAAGCCCAAAGACTGAGCC<br>AACTGGGGTGGCAGAGAGGAGCAGTAAGAACAGAATGAAAAAAAAGAAAGAAGAAAAAACTCCCTCT<br>AGCTAAATGCTATCCTAC | TCTCTTGTATCAAGTTAGAGCAAATCTATTCGCTCGGCGGCTACTGCATCGACTGATTCC<br>CCCACCCCCATTTGACAGATTACCTTAGATCAAATTCGCTACCTATAATCATTGTTATTGA<br>CTAGAGGTTTACCTGTAAAGTAACCGCAGAAATATATAATACATTAAATGGGCCAGGCAG<br>GGAAGGACGCGGGGCAGATCCGTTAGGCCACTTAGACTCTGTATCAGTCGATTTGCCC<br>GTCCACTTATAGAGGTGTAACCTATACACCTTTGCTCCCGACACTCCGGGTATTGCGGA<br>TTCCACTGTCAGACTCATGTCGCCGATCTTGCAGTCGCCGGTAGCAAAGAGGCTTTCAG<br>GAGGACCCGATGTTCCCTTGGACGTAATCCCTGAGAGTTCGGTGGGTTGGAAATTCATA<br>CTGCGGAGGGGCGGGAATTCGTGTTGAGATTGTTAGGGCTGCGACGTCGGCCTGAACT<br>AAGATCTTGTACCTTTTGGTCATGAC |

| ID | Enhancer . | Barcode |
|---|---|---|
| **E41.BC46**<br>Exc-SKOR2.55<br>(spinal) | GAAGATAGTGTTTGTCAAGGATCAGCAACAAAGCCTGGTACATAGTAGGCAGTCAATAATTACTATATCA<br>TATTGATTCTAAAATGCACACTTATTTTCACATTTTAACATCTCTGAAATTGAAATGGATCTATATTGTGTTAT<br>GGTTTAATGACAGTTTTTTCTCTTTCTTAATGTGCATAAAATAACAGTACATCTTATAGTCAGTGGCATCTTA<br>AGTTGATGAAATGCAGTAATTTCTTCTCTCTCTTACTACTGTATCCAAGAACATATTTTACTCCCTCACAAAC<br>CTGACAAGTAACACAGAAAAAATAAAAGCAGCACTATAACAAGTGGTGCCTTACAACCATTTAGAATAAT<br>CAAGAACAATTTTTTCCCATTTCACCTGACTATTTCACTTTTAGAGCTGTGATAAGAGCAACTGAGATTTTT<br>GTGACATAACAGGATTGTTTAAAAGATCTCTTTTTGCCTTTTTCACAGACAGGCCTTGTTTAAATTAA | AGGAGTTCGTAACCGGTTCTTTCGGGCCCAGGTTATGTCATTTTGTGTCTACCTGGAAGTTGA<br>TGTGAAAGGCGCCTGGACAACACAGAAATTGGCATGCCTGTATGATTGCGTCGACAGCTAA<br>GCTCGAGACATAGGATTATAGTCGGAGTAACTACCGTAGAGGGAGCTATATGTCTTGCCGA<br>ACCTTGAGGGGAGGAGATAAGAGGTGCAGGTTTGACGGGTCCCATCCTCAAAGTGCCGCT<br>ATGTTTCACCGGCACCTTCGACAGATCGGATTTCGAAGGATGATCCCCACTGCGGGAATCAT<br>TCGTATTTAATTCCTCCCACCGATCGTGAGAACAAAAGCGACTGTGTGTGAGAGAACTAATTG<br>CCCCGCCCCCACCTAAAGCGCCGAGGTTTCCCTGCGAACCGTGAGTGGGACTGACGGGGG<br>ACTTGATCGGGGGTTGTGCCACAGGTAACGAGACATAGATGTTGGGTATATCATCATCAGTT<br>TCGATATT |
| **E42.BC47**<br>Exc-SKOR2.113<br>(spinal) | AAGATTATTGGCTTTCTGACTAGTGCCTTAAAGCTGGAGTTGGATATATCAAAGCGCTGTTGCATGATTAT<br>TTATGATTTCATTTTACTGCAGTGTAAATTTTTGTAATGTCAGCATTTAAGAAATATTATTCCCGTCAACTTTA<br>CCAAGCCCTGAGTGGCTTCTTATATATTACTTTTCAGCATATTTGCTAATTTCCTCAAGCAGTGCTAGATATT<br>CACTTTGGAATTTGCCTTTTGATGTCTCTTCTGTGATGAGCAGCCACCAAACAGCCCCCACCACATTCTAA<br>TTCTTTATTGCAAACAGTAAAATATGGAGCACTGCATCTGAAAACCACTGTGCGAGACAATCTCATTTTTA<br>CTTTCTTTCTGCTTAAATATGCAATAAATATGCATTCTGATAAGTACAGTTCTAGAATAAGATAGTTTCGTGT<br>AACATCTTGGAGTAAAAAAGAGAAAGAAAAATTTTTTTTTTTTGAGAAGGAGTCTTGCTTTGTTCCC | AAGATAATAAGTTGGGTCCTATAGGGTGCGAACTTCGTTTATGGTGTTACACCAAGGCCCCT<br>GTCTTGACTGCCATAATAAAACATCTGCGAGGGCCATAGACTTTCTCTTTATGGACCTGGCTG<br>TTTGGGTTGCCGATTACGTCGGCTGCGCAAAGGGCCATAATGACCTACCGGTTGTATTTTAG<br>CACTTATCCCGTGAGCCTCGTCGATCAGTTCACAAATACTCCGCTTCGGCCACGAGCTGTGG<br>CCAGGCCGCAAGCTAGCGTACGTGGACACACAGGATGATATTGTAGGGATTCGTTTTGAATT<br>TGGACTCGACTCTGTCCGCCGGCTTGGCAGAACGGAAACACAGTAGGTGATTGCCACCAGT<br>TGGGAACTCTGCAGGCCGCGAGTTGCAAGGTAAAAAAACACAGAGCCCTGCAACCCACTAA<br>GGCCGTGCCATCCTTAGTCGTAAACAGGTGGAGGCAGTGAAACGGATCTCTTTCTAAGGCA<br>CGTGTT |
| **E43.BC48**<br>Exc-SKOR2.103<br>(spinal) | ATGTAACTCAGCTGAAGAATCATTATAGGATTTTTTTAAAAACTGGAAACTTCCCCCCTGCCCCCAGCTGT<br>CTACAGAAATAGATTATTTCTTCTAATTTCCTGATTTAGTTATAGTCATAGTGTACATAAAAACAGATTTGAT<br>TTTCTTGGAATCAAACATAAAATGAAAGCCAAGGCCTTATCATTGAAACAAGACAGCAGTTTTGCAAAGC<br>TGTGTACATCCTTTGAACTCAGATGACATTGTCTCAGAGGAAAAGATGTTTGATGAGTTTTTTACACTTTAA<br>AGCTCTTCCTAAGGAAAATGAGCTGGGAATCTCCTTTGGTAGAATTGAAGGGGGGTGGATCATTTCGTC<br>TTCAAAAGATGAGGGGAAAAGTGTCATTTTCTATTTCCCTTAAGTATCATTCTTTCAAATACTTGTGTGACA<br>AGAGGTTCAACAGGCTTAGATTTGGGTTTTTAGTTTTCCAGAAATGCATGGAGGACTTAAATGGCCCTGT<br>ACA | GCCCGGCCCGTAGCAGGGAGGATCACCCGATATCGCGGCCGAGGGTGTTCTGGAGCGTC<br>TGTGTCCTCGAATCACTTATACACTGGAGCCCGCTGGGCGATACTGGCCGATTAACCTGATA<br>AGAGGCTAATCGGATTGTAGGCTAAAATCATCAACATCCGATGAGGTGTGCGTTCTCTTAGA<br>CAGGTTAAGGCATCACGTACAATAGAAGTGCTCAAGCAGTTGTGTTGTGGATATCTTCGGTG<br>ACCAAAGTTGCCCCCTGGGGCTACTGAAGCCGGTGACCCAGGCTCCACATACAAGCCTGAC<br>GTCAGGCTTCAACCCTCTTCGAGTGGCGTTGAATGTGAAAGTAGGGTCGTTCGGTGCGTAG<br>GGTGAAACCGCGCCGAAACGGTGCCCAGACTACGGTGCAGGTTAGTCGGCGTTGACAAGA<br>TCAATACGCAACTGCCGTTACCGAAAGAGCTCTCCAGCTAGGAACCACCTCCGAGGAAAGTT<br>GATATCACGGC |
| **E44.BC49**<br>Exc-LMO3.15<br>(spinal) | TCGAGAATTTACTACAAAATATACTAGTGTGAGGTGTTTTACAGAATACTCTTTTCCTAGACTCTTAAAAAC<br>AAATCCTTCTGTGTTTTTCCATTTCCCATACAAAGTATAAAAACTTAGAATGGGTGTTCTGAAGTTATTCTGT<br>TCCATGTTCCTCTTCTTAGCTCAAAGTGATTAACAGATATTACTGTAACTAATGAGAACACATTGTGCAGGT<br>CGGAGAATATTAGATAGAAATAGAACATGAATTAATGACATCTGAAAACACTGATGGATGTTCATCAGTT<br>GCTAGGGAAACTTTAGTTTCAGTACTCATGCCAATGAAACAGGAATGCTGTCTATGCACAAGGCAGCTA<br>GGATCTATCTCAGAAATGGAAATGTTGTGCCTGGGAAATGATGATCAACAGGATAAGGCTGTGAATATC<br>AGGATCTTCCAAACAACATAACCCAATAGCTACCCAATATGATGCTGATTGTGTTACCAGAACAGGGTCC<br>TGATC | AGTTTAATCGCTTGCATGCGGCGGCAGGGCGTCATCTGCCCCCCGGGTATCGTAGTGCGTA<br>CGTATGAGCCATAGCATATTGGGCTTCGTCGGCCACTCAGCAGTCTATAAATGTGAGCCACC<br>CTCCGCGAGACAGGCTGTATTGTTGCCTGAACTAACGTTTCTCGGTCTATACAGACCTCAGTG<br>TGGTCTAACTTAATCTAATCAACGTATTATTACCCCTGTCAAGAACAGAAGCAGATCATATAGC<br>CCGAAGTAGCTGGCTACGGCCCGCGCGATAAGGTGAGCGGGACCTGCGGTTAGATTCCGT<br>TTCACCTTGTGATGGAAGATAAAGCGCACGTTCCGATCGGGCGTCCAAAGCGGCCCAACCC<br>TTCAGGGTTGTCAGGAGGGGAGCTCCCTATCCTGAAGCATTTACGACAAAGCAAACGCTCC<br>CACAGATCCGCCCTGTGCAAATGGTGAACCGATAAACCTTGATAGGGGGACTTATATTGAGT<br>CAGTAA |
| **E46.BC50**<br>Exc-LMO3.46<br>(spinal) | GGTTTGAAATGATGGTGAGAAAGCAAGTAAGTAGTGATAACCTAATGACAGGGACCAGATGGAAATTA<br>AAAGGAAGGGATCTGTTGACTCATCTGCATTAAATGATAGAAGTTATTTTTAGATGAAATATTTTATTTAAT<br>TTTAGTGGATAAATTTCTTTTTCAAAAATTAGATTTCTTCTTCACTAAGACAGAACATTGCTTTCAATTCCAAT<br>TTTGGATGCTTTCAATTGCAATTTTGGGAGCCTTAAAAGTTTGAAGTAAATAATTTCATGATCATTTATATAA<br>ATTCACCAGAAAAACATGCAAATCTCATTGAATATATACTTATAATTTGTTTTATTGCTTAAGAGATAATTTTC<br>CTATTACAGTAAGAATAAATAAATACAAATTTAGATTCATTTAGGCAAATTCTCTGCTCTTACTTGGAGTTAA<br>GGATTATTCATATATCACTGTAGTATTGTAAATATTCTGTAGTTTTGCTATTCAATCTGTATCAAT | GGTTTCAACGGGCGGACTTCGGACGTGCTAACTTTAAGGCCGGAAAACATGTGTGGGCGG<br>CCTCTGACGCCCCACGCAAGTACGATCGCAAGACGGGATAAACTCTGTATCTGCCCTGCCCC<br>TCGCATGAACACGATTTCTGGACGAGCGCTCTCGCCAGATCAGATCGATATTTAGCTTGGAC<br>GGGAATTCCGCCGACTATCATCTAGGCATGGTCGTCCGTTTCTCCTGATTGCTACGGTAAACA<br>CCGGACATGTGTCCTAGGTTATTGCTCAACTAATGGCGGTCGTGCGCCCATAGAGCGTCTAG<br>GGGGAACTTTGTCTGAAAGATGCGTCGATGAAGCGAGGAGATAACGAAGCTCATTTACTGC<br>TCCGTTCGGAGTGTACGTACTCTGGGAATACGTGTTATAATGCGGGCCGAATTACGGCGCC<br>CCGGCTTCTTGGGAGCCACAGGCCTGTCAATGATAATACCGCTTAGTATCCGTGACCAACCC<br>GAGCCTC |

| ID | Enhancer | . | Barcode |
|---|---|---|---|
| **E47.BC51**<br>Exc-LMO3.85<br>(spinal) | AATTGGAATCTCTCCCAGAAGCGTTTCCTGTGATGGTCTGAGCCATACCTGTGCCCGTGCTATGTGCAGTTGGAGAAGTCAACAGCCCCTCCCACCCCACCACCAATTCCCTGAATATTTGCAAACTGTGTTTAGAGAGAAAGAGGAAGCTTTCTTGATAACTTCCTTGTTTATACTAATAGCAAAACTCCAACAACTTTGATGGGATCTGCACCAGCAATCTTTCCCTAGGGCTGATGCAGGATATTTAGGGAAGCTCTGTGTTCTAGGCAGGTAAGATAGCCAGGTTCCTAACGGGAATCAACATATCTCTTTAATTCATATTTGAAGTAAAAGAAACATCTCTCTTCCTCCCATTTGGCTCAGGAATTCATCTCTAAAAATCTATCCTGAGACAAAATTTTATGCCCAAAGATAACTCATGCATTATTTTTAATAGCAAAAAAAATGGAAACAATCTTTGCCCAACATTAGCAAGATGATCGAATGAATTATGGCAT | | GGGCATTCACGCCACCATGGTACTCTCCGGCATGTTGTGGTTAGTCCGCCTGGCGGCTACTTGACGATGTTAATACCGGTTGGCCCAAATCATGTCGGTAATGGCGCTTAGACTGCTGTATTCTTCAGAGTTTAATTTGACAGGCTGTTTGTGCTTTCCTGGGCTCGCTCCAGCCATCTGGAACTGGCCGCATTGGGTCATCAGGTCATAGGCGGTAAGTCAATACAGGACGTAATGGCGAGCAGCGCCCGACCTGGTAGGGCACAGTGTCTTGGTGTCTGTCCTACCTTGACTCTAGTGGGATACTAGCATTATGCTGCAAGCTCGCGTCGTGCAACTGTCAGTTCCGATCCTATCGAATCCTTTAGTGCTGACTCGGGCATATTACGAAAAAGTACTTGAAACAGAAATGGCCTAGTTTAGCCCTATCACTCCGTATGGGCCCACCGGTGGCGCCAGAATCTACTCCACTTAGCCGGAGAGCTGCGGATCGCTTAATA |
| **E48.BC52**<br>Exc-LMO3.71<br>(spinal) | AGAACACGTGGATCCGTTAGAGGATGCCTCTTGGTTGCTTAAAAAAAGAGTCAGTCCATTTAAAATTGTTAGTTTTCTATAGTCTTTCAGAACATAGAATTCATTTAGAATTTCCAGCAAAAGTACAGAGAGTCTTCAGGGAAGTTGAAGAAGATAAAATAATTCTGAACACACTGTACACTTTCAGAAGAAAAATATTAAGAGTAAAAATAGTTTCCCAACTCAGTCATTCTTTGGAACTGATAATTTATTCCTGATGGTTCCCCAAAAGAGGGGAAGAGAAGCAAAATAATATATTTGGCTCCTCTAATAGATATTATTCATAGCTCTTTTCCCTCATAGGATAATAGACTATTAGAGCTTCTATAAAATCCCTTTCAGTAATTTTATGGTGGGTCTGCATTTTGTAAGCCATGTTTTATAACTGAGAAAAGAAAAAAAAAAAAACACAACCACCAAGAGAATAATTTTTACAATAATTCATTCCACTGATTTTATTAT | | GGAGCACTGCATTCCTTAGGGCAACAGACCGTGTGCGCACGAGGCAACCTTCTAGGGCTGGCTGTCGTCGACGACCTAGTTCATCAACCTTTTTGTGCTGACGCAGTTAGATCAACCCTGTCTGACGGCGAGAAAGTATGTCGAGGCACACTCGTTCACCCTGGTTTCTAAACGTCTAGTACGAGCGAGAACTCTCCAGGTGAGGATTGCCGCTGATGTACCTTCACTTCTAATGAACATACGACAGTGAACAACGCAGGTACATTCTTGAACCAGCGACGACGAAAAAGGCAACTGCAGGCAATTGCATACTACCTGGCGTAATTGAGTACAAACTTGAGACATCGAGCATATTCATGCTCGGTTACAAGGCGAGGATCAACTACTGGACAGGTTCAGTTTGAGCCTAGGCTGTGCAGGAACGTTAGTCATAGCCTCAGTGTCTATAGTTTGTCGGCTTATTTTTCTCTTGGAGGAGCTTGCGGCACATTCCGCGTAGC |
| **E49.BC53**<br>CS-neg-1:<br>negative control | CGGTCGGCCGGTATGTTGGACCGTCTTTATGTGTGAACAGCATCCGCAAATCTGACTACAGCGAGAGTACGGGGCATTGGTCCGGGAACGGTCGTCAATACTTCTAGCTAACGAGCACTCTTGTTCTCACTAATATGTGCAGCACGAAACATATGTCCAAAAGAACCAAACAACGTGTTCCCAATATTGTATACTCATAGGGGCGCAGAAAGGCCATAATCGATCCTACCTAGCGGTTCTACGTCAGTGCGATCCCCATTATGCCCGACCGTCGCGTAGTATGACGGCACGCTCTGGAGCAAGTGACTTGGGAGAGCACTGCGGACGAGCTACTCTGGTGCCGTCCCGCTAGCAAGTGCGTGACGGCAGTCTTAGCAGTACTTGCTGGCATAGCTTAAAGCTCGCGCTCGACGAGACAGGGCGCTACGCTCGTGCTCACACCGCCACAATTCCTCCGGCAAGGCGTGGTGGAATTGCCAAGCGAGCTTTGGGCAGGCCAA | | CTTTGACCACACACAATACTTCCTCCAGGCGGTCTGGGAAGGTTTAGCATTTTGGGACCAAATATGCCCACACCCATGAAGTAGGCTTGGCGTGACCACGACACCAAGTTGCGCCTAGGACGGTAAACGCCGTGTGGTGTAGCACTATTTTCGTGTGTCGCTACTGCCCTACACGGCAGAGGGCTAGGACTTTAAATTCTATGAACACGTACAGGCGACGAAGACCAATCTGGTGACTGCCCTGAGAGGCCGCCAGCCCGTGGATTTTTGAGAGTTGGAAGTAAGTACCCCCCCCTTAAGTTGCTCTTTTATGGCCCTCAGGTGCTCGTACCCGGGGCCGCATGTAACAGTACTCCACAACAAATAGTCCGGCGGATCGTATTCTACCTAGCAGCTTCCTTTCGTTGACTACCCACTGTGCAGAATGGGCCGCAATAATTTGGGTCTGCCGTAGGGTGACAACTGTCCTTTATGGTGATGTCTATATAGGAAGGCGGGCA |
| **E50.BC54**<br>CS-neg-2:<br>negative control | GGGTTGCGTAGTCGGAGCAACGATTTGCTTCGGAAAATCGAAGACCGCCTTTGCGCGGACGAAGGAAACTTTATGGCCTCAGTCCGTTCTGGTTAGAGCTTCGTCTGCTTAAACATAGTGACGTCAACCCACTACTGGATAAACGTATACATGTGTCTCGGCCTTGTAATTGGATCGGAGGAACAGCTTTTACTCCCAAAGAAAGATCGGCTCTTACCGATCCTCGGCTGGCGGAAAGGGGCTACGGATGACAACGTGGGCAAGCTCCCCCTTGCTAACCTAGGTGACCGAAGGCGCCGTAACACAGTCACGTATCTGTTGCGCCTGAGTGTGTCGTTCATTACTTTCTAGCTCCAACGATCCTGTGGGAACATCCGAGGAACAAGTTGGTCCTCTACAATAAGAACAGAATGGCCAAGTGGGCGCAGATAGCAAACTCCGTTTGGCAGACGCATCCCTTACCCACCATTGGTAGAGGCCCGGAAGGGGCTATTCCGTAT | | CAGTTTCTGTGTGCACTAAGGTAACTCTAAAATCGGTTCTCGGCCTCCGAGCGCTTGTCAAGCACCACCAACAAGCTGTAGATTGTCGTCGACGGGGCCCTCCGCGTTATATTCGGGACAAGTGTGTCTGGGTCTGCCGTTGATCGTATATGTGTGCAGTTAAGTCAGTAAAGAAAACGTGAGGTTTACGGTGACCGACAGTCACATCAACACTGGCACGTAGAAGGGCGAGTGCGTGAATCCGAGTTTTCTGAGCTGTCTGTATCGATAGCGGCCACTGTATAGGTCGTGAGCGCTTTTTATTAGGCCGTGAAGACGAAACCAGCTGCCCGGAATAAGTACTGGTTATAAGCGCCGACCAACTGTTCCGCCGGGAATCAGCCGAGTACTCTGCCTCCTCACAACTTTATAGTCAGTAAAGTAGCTCTGAGGAAATATCGGGGATGTACCTCACCCTCAACCCTTGAGAAGGCCGCGCTAAGGCCCTTTACGACATCAAG |
| **E14b.BC55**<br>SV40e-ext:<br>Ubiquitous | TAAGAGGATCCCTGGTTGAGCGTGCAACCCCTTTCAATTGAAACCCAAAATACACTAGGGTACCGTGGCACGCAAGACCGAGGAAGCAAACAGATTCTGCCGAGGCCGGGGTAGACAGTCTCACGCAACGTGCGCATAAGAGCTTAGCGATTTTACATTAATTATCTTCTCACCCAATGTACTCCCGTAAGTTTCTTGAAATTTATGGGGTCTTCTAAGTCTGGGTATTCTCATCACGAAAGCTATACAAAGGAGCACGAAACTTACTGAAGTACCGGTGTGGAAAGTCCCCAGGCTCCCCAGCAGGCAGAAGTATGCAAAGCATGCATCTCAATTAGTCAGCAACCATAGTCCCGCCCCTAACTCCGCCCAGTTCCGCCCATTCTCCGCCCCATGGCTGACTAATTTTTTTTATTTATGCAGAGGCCGAGGCCGCCTCGGCCTCTGAGCTATTCCAGAAGTAGTGAGGAGGCTTTTTTGGAGGCCTAGGCTTTTGCAAA | | GTCGCGATACCACTAGACGCACCGTCGTTTTGCCCCTGTGATAGATCATGTCGATGACTACCCTTCTTCCCCTTGACAGACGAGTATGTCGTCGAGCGCCGCCCTGCATATTGGGCCTAATACCCGATTGTGGCTTAGGCCTGTAATTCATCCCGCAATACTCTAGCTGCTCTTACTTATCCCAACTGTTCTTCGCCTCAGCATCTCTTACCTTTTTACATTACATTAGGGTCTTAGGCCATCGACGGTAACCGATCACCGGCTAAACAAGAGCTACTACCGAGGGACGAACTCACATGTCGAGAAACACACAGGATATGGTAGCATTTCACGCAGCTAATTGTCGAAGGTTATAGGTCAAGAATATAAAGGAGGTGTAGTGCCTGCACGAAGTCGATGCCTTGTACTCAACGGTTAGTGTAAACCAATTCGTACGAGAACTAGAACCTTACAAGGCGGGTAAAAAGCCCAGTAGCTCAGTGGTCTTGAACTTTATTGTT |

| ID | Enhancer | . | Barcode |
|---|---|---|---|
| **E15.BC56**<br>Inh-PDYN.8<br>(spinal cord) | CCGGGAGTAGTGGACTTATCCAGGTTCCCAGGCTCAGGGAAGAGGCTTACGACAGCCCAGCCTCACACCTGGGCAAGTGGCCCCCATCAACCTGCCCAGCTGCCCTGTGTCTGATTTGCCCCTGGCAGCAACCTAGTTCCCATGCAATTGGCCAGATAGTTTTCCCCAAAAGCAAGAACCAACCCATCCTTCCACTGTGAAATGCCTCCCACCCTTGCTTGGGTGACTCGGTGCCAGGGTAGGCTGATATGCCAAAAACTGAAGCCTAAAGTGTCGCAGAGAACCACAAACGCCATCTGTCAACAGGGAGTTAAGGCACGCTAATCTAAACACATCCCTGGAGATAGCTCTCTACAGCTCATAAATATATACACACAAAACCCTCTCCCAGAGGCAACAAAGCAGGACTGGCTTGAGGCCACGACCAGGCCCGCCTCAGACACCCTGCCTCTCTCTCCTTCCTCTTACCAGGGACTTGGGCACCAGGGACCCTGTCCATT | | AAGCTTTGGAATATAGCCGCCCCCTCCAGACCTGCCTCACGTTTTATATTACATTTAGCTTTTAAACCAAGACGAAGGCGTTCTCGTGATAAGATGGCCGTAGGGTGGATTTAGTGACGAGGGAGTCAGCCTTCTAGGCAAGCGACACGAAAGTGCGATATCAGGTCCTCCTTGTGGTTTCTTCAACCCGCCGGTAGATTTTGACCATTCGACCTCGGGTGGCACGCCGCCCTTGTGTTTATATATTGAGTGCATCGGGAAGCGCCTTGACGGAAAGACGAAAACCTTTACAGAATATAGCAACCCAGCAGTCAAACGTGGGGCTTGATAGTGTGGCTAGAACTTGTATGACAAGGACCATTGAGGCACGCTAGATACCGGCAATGGGAGGATCCAGGATGCCGCGTAGCAGCATCCAATTTACTAGCGTGCACAGCTAAGACTATTAGTGGAACAACGATGGGTCGTAGGAGTCGATTCTTCGTTGACTCTGCGTGGCA |
| **E16.BC57**<br>Inh-PDYN.131<br>(spinal cord) | ATGAAAGACTCAGACTCTTCTCTGGGATCTTAATTTTTATGTGGCTTCTTATTACTTTTTTATCCCTTCTAAAACTCTAAATCCCTTCAACATAGCTTTGGCCCCTTCCACAGCTGAACTGGCAGCAGGCTGATCTTTTTATTGGTGGGTAATGGATTTAGTTTGATTGCTGACACGGGAACAATGAGAGGATAAACAATTCCATTTGTGGCCATGAAATTCCCCATCTGTTGAGCGCGTTCAGGACGGCAGAGCACTGGGCAGAGACAGATGAATGAGCCCGCCACTCCTGATGTTAATGACATCCGTCATACAAGCTGAACATGTGGCTTCATAAATCAACAGCTAATAGCAACTCTGGTTGCTCATGTCATTAATAAGCCATCCGTTTAAATACATAAGAGTTTGCCATAACATTCAGCCTAGTGACCTGACTTGCAATTCTGATCTCACCAAAAAACCAGAAGTAGTAAAGGATGCCGAAACAGCATTCTTCTAAG | | TCTTATTCTCTCGGCCACGATCCATCTAGCACTGGGCGCCATCCGAGAAAACATGAGCCGGGTTGAAGTCGGTGGTTGCCGGTTTGGCTAGACAAGGGCATCTTGTCCACATCTCCCCAACCTCTAAAGCGTTGTCCATGGAGCGGGGTTTTCGTATCAACGACGACTTTAATACCTGCCCGGTGTACCCTTTATCAATAATCGCTGTTACTGTGATCCATGCGATACTCAACATATCCAGGTGTCCGTTGTTATACCCCCTGATCCAGAATACGTGGTGAATGTCTTCTCAGGGGACCCAGGGGTTTTGTTCTGGTTTAGATCAGATTGGCCAGAAGACAGATTCGCGCGTGCGACCTCCTCTCTGTCACCCTCCTGGACTGCCTCCGTCTTAGTTATGTTGAGGTCCTGCATCTGGGAAGGCTACGGACCTGGAGTACTATGAAATGCATTACTCTCCAGTCTTCTTTGTTTATATCTGCAAGTAGTGCTTTGAGATG |
| **E17.BC58**<br>Inh-NPY.17<br>(spinal cord) | GGCCTATAATTTCTAAGAAGCCAGGATACATTTCTCTGACAACAGGCAATTACTTTTGTGCGTGTGTTGAAGGTGACATGATCTTCTTTAATTTAAAGGCACGGGAAGACTGCTTTATGGAAAGGTGAATCACCTTTCCCTTTCCTCCTCCTCGCGACTTGACTTGTACCCCAGATTTGCTTCGCTAATTCATTCCTTGCTTGTCTGTCATGGAAAATGGTTTGTTACGTTAATATCATTCTTATGTTTTTCAGTTATGAATAGCTTTTCAAGGCTTAATGTAGCTTTTGAAAATTCTCTTATTACTGATCTTGGTATTATCACCGTAGGAGTTAAAGTCATGAGACAATTTTATGGGGGTAATTTCTTAGTAAAGTAAAATTTTGTGCTGAAATCCTGCTGGGTTCTACACATGTTGATGGATGACCCACTGTTTAATAGGACATTTTACTGTTCATATATGTGAGGAATGGGTGTGCTAATTAGAGACTTAGAAGACA | | TGTCGTGATTGACTCTCGAGCTTAAGGGATTAATATCTTGCGTTCTGCCAGACGTGCGCAGATACGTACTCTATAGCCGGTAACCATCTGATCTACTCACTGCGCACAAATGTGCGTTGCCACTTCCCGAAGGAAGCTGTTTCGACGAGATTTGACTGAAACTCAGGCTCCCGAGCTCTCGTTCATCCAAGGCGTCTTGATTCATACTCCGAAATCTTAAGCGTGCAATTCTAAGCAATCGTGCGTTCGGGGGTCCGAATTCCGTCTGACTGGAGATCCCATTCACAAACGATAAAGTATCTACCTCCGGTCCTCGAACCGTTGTTCGGTAACATGGTGCTAATACTGAACTCGTGCATGCTTCGATTAGCGGGTCGCTTCAGACCTCAGACCCTGTTCCAGAGGCAAAATAGGCTCATACGCTATGATCCCAGTTTTCACTAAACGCTGTTTAGTGTAGAACATTTCCGGTAACTGAAGTTGACTGTGACCCGAGCCTA |
| **E18.BC59**<br>Inh-NPY.0<br>(spinal cord) | GTCTCCAGATCTTCTCAGCACCCCCATCCTCCACCCCCACTCCTGGCTCAGTGTTGGCGTTCAGAAATGTTAAATAACAATAATAATAATAACACATCTGAGTGTGACGCTTTAATAGCCAATCTGTCATTTTCCTACTTTATTTCTGAGATTTTAAGAACAGAGCCCTTCCTGCACAAAACATCAGGGTGCATTTATATTAAAAAATGAAACTAATTAAAATGTTTGAAAACCATAAAGCACACCACCATAACACAGCAGCAGCCCTATTTTTCCATTCGAAAAAGATTTTTCTTCTTCATTTTGTGGCCTGCCCTGATGAGAAAACTGTTGGCGCTGGGAGTCACTACAGGTGTGAGGCTGATAGGAGCCGTTTTGCCCCTGAGCACTGCCTTTGCTCCTAGACCCTGGCATGGCGTCTGAGGCCACAAAGCTGGAGCAGCCAGAGGGTTGGACTTGGATGGAGAAGGCTGGAGGACAAGCTACCCTCCAGGTAATCA | | AATATCGATAAGTTACAGTCTCTTTTCGTCAACGTTGGCAAAGGATGCTAGGCGTATCTCTAAATGTCTCGGTGTTGAGTGCAGCTTCGAGATTTGATATTTGGGGCTCGCGCCGGGTTAATAGAGTGCGTAAGGTTTATTCCCAACTCTTTGCGATGTTAGCAGACGCATAGTCCCCGAACGTGGCGGAGTACATCGAAGTCAGTGGCCGTCCTCTCGGACGACCGTTGCTTGTCCTTAGTGTTAGATGACCTGTTTGCGTCGACGGTGGGCACGTAGCTATAACCTCTGCTCGCAAACAAAGAATTAGTCGGACCGGAATTGATCGCTCACAATAGTGAGTCCTTCGACGTTTGTTGCGTACCACGCATGCTATCCGCGAACCGGCAAGCACGCGTCTGGCTACAGTTTCCCGGAAAATCAACATGTCAATACGCGGACTAAAGGCGCCCACAGGCTAAGCGTGTTAAATCTATATCCCTATGTATAAATCGGACCAG |
| **E19.BC60**<br>Inh-NPY.49<br>(spinal cord) | GTCTAAGGCACATCTTCTCCAAATGAGAATTGTATTCATTTTTATTAGTAGTAGGATAGTTAAATCACCTTCATAGAATAACTTACGGGTTTTTAAAACAAATCTGAAAATTTATTTGAAAGTTATTAATGCTAATTTTATTATTAAAATTGTAATAGTCTTACTCTAAAACATTGATGATATGGTGCATGACCAATATTATCATAGCTGCAAGATTAATTCTACAGTCACTGTATTAAACATTGGCAACAGTACTAAATAGACACACTCAAGTGGCACATTAATGCTATTGCTAACATTTTTGTCTATAATAGAATTTGAATAGGAAGAAAAATTTAAATTACATGTTTTCTAAAAGAAGGTAGAAGCAGGGATTGTAAATTGGATAAATATAAAAAATCCTTCAACTACTGAAAATGGTTAGCAGTGTACAAAATGATTTTGACAAGCGTGGTGTTTAAAATGGTAACTTCTGTAGACTCACACCTATAAGGAAATGT | | GACTCTGCGAGAGGTACACTCCCCACCGGGGGCCAATTGCGAATGGGTCAAGTGCTACCATCTATACACGGGACAAGAGTACTTGCAGTCTTCTCCCTTCTTACTATGGCACGGCTGATGGTATCGGTGGTGACAGTCTACCACATGCCATAGCATAAACTATAGGGTGGACGATTATGGCTAGCTAAGTGGAGTCGTGAGGGTACATAATACAAATACTAACCCCTTATGGACAAAAGCGTTAGAAATCCCCACTATCATCTGATTGATGTATATTGTAGAGAGACAAACACGCCTGGGCGCCAGTGTGCCCCGGAGCATAGCCACGGTAAGACCTGATGTATAGAGGAAAGTACATATACATAAACAGTTCGGGACGAGGCAGTCCTCGTATAGGGTAGAGTTTGTGCATTGTTAGTCCGTGCCCCTGAGGCGGTGATATGATCCAGTTAATAGTCTTTCTGCCGCGCAGGGAAAATTCAATGACATGCGAAATTAGG |

| ID | Enhancer | . | Barcode |
| --- | --- | --- | --- |
| **E20.BC61**<br>mDlx-prom (500bp): Inhibitory neurons | CTTTGGGTTCGTAGCTCCCCAGGATCAATTCTGAACAAAGCCTCCAGCTGCAGTGCCATCCAATTTGAAG<br>CAGACATTGGGGACAATTTAAGGTTTTTATCCACAAGAAGGTTTTTTTCCATTCTCTTAAATGCAGCCATAA<br>TTAGAGTAATTTTTCATGTAGCCCGCTGATTACAGCGTTTTTACCGTCAAAGATAATTACCTGTAATTTTCTT<br>CCACTTTTAATACTAAAAAGCCATCTTTATTTAGATTCAGGAACAGGAAAGGCGAAACAAAAGAGGGAAA<br>TTATTCTGTTATTCATACACAAATTGCAGAGACATAGGACCTAAAATTGAAAATTAACCAAAATTATAATGC<br>TGGAAAGAATGGAAGAGGCTGCAGAAGTACAGCTGGTAGTAGGGCTTTGCTGAATTATTCAAATTATGT<br>TAGAGAAAAAGCTACCATATATCGAAACCAACACTAATTTTTCCTTAAAAAAATTTCACCAAATATATGCCAA<br>AC | | AACTCGTTTGGCGGACGCGCATCGTATTCTGCAAACCGACATCCGTCCGCCATCCTGCCGCT<br>TAGCCCACTGGCTGGAGCCACAGGTTGCATGACTATAGGAGCCACGCGTAATTGATATTTTG<br>CCCTTCTTCGGAATAAATGTAGTAATCGCTAACATGGCCTTATAGATCTATTGGCTGAGGCTG<br>CTCGATTTTGTTTCTACCTGGTTTCTATGGGCCGTAGTTAGCCACACACCGAGGGTCAAAGCC<br>TCCCTGAGCCGGTACAAAGGAGTCGCAGTCACACCCGCCGGATATGCATGGGGAGGCGAA<br>GGGTCATCCTGATTTATCCGGGACCGCTTGGCCTAAAGCCAGGACTGAGGATACGCCCTAT<br>GACCGCGGCGCTTCCACGCACCACTGCAACTTAGGCAGAGAATTTGGTTAGCCACAACTTG<br>AGATACTAAGTACTCAATTAACCTGCCAAAGTTACACTACTCCCAGCTCAGTCACTACGCTGG<br>ATGCC |
| **E21.BC62**<br>MN.098: motor neurons-Harvatin (spinal cord) | GGCTTCATTTCTGAGCTGCAAACTGCCTGGGTGTCTGGAGAAGCAGGTTGGCGTGGTGGTTAGCAGTG<br>CGTGGGCGGGGTTGCCCGCTCTTGATTTATGATTTCTTTGTCTCTGTGGAAGCACTTAAGTGCAGGCTTT<br>AGTTCCAATGACACTCAGGAGCCTCTGGATTCCAGCACTGGGGATGGGGGTGGGGTAGAACGTTCTCA<br>GGCCTCACCAACCCCTCCCCTGTGTGCTGCCTTTGGGAGAGTCCCAAGGCTTCAGCATTACTTAATTAAT<br>TAGGCCTCTACTGCTACATAGGCTCAGATTCAAAAGAACAGAGTGGCCCACGTCAGCCATTCCCGGAAA<br>AGTCTGATGGCTGGAAGCCAGAGGACTATGTGTCTGCCTTGCTGCCCTTGGCCAGCCCATCCTGAATGC<br>CCAGACTCGGACAATGGAGTAGGTACAGAAGGGTAAAGACAGTGTCTTCTGTACCAGTAAGTGGGCCC<br>TGATCTGCTCTCTACAGC | | CCAGACCTCTTGTCAGTACAAGTCAACCTCCTATCGACCGAAGAGTTATGCAACACCTAGCG<br>GGCCAACATACGGTCCGTGTAAGCAACAAATCCCCGAAGGGATCATTGAGTGGCAAATGTC<br>CGCATACGATACCCGCGATCACTGGATTAGTCCCCGAGGGACGCTCCAGCATCTAAAGATTA<br>AGATCCCGAAAGAAACGGCTGCCTTGGATCGATGCCGAACACCTTGGCAGGGACGGACCT<br>ACGTACACGTTCGCTGCGTTGATCTCCGGATAGGGAACGTAATACGGGGACGTCGTTGATG<br>AAAATATTAGACCCGAACGAATGCTTTGTCTATGGGCACTGCAGCTTGCGTGAGGTCCCATA<br>GTCGCTTGAAACGATTGGTCCTCCAATAGTATTACCGGCCTTACCTCTATCGTTTCGGACCAAT<br>CCGGGTGCTGCTTACGGTATGATTCAACTCAACTTGCGAATATGCCATCCCAGGGTACAAGT<br>GTGTGC |
| **E22.BC63**<br>Pdyn-prom (500bp): Inh and Exc PDYN (spinal cord) | AGGACTGTTCAGGGCATGGTTTCCAGAGGACAGGCAGCTCTTTGGACTCTCTGAGCTTCTGTAAGCTCA<br>CCTTGAAACTATGATTTCCCTGTGCTGAGTAACCAGACCCAAACCTAACTTCCCCCGTCCAAGCTCCCACC<br>GGCGCCCCTGCATAGCACCAGGAGCACATGCATAAAGCCAGAGAGCCTGAGCGCCCTGCGGATCTTA<br>CCTGCTTACTGCTTTGTCCCTGGCAGGCTTCTGGCTGATCAGGTGGTAGGGTGGCCGATCCAAGATTCA<br>GAAGGTACCCTGACTCTAGCTCAGAGAAGTGCCACAGAGGAAGCGACGAGGGCTACCAAATCCCCCT<br>ATGGCAGTTCTGGAGCCTTATTCCCCTTTTATAGATGAGAGTCTCCCCCTGTCGCTCAGCCCTCTCAGCG<br>CTCGCTCACAGAGCACAGGCTGCCAGGAGCCAATGGGGGAGTGGCAGAGCAGTCAACGCTCAGACC<br>CCCGGGTCCTTCCACTCAACT | | GAACAGCCATCCACCGGACCCACTTATTTCGAACGTGAGTGCTCCATCAGAATTTGATTGACC<br>AAGCAGACTGCGATTCCAGAGAATAACAAGCTGTAACAGTTCCGGAGCCCCATTCCCCCCAT<br>TCACGGATCTCTATTGGGAATTGGGGATATCCGCACCAGTATTAAGTGAGGGTAAGGGTCA<br>CTACAATCTTGTCTGGACATCTGACGAGCTCGTACATGGCACTGCTTCACGGGGGAAGCAAC<br>TTTCTAATTGCGAAAGGTCCTCCGATACTTCCGGAGCTAAGCCTCATGAGGAATCGACCTGG<br>ACGGTCAATTTTGGGATACACCCCTTGCGTAGAGACAACGAGAAGAGACACGTCAAACGCA<br>GGCAAGCGGCTGTCGGCAGAGAAGTTTACCTGTAGAGAAGTTAGAGTAAAGAGGTTGCGT<br>GCTCGATCAATCAGTGTCACGAGCGTAACCCACCAGTGTCCGTTGTGCGTAGTCGTCCATTC<br>ATCAAAG |
| **E23.BC64**<br>Npy-prom (500bp): Inh and GABAergic (spinal cord) | AAGCCACTGGGACCTTGCGGTCCCGCACAGCTGCACCGGGAGCGGGAAAAACCTCGCTCGCCTGCCT<br>CCCTCCCAGAGCGCGCAGTTATCCCGCGGGCCCCAGATATGCTCCCCAAGTAGAATATTCAGGTCCCTA<br>CAGACCCGCACGCAGAGAGCTGACAATTCCCGCGACAGGCAACCTAAGCGGTCCCGCTTCATCTCCCT<br>AGCGGGACTCGACGGTCGGGGATAGAGAGAGGGCTCCGAGGGTGCTCACTCTTGACTGTTCCCTTCT<br>CCCCGCAGAGCGGGCCGCTTAGAATTGGGGTGTGGGTGGCTCCAGACGCCGCCACTCGAGCGGCAG<br>TGGCTCCAGCCTCCTCCCCCGCCGCTGGGGGCGGGAAGTGGCTGTGGGAGTCACCCGGGCGTGACT<br>GCCCCCGAGGCCCCTCCTGCCGCGACAAGGGCGCTCCATAAAAGCCTGTGGGCGACCCGCTCTCCGC<br>ATCCCACCGGTGGATCTCTTCTCTCACAGA | | CTTTAAAGTCACGATTTGGCTGTTAAACACAGAGCGGTGGTGGAAGGGTCAACCCTGATGG<br>TCGTATACCTCCCAGCAAGTCTTACGTAAACTGCTAGGTGCTCGCTCGTAAGCTGTGCTGCTG<br>AAAGTCGACCAATCGGGCGTACGGACTGCGTCTAGGAATATCTACTATCCGGTTTTGGGGA<br>ATGATCTGATGTTGGGAGTAGAAGTGATTTAGCGGAAGGTGGGTATTCTGGACTGTTTGCTC<br>TCGCCATCCGCATCCCATGTTTCTTGATGCCATTACTTGACGACCTCTACCAGGTGGCCTTGG<br>CCTTGTCTTGAACTGTATAAAATAAAAAGGCAGTAAACCGAGTAAAAACCGAGGTAATGACC<br>GAATGGGTGGCCACGCGAACGTCCCCCATTAGCAAGTAAAGGACATTCGTACATCCCGGGC<br>CACCCCATCTCAGTCCGTCCTTGAGTAAAGTACTGGTGAAGGCCTCAGAGGAATGAACGTAT<br>TGCCG |
| **E24.BC65**<br>MN_Hb9: motor neurons (spinal cord) | AGAGTGGTTAGCTGATGAATTGACAAAAACTAATCAGCTTTATTGGGAAACAGGTTTAAGGGCACGGAC<br>GTGTCAATAACGCTCAGCCTGACCCCCTCTTCCATTAGCTAGGCAGGCTGATTAGAGTAAGAAGACCCT<br>CTCGCCGATTGCACCGAAATTGCTTTCATAAACTCACAATATTAGCAGTCGATGTGAGCAGCTAATTTAGA<br>GAAGACATCCAGGCCCGTCACATCGATCATCGGGTTTATGTATCATTATCTCATTTACAGAATGTCGCAGT<br>GTAATACATGTTTTTGGCAATAGGCTTGGCCGCTCGCCGGAGCCGCGCGCTGTTCCTTTCGGCCACACT<br>GCAGGGGACTTCCTCAACAATCTACGATGTCTCTTCGCGTGAACAAGGAGGCACTTTGCACACATATGC<br>ACACTTGCGCCCAGGCTCTCAGTGCGCGCTCAGTGCACACATGCACGTGGACACATGCTCCGACATACT<br>ACCTGAGCTGCCT | | GGTCAACTGCACTATCTATTTGTTAGTCGATACAGGCGCTCTAACAGCCATTGGGGCCCTACC<br>CTAGAGGTTGCGGGCCGGTAGTTGCCTGTGTTCAGCCTGTTCCGCCAATCACGTATTGCTAG<br>AGGTAAGGGATTCCAGCTCTGCACTTTTGTAGCATCCCTCTAGAGGTGTTTATAAACATAATTA<br>CAGCGATAGGTCGTCTTGAGTCTTACATTAAATCGTTCGGAGAGACATTTGGCATTCGAAATG<br>CCCCTGACCGTCCGGTCCCCATTTGCGGTTGTTATACCCTGGCAGCGAAACCGACTCTATGC<br>ATAGATGTCCTGTATAACATTATAGCGCGATTTGACTACCCAAAGCAGGATTATGGTTGTAAA<br>CGACCGGTATCAGGATGTCATGAATCCTGATCGTCTTCACAAACTCACTGCTGTCATCTAATC<br>GGAACCCGCATGGGGTCTACTCTTAGTACATCATGATTGTAAAGATATACTGCCGAAACT |

| ID | Enhancer | . | Barcode |
| --- | --- | --- | --- |
| **E25.BC66**<br>mm_oligo.18: oligodendrocytes | ATTAGATATTTTCTTTACTTACATTTCAAATTTTATCCCCTTTCCTCATTTCCCCTTTGAAACTCCCCCATCCCA<br>TCCCTCTTCCCCTGCTCACTAGCCCACCCACTCCTGCTTCTAGCACCACTCCTTGAAAAGACTTGCTCCAC<br>AGATCTGCTTTTGTTCCCTTGTCAAGGAATAATTTTCCACAGAGACTTTGTTTAGGAACTCACTATTCTCGT<br>CTATTGATCTTTGTGTCTGTCCTTTCAGTAGACACTAATACAATTGTTGCATCTGTCCAGTGAGCTTCGAAG<br>TGTATTTGTTTCCATTGTTCTTTTGCAATACTGAATGTCCATTATATAGGATTAGTGTACTGTGCTGCTGATC<br>TGCAAAAAATGAGCAGTAACTGATAACTGAATTCTCAACATGGATTTTCTTGGTGACCTGTCTAGAGATAC<br>TGAACATGATACTCTGAAATGTTTTCTGCCCTAGGAGAAGGAATGAGGTGATTTGTTATGGGGTGA | | CCAACTTGATAAAGACTCTGAAGGTGTATGGGTGATGTAGAAGGGCCATCCGCGATGTGAA<br>GTGTCACTGTGTCTATTTCCCCCCATGGACTCAAGTATGGGGTTCGGACCTGCGAGATGCCC<br>TTGCTGAACGCTAACGCAGCAAATAGACAATGAGTCCGCTACTGCGCACGCTTGAGCCCTAC<br>TGACGTCAAATAGTTCGAAGCTATATCTGTCGGGAGTCTATCGATCTCATGATACAGCCACGC<br>GACATAACCTAGCCTCTTTTGGTGTCCAGTCTTACACCCTCAGAGAGGACACCGCTGCGTGG<br>CCTATCCGACCGGCGTCGTATCTATGGTGGATGCTACTTAGAGAAAACGCCGCCCAGAAGA<br>GATTCACTTAACGCCTAAGGCGGGTGCAGCCGTTGCAAGAATCAGAAATAGGCACTTAAAG<br>AGAAGTATACCATCTTTCCCCATGTCAAGCCATGTTTTGCGGTCTTTCAGCCTAATCTACGGAA<br>TAGC |
| **E27.BC67**<br>hu_oligo.41: oligodendrocytes | AGGTCAGTCCTGGAAGAAGTAAAACCATGCCAAGCTGTCTTCTACCCCAAATGCTTGTCCTGTGCATCAG<br>TCGCCATTCTATTGTGTCCCTGAGCCTTGAGCGCACCTCCTTTCTCCTGCCCAAAGTGGGGGGTAGAAG<br>AAGCCCCCTGCATGGCAGCCTGCTGCCAGCACTGAATGGCTCATGCTTAGATCCACTTGGCACACTGGC<br>ACACAGCATACACAATGTCCGCTGTTGTGCCTACTGTCCCTGCTTGGTGACAGTACCGCAGGGAACTCT<br>GTCCCCATTGTGTCTGTCCAGTGCTTTGTGAGCATGTGTTTGTTTAGGATAGAATGAAGGGTTTTTCCTCA<br>CTCATGAGACTGGGAGAAAATCAGGCAATCTAATAAGCCTATAGTCTCTTCCCTGCAGCACTCAGACCTG<br>CAAGACACAAACAAGACCTTTCTTACCCGCCTGGCTTCTGGCAGCTGCCACAGAACCTGCATGGTGGCC<br>CTGCCAAACTAAC | | CCCTGTAAATGTGGATAATGTTGCATTGACCGGACTCAGGAACGCTTGGTAGAAGAAGATTA<br>ATCTGACGCTAACAAGGACTAACGCCGGAAAGTTTCTCAAATCGGTGAGGTGCGTGGGCAT<br>CTATCAGCCCTCTCTTCACTATGAGCGGGCAGCACTACTGCGCTTGACATCCACGGGTTGGA<br>AGTCATCCAGGGGGCTAGCGCTGTCCACAGTATACCTCCGTGAAAAGAGTTTCAAAAGGTC<br>GCGGGTACATTATACATCGAACAGACCATGCGCCCGTTCGCTCTATGGTGGTCCATTCTCAC<br>CAATTCTCTTCGGACAACAAGGATGTGTCACTGAGTAACACTGGACGATTCGCAGCGTAGAG<br>TACGGTTCCGCCCGTCCTGAAGTAAATCTGTTTGGCTACCGCCGCGGTGCATAGGCAGCGC<br>GGATTGCGAAGGCCGTAAGGGGCGATTTGAAGAGGATGGTAGGAGTATGTCATTAATTATC<br>TGGATCTA |
| **E28.BC68**<br>MN.187: motor neurons-Harvatin | CAAAGCAGCTGCTTCCCACACTCAGAGTGCAAGAGCTCACACCTCCCCCAGCTCTGGGGAAGGACAAT<br>GACATTTTGAAAGTACTGTAGCAGCTACTCACTGTTCCTGAAGCTTCCACCCTTCTGCCCTGAACCCTGAA<br>CTCTTTTTTCTTTGCTAGCAGTTTTCCTAACATCATTCAAACCAGATGGATTTTTCATCAATTATGTGACATAT<br>CACATTGCCCTCATATGTCATAAATTTCATGGATTTCTCAATATCTTGTCATGATGAACCCAAATTTAGCAAT<br>ATTATCAGACAGTGGCTCTGCATATCAAGACACTTAATGTCTGTGCCAATAACCAAGATTTATCATTTCTTG<br>AATTTAACAGCCAGTGCCATTGGTTCTTTTAAATTGTTTCAGCGTGGGCTAAGCTTTCAGAAATAAAGAAA<br>TTAAACTTTTACTGCTCTCTCTCTTATCTCTCTCTCTCTCTCTCTCTCTCTCTCTCTCTCTCTCTCTCT | | CATCTGTCTCACGGGTGATGACGGTCTTAGTAAACGGTTGCCGGGGAGGATTCTCAACATCA<br>CTCCTCAGCAGATGACGGGCACGGACTACTAATCACACGTCTTGGGTGAGTGGTGAAATGA<br>CGACCAAACAGATATTTGTATCGACATATAGTTACGAGCAAATCGTTGTGCCCAGCCGCAAG<br>GTTATCTCTCTGTACGTGGATCAGAGATGGGCAGATGGACCCTCTTCTGAGAACATATCCCT<br>GTTTCTCGTGCGAAGGTCCGACAAAGTTGGACCCGGTGCAAGTTTCAGCGACTGTTCCCGG<br>AGAGCATCCTGACGCTGGGCTGGATCCAGATCACTAAGGCGGGTGCCGCTCTAAATGCACT<br>CACGAAGCGATTTTTTTGCCTCAGATTCGAGGGTTTACAATCGAGCCAGCCCGGATATGCTA<br>CTTTCAGAGGACCTCCAGAGTGAGTTAGAGGGGTACTTCAGGTTGTTCCGTTACACCACCGC<br>TCCAAAG |
| **E29.BC69**<br>Inh-PDYN.5 (spinal cord) | CATGCTTCCTCCTCCGTGCCCTCCTTCCATGCCCTCCAAGGAGAGCACGTAGGGCCGTGGAAGGGGCC<br>TCGGCAAGTAAGGGACCTGCTGCTTAGGCTGCATCAGCAGATGGCATCTGCTTACAAAGCCATCCTTGG<br>CTGTCATGACCCTTTCTGAAAGAGAAGACGGATGAGGGATGTTGTTTTTCTTTCCTGAGCCTATAATACA<br>GCAACAGAGATGAGATGTGTAAATGCCTAGCCTAGAGTCTGTCTCATGGTAACTGTTCAGTGACTGGCA<br>GCTACTATGGTTATTAGTAAATTACCATCTGCTTTAATCATTCTCCTGTGAGTTTCATCTGAACCTATTAGTTT<br>CCCAGAGCCAGCTTGCCCTGGGGAAAAGTCACAAAGCTAAGAACAACTTTAATGTTGGCCTCCAGACTC<br>AGTCAGATACCTGTTTGCTAGAACACTTTGAACCAGGGTGTGGGTCCTGCTCCATGAACATGGTGTGAA<br>TTACCCTGTAAA | | CATAGAGCTGGCTCCCAGTGGAACCATGAGACTTTTGAAAACCATCTATAGTCCCGGGCTCT<br>TCGACAACTCGGAGTAGAATATACACCTGGACGCCTCCTAGGCACAGATAGTAACCCATGC<br>GCAATATTTGCGGTGACGGTCAGATTGCTGGATCTCGATGGGAAGTCGTATGAGCGGTTGT<br>GTGGTTATTGAGACTTAAAGAACCAGGGGCCAAGTTTGGTTGGGACATCAGAACAACGGAA<br>GATGTTCTTAACCTGATATAGCCCATTCGGTAGCACTCACCGGGAAACACCGGTACGACCCT<br>AAGTGAAATACGCTCATCTGACGTATTGCCGATAGGTTATTAAGTGATCGCCGCTATAGCCCT<br>GGCCCTACTCACGCAGCACAGGCAGTACATGACGTAACCTGTGAAAATCATGGTAGCCAAT<br>GAAGTGGCCGGACGTGAGAGGTAAAACCCCTCAACCTGGAAACTCAGCGGATTTAAGTCC<br>GTAGTCAAA |
| **E31.BC70**<br>hsyn-prom (500bp): neurons | AGTATCTGCAGAGGGCCCTGCGTATGAGTGCAAGTGGGTTTTAGGACCAGGATGAGGCGGGGTGGG<br>GGTGCCTACCTGACGACCGACCCCGACCCACTGGACAAGCACCCAACCCCCATTCCCCAAATTGCGCAT<br>CCCCTATCAGAGAGGGGGAGGGGAAACAGGATGCGGCGAGGCGCGTGCGCACTGCCAGCTTCAGC<br>ACCGCGGACAGTGCCTTCGCCCCCGCCTGGCGGCGCGCGCCACCGCCGCCTCAGCACTGAAGGCGC<br>GCTGACGTCACTCGCCGGTCCCCCGCAAACTCCCCTTCCCGGCCACCTTGGTCGCGTCCGCGCCGCCG<br>CCGGCCCAGCCGGACCGCACCACGCGAGGCGCGAGATAGGGGGGCACGGGCGCGACCATCTGCGC<br>TGCGGCGCCGGCGACTCAGCGCTGCCTCAGTCTGCGGTGGGCAGCGGAGGAGTCGTGTCGTGCCTG<br>AGAGCGCAGCTGTGCTCCTGGGCACCGCGCAGTCC | | TGACGGGGGTTAGAGAACACGCATGCTCTGTCAATATCACAAATTGTTAAACAGCACCTGTG<br>GCAAAACGGCGGTTCTAATTCTCCCCATACGCATCAACAGCTGTCTCGAGGACTTGATTAACA<br>TCTGAGCGACGATGATACCAAGACTGGTTAGTGCGACCGCTAGCGACACGAGGAACTATTC<br>CATCGGGTACACCATGTACCTGCGTGATTATACTTTGATAGAACAAAATATCTCAAGCCCTCCT<br>AATCCCGTCTGGTGCCCCGCAATGAAAAGGCTAACGAGGATTATCTCCCAGCGCGAATATG<br>ACCCTAATAGGATTGCTGCACGAATGAAACCTATCAGTATACTTTGCCGTAAAAGAGAACGG<br>TGCTCTCACCACGGCACGAAGCAGGAACCCTTAAGAAGAGGCCATCTGATGCAACATTCCT<br>GATCTGCCTCGGCGGGTTGGTTACTACGTCGCTGCCTTCGTGCCATTCGCCGTCGTTACTCT<br>GCGG |

| ID | Enhancer | . | Barcode |
|---|---|---|---|
| **E39.BC71** Exc-SKOR2.8 (spinal) | AGAGCCTTCAGGTGTCTCCTTCTTACTCAGAGTTGCTGAAAGAAGTAGGAAAACACATGGGGTTCTATATACGTTTTGGGGATTGCTGATGTGATTTATTGAAGATTTTCGTAGCCTTGAAAAATTCAATTTTATACAAAATCAAGTATTTTGCATAATAAAACTGGTAGAAAATGTGCATATGTTTTACTTTTATGTTCCGCTTTTATGAACACTGTCAGACTGAAAACTGAATTTATGAAGAATTTACCCAACACACCAAGAAAATTGCTGCCAAGAAATCTTGTATTTACACAGTGTGATAGAAAGAGGCTGAGAGGCATTTAACACTGCAGTCCAGTTTATATTCAGCCACACTCTATATTTAACTTTTTCTTTTCCAAAACACGAGAAGCTAAAGGAAAAAATCTTTATAGAAGAAAAAGGCCTTCTTTGAGATAAAAATTGTATAATTCAGAATAAAACTGCAAAGGAGAGAGAAAGAATAATCTCAAGACGAT | | AGTTGAATGACATCAGCGGCCGGCCCGGGTAGAATACCAATTCCAGAGGGTCGCTCCCAAGCTTGAGGATTCCTTTGTTGCGGAAGATTTGCAGCCTTCGATCTAAAACTCTCGTGGCAGTTCAAGAACATCTTCGAGCGTCTTGATAGGTAGACTAGCATGGATTGTTTTTTTTGGTTACGGACGGGGGGTAGAGCGCTTCATGTCAATATTCAAATAGACTATATCTAGGAATTAATTAATATCCCGAGACTGCGCCGTATGTAGTATGTTGACTTGCAGCAAGAATGTTTCAGCGGAGTGCGGGCTAACATCTGACCACCACGAGTACATGCTCCATGATGAGGGGTATAGGGGACACCAGGCAATTGCGAGATTTTTATCTTGTGAGGAAAGCAAGACAGATGCGAACCGCGGGAGTTCTAAGACCATATTATTGCGTGGCCAGGCTGTTACAAAACGTTGCATGTTTGACAAGGAACTTTCGATCGTCGCGTAAG |
| **E60.BC72** Exc-PDYN.0 | TTGACCATTTGACACACAGCACTGCAATGGGTGATGGGTAGtttttttttAtttttttttcttAAAGTGTAGCTTGTTAAATTCTTACTGAAAATAGACTCCAAACTACATTCTTGGCAGGGCACGAGAGAAAGGCCTGACTGTTATCTGCTGAAGCCATGAAACTGCTCTCAGGAAGCTGTTGGCTGCCAGGAGAGCCCCAGCTGAGGAGAGCGGCTGAGAGAATTACCTCCTCGGGCCCACAGAGGGAGGGAGGCCTGCAGGTCGAAGCCAGCCAATGCTCAGAGCAGCGTGGGCGGCCTGGAGGGTCCACACAGCACTTGGCTACCGAGTGGCAGCATCATTCCTAGCAGGCCATCTTCTCATTCCACTTGTCTGGCATCGGCAGGGCTGCACAGAAGCCCAAAGACTGAGCCAACTGGGGTGGCAGAGAGGAGCAGTAAGAACAGAATGAAAAAAAAGAAAGAAGAAAAAACTCCCTCTAGCTAAATGCTATCCTAC | | ACTTAGGTTCATTGGGCTTGCCGTGGGACAGAGTTCTAATTGGCGAAACGTACCCCAGAATCTTCTTCGCGGTCCACTTTTAACTTTCGAGTTGCTACAGCATATAGCGATTCCCTTCTTTAAGCCCCCACTACTCAGACATGCGGCAATAAACGCGCAAAAGGCCTGCCAAGGTGTAGTGTTCTATGCCCGGGAAATGGGGGGCGTTCTGGCATAATATGAGCCAGCTAGCTAAGGTTTTTCGCCAACCCACCTTGAACACGGTGAGGACCCGGTCCGCTTACTCTGTTCTGAATCGTTACGTTGCCCAAAGGCATATTAATTCATTTCCATCCTTTATCTTTGAGTTGTCGGGCATCCCGCCACCTTGTCAATTCTCAATTCGTTGTAACGCTAAAGACACCGCCCTTCTCTCAATACTCGAATATCCTATCTCGTGAGACATGCGACTATCTGCTGATTCAAATGAAAAGCGTCACGAGGACTTCCCTTCATCGCAA |
| **E41.BC73** Exc-SKOR2.55 (spinal) | GAAGATAGTGTTTGTCAAGGATCAGCAACAAAGCCTGGTACATAGTAGGCAGTCAATAATTACTATATCATATTGATTCTAAAATGCACACTTATTTTCACATTTTAACATCTCTGAAATTGAAATGGATCTATATTGTGTTATGGTTTAATGACAGTTTTTTCTCTTTCTTAATGTGCATAAAATAACAGTACATCTTATAGTCAGTGGCATCTTAAGTTGATGAAATGCAGTAATTTCTTCTCTCTCTTACTACTGTATCCAAGAACATATTTTACTCCCTCACAAACCTGACAAGTAACACAGAAAAAATAAAAGCAGCACTATAACAAGTGGTGCCTTACAACCATTTAGAATAATCAAGAACAATTTTTTCCCATTTCACCTGACTATTTCACTTTTAGAGCTGTGATAAGAGCAACTGAGATTTTTGTGACATAACAGGATTGTTTAAAAGATCTCTTTTTGCCTTTTTCACAGACAGGCCTTGTTTAAATTAA | | GAAAATGCGATCACTTGGCTCGTCGGGTTGTATATTAATCAGTATACGCAGGCGTCACCGCGGTTAATTCTAGGTATTGTGTGGTTTTTAGTTCCCTATCTGCCCCTCCTCCCCGTACGATTTTATCTCCCCCGTCGGTCGTGTCGATGGTTTCAGGGTCGGCTAGTCTGAGAATCGTGTGCTGGAATATAAGACGCCAGCCCCCACTCCAGTCTAGATGTATAGTCTTAACTGGGTAGCCAATTGTGAAGTAGTGTGTCCAGGTACTAAAAACTCTCGCGCACGGTCCGGTGGCTTGCAATATCGAAAAGGTGTTTTGACTGGGAGACAATCCTCTCAGTGAGTTAAACGAGTAATCGACGGGCTATCATTATGGCGTAGCACTCTATACCTACCATCCCTGCATCTAACACATACCAGTTTGGAAGACTGTTACCGTGTAACTGTCACCTCGACCCCGTCTGCCGTATAAGTTCCTCGCCATACAAAATTAGGCTATA |
| **E42.BC74** Exc-SKOR2.113 (spinal) | AAGATTATTGGCTTTCTGACTAGTGCCTTAAAGCTGGAGTTGGATATATCAAAGCGCTGTTGCATGATTATTTATGATTTCATTTTACTGCAGTGTAAATTTTTGTAATGTCAGCATTTAAGAAATATTATTCCCGTCAACTTTACCAAGCCCTGAGTGGCTTCTTATATATTACTTTTCAGCATATTTGCTAATTTCCTCAAGCAGTGCTAGATATTCACTTTGGAATTTGCCTTTTGATGTCTCTTCTGTGATGAGCAGCCACCAAACAGCCCCCACCACATTCTAATTCTTTATTGCAAACAGTAAAATATGGAGCACTGCATCTGAAAACCACTGTGCGAGACAATCTCATTTTTACTTTCTTTCTGCTTAAATATGCAATAAATATGCATTCTGATAAGTACAGTTCTAGAATAAGATAGTTTCGTGTAACATCTTGGAGTAAAAAAGAGAAAGAAAAATTTTTTTTTTTTGAGAAGGAGTCTTGCTTTGTTCCC | | AAGCCGTGATTTGAACAACTAGATTGGTCTTTAACGTGCTTTATTTCTCTAACCAAAGACTTTAAAATTGGAATGGGTGTTACCAGTGCTTAGGCGGCCTTGCTTTCGCACGCTCCCTCGGCACTTAACCAGTCCGACATAGGAAGGCGAGTGAATAGGTATCCGTTTTAACTTGATGGACCCCGGACCCACCCCGCGTATGACCTTTAAATTCGGAATCTACGACTCGTCCAAAGAATCCGATCGCGATCTCAGGGAGTCCACCATAGCTCTACCTAGCTCCAGATCTAGCAAAGGGTTTCAAAGTGACTATCGACGCAGGAGAGGTCTTATGTGTCCTAATGCAAAAGTGCTCAGAGTAGACCACTAGAAGCTTAGCCATGGGTCCAGACGCTGTAGTCACACTCGCTAGGCGCTCCAAAAGGGAAGTGTATACAAATAGCAGATAGCACGCAACAGACTGACAAGGTGGTCTATTGTAATGACTCGATATATTCTAG |
| **E43.BC75** Exc-SKOR2.103 (spinal) | ATGTAACTCAGCTGAAGAATCATTATAGGATTTTTTTAAAAACTGGAAACTTCCCCCCTGCCCCCAGCTGTCTACAGAAATAGATTATTTCTTCTAATTTCCTGATTTAGTTATAGTCATAGTGTACATAAAAACAGATTTGATTTTCTTGGAATCAAACATAAAATGAAAGCCAAGGCCTTATCATTGAAACAAGACAGCAGTTTTGCAAAGCTGTGTACATCCTTTGAACTCAGATGACATTGTCTCAGAGGAAAAGATGTTTGATGAGTTTTTTACACTTTAAAGCTCTTCCTAAGGAAAATGAGCTGGGAATCTCCTTTGGTAGAATTGAAGGGGGGTGGATCATTTCGTCTTCAAAAGATGAGGGGAAAAGTGTCATTTTCTATTTCCCTTAAGTATCATTCTTTCAAATACTTGTGTGACAAGAGGTTCAACAGGCTTAGATTTGGGTTTTTAGTTTTCCAGAAATGCATGGAGGACTTAAATGGCCCTGTACA | | CAAAATGTGGTTTAATTCATTCGATCTGAGATCAATGCCGCGGTAAGGGAATGAGTCAGACGGGGGCATCGGGCGGTGAGTGAGCAACAAGATTCGTAAGTTTGACACGCTCCGGTATAGACATGTACTGCGGAGCGCTGTCCCGAGAGTTGCACCGATCATGATCGCCGACCTCTAAATGGTGCCCGTTCCGTACACCACATCAAACATTGGCACATCATTATCGTTACTTCATTAAATCGTTGAATCCACTACAAGATAACCTTCCCGTTAGGTCTAGCCCTGTTCCTGATTCCAGGGATACGTCCATACACAGCTTTTCCGCCACCAGGACAAAGGAGGCGCGGTGATTCGCTCAGGTCGCTCGGCCACTGGCTCGCGATATACAAGCCGGTTTGACTCACAACATTAGGATCACTTGCGTCCGCTCTTTGGTCTTAGAATGCCAATCTTCGTGAAACTTTGTTAAAACGAGCGTCATTGATCTGATATGTGGTCAA |

| ID | Enhancer | . | Barcode |
|---|---|---|---|
| **E44.BC76** Exc-LMO3.15 (spinal) | TCGAGAATTTACTACAAAATATACTAGTGTGAGGTGTTTTACAGAATACTCTTTTCCTAGACTCTTAAAAACAAATCCTTCTGTGTTTTTCCATTTCCCATACAAAGTATAAAAACTTAGAATGGGTGTTCTGAAGTTATTCTGTTCCATGTTCCTCTTCTTAGCTCAAAGTGATTAACAGATATTACTGTAACTAATGAGAACACATTGTGCAGGTCGGAGAATATTAGATAGAAATAGAACATGAATTAATGACATCTGAAAACACTGATGGATGTTCATCAGTTGCTAGGGAAACTTTAGTTTCAGTACTCATGCCAATGAAACAGGAATGCTGTCTATGCACAAGGCAGCTAGGATCTATCTCAGAAATGGAAATGTTGTGCCTGGGAAATGATGATCAACAGGATAAGGCTGTGAATATCAGGATCTTCCAAACAACATAACCCAATAGCTACCCAATATGATGCTGATTGTGTTACCAGAACAGGGTCCTGATC | | GCTGGAGAGTGTAGGATCGCTCACCCCAGCCCCACGACCTGTACGTAACTGCGCGAAGTTCTAGCCATGTAGTGAATCTGAGAACAGTGCAGAGTAGGCTCTTACTCCGTCCCCCTCTCGTGATCGTAGGGACGCGGTCATGCCTCCCGTGTGGGTTGCGTACGCAAGCTTAGCCCTCCCGCCAGAAACCCAGTGCGACGTAACGATAAGACACTCCTGTCATCTTGGAACGCTCGTCGGAGGTGCGTGAGATGTGAGTCTCTGCAAGTTCATCTAGGGTAGATGTGATTTAGGCAACAAGTCATCAGTGGCAGGCAGAATTTTGACCCATCTGATCAGCCTATATTTTCCCCGAGTCTCTCGTCACGCAGAGTCTTACAAAGGGACACTTATGCGGTTGTAACGACAGCTAAGGTGTTTTTTATCACTTTGGACTTCTTGCAATAGTGATACTAATAATTAGCGACGGCCCCAATATATCCACAAGTTACACGTCTAAT |
| **E46.BC77** Exc-LMO3.46 (spinal) | GGTTTGAAATGATGGTGAGAAAGCAAGTAAGTAGTGATAACCTAATGACAGGGACCAGATGGAAATTAAAAGGAAGGGATCTGTTGACTCATCTGCATTAAATGATAGAAGTTATTTTTAGATGAAATATTTTATTTAATTTTAGTGGATAAATTTCTTTTTCAAAAATTAGATTTCTTCTTCACTAAGACAGAACATTGCTTTCAATTCCAATTTTGGATGCTTTCAATTGCAATTTTGGGAGCCTTAAAAGTTTGAAGTAAATAATTTCATGATCATTTATATAAATTCACCAGAAAAACATGCAAATCTCATTGAATATATACTTATAATTTGTTTTATTGCTTAAGAGATAATTTTCCTATTACAGTAAGAATAAATAAATACAAATTTAGATTCATTTAGGCAAATTCTCTGCTCTTACTTGGAGTTAAGGATTATTCATATATCACTGTAGTATTGTAAATATTCTGTAGTTTTGCTATTCAATCTGTATCAAT | | AAATTAGTACCGGATGCTCGAGCGCTTTGATTTTCAGGCGGATTTGGAACTGACAAAGAGGGTGAAATGAGCTCCCATCCCGATAACAACTTGAATTGCGAATACCTGTACGACTGTTCGTCAGTGCGGCATTCGACTATGACTCGACGAACGTGTTCGAGAATATTTTTACGATGCACGGCGGGTTTGGTTGGATTCCCTGAACCTAGCATAAGATCTACATTAATGGAACGTAAAGATAACTAGCCGTAACCGAACACACAATTTCTGATGGACAGTACGTCAAATGAAAGCAAAGTTAGAAATTTGTAGCCCTGAACGCAATTCCCCCGTCACCGTCCCGTATTTCTGCCCTTTACTGCGATTCGCAGTCTTCGCCTCCAGACGCGATGTAGTGTCAAGGAAGTGCACCACGCTTTCGGACTTATATTCCTGAACTAGTCGCTAACGGGGACATTGGTTGAGCAGCAAAGGCGCTAGCCGGGGGAATGACGGAGTCA |
| **E47.BC78** Exc-LMO3.85 (spinal) | AATTGGAATCTCTCCCAGAAGCGTTTCCTGTGATGGTCTGAGCCATACCTGTGCCCGTGCTATGTGCAGTTGGAGAAGTCAACAGCCCCTCCCACCCCACCACCAATTCCCTGAATATTTGCAAACTGTGTTTAGAGAGAAAGAGGAAGCTTTCTTGATAACTTCCTTGTTTATACTAATAGCAAAACTCCAACAACTTTGATGGGATCTGCACCAGCAATCTTTCCCTAGGGCTGATGCAGGATATTTAGGGAAGCTCTGTGTTCTAGGCAGGTAAGATAGCCAGGTTCCTAACGGGAATCAACATATCTCTTTAATTCATATTTGAAGTAAAAGAAACATCTCTCTTCCTCCCATTTGGCTCAGGAATTCATCTCTAAAAATCTATCCTGAGACAAAATTTTATGCCCAAAGATAACTCATGCATTATTTTTAATAGCAAAAAAAATGGAAACAATCTTTGCCCAACATTAGCAAGATGATCGAATGAATTATGGCAT | | ATTTTGAACCGAGCACCTCACGAAGTTGCGATCACACACAGGAGGTAGGTGTAAGTACTGCGCCTGAGAGGGGCCGGAACAATGCGATACCTTTGGCGGCATCAAGGCGGATGCGATTGACAGTTAGTGCGGCTGCAGGACGCTGCGGAAGGTTTTCACTCCAAAATGCGCAGGTAATCTTGAGAAGATAGAGCCTGACGGTCTGCCGTTCTACTTCACCATCGTAGCCCGTATTCAAGGTAGTGGTGCACGACACACAACAACCTGTCTTTCGACCCCTGAGCCAGCTAGAAGTCCTAAGTTGAATCTACAGTACCAAGTAGTGTAACGATGACGGAACGATCGGTCATATAAAGTCACATAATTGGCGCTTTGCAATTGAGTGCCAAGCCTCGTTTAATCAAATATCGAGTGACATGAAACCACTAGGATCGAGGTACTAGATTAATGTCGAGCTCATTGACTCGTGCTGCAGACAGGAATGACTTTAAGATAGCAAT |
| **E48.BC79** Exc-LMO3.71 (spinal) | AGAACACGTGGATCCGTTAGAGGATGCCTCTTGGTTGCTTAAAAAAAGAGTCAGTCCATTTAAAATTGTTAGTTTTCTATAGTCTTTCAGAACATAGAATTCATTTAGAATTTCCAGCAAAAGTACAGAGAGTCTTCAGGGAAGTTGAAGAAGATAAAATAATTCTGAACACACTGTACACTTTCAGAAGAAAAATATTAAGAGTAAAAATAGTTTCCCAACTCAGTCATTCTTTGGAACTGATAATTTATTCCTGATGGTTCCCCAAAAGAGGGGAAGAGAAGCAAAATAATATATTTGGCTCCTCTAATAGATATTATTCATAGCTCTTTTCCCTCATAGGATAATAGACTATTAGAGCTTCTATAAAATCCCTTTCAGTAATTTTATGGTGGGTCTGCATTTTGTAAGCCATGTTTTATAACTGAGAAAAGAAAAAAAAAAAAACACAACCACCAAGAGAATAATTTTTACAATAATTCATTCCACTGATTTTATTAT | | CCTATTGTGGGGACTCCCACATGTTCAAATTACACTCGTTACGGTACTGGGCCACTTGCCCGCTTCCATTAATGGAGTCGTCTGATTACGCCCTGTGTTGAAAGTAGTATCTAGCGCGATGCCTAACCGTAAGGGCCCCGGATTCTTTATGTCAATATCACCCCGCGGCACACGTTGGCCAAGGTAAGCCTGCAGTCATTCTGATCGAAGACCAACTGGTGTTTAGATGGCCTTCACCCTGAACGTGGTTGTCTTCATGATGGCTGGGCCAGAAGGCTGCTCTCTCGAAGGGAGGAAGTGGGGTATTCTGTATAGAAATCGACGCTCGAAGGGCAGTACCACTCGTCCAATTCACTCGGGGTCCCACGATTCAAGGGGAAGAGATTGGCAGTAGGTCCCTGTCAATGTAACGTGGTGGTACCGACATACGTGCGTCGAAATTGTGTCGACTCTTTCCAGGCATGGTCTGGATGATCCAACGCCTAAGAGCACACACAACT |
| **E49.BC80** CS-neg-1: negative control | CGGTCGGCCGGTATGTTGGACCGTCTTTATGTGTGAACAGCATCCGCAAATCTGACTACAGCGAGAGTACGGGGCATTGGTCCGGGAACGGTCGTCAATACTTCTAGCTAACGAGCACTCTTGTTCTCACTAATATGTGCAGCACGAAACATATGTCCAAAAGAACCAAACAACGTGTTCCCAATATTGTATACTCATAGGGGCGCAGAAAGGCCATAATCGATCCTACCTAGCGGTTCTACGTCAGTGCGATCCCCATTATGCCCGACCGTCGCGTAGTATGACGGCACGCTCTGGAGCAAGTGACTTGGGAGAGCACTGCGGACGAGCTACTCTGGTGCCGTCCCGCTAGCAAGTGCGTGACGGCAGTCTTAGCAGTACTTGCTGGCATAGCTTAAAGCTCGCGCTCGACGAGACAGGGCGCTACGCTCGTGCTCACACCGCCACAATTCCTCCGGCAAGGCGTGGTGGAATTGCCAAGCGAGCTTTGGGCAGGCCAA | | TCACACCTTGCCCGAGGCAGTGTTGCGACTTAACGCGTCAATAGGGACACGTCGTATTCCGCCAGACTACCCGTGGAAGGCTATAGGGCGCAAGAAGAAGCTAACACAAACGACCTCTAGTTTCTAAGTCTTGTAGTACAAAATTTGTCCTTGCGCCATGCGAAGAACGGAAAGCTCGCCGGGGCTCGGAGAGCTGACGTGCCACGTTACGAGCCCGGGCGCTCTGTACCTTGGCTTATCGAACAGGTTAGGAAATACGTCACGATTCGTCGTATTTAACTTAACGCAACATCACGGGAATTAGCTGTATTGACGCTAAAATGAAAAATAAAGCTAGCCGGTAGGTCGGTACAGACTGCTCGCTGGAGCCTCCCGCCACGAGTATTGAGCAGCAGAGGGCAGTTGTATTCGCGGACGGCCACGTTAGACCCAGTTTTATGATCACGGGCCCATTCCGATTGGAAGATTTGCTCACCATGCAACATAGGATGATTAGAATG |

| ID | Enhancer | Barcode |
|---|---|---|
| **E50.BC81**<br>CS-neg-2:<br>negative control | GGGTTGCGTAGTCGGAGCAACGATTTGCTTCGGAAAATCGAAGACCGCCTTTGCGCGGACGAAGGAAACTTTATGGCCTCAGTCCGTTCTGGTTAGAGCTTCGTCTGCTTAAACATAGTGACGTCAACCCACTACTGGATAAACGTATACATGTGTCTCGGCCTTGTAATTGGATCGGAGGAACAGCTTTTACTCCCAAAGAAAGATCGGCTCTTACCGATCCTCGGCTGGCGGAAAGGGGCTACGGATGACAACGTGGGCAAGCTCCCCCTTGCTAACCTAGGTGACCGAAGGCGCCGTAACACAGTCACGTATCTGTTGCGCCTGAGTGTGTCGTTCATTACTTTCTAGCTCCAACGATCCTGTGGGAACATCCGAGGAACAAGTTGGTCCTCTACAATAAGAACAGAATGGCCAAGTGGGCGCAGATAGCAAACTCCGTTTGGCAGACGCATCCCTTACCCACCATTGGTAGAGGCCCGGAAGGGGCTATTCCGTAT | ACCAGACGGGAACATATTGCGCGGAGATACTTATGGCGGATTCACGGTAGACTTAGGAGGTCTTCCACCTTCTCGGTTAACAAACACCTCGTTTAAAATCCGTTCATGACTTTACGCGTGGACAACATAGACGTGGCCTAATAGGGCAGCCCGCAATAGTACTTGGACTTAAAGCAGCCTGGAAGCTGTAAGGCCTGACGAGACTTCCATTTCCAGGAGGCGGTTACGTTTTTAGAATAGAAAATGGAGCTTGCGAATACCCTCAGTTTCGAGGGATTTGGGGTTGCGCTAAACTGCCAGTGCCCTAATGCACGCGATATCGTTAATCATCCGTGGCCAGGGGCGACAAGAGCAAATACCTGTGAGTATGTTAGCTCCGTCATCGGGGAGGAGATTGGGTGAGGTAATCTTGCGAGGCATTGGTCGCGTGGAGCCCTGGATTTGTCTATGGCGGGTTATACTTAACAGCAGTTTTACTTGAATCGACCAGGGCTTGCGCG |

Table S12. **Library Enhancer and Barcode sequences that are used in this study.**

| Name | Sequence | Description |
|---|---|---|
| 280-NGS_BC_Fw_3 | GTCTCGTGGGCTCGGAGATGTGTATAAGAGACAGCGACTAGCGGATCCAAAAA | Binds at the beginning of the barcode region to prepare a sequencing library in the final vector (pAAV_B). Should be paired with 281-NGS_BC_Rv_4 |
| 281-NGS_BC_Rv_4 | TCGTCGGCAGCGTCAGATGTGTATAAGAGACAGAGCTTGATATCGAATTCCGC | ESCARGOT sequencing primer for the barcode of the step 2 plasmid to be paired with #280 or #282 (to be followed with v2Ad1.x and v2Ad2.x) |

Table S13. **Primer sequences that are used in this study.**

**RESSCU_data_combined.xlsx**

Table S14-S17**. RESSCU model coefficients, performance statistics.** S14 sheet 1: cell-type coefficients; S15 sheet 2: neuron subtype coefficients; S16 sheet 3: cell-type model fit statistics; S17 sheet 4: neuron model fit statistics
